# Dual regulatory role of IS*91*-encoded Orf121 in IS*91* transposition

**DOI:** 10.1101/2025.01.24.634351

**Authors:** Aurélien Fauconnier, Sandra Da Re, Margaux Gaschet, Thomas Jové, Marie-Cécile Ploy, Cécile Pasternak

## Abstract

Prokaryotic insertion sequences (IS) are pivotal in the propagation of bacterial multidrug resistance, with IS*91* notably linked to virulence and antibiotic resistance genes. However, the precise mechanism by which IS*91* contributes to gene dissemination remains elusive. Unique among its family, IS*91* features a small open reading frame (orf) upstream of the *tnpA* transposase gene, potentially encoding a 121-amino acid protein, *orf121*, which may be translationally coupled with *tnpA*.

Using a genetic system based on the mating-out assay in *Escherichia coli*, we explored the role of *orf121* in the *in vivo* transposition of IS*91*. Our findings indicate that the overlap between *orf121* and *tnpA* is crucial for *tnpA* transcription, with both being primarily transcribed as bicistronic mRNAs from the P*_orf121_* promoter. Additionally, the expression of *orf121* (whether *in cis* or *trans*) significantly reduces the frequency of IS*91* transposition and the rate of one-ended transposition. Furthermore, only the single-stranded DNA circles of IS*91* intermediates can integrate into new target sequences, and Orf121 negatively influences this insertion step.

In summary, *orf121* acts as a negative regulator of transposition while ensuring the expression of *tnpA*, thus providing insights into the complex mechanisms underlying IS*91*-mediated gene dissemination and its potential role in antibiotic resistance propagation.

## Introduction

Mobile genetic elements (MGEs) are pivotal in the propagation of multidrug-resistant genes, presenting a formidable threat to global health across human, animal, and environmental domains (for recent reviews, see ^1,2^). These elements encompass genetic entities that facilitate intercellular transfer (e.g., transmissible plasmids and integrative conjugative elements) as well as transposable elements (TE) that drive intracellular mobility, including insertion sequences (IS), transposons, and gene cassettes from integrons. TEs not only enhance the dissemination but also activate proximal antibiotic resistance genes (ARGs).

Among these, the IS*91* family is particularly atypical, with only four members confirmed to be actively transposing: the canonical IS*91* ^3^, IS*801* ^4^, IS*1294* ^5^, and IS*1294b* ^6^, closely related to IS*1294*). They encode transposases that belong to the HUH superfamily of single-strand nucleases^7^, which are presumed to transpose via a rolling-circle (RC) mechanism^8^, yet their exact role in gene dissemination remains elusive.

Unlike traditional IS that feature terminal inverted repeats (IR) flanking the transposase gene *tnpA*, IS*91* family members possess two functionally distinct ends^9^: the *ori*IS sequence, initiating transposition, and the *ter*IS end, terminating transposition. However, in IS*91*^9^ and IS*1294*^5^, termination often fails at a frequency ranging from 1 to 10% for IS*91* and from 1.7% to 14% for IS*1294b*^6^, leading to the mobilization of adjacent DNA fragments, a process known as one-ended transposition (OET). Consequently, as IS*91*-like elements are often linked with virulence and antibiotic resistance genes, they could significantly influence the spread of such genes^9–11^.

IS*91* family members do not generate target site duplications (TSD) upon insertion but display target sequence specificity. The recognized target for the insert is a tetranucleotide which is closely related to the elements active in transposition, IS*91* (5’-CTTG)^12^, IS*801* (5’- GTTC)^4^, IS*1294* (5’-GTTC)^5^ and IS*1294b* (5’-GTCC)^6^. The *ori*IS region of IS*801* and IS*91* contains two imperfect subterminal palindromes, whereas IS*1294* and IS*1294b* feature a single presumed palindrome (Fig. 1). The *ter*IS region contains a single imperfect subterminal palindrome adjacent to the cleavage site, differing between IS*91* (5’-CTCG) and the other three elements (5’-GTTC).

**Figure 1.**
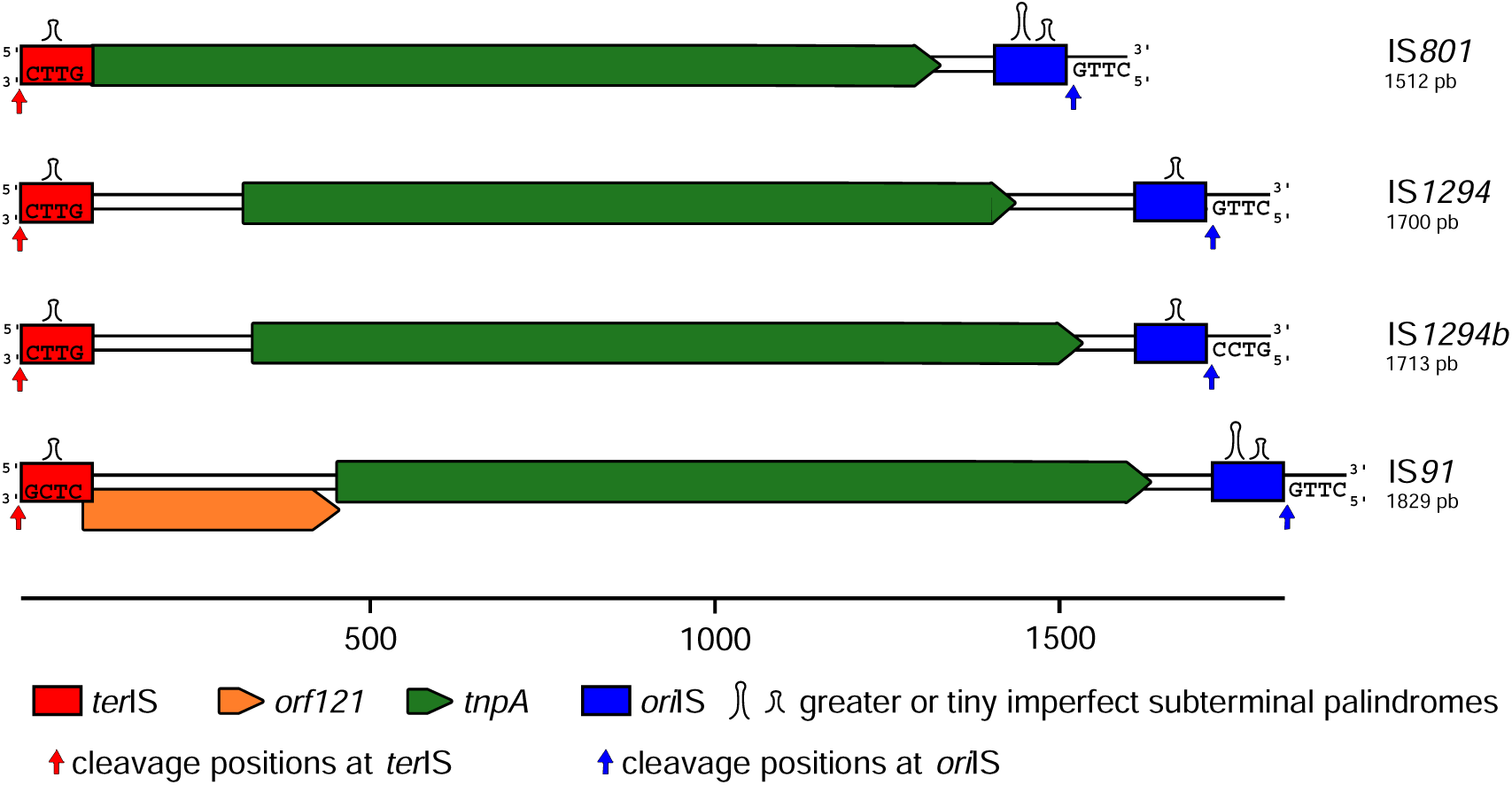
Genetic organization of members of the IS*91* family (IS*801*, IS*1294,* IS*1294b* and IS*91*) active for transposition. *ter*IS and *ori*IS ends carrying the palindromic sequences are depicted as red and blue boxes, cleavage positions at *ter*IS and *ori*IS ends are shown by red and blue vertical arrows, respectively. The cleavage site at the *ter*IS end and the tetranucleotide target site adjacent to *ori*IS are indicated. *orf121* and *tnpA* encoding sequences are shown as orange and green arrows, respectively.

Previous investigations have documented the *in vivo* formation of both single-stranded (ss) and double-stranded (ds) IS*91* circular intermediates during transposition, with both forms bearing a junction where *ori*IS*91* and *ter*IS*91* are juxtaposed^13^. However, it remains to be determined which intermediates play a functional role in the IS*91* transposition pathway.

Distinct from other IS*91* family members, the IS*91* features an additional open reading frame called *orf121* upstream of the *tnpA* transposase gene, which encodes a 121 amino acid polypeptide. The termination codon of *orf121* overlaps with the initiation codon of *tnpA*, implying a potential coupling in their expression. Prior research indicated that mutations in *orf121* do not impact transposition frequency but do alter the insertion target choice^14^. The specific role of the *orf*121 in IS*91* transposition has yet to be elucidated.

We aimed to determine the involvement of *orf121* in IS*91* transposition within *Escherichia coli*. Using a genetic system based on the mating-out assay, we observed that *orf121* expression induction reduces IS*91* transposition *in vivo*. Our findings indicate that *orf121* and *tnpA* are co-expressed, with their overlapping regions being critical for *tnpA* transcription. Furthermore, we established that only single-stranded DNA circular intermediates are inserted into new target sequences, and Orf121 negatively influences this step. Additionally, *orf121* expression significantly reduces the OET rate, suggesting that Orf121 is necessary for the precise recognition and cleavage of the *ter*IS*91* end.

## Results

### *orf121* is a conserved feature in the IS*91* element

To study if *orf121* plays a specific role in IS*91* transposition, we first wanted to address if *orf121* was conserved in different isoforms of IS*91*, the paradigm of the IS*91* family. As there was no available database analysis to assess the diversity and characteristics of IS*91*, we performed an *in silico* analysis using the amino acid sequence of the TnpA transposase encoded by IS*91* (accession number CAD30457.1) as a reference for the first described isoform. Sequences were filtered to retain those with an amino acid identity of 95% or higher and not truncated, identifying 134 distinct isoforms. Most of the isoforms were mainly identified in Enterobacterales, and as single copies on plasmids, with over half of the identified plasmids belonging to the IncFII type.

For all isoform sequences described and analyzed in this study, refer to Supplementary File 1 and File 2. The overlap between the *orf121* and *tnpA* genes was highly conserved, in 79% of cases. In other isoforms, the overlap loss was due to insertions or deletions within the homo- polymeric regions of the *orf121* sequence. The target insertion sites showed a consistent preference for 5’-GTTC or 5’-CTTG, followed by 5’-CTCG, with a T consistently in the second position. The predominant cleavage site tetranucleotide was 5’-CTCG. This result is in aligning with previous reports^9,12^.

### P*_orf121_* drives expression of both *orf121* and *tnpA* genes

As mentioned above, the termination codon of *orf121* overlaps with the initiation codon of the *tnpA* gene in IS*91,* suggesting that the two genes could be expressed from the same promoter. For this reason, we have analyzed the transcriptional expression level of *orf121* and *tnpA* in their natural configuration within the IS*91* element.

Using BPROM (http://www.softberry.com)^15^, we identified putative σ70 promoters and Shine-Dalgarno (SD) sequences for both genes. Notably, the putative *tnpA* promoter P*_tnpA_* was located within the coding sequence of *orf121* and P*_orf121_*in the *ter*IS*91* region of the IS*91* element (Fig. 2a). The activity of the predicted promoters P*_tnpA_* and P*_orf121_*was assessed by generating transcriptional fusions between each putative promoter and the *lacZ* gene, including the *lacZ* ribosome binding site (RBS) (Fig. 2b). β-galactosidase assays revealed that both promoters are functional, with P*_orf121_* exhibiting significantly higher activity than P*_tnpA_* (62.8-fold; Fig. 2c).

**Figure 2.**
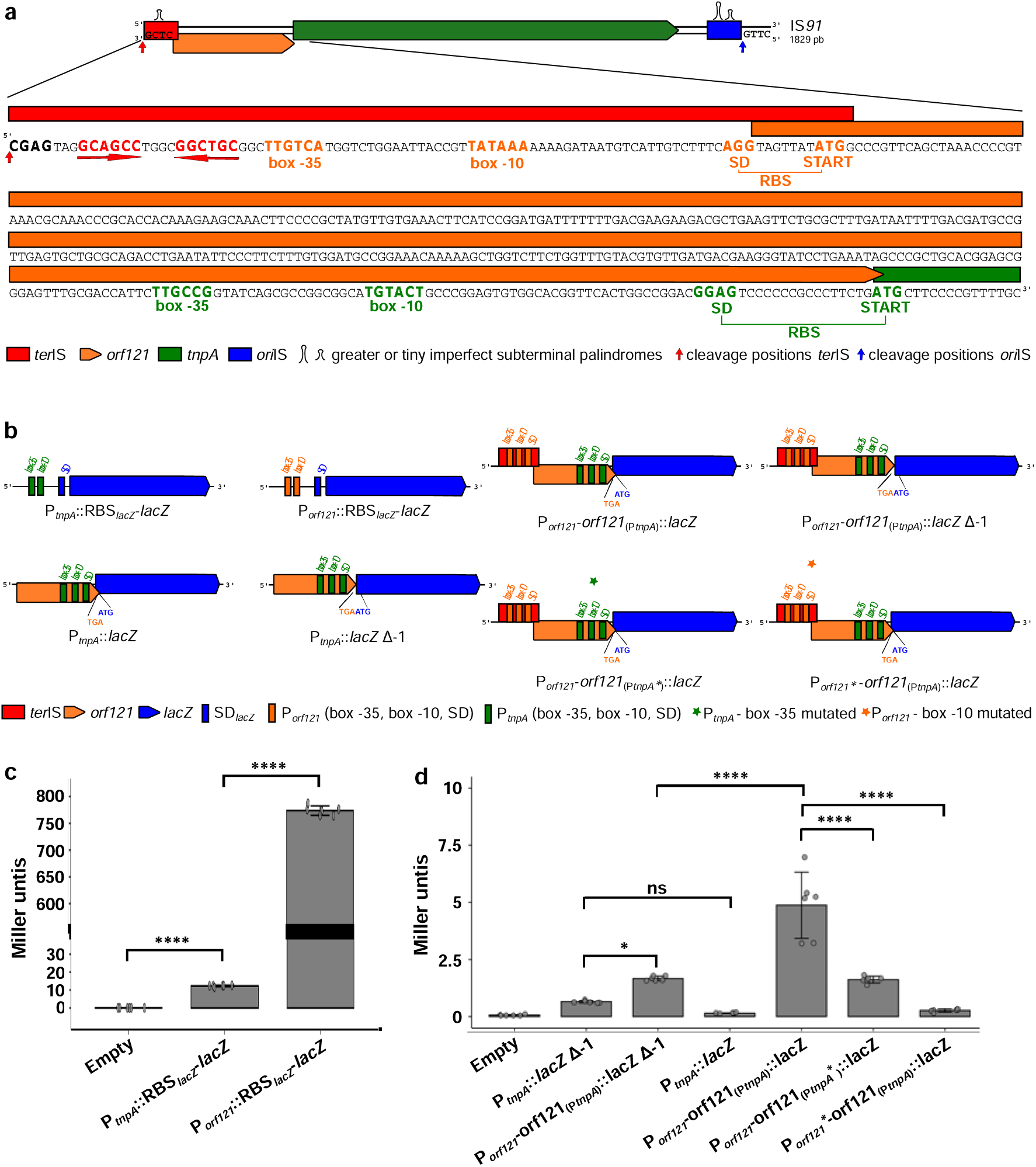
Sequence of the IS*91* from *ter*IS*91* to the start of the *tnpA* gene and estimation of the P*_orf121_* and P*_tnpA_* promoters activity. a) Sequence of IS*91* accession number X17114.5. The *ter*IS region, the *orf121* and *tnpA* genes are indicated by a red rectangle, an orange arrow and a green arrow above the nucleotides, respectively. Red arrows represent the *ter*IS terminal palindrome. The cleavage tetranucleotide is in bold with a red arrow indicating the cleavage site. The potential -35, -10, SD and start codon regions are shown in bold (orange for P*_orf121_* and green for P*_tnpA_*). b) Scheme of the P*_orf121_* and P*_tnpA_ lacZ*-transcriptional fusion constructs described in Table 1. c) Β-galactosidase activities of the transcriptional fusions between *lacZ* and the two predicted promoters: P*_orf121_* and P*_tnpA_* with lacZ RBS. d) Β-galactosidase activities of the transcriptional fusions between *lacZ* and P*_tnpA_*with its native RBS in the presence or absence of the +1 overlap and between *lacZ* and P*_orf121_* with its native RBS and *orf121* gene in the presence or absence of the +1 overlap. Beta-galactosidase activity is expressed as Miller units. Empty: empty transposase expression vector. The results are the average of at least 3 independent experiments. Ns = non-significant; p = 0,0332 (*) and p <0.0001 (****).

**Table 1.**
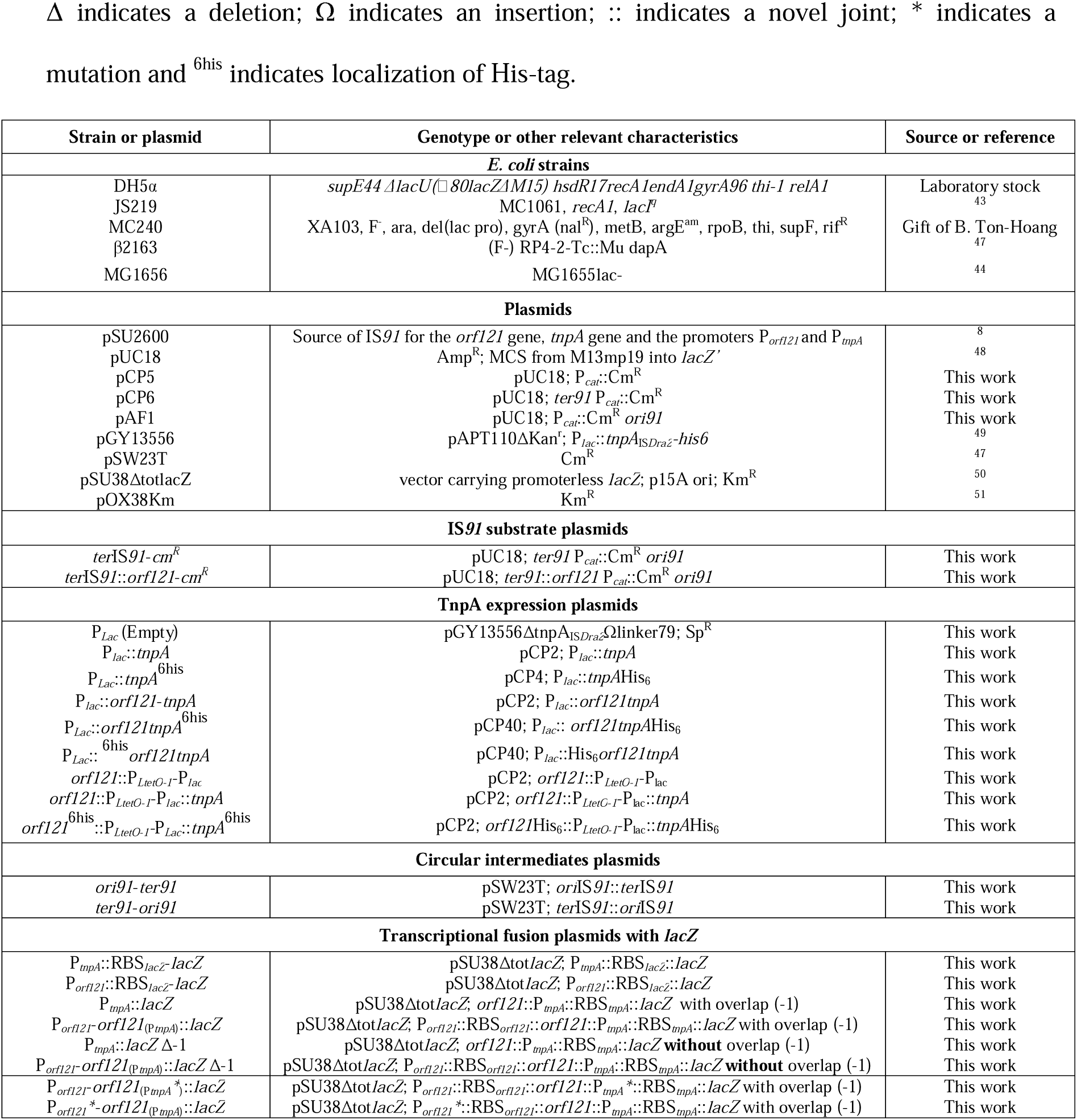
Bacterial strains and plasmids. Δ indicates a deletion; Ω indicates an insertion; :: indicates a novel joint; * indicates a mutation and ^6his^ indicates localization of His-tag.

To investigate the expression of *orf121* and *tnpA* in their native configuration to determine whether *tnpA* expression is coupled to *orf121* and to assess the significance of the overlap between the *orf121* stop codon and the *tnpA* start codon. We created two new *lacZ* transcriptional fusions with P*_tnpA_*, including its native RBS, either retaining or removing the +1 overlap (Table 1, Fig. 2b). Additionally, we developed two translational fusions between *lacZ* and the *orf121* gene with its native promoter, P*_orf121_*, resulting in P*_orf121_*- *orf121*(P*_tnpA_*)::*lacZ*, with and without the +1 overlap (Table 1, Fig. 2b). Globally, the estimated β-galactosidase activities were much lower in constructs compared to the ones with the lacZ RBS. The activity of P*_tnpA_* was completely abolished when maintaining its native RBS compared to the construct with the *lacZ* RBS (compare Fig. 2c and Fig. 2d). Although still very low, the activity of P*_tnpA_* increased significantly, by a factor of 4.1, when the native +1 overlap was removed. The β-galactosidase activity of P*_orf121_*-*orf121*(P*_tnpA_*)::*lacZ* was low, indicating that P*_orf121_* has a weak activity. In contrast to P*_tnpA_*, the removal of the *orf121*/*tnpA* overlap resulted in a 2.9-fold decrease in β-galactosidase activity. Although P*_tnpA_* activity was minimal (<1 MU), it increased when the P*_orf121_*promoter was present upstream (P*_orf121_*- *orf121*(P*_tnpA_*)) in both configurations, with and without the *orf121*-*tnpA* overlap (Fig. 2d). In the absence of the overlap, the activity of P*_orf121_*-*orf121*(P*_tnpA_*)Δ-1 was reduced by 2.91-fold compared to the construct with the overlap. To further investigate the contributions of the P*_orf121_* and P*_tnpA_* promoters to *tnpA* expression, we engineered mutants of these promoters within the P*_orf121_*-*orf121*(P*_tnpA_*)::*lacZ* construct. Specifically, we mutated P*_orf121_* in the -10 box (TATAAA to CGCGAA) and P*_tnpA_* in the -35 box (TTGCCG to TGGCGG) without altering the amino acid sequence of Orf121. The activities of the mutated promoters showed significant reductions, experiencing a 18-fold and a a 3-fold decrease respectively, compared to P*_orf121_*P*_tnpA_*(Fig. 2d).

Altogether, these results suggest that *tnpA* is predominantly expressed from the P*_orf121_*promoter and that its expression is heavily dependent on the overlap between *orf121* and *tnpA*.

### Inhibitory role of *orf121* in IS*91* transposition

After assessing the expression of *orf121* and *tnpA* genes within the IS*91* element, we explored the potential regulatory role of *orf121* in IS*91* transposition *in vivo*. For this purpose, we utilized a genetic system using a mating-out assay in *E. coli* containing three plasmids to assess the mobility of IS*91* derivatives into the pOX38Km conjugative plasmid, upon expression of *tnpA* alone or with *orf121 in trans* (relative to IS*91*) (see Fig. 3, and Materials and Methods). Two different IS*91* derivatives were cloned into a high-copy number plasmid. Both derivatives include a sequence of 89 bp from the left-end of IS*91* at the *ter*IS*91* cleavage site (5’-CTCG) and the final 82 bp of the right-end, shown to be sufficient for functional transposition^8^, encompassing the *ori*IS*91* as well as the essential 5’-CTTG tetranucleotide adjacent to the *ori*IS*91* end (Fig. 3a). These *ter*IS*91* and *ori*IS*91* sequences flanked a Cm^R^ cassette (for selection of transposition events) in the IS*91* derivative termed *ter*IS*91*-*cm^R^*. Another derivative, *ter*IS*91*::*orf121*-*cm^R^*, included the Cm^R^ cassette along with *orf121* expressed *in cis* relative to IS*91* (Table 1 and Fig. 3a). TnpA was expressed *in trans* from a compatible plasmid under the inducible P*_lac_* promoter, either alone called P*_lac_*::*tnpA* or with *orf121* termed P*_lac_*::*orf121*-*tnpA* (in its native configuration with overlapping *orf121*- stop/*tnpA*-start) (Table 1, Fig. 3a). Transposition frequencies of IS*91* derivatives into pOX38Km were evaluated as illustrated in Fig. 3b. With *orf121* expressed *in trans*, we observed a dramatic reduction in transposition frequency (8045-fold) with *ter*IS*91*-*cm^R^* upon IPTG induction when both *orf121* and *tnpA* were expressed (P*_lac_*::*orf121*-*tnpA*), compared to expression of tnpA alone (Fig. 3c). In contrast, under basal conditions of *tnpA* expression (i.e., without IPTG induction) from P*_lac_*::*tnpA*, expression *in cis* of *orf121* from its native promoter (*ter*IS*91*::*orf121*-*cm^R^*) resulted in a significant 11.1-fold decrease in transposition frequency compared to *ter*IS*91*-*cm^R^*. However, in the presence of IPTG, no significant impact on transposition frequency was observed when *orf121* was expressed *in cis* from its native promoter (*ter*IS*91*::*orf121*-*cm^R^*) compared to the construct lacking *orf121* (*ter*IS*91*-*cm^R^*) in cells expressing only *tnpA* (P*_lac_*::*tnpA*).

**Figure 3.**
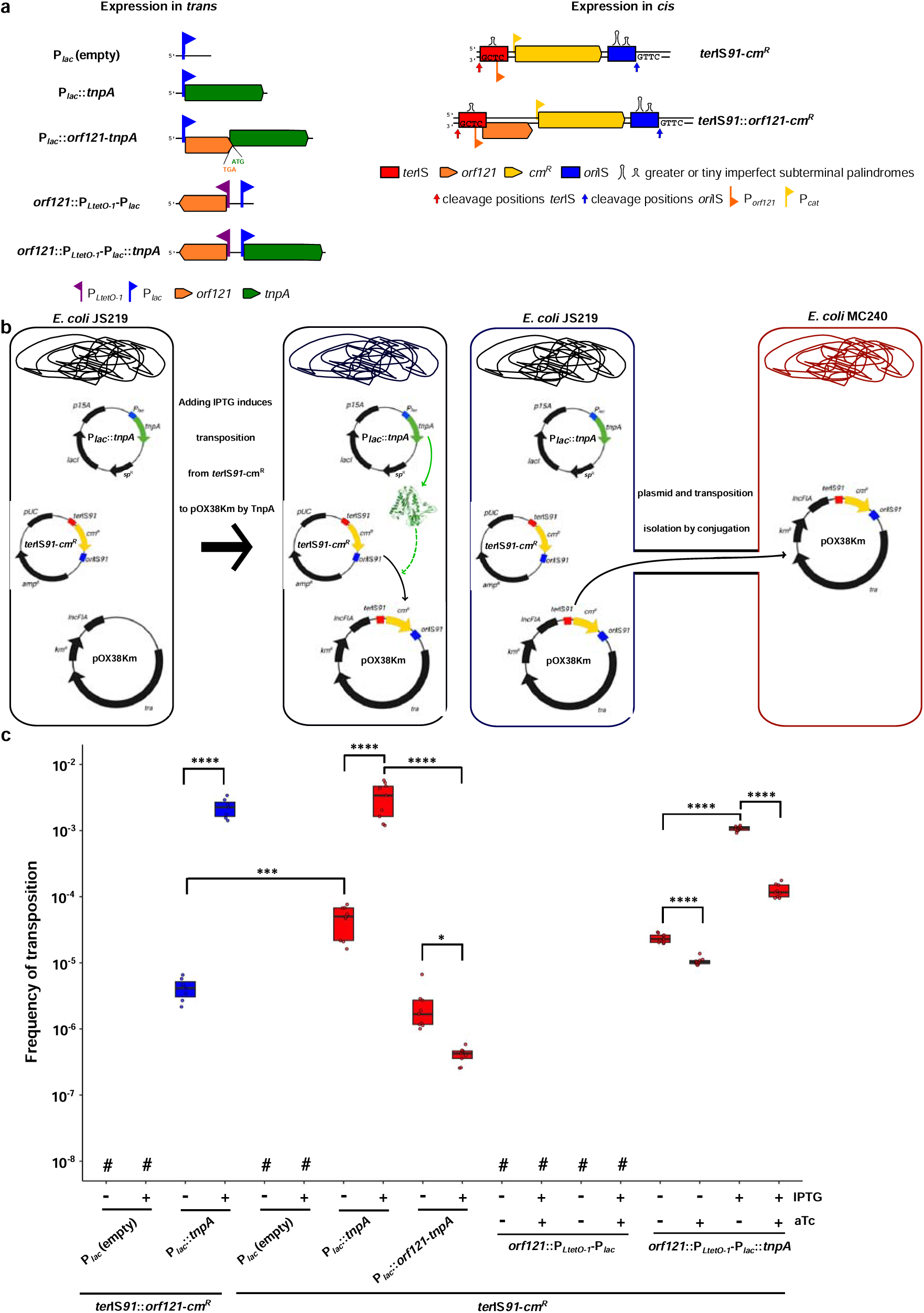
Expression of *orf121* decreases IS*91* transposition frequency. (a) The IS*91* transposase donor plasmids (*expression in trans*) feature the P*_lac_* promoter (blue), which controls the *tnpA* gene (green) with the upstream presence of *orf121* (orange). When the two genes are expressed under the control of two promoters, the *orf121* gene is under the control of the P*_LtetO-1_* promoter (purple). These plasmids are selected by the *sp*^R^ resistance gene. IS*91* substrate plasmids are derivatives of the pUC18 and confer ampicillin resistance (*amp*^R^). They carry the mini-IS composed of the *ori*IS*91* (blue) and *ter*IS*91* (red) flanking the *cm*^R^ mobility reporter gene (Chloramphenicol resistance; yellow) for *ter*IS*91*-*cm^R^*and, the *ter*IS*91*::*orf121*-*cm^R^* composed of the *ori*IS*91* (blue) and *ter*IS*91*::*orf121* region (native IS*91* configuration, red and orange) flanking the *cm*^R^ mobility reporter genes, used for detecting the OET mechanism. (b) Schematic of the mating-out assays, the donor *E. coli* JS219 (black) carries three plasmids: the IS donor plasmid, a derivative of pUC18 carrying *ter*IS*91*-*cm^R^*; a second plasmid with an origin of replication p15A and expresses the IS*91* transposase (*tnpA*) under the control of the IPTG-inducible P*_Lac_* promoter, called P*_lac_*::*tnpA*. The conjugative pOX38Km (a derivative of plasmid F) is the target for transposition. We use conjugation to isolate transposition events on the pOX38Km plasmid in the *E. coli* MC240 recipient cell. (c) Transposition frequency of IS*91* derivatives (*ter*IS*91*-*cm^R^* and *ter*IS*91*::*orf121*-*cm^R^*) estimated by mating-out assays (see (b)) when *tnpA* is expressed *in trans* alone or with *orf121* from the same or independent promoters (box red) or with *orf121* expressed *in cis* from *ter*IS*91*::*orf121*-*cm^R^* (box blue). The – and + indicate the presence or absence of inducers, IPTG (0.5 mM) and aTc (50 ng/ml). Data are represented as box-plot. Experiments were performed at least 5 time. P = 0,0332 (*); p = 0,0002 (***) and p <0.0001 (****). #, undetectable transposition (< 3.35 x 10^−8^).

Then, we explored whether separate expression of *orf121* and *tnpA* genes could impact the mobility of the IS*91* derivative, hypothesizing that Orf121 might modulate transposition frequency. We assessed the transposition frequency of the *ter*IS*91*-*cm^R^*derivative in *E. coli*, where *tnpA* and/or *orf121* were expressed independently from the inducible promoters P*_lac_*and P*_LtetO-1_*, respectively, designated as *orf121*::P*_LtetO-1_*-P*_lac_*::*tnpA* (Fig. 3c). In the absence of IPTG and anhydrotetracycline (aTc) inducers, the basal transposition frequency of *ter*IS*91*- *cm^R^*was observed in cells containing *orf121*::P*_LtetO-1_*-P*_lac_*::*tnpA*, due to promoter leakage. With aTc leading to the expression of *orf121* from P*_LtetO-1_* we observed a significantly decrease of the transposition frequency by 2.3-fold compared to the condition without aTc. When *tnpA* expression was induced with IPTG alone, transposition frequency increased by 45.1-fold relative to the basal level. However, co-expression of *orf121* with *tnpA* in presence of both inducers aTc and IPTG, led to an 8.6-fold reduction in transposition frequency relative to IPTG alone. This result reveals that, aside from the inherent configuration of the *orf121* and *tnpA* genes impacting element mobility, the Orf121 protein modulates transposition frequency, independently of the *orf121*-*tnpA* overlap.

### Impact of *orf121* on *tnpA* transcription and translation

In conjunction with the mating-out assay, we quantified the transcripts of *orf121* and *tnpA*. Our findings revealed that co-expressing *orf121 in trans* with *tnpA*, whether from the same or independent promoters with *ter*IS*91*-*cm^R^*, resulted in a significant reduction in the mRNA copy number of *tnpA* by 6.9-fold and 1.8-fold, respectively (Fig. 4a). Similar trends were observed when *orf121* was co-expressed *in cis* (*ter*IS*91*::*orf121*-*cm^R^*) with *tnpA* (P*_lac_*::*tnpA*), with a 2-fold decrease in *tnpA* transcript levels compared to *ter*IS*91*-*cm^R^*. The decline in *tnpA* transcripts was inversely proportional to the number of *orf121* transcripts. There is no impact of *tnpA* expression on the mRNA copy number of *orf121* (Fig. 4b). These observations suggest that *orf121* exerts a negative regulatory effect on *tnpA* transcript levels.

**Figure 4.**
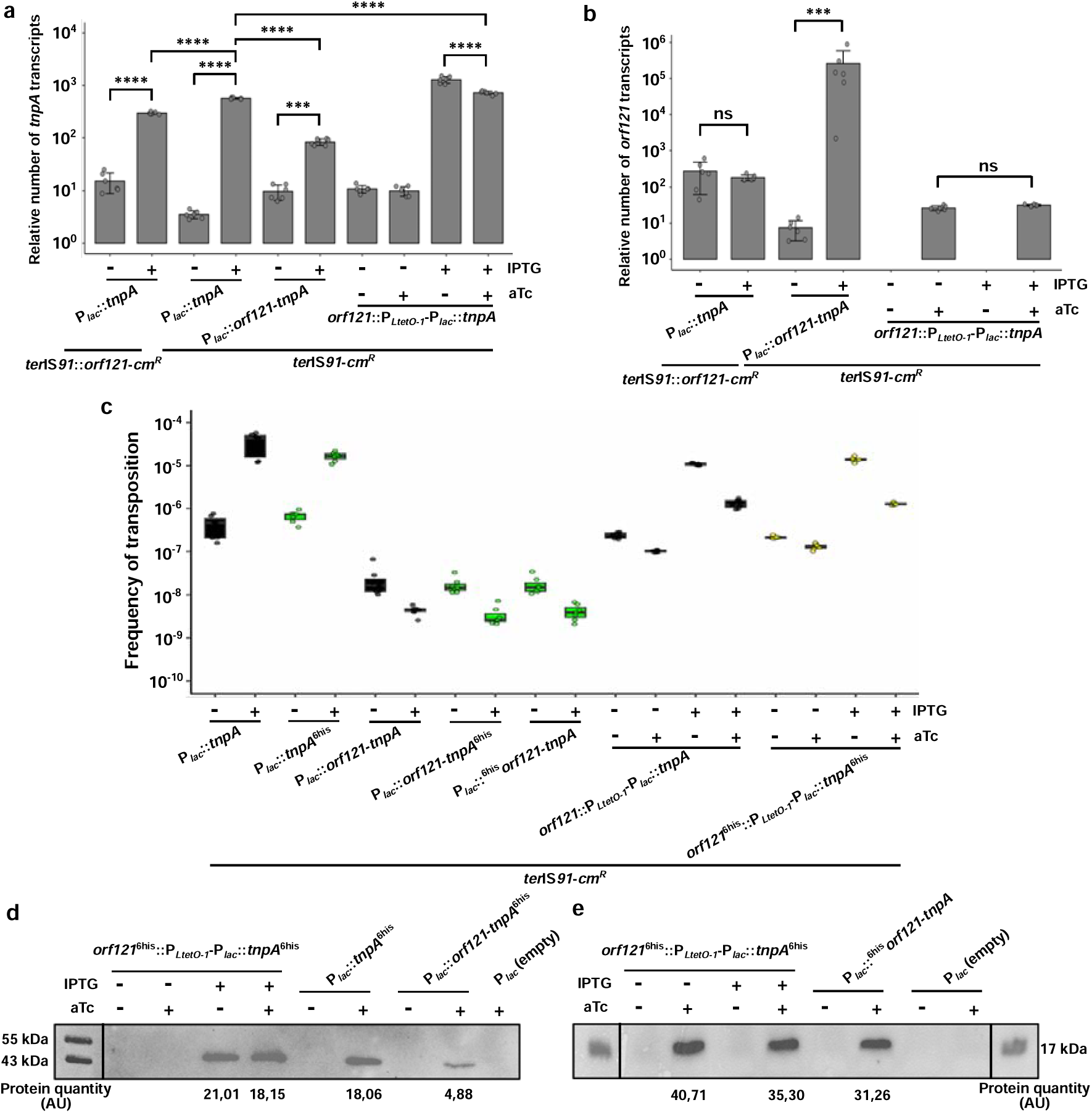
*orf121*-driven changes in *tnpA* transcription and translation dynamics. (a and b) Transcript levels of *tnpA* (a) and *orf121* (b) in mating-out experiments. Transcript levels of *tnpA* and *orf121* were normalized to the housekeeping gene *dxs*. Data are the average of transcripts levels measured from at least 5 independent mating-out experiments. Error bars indicate the SD. Asterisks indicate significant difference: ns = non-significant; p = 0,0002 (***) and p <0.0001 (****). Symbols and constructs are detailed above (Fig 4a). (c) Transposition frequency of the IS*91* derivative (*ter*IS*91*-*cm^R^*) estimated by mating assays when *tnpA* is expressed *in trans* alone or with *orf121* from the same (box green) or independent (box yellow) promoters, when *tnpA* and *orf121* are tagged with a His-tag (C-term for *tnpA*\* or N-term for \**orf121*) compared construction without tagged (box black). (d and e) Detection of TnpA (d) and Orf121 (e) protein by western immunoblotting. His- tagged TnpA (C-term; tnpA*; expected size 47 kDa) and His-tagged Orf121 (N-term; *orf121; expected size 17 kDa) were extracted from the mating-out experiment culture. The - and + indicate the presence or absence of inducers, IPTG (0.5 mM) and aTc (50 ng/ml). *, His-tag position; AU: Arbitrary Unit of quantity of protein. Symbols and constructs are detailed above.

In parallel, we assessed the protein levels of Orf121 and TnpA to complement the mating assay results that highlighted the impact of Orf121 on IS*91* mobility. Using C-terminal His-tagged TnpA (*tnpA*^6his^) and N-terminal His-tagged Orf121 (^6his^*orf121*) proteins in mating-out assays, we performed Western-immunoblotting (Table 1). It was confirmed that His-tags did not influence transposition frequencies (Fig. 4c). Orf121 was exclusively detected in the soluble fraction of the bacterial lysate, whereas TnpA was present in the pellet as inclusion bodies. The identities of ^6his^Orf121 and TnpA^6his^ were verified via mass spectrometry. As shown in Fig. 4d, the quantity of TnpA remained stable, around 20 AU (Arbitrary Units), regardless of whether *tnpA* was expressed alone or with *orf121* when their expression was independent (*orf121*^6his^::P*_LtetO-1_*-P*_lac_*::*tnpA*^6his^). However, when *tnpA* and *orf121* were expressed from the same promoter (P*_lac_*::*orf121*-*tnpA*^6his^), the quantity of TnpA was 4.1-fold lower. Orf121 production remained similar across different expression conditions (Fig. 4e). These findings indicate that the reduced production of the TnpA protein is primarily due to the coupled transcription of *orf121* and *tnpA*. This correlates with the decreased number of *tnpA* transcripts observed (P*_lac_*::*orf121*-*tnpA*).

Collectively, the transcript and His-tag analyses suggest that the regulation of coupled *orf121* and *tnpA* expression operates at both transcriptional and translational levels.

### Effect of Orf121 on integration of IS*91* circular intermediate forms in *E. coli*

It has been previously established that two distinct DNA species, single-stranded (ss) and double-stranded (ds) IS*91* circular intermediates, are independently formed *in vivo* post-induction of transposase expression, with *ori91* and *ter91* fusion^13^. Nevertheless, the functional intermediates facilitating IS*91* insertion have yet to be clarified. To address this, we employed a suicide mating assay *via* transformation or conjugation of the *ter91*-*ori91 or ori91*-*ter91* junction into *E. coli* DH5α strains harboring various *tnpA* and/or *orf121* expression plasmids (P*_lac_*::*tnpA* or P*_lac_*::*orf121*-*tnpA*). The sequence of the *ter91*-*ori91* junction and the ds or ss insertion assays are presented in Fig. 5a and Fig. 5b (Table 1).

**Figure 5.**
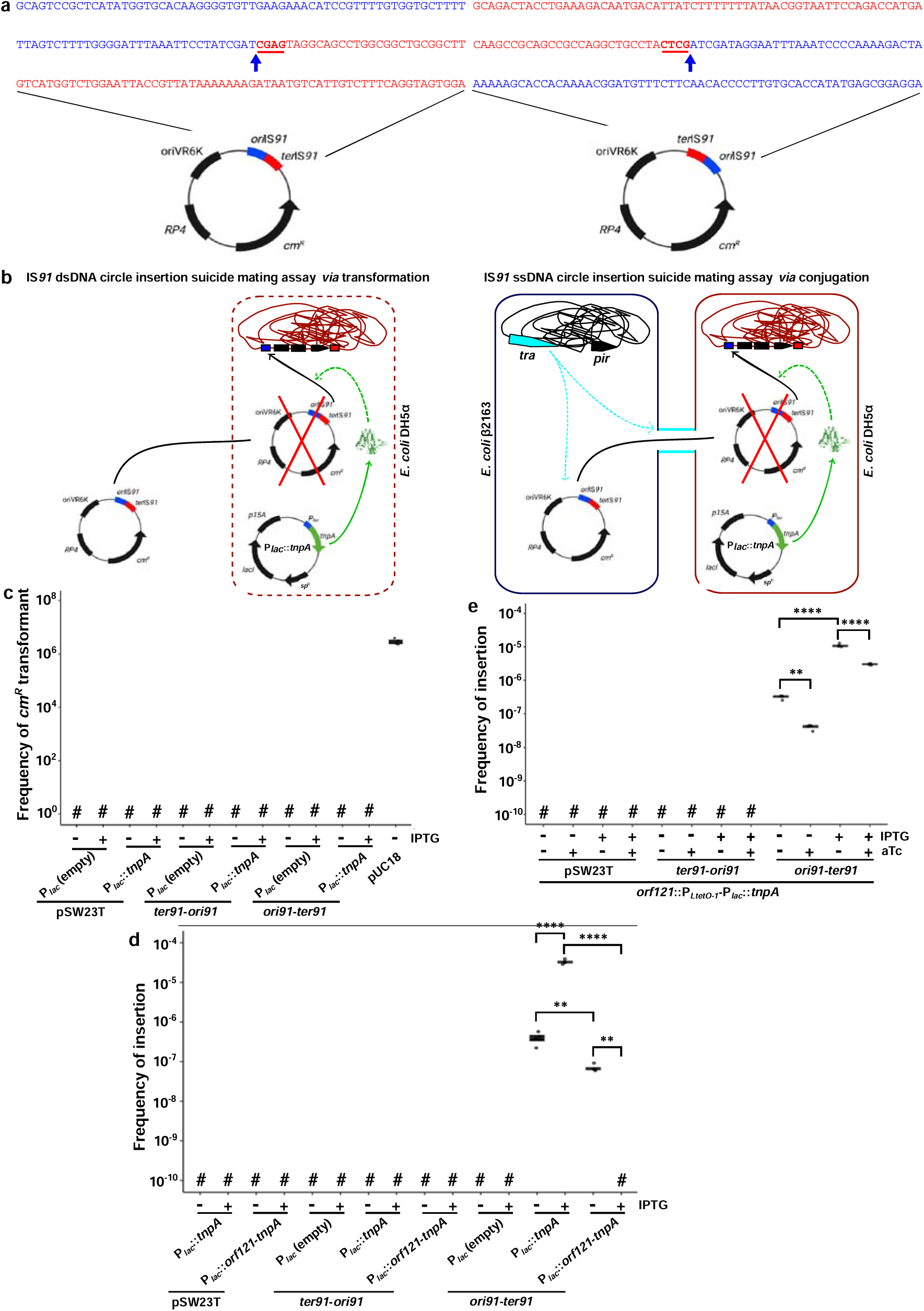
Only ssDNA circular intermediates are successfully inserted into a new target sequence. (a) Sequence of the *ori91*-*ter91* or *ter91*-*ori91* junction present in the substrates used for insertion analyses. *ter*IS and *ori*IS ends are depicted as red and blue, cleavage positions are shown by blue vertical arrows, respectively. The cleavage site at the *ter*IS is indicated (bold red underline). (b) Principe of suicide mating assay via transformation or conjugation of the *ter91*-*ori91* or *ori91*-*ter91* junction into *E. coli* DH5α strains to test ds- and ss-circular intermediates. (c) Transformation assays assessing *ter91*-*ori91* or *ori91*-*ter91* dsDNA-circular IS*91* intermediates insertion into *E. coli* chromosome when TnpA was expressed alone or with Orf121 from the same promoter. The pUC18 plasmid was used as a positive control of transformation efficiency. Experiments were performed 4 times. #, undetectable transformants (< 2.63 x 10^−9^). (d and e) Suicide conjugation assays assessing *ter91*-*ori91* or *ori91*-*ter91*ssDNA-circular IS*91* intermediates insertion into *E. coli* chromosome when TnpA was expressed alone or with Orf121 from the same promoter (d) or with Orf121 from independent promoters (e). Experiments were performed 4 times. p = 0,0021 (**) and p <0.0001 (****). #, undetectable exconjugants or transformants (< 2.93 x 10^−9^).

For dsDNA experiments, the positive control transformation with the pUC18 plasmid yielded the expected transformation frequency with 100 ng of pUC18 (Fig. 5c). However, no transformants were detected with any ds-circular *ter91*-*ori91 or ori91*-*ter91* junction intermediate, suggesting the dsDNA intermediate is not a viable substrate.

In contrast, for ssDNA experiments, the absence of Cm^R^ exconjugants was observed with donor cells carrying the empty plasmid pSW23T (lacking *ter91*-*ori91 or ori91*-*ter91* junction) and recipient cells expressing P*_lac_*::*tnpA* or P*_lac_*::*orf121*-*tnpA* as excepted (Fig. 5d). Likewise, the frequency of Cm^R^ exconjugants was undetectable when the donors possessed the DNA junction in the *ter91*-*ori91* orientation. Conversely, when the donor cells carried the *ori91*- *ter91* junction, we observed a significant reduction in transposition frequency in conditions without IPTG (promoter leakage) when both *orf121* and *tnpA* were expressed from the same promoter (P*_lac_*::*orf121*-*tnpA*) compared to *tnpA* alone (P*_lac_*::*tnpA*) (Fig. 5d). Upon IPTG-induced expression, *ori91*-*ter91* junction transposition frequency was detectable in cells expressing *tnpA* alone but became undetectable in cells expressing both *orf121* and *tnpA*, with a substantial 12347-fold reduction. Further experiments with recipient cells expressing *tnpA* and *orf121* under distinct inducible promoters (*orf121*::P*_LtetO-1_*-P*_lac_*::*tnpA*) verified the negative impact of Orf121 on *ori91*-*ter91* junction in insertion (Fig. 5e).

Molecular analysis via arbitrarily primed PCR (AP-PCR) of numerous exconjugant clones confirmed all Cm^R^ exconjugants originated from *ori91*-*ter91* junction insertion into the recipient cell’s chromosome (Supplementary Table 2).

Our results indicate that the single-stranded circular intermediate is the exclusive form capable of initiating insertion, and that only the bottom strand is engaged in the process. Additionally, Orf121 acts as an inhibitor, reducing the insertion efficiency of the circular intermediate mediated by TnpA.

### Lack of Orf121 influence on IS*91* target site insertion

Earlier research indicated that Orf121 might influence insertion specificity, as *orf121* mutations affected target selection without impacting transposition^14^. To confirm that the observed Cm^R^ exconjugants were due to transposition of the IS*91* derivative into the pOX38Km plasmid, rather than spontaneous mutations, AP-PCR analysis was conducted on numerous transposition events to identify target insertion sites.

Given our findings, we performed a χ² test to evaluate whether the presence or absence of *orf121*, as well as its *cis* and *trans* expression, affected the target site preferences at 5’-GTTC and 5’-CTTG. The data revealed that the target site specificity was consistent across conditions, with *orf121* expressed either *in cis* (*ter*IS*91*::*orf121*-*cm^R^*) or *trans* (P*_lac_*::*orf121*- *tnpA*) and was similar to that observed with *tnpA* alone (P*_lac_*::*tnpA*) (Table 2) (χ² value = 22.20; p-value = 0.45). Additionally, no significant differences in target specificity were detected when *orf121* and *tnpA* were regulated by distinct promoters (*orf121*::P*_LtetO-1_*- P*_lac_*::*tnpA*) (Table 2) (χ² value = 3.77; p-value = 0.29). These results suggest that the expression of Orf121, whether in *cis* or *trans*, does not alter the specificity of target selection or insertion sites.

**Table 2.**
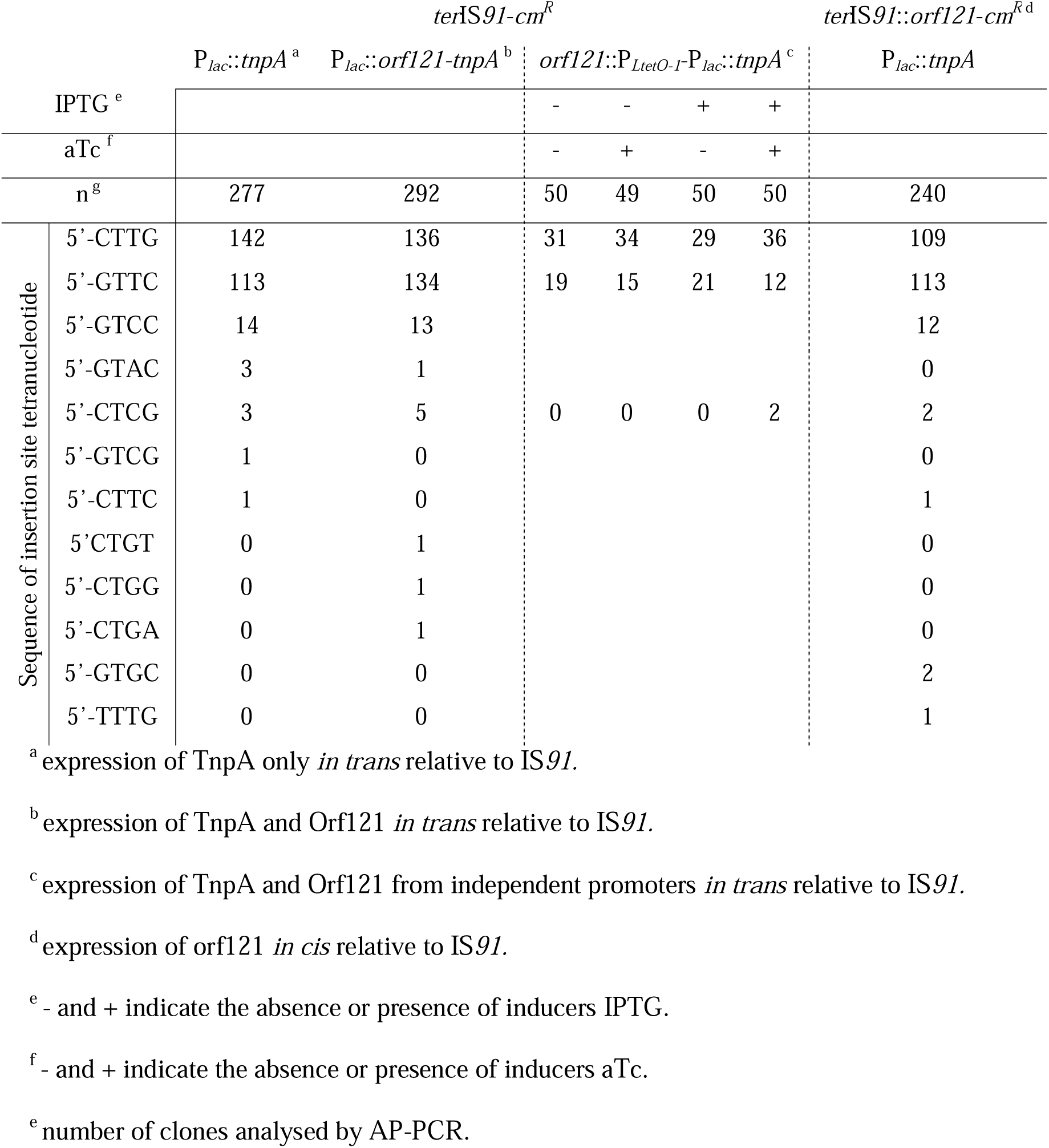
Identification of the tetranucleotide insertion target in the different mating-out assays.

### *orf121* is a key factor in precise *ter*IS*91* end recognition and cleavage

We explored the involvement of Orf121 in the specific recognition and cleavage of the *ter*IS site by analyzing the tetranucleotide cleavage patterns at the *ter*IS*91* end across multiple transposition events, utilizing AP-PCR. Our findings consistently identified a single cleavage site (5’-CTCG) in all instances, whether the transposase was expressed independently (P*_lac_*::*tnpA* with *ter*IS*91*-*cm^R^*; 43 events) or alongside *orf121 in trans* (P*_lac_*::*orf121*-*tnpA* with *ter*IS*91*-*cm^R^*; 68 events).

Subsequently, we investigated the proportion of one-ended transposition (OET) in different mating-out assays, given the known inefficiency of IS*91 ter*IS*91* end. As detailed in Table 3, expressing *tnpA* alone (P*_lac_*::*tnpA* or *orf121*::P*_LtetO-1_*-P*_lac_*::*tnpA* with IPTG only) led to approximately half of the events being OET. In contrast, the co-expression of *orf121 in trans* P*_lac_*::*orf121*-*tnpA* and *orf121*::P*_LtetO-1_*-P*_lac_*::*tnpA* with IPTG and aTc significantly lowered OET rates by 2.94- and 1.5-fold, respectively (relative to the conditions without *orf121*). Similarly, *orf121* expression *in cis* (P*_lac_*::*tnpA* with *ter*IS*91*::*orf121*-*cm^R^*) reduced OET incidence to 28%, marking a substantial 1.89-fold decrease compared to the condition without *orf121* (P*_lac_*::*tnpA* with *ter*IS*91*-*cm^R^*). Notably, even without *tnpA* induction, the Orf121 negative effect on OET was evident, as shown by a 2.64-fold reduction in OET (11% with aTc; 29% without aTc, Table 3) when the P*_LtetO-1_* promoter with *orf121*::P*_LtetO-1_*-P*_lac_*::*tnpA* (addition of aTc) was derepressed. These results highlight that Orf121 is crucial in ensuring precise recognition and cleavage at the *ter*IS*91* boundary.

**Table 3.**
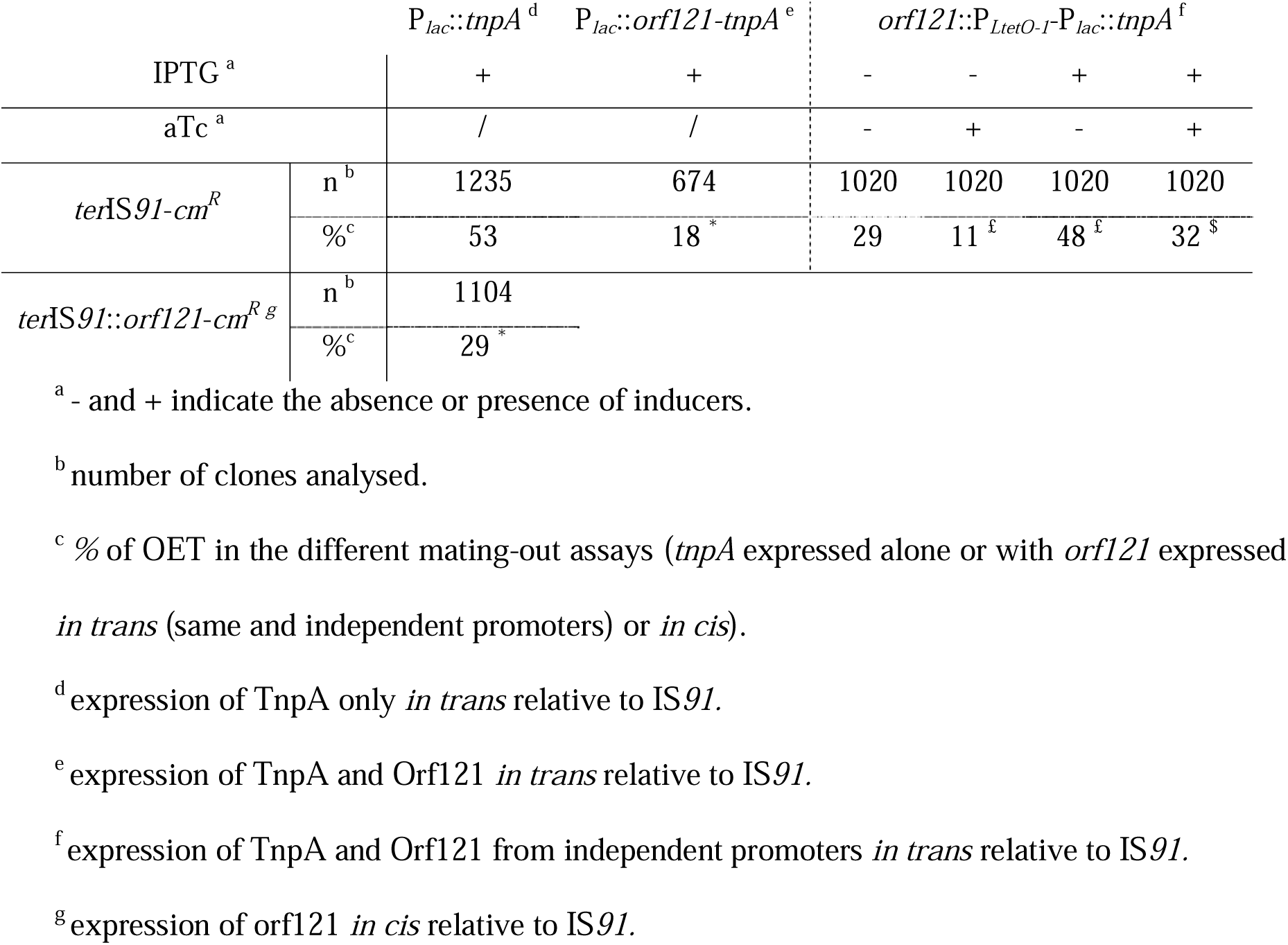
Estimation of the % of OET depending on the presence of Orf121. The n value indicates the number of clones analysed, the - and + indicate the presence or absence of inducer. Impact of *orf121* expression *in cis* or *in trans* on recognition and cleavage of the *terIS* site. The exponents * indicate that the difference between the percentages is significant compared to the condition without *orf121* expression, i.e. the observed difference cannot be due to random sampling. The exponents ^£^ indicate that the difference between the percentages is significant compared to the condition without inducer, i.e. the observed difference cannot be due to random sampling. The exponents ^$^ indicate that the percentage difference is significant compared to IPTG-only.

## Discussion

IS*91* as the prototype for both the IS*91* family and the related IS*CR*s family has often been associated with or located near antibiotic resistance genes^16–20^, suggesting that these IS elements might play a role in the mobilization of antibiotic resistance. Despite its biological and clinical relevance, this family remains poorly characterized, necessitating further studies into its regulation and transposition mechanisms. In this study, we mechanistically explored the IS*91* transposition and showed the crucial role *orf121* plays in the transcriptional and translational regulation of the IS*91* TnpA transposase.

Our findings reveal that *orf121* is a conserved feature across most IS*91* isoforms, with its stop codon overlapping the start codon of *tnpA*. This configuration establishes a transcriptional coupling mechanism wherein orf121 ensures *tnpA* transcription while negatively regulating transposition frequency. The intrinsic P*_orf121_* promoter is significantly stronger than P*_tnpA_*, driving the expression of both genes. The transcriptional coupling, supported by the identification of a potential slippage region in *orf121* (5’-A_192_AACAAAAA_200_), aligns with previous models of RNA polymerase-mediated slippage observed in other insertion sequences^21,22^. Interestingly, isoforms with disrupted overlap due to homopolymeric insertions or deletions may exhibit altered regulatory dynamics, leading to a truncated *orf121* gene and loss of overlap with *tnpA*, which may alter *tnpA* expression levels, highlighting the evolutionary flexibility of this mechanism. Further studies are required to analyse in depth the overlap between the two genes and the effect of the potential slippage region in *orf121* on *tnpA* transcription, thereby determining which of the IS*91* isoforms identified can activate transposition. This process involves a -1 frameshift that allows the ribosome to slip one base upstream and continue translating in an alternative reading frame, as shown in other IS elements^23–26^. The mechanism of transcriptional coupling observed in *orf121* and *tnpA* resembles that described for other insertion sequences, such as IS*10*^27^ and IS*50*^28^, overlapping regulatory regions introduce modulate translation efficiency.

Orf121 demonstrates a dual regulatory function: (i) facilitating *tnpA* transcription while reducing TnpA protein levels and (ii) limiting transposition frequency and insertion efficiency. Co-expression of *orf121* with *tnpA* significantly decreases transposition frequency, irrespective of whether the genes are expressed *in cis* or *trans*. This effect of Orf121 on transposition can be explained by: i) a direct interaction between the Orf121 and TnpA proteins, akin to what has been reported for the IS*911* element^29^, ii) considering the impact of *orf121* expression on *tnpA* mRNA levels, we cannot exclude the possibility of mRNA-mRNA interactions, mRNA-protein interactions, or a role for *orf121* in stabilizing *tnpA* mRNA, similar to what has been suggested for IS*10*^27^. Indeed, we could have the presence of a third partner protein in the regulation, the strand opposite the sequence of *orf121* and *tnpA* has never been studied, although the presence of two other potentials ORFs has been suggested^9^. Additionally, our *in vivo* mating-out assays indicate that Orf121 exerts a negative control over transposition frequency when the two genes are expressed under the control of separate promoters. This observation raises the possibility that the Orf121 protein interacts directly with TnpA, or alternatively, that Orf121 competes with TnpA for binding to the *ori*IS*91* and *ter*IS*91* by forming a dimer. An analysis of the Orf121 peptide sequence identified a potential leucine zipper motif (V49-X_5_-L55-X_5_-V61 or V61-X_6_-L68-X_6_-V75) that could be involved in DNA binding, along with a classic zinc finger motif, CxxC, which might facilitate transposase interaction or dimerization (C105-X_2_-C108) (Fig. 6a). The element appears to function as a dimer, with cysteines 105 and 108 being crucial for its activity and possibly for the regulation of IS*91* transposition^30^. Further investigation into the Orf121 protein could significantly advance our understanding of its role and the mechanisms underlying IS*91* transposition.

**Figure 6.**
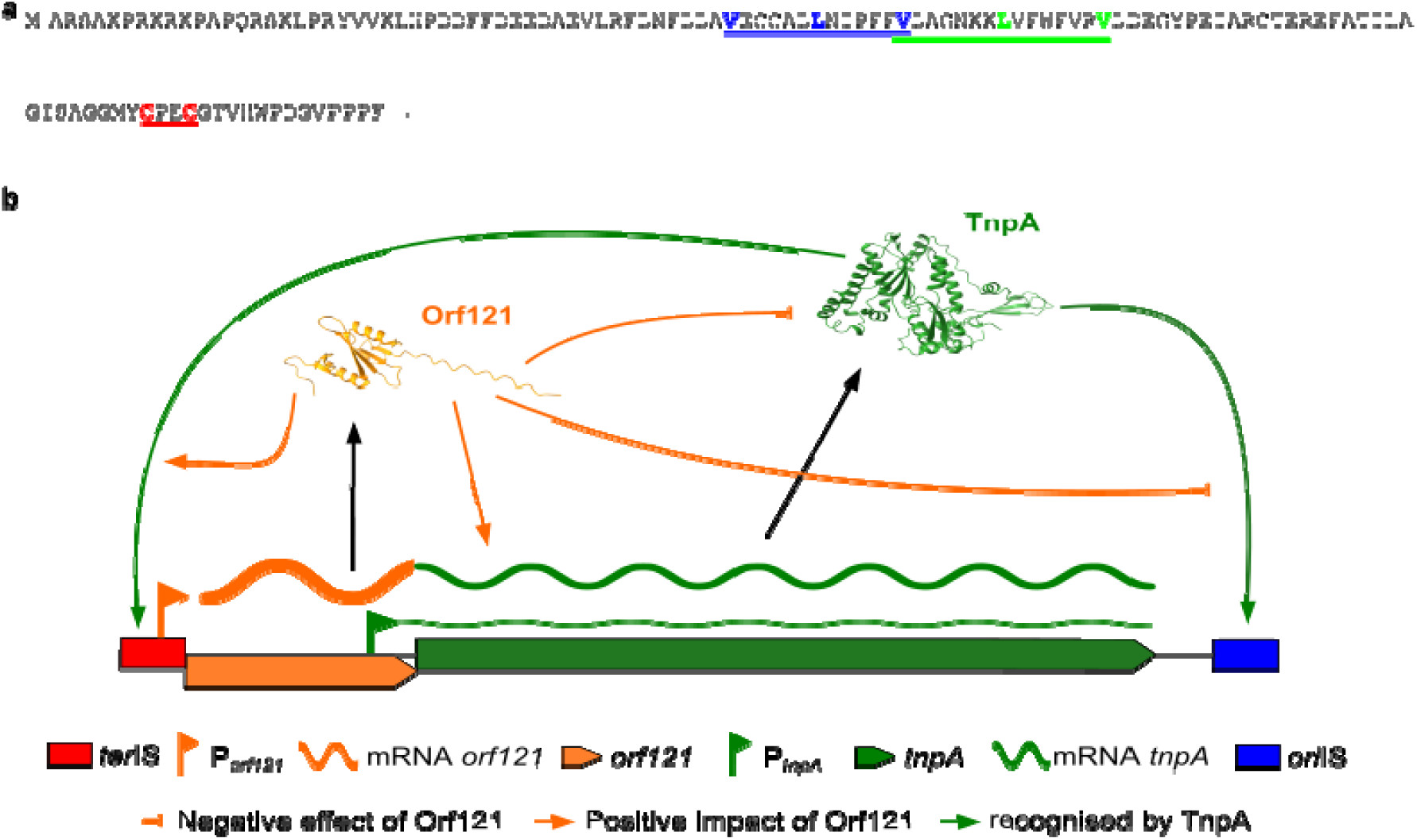
Global diagram of the role of Orf121 in IS*91* mobility. a) Amino acid sequence of Orf121 (accession number CAC14584.1). In blue, bold and highlight or in green, bold and highlight, the first and second potential leucine zipper interacting with DNA. In red, bold and highlight the CxxC domain of zinc fingers enabling dimer formation or interaction with TnpA. b) Global IS*91* interaction model involving Orf121. *orf121* and *tnpA* mRNA are primarily transcribed from the P*_orf121_* promoter. The overlap between the two genes reduces the rate of *tnpA* transcript via the -1 shift between the STOP codon of *orf121* and the start codon of *tnpA*. Orf121 can interact directly with TnpA or *tnpA* mRNA, negatively regulating its activity in the element’s mobility capacity (cyan arrow). Orf121 via dimer formation could bind to the *ori*IS end and compete with TnpA for site binding, reducing element transposition (red arrow). Orf121, through interaction with TnpA or by binding to the *ter*IS end directly, enables better recognition of the *ter*IS end and clean excision of the element (green arrow). The 3D structures of Orf121 and TnpA proteins were modeled with AlphaFold3.

Moreover, the Western-immunoblotting analysis demonstrated that the quantity of TnpA protein is significantly lower in the natural gene configuration than when the tnpA gene is expressed alone. These results suggest that the natural configuration might sequester translation initiation signals, leading to a reduction in protein production. This hypothesis is supported by studies on IS*10*^31^ and IS*50*^28^, where it has been proposed that secondary mRNA structures could sequester the translation initiation signals of the *tnpA* gene, reducing protein synthesis. Consequently, while the natural gene configuration is essential for *tnpA* transcription, it simultaneously acts as a negative regulator of TnpA protein production.

Our findings demonstrate that Orf121 plays a critical role in regulating the insertion efficiency of single-stranded circular intermediates during IS*91* transposition. Specifically, Orf121 negatively impacts the transposition process by significantly reducing the efficiency of bottom-strand intermediate insertion. This regulatory effect indicating a robust mechanism of control independent of the gene’s positional context. Mechanistically, this inhibition may involve Orf121 competing with TnpA for binding to the insertion machinery or interfering with the proper recognition of target sites by forming specific protein-DNA or protein-protein interactions. Additionally, the interaction between Orf121 and the transposition complex could restrict the availability or conformation of the bottom-strand circular intermediate, thereby limiting its ability to integrate into the recipient DNA. These findings highlight the dual functionality of Orf121, not only ensuring accurate excision of the circular intermediate but also finely tuning its insertion frequency to prevent potential genomic instability.

Orf121 ensures accurate excision and processing of the circular intermediate by facilitating precise cleavage at the *ter*IS*91*, our data demonstrate that Orf121 enhances the accurate recognition and cleavage of the *ter*IS*91*, reducing the occurrence of OET and limiting the mobilization of adjacent genes. This precision is crucial, as unchecked OET could facilitate the dissemination of resistance and virulence determinants. Indeed, it has been shown that IS*91*-type elements, such as IS*801*, can mobilize DNA fragments as large as 40 kb^32^. The exact molecular mechanism underlying this regulation remains unknown but may involve Orf121-TnpA interactions or the modulation of target DNA recognition through protein-DNA complexes.

This study highlights the pleiotropic role of *orf121* in IS*91* transposition, underscoring its triple function (Fig. 6b): (i) ensuring transcription of *orf121* and *tnpA via* the P*_orf121_* promoter; (ii) modulating tnpA transcript and protein levels to control transposition frequency; and (iii) enhancing precise cleavage at *ter*IS*91*, limiting the spread of adjacent genes. These findings not only expand our understanding of IS*91* biology but also offer broader insights into the regulation of transposable elements associated with resistance gene dissemination.

## Materials and Methods

### Bacterial strains, media and growth conditions

Bacterial strains and plasmids used in this study are listed in Table 1 and supplementary Table 3. All *E. coli* strains were grown in Lysogeny Broth (LB) at 37°C under agitation at 300 rpm. Media were supplemented, when necessary, with the appropriate antibiotics used at the following concentrations: rifampicin 50 μg/mL, spectinomycin 50 μg/mL, kanamycin 25 µg/mL, chloramphenicol 25 µg/mL, nalidixic acid 25 µg/mL, and ampicillin 100 µg/mL. The expression of IS*91 tnpA* alone or the co-expression with *orf121* from the P*_lac_* promoter was induced by adding 0.5 mM IPTG to the media. The expression of *orf121* from the P*_LTet-O1_* promoter was induced by adding 50 nM anhydrotetracycline (aTc) to the media. Transformation of *E. coli* with plasmid DNA was performed as previously described ^39^. All plasmid constructions were verified by DNA sequencing.

### DNA manipulations

PCR reactions were carried out using Phusion DNA Polymerase (ThermoScientific) to amplify the fragments used for cloning and Quick-Load® Taq 2X Master Mix (NEB) for all other applications. All PCR reactions were purified using MACHEREY-NAGEL’s NucleoSpin Gel and PCR Clean-up according to the manufacturer’s recommendations.

IS*91* derivate insertion sites in the pOX38Km reporter plasmid or the *E. coli DH5α* genome, were mapped by arbitrary priming PCR (AP-PCR). The first PCR round was performed in a final volume of 50 μl containing 0.8 μM of primers (arbitrary primer, ARB1 and chloramphenicol resistance gene-specific, Cm-1005) and 15 μl of template (a single Cm^R^ colony lysed in 20 µl). PCR was performed as follows: 2 min 95°C, 6 cycles of 30 sec 95°C, 30 sec 30°C, 1 min 30 sec 72°C; 30 cycles of 30 sec 95°C, 30 sec 45°C, 2 min 72°C; and finally, 72°C for 5 min. The second PCR cycle was performed in a final volume of 50 μl containing 0.8 μM of primers (arbitrary primer ARB2 and Cm-1049) and 5 μl of the purified PCR product from the first cycle as template. PCR was performed as follows: 2 min 95°C, 30 cycles (30 sec 95°C, 30 sec 57°C, 2 min 72°C) and a final step 72°C for 5 min. The PCR products were directly sequenced using the Ori91-1104 primer.

The *ter*IS cleavage site was mapped using the same procedure with primer pairs ARB1/Cm- 473 for the first cycle of PCR and ARB2/Ter91-337 for the second cycle, and the PCR products were directly sequenced with the Ter91-276 primer.

Oligonucleotides used for PCR amplification of DNA fragments required for plasmid construction, diagnostic PCR or sequencing are described in supplementary Table 3.

### *In silico* analysis of IS*91* DNA sequences from the public database

The amino acid sequence of the TnpA transposase isoforms encoded by IS*91* (accession number CAD30457.1) were blasted with BLASTp (NCBI). The matching sequences of TnpA were filtered to retain only those sequences for which the level of amino acid identity was higher (or equal) than 95%. Recovered nucleotide sequences in which the IS and TnpA were partial or truncated were discarded. For each accession number recovered, we analysed the target insertion and cleavage sites and data related to the bacterial host, the host of the IS and the genetic support. Genetic support was analysed using the PLSDB site ^40,41^. When the PLSDB site gave no results, we searched for plasmid-specific genes (such as *PsiA*-*B*, *StbA*-*B*, *RepA…*) or chromosome-specific genes (such as *DNAa*, *SeqA*, *DNAt*) to identify the genetic support. To define each isoform, we used a 95% nucleotide identity threshold over the entire IS with at least one nucleotide that differs. This data extraction was carried out on 2020-10-15 for IS*91*.

### Estimation of *in vivo* transposition frequencies of IS*91* derivatives into *E. coli* by mating- out assay

Mating-out assays were performed as previously described^42^. The donor strain was *E. coli* JS219^43^ carrying three plasmids: i) *ter*IS*91*-*cm^R^* or *ter*IS*91*::*orf121*-*cm^R^* (used as IS*91*-Cm^R^ derivatives donor plasmids P_IS_; Amp^R^), ii) P*_lac_*::*tnpA*, P*_lac_*::*orf121*-*tnpA* or *orf121*::P*_LtetO-1_*- P*_lac_*::*tnpA* (used as transposase +/- *orf121* expression plasmids; Sp^R^) and iii) pOX38Km, the conjugative target plasmid (Km^R^) (Table 1, Fig. 3.b and supplementary Table 3). The recipient strain was *E. coli* MC240 (Rif^R^, Nal^R^). Following mating of the donor with the recipient strain, cells were plated on LB plates supplemented with rifampicin and kanamycin (count of exconjugants i.e. all recipients carrying pOX38Km without and with IS*91*-Cm^R^ derivatives) or nalidixic acid, kanamycin and chloramphenicol (count of transposants i.e. recipients carrying pOX38Km with IS*91*-Cm^R^ derivatives). Transposition frequency was determined by dividing the CFU of transposants (Nal^R^, Km^R^ and Sp^R^) by the CFU of exconjugants (Nal^R^, Km^R^). Transposition event frequencies were estimated from more than 5 independent experiments. These experiments were used to calculate mean values and standard deviations.

### Estimation of the proportion of one-ended transposition events in mating-out assay

Several hundred clones resulting from transposition were subcultured onto both LB plates supplemented with kanamycin, ampicillin and nalidixic acid (numeration of recipient cells with pOX38Km carrying the whole derivatives donor plasmids P_IS_ resulting from OET), and LB plates supplemented with kanamycin, chloramphenicol and nalidixic acid (Total population of recipients carrying pOX38Km with IS*91*-Cm^R^ derivatives). The percentage of OET was determined by dividing the number of (Km^R^, Amp^R^ and Nal^R^) clones by the number of (Km^R^, Cm^R^ and Nal^R^) clones and multiplying by 100.

### Estimation of *in vivo* insertion of IS*91* circular intermediates into *E. coli*

<u>To test the double-stranded circular intermediate</u>, *E. coli DH5α* cells were rendered competent in the presence or absence of the inducer IPTG (0.5 mM) containing the plasmid P*_lac_*::*tnpA* or P*_lac_*::*orf121*-*tnpA*. These competent cells were transformed with 500 ng of suicide plasmids *ori91*-*ter91* or *ter91*-*ori91-*containing the junction in the pSW23T derivative (Table 1, supplementary Table 3). Cells were heat-shocked at 42°C for 1 min, followed by 5 min on ice. The cells were then placed in 1 ml LB with or without inducer for 1 hour at 37°C with agitation, then plated. Determination of the total number of viable cells was performed on LB plates supplemented with spectinomycin (Sp^R^), and insertion of junctions into the *E. coli DH5α* genome was selected on LB plates supplemented with spectinomycin and chloramphenicol (SpR, Cm^R^).

<u>To test the single-stranded circular intermediate</u>, overnight cultures with Cm^R^ + DAP (0.3 mM) for donors (*E. coli* β*2163* (*pir-*) carrying the suicide plasmid (pSW23T (Empty), *ori91*- *ter91* or *ter91*-*ori91*; Table 1) and Sp^R^ for recipients (*E. coli DH5α* target strain containing different *tnpA* and/or *orf121* expression plasmids (P*_lac_*::*tnpA*, P*_lac_*::*orf121*-*tnpA* or *orf121*::P*_LtetO-1_*-P*_lac_*::*tnpA*; Table 1) were diluted 1:100, without antibiotics, and further grown to OD_600_ = 0.3. Expression of *tnpA* and/or orf121 was then induced by adding respectively 0.5 mM IPTG and/or aTc at 50 ng/ml in the recipient cells cultures. The strains were left for a further 1 h at 37°C with agitation. Filters containing the mixture of donor and recipient strains in a 1:8 ratio were incubated for 3 h at 37°C on LB plates supplemented with 0.3 mM DAP and 0.5 mM IPTG, or 0.3 mM DAP, 0.5 mM IPTG and 50 ng/ml aTc. Cells were then suspended in 2 mL LB by vortexing the filter, and appropriate dilutions were plated on selective LB plates (Sp^R^) for recipients and LB plates (Sp^R^ + Cm^R^) for transconjugants. The frequency of Cm^R^ transconjugants was calculated as the number of Cm^R^-tagged recipient cells relative to the total number of recipient cells. The frequencies of the insertion event per viable cell from 4 independent experiments were used to calculate mean values and standard deviations.

### Quantification of *orf121* and *tnpA* transcripts

For mating-out transcripts, overnight cultures were diluted to OD_600_ = 0.2 and grown to the mean log phase (OD_600_ = 0.5). Cells were pelleted and total RNA was extracted with the NucleoSpin® RNA Extraction Kit (Macherey-Nagel Inc.). Contaminating DNA was removed from RNA samples by using the Turbo DNA-free Kit (Ambion). RNA integrity was check and quantified. cDNAs were synthesized from 1 μg of DNase-treated total RNA by using PrimeScriptTM RT Reagent kit (TaKaRa Clontech) following manufacturer’s instruction. cDNA of genes *tnpA*, *orf121* and *dxs* was quantified using the PerfeCTa® SYBR® Green FastMix® Kit (Quanta BioSciencesTM) with appropriate oligonucleotides (supplementary Table 3). Relative expression of *orf121* (Primers ORF121 F and ORF121 R) and *tnpA* (Primers TnpA F and TnpA R) genes was estimated by normalizing copy number of transcripts to that of the housekeeping gene *dxs* (Primer dxs-LC3 and dxs-LC4). The *tnpA*/*dxs* or *orf121*/*dxs* ratio was obtained from 3 independent experiments, which were used to calculate mean values and standard deviations.

### **β**-galactosidases assays

β-galactosidase assays were performed with the *E. coli* MG1656 strain carrying various plasmids (Table 1 ; ^44^). Overnight cultures were diluted 1:100, and further grown to mid-log phase (OD_600_ = 0.4-0.6). The β-galactosidase assay was performed as previously described^45^. β-galactosidase activity was obtained from 3 independent experiments, which were used to calculate mean values and standard deviations.

### Estimation of His-tagged ORF121 or TnpA_IS91_ protein expression by western- immunoblotting

Overnight cultures were diluted to OD_600_ = 0.2 and grown to mid-log phase (OD_600_ = 0.5). Gene expression was induced by adding 0.5 mM IPTG (*tnpA* alone or with and *orf121*) and/or 50 nM aTc (*orf121*) in the media. Cultures were incubated for further 3 h at 37°C, and 1 mL of each culture was centrifuged and stored at -20°C. Cell extracts were prepared using the B-PER bacterial protein extraction kit according to the manufacturer’s recommendations at a rate of 100 μL of reagent per OD unit (Thermo Scientific, Ref. 90078) and inclusion bodies proteins using the solubilization reagent used according to the manufacturer’s instructions (Thermo Scientific, Ref. 78115). Cell extract samples were subjected to 15% SDS-PAGE electrophoresis, followed by a transfer to a PVDF membrane according to the manufacturer’s instructions (Trans-Blot Turbo Transfer System, BioRad). The membrane was incubated in a 1:3000 dilution of mouse anti-His-tag antibody (Invitrogen, Ref. MA121315) overnight at 4°C, then washed 3 times for 10 minutes with PBS. Next, the membrane was incubated in a 1:3000 dilution of anti-Mouse IgG, HRP antibody (Invitrogen, Ref. A28177), then washed 6 times for 5 minutes with what. The membrane was revealed with SuperSignal West Pico PLUS (Thermo Scientific, Ref. 34577). The amount of protein was determined in arbitrary units (AU) by calculating the value of the area under the curve using ImageLab software.

### Analysis of His-tagged ORF121 or TnpA_IS*91*_ proteins by mass spectrometry

Peptides from SDS-PAGE were analyzed by micro-LC-MS/MS using a nanoLC 425 system in micro-flow mode (Eksigent, Dublin, CA, USA) coupled with time-of-flight (TOF) (TripleTOF 5600+ Sciex, Framingham, MA, USA) operating in high-sensitivity mode. Reverse-phase LC was performed via a trap and elute configuration using a trap column (C18 Pepmap100 cartridge, 5 µm pore size; Thermo Fisher Scientific) and an analytical column (ChromXP C18 column, 12 nm, 3 µm pore size, Sciex) with the following mobile phases: loading solvent (water/ACN/TFA 98/2/0.05% (v/v)), solvent A (0.1% (v/v) TFA in water) and solvent B (water/ACN/TFA 5/95/0.1% (v/v)). All samples were loaded, trapped, and desalted using a flow rate of 10 μL/min with loading solvent for 5 minutes. The chromatographic separation was performed at a flow rate of 2 µL/min as follows: initial, 5% solvent B, increased to 25% for 90 minutes, then increased to 95% B for 10 minutes, maintained at 95% for 5 minutes and, finally, decreased to 5% B for re-equilibration. The peptides were then reprocessed via ProteinPilot with the Mascot module using *Escherichia coli* Uniprot 2022_03. The following parameters were entered: trypsin 1 maximum missed cleavage, cysteine carbamidomethylation (fixed), methionine oxidation (variable), Precursor Tolerance 0.01Da; MS/MS fragment tolerance: 0.05 Da, a p<0.05 - peptide score > 25 - bold red required and a protein score > 100.

### Statistical analyses

All analyses were performed using R Statistical Software (v4.4.1) and RStudio (v 2024.04.2+764). ’s Statistical analyses include a Mann-Whitney U-test and a Chi-square test to compare percentages and insertion target sites. Graphics were rendered via the ggplot2 R package (v3.5.1)^46^. All figures were prepared using Inkscape 1.2.2 (https://inkscape.org/).

## Acknowledgements

A. Fauconnier acknowledges the French ministère de l’Enseignement Supérieur, de la Recherche (MESR) for his doctoral training grant. This work was supported by fundings from the French research institute Inserm.

The authors thank Emilie Pinault from BISCEm unit (Univ. Limoges, UAR 2015 CNRS, US 42 Inserm, CHU Limoges) for technical support regarding mass spectrometry.

The authors are grateful to Céline Loot (Institut Pasteur, Université Paris Cité, CNRS UMR3525, Unité Plasticité du Génome Bactérien, 75724 Paris, France) and Vincent Burrus (Département de biologie, Université de Sherbrooke, Sherbrooke, Québec, Canada) for critical review of this article.

## Supplementary tables

**Supplementary Table 1.** Host of bacteria carrying IS*91* elements in GenBank®

**Supplementary Table 2.** Analysis of the effect of Orf121 expression on the IS mimicking IS*91* target insertion sites in the *E. coli* chromosome

**Supplementary Table 3. Oligonucleotides used in this study**

## Supplementary Files

**Supplementary File 1. Sequences of the different IS*91* isoforms described in this study.** IS*91*-V1 corresponds to the first sequence of the IS*91* element identified (number accession X17114.5).

**Supplementary File 2.** Description of the different IS*91* isoforms described in this study. Supplementary File 3. Complete data sets.

