## Supplementary Table 1 for "Dual regulatory role of IS*91*-encoded Orf121 in IS*91* transposition"

**Supplementary Table 1. Host of bacteria carrying IS*91* elements in GenBank®**

The n value indicates the accession number analysed. X indicates the nucleotide unknown

|  |  | **IS*91*** |
| --- | --- | --- |
| **Host of bacteria carrying IS** | n | 924 |
|  | Animals | 451 (48.8%) |
|  | Homo Sapiens | 219 (23.4%) |
|  | Plants | 1 |
|  | Environment | 19 |
|  | Food | 1 |
|  | Unknown | 239 (25.5%) |
| **Bacteria carrying IS** | n | 924 |
|  | *Escherichia coli* | 693 (75%) |
|  | *Salmonella enterica* | 222 (24%) |
|  | *Shigella sonnei* | 6 |
|  | *Shigella flexneri* | 0 |
|  | *Shigella dysenteriae* | 0 |
|  | *Pseudomonas savastanoi* | 0 |
|  | *Klebsiella pneumoniae* | 1 |
|  | *Pseudomonas syringae* | 0 |
|  | *Shigella boydii* | 0 |
|  | *Pseudomonas amygdali* | 0 |
|  | *Pseudomonas avellanae* | 0 |
|  | *Pseudomonas ficuserectae* | 0 |
|  | *Providencia sp.* | 1 |
|  | *Zoogloea sp.* | 1 |
|  | *Escherichia marmotae* | 0 |
|  | *Citrobacter farmeri* | 0 |
|  | *Enterobacter cloacae* | 0 |
| **Genetic support of IS** | n | 924 |
|  | Plasmid | 671 (72.6%) |
|  | Chromosome | 26 |
|  | Unknown | 227 |
| **Number of copies of IS per replicon** | n | 829 |
|  | 1 | 790 (95.3%) |
|  | 2 | 12 |
|  | 3 | 11 |
|  | 4 | 11 |
|  | 5 | 2 |
|  | 6 | 0 |
|  | 7 | 2 |
|  | 8 | 0 |
|  | 9 | 1 |
| **Origin of replication of plasmids carrying IS** | n | 671 |
|  | IncFII | 366 (72.6%) |
|  | R64 / IncI1 family | 38 |
|  | ColE1 | 16 |
|  | IncI1-I(alpha) | 10 |
|  | IncFIB | 3 |
|  | IncHI | 2 |
|  | IncC | 1 |
|  | IncN | 2 |
|  | IncI2 | 1 |
|  | IncY | 9 |
|  | Inc | 4 |
|  | RK2 | 4 |
|  | IncP | 2 |
|  | IncHI2 | 1 |
|  | Col | 0 |
|  | RepHI2 | 0 |
|  | unknown | 212 |
| **Insertion target sites** | n | 924 |
|  | 5’-GTTC | 590 |
|  | 5’-CTTG | 208 |
|  | 5’-CTCG | 64 |
|  | 5’-GTCC | 20 |
|  | 5’-CTGG | 4 |
|  | 5’-GTAC | 5 |
|  | 5’-GTGC | 3 |
|  | 5’-GTTG | 2 |
|  | 5’-TTGC | 1 |
|  | 5’-TTTA | 1 |
|  | 5’-TTAT | 0 |
|  | 5’-CTAX | 0 |
|  | 5’-XXXX | 26 |
| **Cleavage sites** | n | 924 |
|  | 5’-CTCG | 923 |
|  | 5’-GTTC | 0 |
|  | 5’-CTCT | 2 |
|  | 5’-GTAC | 0 |
