## Supplementary Table 2 for "Dual regulatory role of IS*91*-encoded Orf121 in IS*91* transposition"

**Supplementary Table 2. Analysis of the effect of Orf121 expression on the IS mimicking IS*91* target insertion sites in the *E. coli* chromosome**

|  |  | P*_lac_*::*tnpA* ^a^ | P*_lac_*::*orf121tnpA* ^b^ | *orf121*::P*_LtetO-1_*-P*_lac_*::*tnpA* ^d^ | | | |
| --- | --- | --- | --- | --- | --- | --- | --- |
| IPTG ^e^ | | - / + | - / + | - | - | + | + |
| aTc ^e^ | |  |  | - | + | - | + |
| n ^c^ | | 50 | 20 | 57 | 26 | 58 | 58 |
| Sequence of insertion site tetranucleotide | 5’-CTTG | 24 | 15 | 24 | 8 | 19 | 21 |
|  | 5’-GTTC | 13 | 2 | 29 | 18 | 27 | 27 |
|  | 5’-CTCG | 8 | 1 | 3 | 0 | 8 | 10 |
|  | 5’-GTCC | 4 | 2 | 1 | 0 | 2 | 0 |
|  | 5’-GTAC | 1 | 0 | 0 | 0 | 2 | 0 |

^a^ P*_lac_*::*tnpA*^:^ expression of TnpA only

^b^ P*_lac_*::*orf121-tnpA*^:^ expression of TnpA and Orf121

^c^ n value indicates the number of clones analysed by AP-PCR

^d^ *orf121*::P*_LtetO-1_*-P*_lac_*::*tnpA*: expression of TnpA and Orf121 from independent promoters

^e^ - and + indicate the absence or presence of inducers.
