## Supplementary Table 3 for "Dual regulatory role of IS*91*-encoded Orf121 in IS*91* transposition"

**Supplementary Table 3. Oligonucleotides used in this study**

| **Oligonucleotide name** | **Oligonucleotides sequences (5' ⭢ 3')^a^** | **Use** |
| --- | --- | --- |
| **Linker formation** | | |
| CP17 | TATG**GGTACCGTCGACGGATCC**GCATG | Formation of linker79 (P*_Lac_* (Empty)) |
| CP18 | C**GGATCCGTCGACGGTACC**CA |  |
| CP21 | **AATTC**CGAGTAGGCAGCCTGGCGGCTGCGGCTTGTCATGGTCTGGAATTACCGTTATAAAAAAAGATAATGTCATTGTCTTTCAGGTAGT**G** | Formation of linkerTer91 (*ter*IS*91*-*cm^R^*) |
| CP22 | **GATCC**ACTACCTGAAAGACAATGACATTATCTTTTTTTATAACGGTAATTCCAGACCATGACAAGCCGCAGCCGCCAGGCTGCCTACTCG**G** |  |
| CP19 | **G**TCCGCTCATATGGTGCACAAGGGGTGTTGAAGAAACATCCGTTTTGTGGTGCTTTTTTAGTCTTTTGGGGATTTAAATTCCTATCGATCAAG**A** | Formation of linkerOri91 (*ter*IS*91*-*cm^R^, ter*IS*91*::*orf121*-*cm^R^*) |
| CP20 | **AGCTT**CTTGATCGATAGGAATTTAAATCCCCAAAAGACTAAAAAAGCACCACAAAACGGATGTTTCTTCAACACCCCTTGTGCACCATATGAGCGGA**CTGCA** |  |
| **Plasmid construction** | | |
| CP23 | TACT**GGATCC**TGAGACGTTGATCGGCAC | Amplification of the P*_cat_*:Cam^R^ cassette (*ter*IS*91*-*cm^R^* intermediate) |
| CP24 | TGAG**CTGCAG**TGCCACTCATCGCAGTAC |  |
| CP56 | CCG**GAATTC**GAGTAGGCAGCCTGGCGGC | Amplification of *terIS91*-*orf121*-P_tnpA_ (*ter*IS*91*::*orf121*-*cm^R^*) |
| CP75 | CGC**GGATCC**TCAGAAGGGCGGGGGGACTCC |  |
| CP124 | CCG**CTGCAG**TTGCTTTCAGGAAAATTTTTCTG | Addition of the tet^R^ gene to repress the P*_LtetO-1_* promoter in plasmids *orf121*::P*_LtetO-1_*-P*_lac_*, *orf121*::P*_LtetO-1_*-P*_lac_*::*tnpA, orf121*^6his^::P*_LtetO-1_-*P*_Lac_*::*tnpA*^6his^ |
| CP125 | CCG**CTGCAG**ATTCTCACCAATAAAAAACG |  |
| CP56 | CCG**GAATTC**GAGTAGGCAGCCTGGCGGC | Amplification of *ter*IS*91*-*orf121* (*ter*IS*91*::*orf121*-*cm^R^*) |
| CP75 | CGC**GGATCC**TCAGAAGGGCGGGGGGACTCC |  |
| CP15 | TATA**CATATG**CTTCCCCGTTTTGCCGAC | Amplification of the CDS of *tnpA*_IS_*_91_* (P*_lac_*::*tnpA*) |
| CP16 | GCGT**GGATCC**TATTAAACAGGATGACACTG |  |
| AF4 | ATAT**GGATCC**TCAATGATGATGATGATGATGGCCGCACACCTGCTGTCG | Amplification of the *tnpA*_IS_*_91_* CDS with a C-terminal His6 tag (P*_Lac_*::*tnpA*^6his^) |
| CP58 | TATA**CATATG**GCCCGTTCAGCTAAACC |  |
| CP58 | TATA**CATATG**GCCCGTTCAGCTAAACC | Amplification of the CDS of *orf121*- *tnpA*_IS_*_91_* (P*_lac_*::*orf121*-*tnpA*) |
| CP16 | GCGT**GGATCC**TATTAAACAGGATGACACTG |  |
| AF4 | ATAT**GGATCC**TCAATGATGATGATGATGATGGCCGCACACCTGCTGTCG | Amplification of the CDS of *orf121*- *tnpA*_IS_*_91_* with a His6 tag at the C-terminus of *tnpA*_IS_*_91_* (P*_Lac_*::*orf121tnpA*^6his^) |
| CP58 | TATA**CATATG**GCCCGTTCAGCTAAACC |  |
| CP16 | GCGT**GGATCC**TATTAAACAGGATGACACTG | Amplification of the CDS of *orf121* - *tnpA*_IS_*_91_* with an N-terminal His6 tag of *orf121* (P*_Lac_*:: ^6his^ *orf121tnpA*) |
| CP86 | TATA**CATATG**CATCATCATCATCATCATGCCCGTTCAGCTAAACC |  |
| CP88 | CTGCCCGCTTT**CCTCGAGT**CCCTATCAGTGATAGAGATTG | Amplification of the P*_LtetO-1_* portion portion (170 bp) |
| CP89 | CCATATAACTACCTCGGTCAGTGCGTCCTG |  |
| CP90 | TCTA**GAATTC**TCAGAAGGGCGGGGGGACTC | Amplification of the *orf121* portion (386 pb) |
| CP91 | GGACGCACTGACCGAGGTAGTTATATGGC |  |
| CP88 | CTGCCCGCTTT**CCTCGAGT**CCCTATCAGTGATAGAGATTG | Amplification of the *orf121*::P*_LtetO-1_* (*orf121*::P*_LtetO-1_*-P*_lac_*) |
| CP90 | TCTA**GAATTC**TCAGAAGGGCGGGGGGACTC |  |
| AF68 | CGG**GGTACC**ATGCTTCCCCGTTTTGCCGAC | Amplification of the CDS of *tnpA*_IS_*_91_* (*orf121*::P*_LtetO-1_*-P*_lac_*::*tnpA*) |
| CP16 | GCGT**GGATCC**TATTAAACAGGATGACACTG |  |
| AF69 | ACACTCCGGGCAGTACATGC**CGCCGGCG**CTGATACCGGCAAGAATG | Amplification of the CDS of *orf121*::P*_LtetO-1_*-P*_Lac_*::*tnpA*_IS_*_91_* with a His6-tag at the N-terminus of *orf121* and at the C-terminus of *tnpA*_IS_*_91_* (*orf121*^6his^::P*_LtetO-1_-*P*_Lac_*::*tnpA*^6his^; *in fusion*) |
| AF71 | GCACTGACCGAGGTAGTTATATGCATCATCATCATCATCATGC |  |
| AF70 | ATGATGATGATGATGATGCATATAACTACCTCGGTCAGTGC |  |
| AF73 | GTCGGCAAAACGGGGAAGCATATGTATATCTCCTTCTTAAA |  |
| AF72 | TTTAAGAAGGAGATATACATATGCTTCCCCGTTTTGCCGAC |  |
| AF74 | TCTGTATGGAACGGGCATGC**GGATCC**TCAATGATGATGATGATGATG |  |
| CP79 | GCTGCCTACTCGATCGATAGGAATTTAAATCC | Amplification of the *ori*IS*91* portion (159 bp) |
| Cm-1049 | GCTTCCATGTCGGCAGATGC |  |
| CP80 | ATTCCTATCGATCGAGTAGGCAGCCTGGCGGCTG | Amplification of the *ter*IS*91*portion (113 bp) |
| Ter91-337 | CAACGTCTCAGGATCCACTACCTG |  |
| CP79 | GCTGCCTACTCGATCGATAGGAATTTAAATCC | Amplification of the *ori*IS*91*-BamHI portion (110 bp) |
| CP81 | ATAT**GGATCC**TCCGCTCATATGGTGCACAAG |  |
| CP80 | ATTCCTATCGATCGAGTAGGCAGCCTGGCGGCTG | Amplification of the *ter*IS*91*-PstI portion (107 bp) |
| CP82 | GTGG**CTGCAG**ACTACCTGAAAGACAATGAC |  |
| CP81 | ATAT**GGATCC**TCCGCTCATATGGTGCACAAG | Amplification of the Junc *ter*IS*91*- *ori*IS*91*-INV fusion fragment by junction PCR (*ori91*-*ter91, ter91*-*ori91*) |
| CP82 | GTGG**CTGCAG**ACTACCTGAAAGACAATGAC |  |
| CP110 | CGC**GAATTC**ATGGCCCGTTCAGCTAAAC | Amplification of the fragment containing the P*_tnpA_* promoter (boxes -35 and -10) with RBS*_lacZ_* (P*_tnpA_*::RBS*_lacZ_*-*lacZ*) |
| CP116 | GCG**GGATCC**ATGGCTGTTTCCTGTCCGGCCAGTGAACCGTG |  |
| CP95 | CGC**GAATTC**GAGTAGGCAGCCTGGCGGC | Amplification of the fragment containing the P*_orf121_* promoter (boxes -35 and -10) with RBS*_lacZ_* (P*_orf121_*::RBS*_lacZ_*-*lacZ*) |
| CP96 | GCG**GGATCC**ATGGCTGTTTCCTGAAAGACAATGACATTATC |  |
| CP110 | CGC**GAATTC**ATGGCCCGTTCAGCTAAAC | Amplification of the fragment containing the P*_tnpA_* promoter (box -35 and -10) set with RBS*_tnpA_* and an overlap between the stop codon of *orf121* and start of *lacZ* (P*_tnpA_*::*lacZ*) |
| CP121 | GCG**GGATCC**ATCAGAAGGGCGGGGGGA |  |
| CP95 | CGC**GAATTC**GAGTAGGCAGCCTGGCGGC | Amplification of the fragment containing the P*_orf121_* promoter (boxes -35 and -10) set with RBS*_orf121_* (+ P*_tnpA_* promoter in CDS of *orf121*) and an overlap between the stop codon of *orf121* and start of *lacZ* (P*_orf121_*-*orf121*_(P_*_tnpA_*_)_::*lacZ*) |
| CP121 | GCG**GGATCC**ATCAGAAGGGCGGGGGGA |  |
| CP110 | CGC**GAATTC**ATGGCCCGTTCAGCTAAAC | Amplification of the fragment containing the P*_tnpA_* promoter (box -35 and -10) set with RBS*_tnpA_* and WITHOUT an overlap between the stop codon of *orf121* and start of *lacZ* (P*_tnpA_*::*lacZ* Δ-1) |
| CP128 | GCG**GGATCC**ATTCAGAAGGGCGGGGGGAC |  |
| CP95 | CGC**GAATTC**GAGTAGGCAGCCTGGCGGC | Amplification of the fragment containing the P*_orf121_* promoter (box -35 and -10) set with RBS*_orf121_* (+ P*_tnpA_* promoter in CDS of *orf121*) and WITHOUT an overlap between the stop codon of *orf121* and start of *lacZ* (P*_orf121_*-*orf121*_(P_*_tnpA_*_)_::*lacZ* Δ-1) |
| CP128 | GCG**GGATCC**ATTCAGAAGGGCGGGGGGAC |  |
| CP110 | CGC**GAATTC**ATGGCCCGTTCAGCTAAAC | Insertion of a mutation in the -35 box of the P*_tnpA_* promoter left portion (305 bp) |
| AF80 | CTGATACCCGCCAGAATGGTCGCAAAC |  |
| CP121 | GCG**GGATCC**ATCAGAAGGGCGGGGGGA | Insertion of a mutation in the -35 box of the P*_tnpA_* promoter on the right-hand side (102 bp) |
| AF79 | CGACCATTCTGGCGGGTATCAGCGCC |  |
| CP110 | CGC**GAATTC**ATGGCCCGTTCAGCTAAAC | Amplification of the Junc fusion fragment mutated in box -35 of the P*_tnpA_* TTGCCG promoter into TGGCGG (P*_orf121_*-*orf121*_(P_*_tnpA_**_)_::*lacZ*) |
| CP121 | GCG**GGATCC**ATCAGAAGGGCGGGGGGA |  |
| CP95 | CGC**GAATTC**GAGTAGGCAGCCTGGCGGC | Insertion of a mutation in the -35 box of the P*_tnpA_* promoter, left portion (392 bp) |
| AF80 | CTGATACCCGCCAGAATGGTCGCAAAC |  |
| AF79 | CGACCATTCTGGCGGGTATCAGCGCC | Insertion of a mutation in the -35 box of the P*_tnpA_* promoter on the right-hand side (102 bp) |
| CP121 | GCG**GGATCC**ATCAGAAGGGCGGGGGGA |  |
| CP95 | CGC**GAATTC**GAGTAGGCAGCCTGGCGGC | Amplification of the Junc fusion fragment mutated in box -35 of the P*_tnpA_* TTGCCG promoter to TGGCGG (P*_orf121_*-*orf121*_(P_*_tnpA_**_)_::*lacZ*) |
| CP121 | GCG**GGATCC**ATCAGAAGGGCGGGGGGA |  |
| CP95 | CGC**GAATTC**GAGTAGGCAGCCTGGCGGC | Insertion of a mutation in the -10 box of the P*_orf121_* promoter, left portion (72 bp) |
| AF78 | TTATCTTTTTTCGCGACGGTAATTCCAG |  |
| AF77 | AATTACCGTCGCGAAAAAAGATAATGTC | Insertion of a mutation in the -10 box of the P*_orf121_* promoter right portion (424 bp) |
| CP121 | GCG**GGATCC**ATCAGAAGGGCGGGGGGA |  |
| CP95 | CGC**GAATTC**GAGTAGGCAGCCTGGCGGC | Amplification of the Junc fusion fragment mutated in box -10 of the P*_orf121_* TATAAA promoter into CGCGAA (P*_orf121_**-*orf121*_(P_*_tnpA_*_)_::*lacZ*) |
| CP121 | GCG**GGATCC**ATCAGAAGGGCGGGGGGA |  |
| **Arbitrary-primed (AP)-PCR** | | |
| ARB1 | GGCCACGCGTCGACTAGTACNNNNNNNNNNATCGG | First round of arbitrary PCR to map IS*91* insertion sites into pOX38Km |
| Cm-1005 | GCAGATTACGCGCAG |  |
| ARB2 | GGCCTCGCGTCGACTACTTC | Second round of arbitrary PCR to map IS*91* insertion sites into pOX38Km |
| Cm-1049 | GCTTCCATGTCGGCAGATGC |  |
| ARB1bis | GGCCTCGCGTCGACTACTTCNNNNNNNNNNGACTG | First round of arbitrary PCR to map IS*91* insertion sites into *E. coli* chromosome |
| Cm1005-Bis54 | CCCTTATTCGCACCTGGCGG |  |
| ARB2 | GGCCTCGCGTCGACTACTTC | Second round of arbitrary PCR to map IS*91* insertion sites into *E. coli* chromosome |
| Cm1049-Bis54 | GGCGTCGACGGTATCGATAAGC |  |
| Ori91-1104 | GCAGTCCGCTCATATGGTGC | Sequencing of arbitrary PCR products |
| ARB1 | GGCCACGCGTCGACTAGTACNNNNNNNNNNATCGG | First set of arbitrary PCRs on the *ter*IS*91* side in pOX38Km with ARB1 |
| Cm-473 | ATCAACGGTGGTATATCCAGTG |  |
| ARB2 | GGCCTCGCGTCGACTACTTC | Second set of arbitrary PCRs on the *ter*IS*91* side in pOX38Km with ARB2 (*ter*IS*91*-*cm^R^*) |
| Ter91-337 | CAACGTCTCAGGATCCACTACCTG |  |
| ARB2 | GGCCTCGCGTCGACTACTTC | Second set of arbitrary PCRs on the terIS91 side in pOX38Km with ARB2 (*ter*IS*91*::*orf121*-*cm^R^*) |
| Orf121-484 | GTCTGCGCAGCACTCAACG |  |
| Ter91-276 | TCCAGACCATGACAAGCCGC | For Sanger sequencing of the *ter*IS*91* side (*ter*IS*91*-*cm^R^*) |
| Ter91-318 | CTACCTGAAAGACAATGAC | For Sanger sequencing of the *ter*IS*91* side (*ter*IS*91*::*orf121*-*cm^R^*) |
| **qRT-PCR primer** | | |
| ORF121 F | GTGCTGCGCAGACCTGAATAT | *orf121* |
| ORF121 R | ACTCCGGGCAGTACATGCC |  |
| TnpA F | TCCCTGGTGTTCCACAACCGGT | *tnpA* |
| TnpA R | CACACCACCGGCAGTTGTCG |  |
| dxs-LC3 | ATGACGTGGCGATTCAAAA | housekeeping gene *dxs*^1^ |
| dxs-LC4 | AGCCGGTATAGAGCATCTGG |  |

^a^tags with restriction site are in bold.

^1^ Baltazar M, Bourgeois-Nicolaos N, Larroudé M, Couet W, Uwajeneza S, Doucet-Populaire F, Ploy MC, Da Re S. Activation of class 1 integron integrase is promoted in the intestinal environment. PLoS Genet. 2022 Apr 28;18(4):e1010177. doi: 10.1371/journal.pgen.1010177. PMID: 35482826; PMCID: PMC9090394.
