## Supplementary File 1 for "Dual regulatory role of IS*91*-encoded Orf121 in IS*91* transposition"

>IS91-V1

CGAGTAGGCAGCCTGGCGGCTGCGGCTTGTCATGGTCTGGAATTACCGTTATAAAAAAAGATAATGTCATTGTCTTTCAGGTAGTTATATGGCCCGTTCAGCTAAACCCCGTAAACGCAAACCCGCACCACAAAGAAGCAAACTTCCCCGCTATGTTGTGAAACTTCATCCGGATGATTTTTTTGACGAAGAAGACGCTGAAGTTCTGCGCTTTGATAATTTTGACGATGCCGTTGAGTGCTGCGCAGACCTGAATATTCCCTTCTTTGTGGATGCCGGAAACAAAAAGCTGGTCTTCTGGTTTGTACGTGTTGATGACGAAGGGTATCCTGAAATAGCCCGCTGCACGGAGCGGGAGTTTGCGACCATTCTTGCCGGTATCAGCGCCGGCGGCATGTACTGCCCGGAGTGTGGCACGGTTCACTGGCCGGACGGAGTCCCCCCGCCCTTCTGATGCTTCCCCGTTTTGCCGACATTTTTCAGCAGGGTAACCGCTGGCTTAACTGGCTGGAGAAACAGCCGGAAGGTTCAGTGCGTCCGGTGGTGACTGAGTCAGTGACAAAAATCATGGCATGCGGGACCACGCTGATGGGCTACACGCAATGGTGCTGTTCGTCACCGGACTGTTGCCACACCAAAAAGGTCTGCTTCCGGTGTAAAAGCCGCTCCTGTCCGCACTGCGGGGTGAAGGCTGGCGCACAGTGGATACAGTATCTGCTGAGCCTGGTCCCCGACTGCCCGTGGCAGCATATTGTGTTCACACTTCCCTGCCAGTACTGGTCCCTGGTGTTCCACAACCGGTGGTTACTGGCAGAGATGAGCCGCATTGCAGCGGATGTGATACTGGAAATCTGCCATCAGACAGATGTGGAGCCGGGGATATTCACGGTGATCCACACATGGGGGCGTGACCAGCAGTGGCATCCGCATATCCATTTATCGACAACTGCCGGTGGTGTGACGTCGGGCCACACCTGGAAAAATCTTCATTTTTACGCCCGTAAGGTGATGAGCATGTGGCGTTACCGGATAACGCGGCTACTGTCCCGGAAATACCCGGAGCTGGTGATACCGGATGAACTGGCAGTGGAAGGAAACAGCAAACGGGACTGGAATTGCTTCCTGGACACGCATTACCGCCGCGGCTGGAATGTCAACATATCCAGGGTGATGGATAACGCCACACATGTGGCGGTGTACTTCGGCTCTTACCTGAAAAAGCCACCGGTGCCGATGAGTCGGCTGGAGCATTATGCCGGTCAGGATGAAATCGGTCTGCGTTACAACAGTCACCGTACAAAACGGGAAGAATACCTGTTGATGAGTGGAGATGAGTTCATGGAAAGGTTCTCCTGGCATGTAGCAGATAAGGGGTTCCGTATGGTGAGGTACTACGGTTTCCTGAGTCCGGTGAAGCGCCGGTTACTGGAAGAAGTTGTGTACGTCATAACGGAGACGGTGAGAAAAACGGCGATGCAAATCAGGTGGAGAGGGATGTATCAGCGGTTACTGAAGGTTGACCCGCTGAAGTGCATTCTGTGCGGAAGTCAGATGCGTTTTACGGGGCTGAAGCGGGGTTACCGACTGGCAGAGCTGGTCCTGATGCATGAGCGACTGGCACGACAGCAGGTGTGCGGCTGAGAGCCGCAGAGGGGAAGTTGCGTCCATTTTACCGGAAACGGAGCAAAAAACCGCCATTCATACCCTGTATCAATCAGTGTCATCCTGTTTAATAGTCGTTTCCGCTCATATGGTGCACAAGGGGTGTTGAAGAAACATCCGTTTTGTGGTGCTTTTTTAGTCTTTTGGGGATTTAAATTCCTATCGAT

>IS91-V2

CGAGTAGGCAGCCTGGCGGCTGCGGCTTGTCATGGTCTGGAATTACCGTTATAAAAAAAGATAATGTCATTGTCTTTCAGGTAGTTATATGGCCCGTTCAGCTAAACCCCGTAAACGCAAACCCGCACCACAAAGAAGCAAACTTCCCCGCTATGTTGTGAAACTTCATCCGGATGATTTTTTTGACGAAGAAGACGCTGAAGTTCTGCGCTTTGATAATTTTGACGATGCCGTTGAGTGCTGCGCTGACCTGGGTATTCCGTTCTTTCTGGATGCAGGAAACAAAAAGCTGGTCTTCTGGTTTGTTCGTGTCGATGACGAAGGGTATCCGGAAATAGCCCGCTGTACGGAGCGGGAGTTTGCAACCATTCTTGCCGGTATCAGTGCCGGTGGTATGTACTGCCCGGAATGCGGCACAGTTCACTGGCCGGATGGCGTTACCCCCCCGTCTGATGCTTCCCCGTTTTGCCGATATTTTTCAGCAGGGTAACCGCTGGCTTAACTGGCTGGAGAAACAGCCGGAAGGTTCAGTGCGTCCGGTGGTGACTGAGTCAGTGACAAAAATCATGGCATGCGGGACCACGCTGATGGGCTACACGCAATGGTGCTGTTCGTCACCGGACTGTTGCCACACCAAAAAGGTCTGCTTCCGGTGTAAAAGCCGCTCCTGTCCGCACTGCGGGGTGAAGGCTGGCGCACAGTGGATACAGTATCTGCTGAGCCTGGTCCCCGACTGCCCGTGGCAGCATATTGTGTTCACACTTCCCTGCCAGTACTGGTCCCTGGTGTTCCACAACCGGTGGTTACTGGCAGAGATGAGCCGCATTGCAGCGGATGTGATACTGGAAATCTGCCATCAGACAGATGTGGAGCCGGGGATATTCACGGTGATCCACACATGGGGGCGTGACCAGCAGTGGCATCCGCATATCCATTTATCGACAACTGCCGGTGGTGTGACGTCGGGCCACACCTGGAAAAATCTTCATTTTTACGCCCGTAAGGTGATGAGCATGTGGCGTTACCGGATAACGCGGCTACTGTCCCGGAAATACCCGGAGCTGGTGATACCGGATGAACTGGCAGTGGAAGGAAACAGCAAACGGGACTGGAATTGCTTCCTGGACACGCATTACCGCCGCGGCTGGAATGTCAACATATCCAGGGTGATGGATAACGCCACACATGTGGCGGTGTACTTCGGCTCTTACCTGAAAAAGCCACCGGTGCCGATGAGTCGGCTGGAGCATTATGCCGGTCAGGATGAAATCGGTCTGCGTTACAACAGTCACCGTACAAAACGGGAAGAATACCTGTTGATGAGTGGAGATGAGTTCATGGAAAGGTTCTCCTGGCATGTAGCAGATAAGGGGTTCCGTATGGTGAGGTACTACGGTTTCCTGAGTCCGGTGAAGCGCCGGTTACTGGAAGAAGTTGTGTACGTCATAACGGAGACGGTGAGAAAAACGGCGATGCAAATCAGGTGGAGAGGGATGTATCAGCGGTTACTGAAGGTTGACCCGCTGAAGTGCATTCTGTGCGGAAGTCAGATGCGTTTTACGGGGCTGAAGCGGGGTTACCGACTGGCAGAGCTGGTCCTGATGCATGAGCGACTGGCACGACAGCAGGTGTGCGGCTGAGAGCCGCAGAGGGGAAGTTGCGTCCATTTTACCGGAAACGGAGCAAAAAACCGCCATTCATACCCTGTATCAATCAGTGTCATCCTGTTTAATAGTCGTTTCCGCTCATATGGTGCACAAGGGGTGTTGAAGAAACATCCGTTTTGTGGTGCTTTTTTAGTCTTTTGGGGATTTAAATTCCTATCGAT

>IS91-V3

CGAGTAGGCAGCCTGGCGGCTGCGGCTTGTCATGGCCTGAAATTACCGTTATAAAAACAGACAATATCATTGTCTTTCAGGTAGTTATATGTCCCGTTCAGCTAAACCCCGTAAACGAAAACCTGCCCCTCAAAGAAGCAAACTTCCCCGCTATGTCGTGAAGCTTCACGACGATGACTTCTTTGACGAAGAAGACGCAGAAGCTCTGCGCTTTGATAATTTTGACGATGCCGTTGAGTGCTGCGCAGACCTGAATATTCCCTTCTTTGTGGATGCCGGAAACAAAAAGCTGGTCTTCTGGTTTGTTCGTGTCGATGACGAAGGGTATCCTGAAATAGCCCGCTGCACGGAGCGGGAGTTTGCGACCATTCTTGCCGGTATCAGCGCCGGCGGCATGTACTGCCCGGAGTGTGGCACGGTTCACTGGCCGGACGGAGTCCCCCCGCCTTCTGATGCTTCCCCGTTTTGCCGACATTTTTCAGCAGGGTAACCGCTGGCTTAACTGGCTGGAGAAACAGCCGGAAGGTTCAGTGCGTCCGGTGGTGACTGAGTCAGTGACAAAAATCATGGCATGCGGGACCACGCTGATGGGCTACACGCAATGGTGCTGTTCGTCACCGGACTGTTGCCACACCAAAAAGGTCTGCTTCCGGTGTAAAAGCCGCTCCTGTCCGCACTGCGGGGTGAAGGCTGGCGCACAGTGGATACAGTATCTGCTGAGCCTGGTCCCCGACTGCCCGTGGCAGCATATTGTGTTCACACTTCCCTGCCAGTACTGGTCCCTGGTGTTCCACAACCGGTGGTTACTGGCAGAGATGAGCCGCATTGCAGCGGATGTGATACTGGAAATCTGCCATCAGACAGATGTGGAGCCGGGGATATTCACGGTGATCCACACATGGGGGCGTGACCAGCAGTGGCATCCGCATATCCATTTATCGACAACTGCCGGTGGTGTGACGTCGGGCCACACCTGGAAAAATCTTCATTTTTACGCCCGTAAGGTGATGAGCATGTGGCGTTACCGGATAACGCGGCTACTGTCCCGGAAATACCCGGAGCTGGTGATACCGGATGAACTGGCAGTGGAAGGAAACAGCAAACGGGACTGGAATTGCTTCCTGGACACGCATTACCGCCGCGGCTGGAATGTCAACATATCCAGGGTGATGGATAACGCCACACATGTGGCGGTGTACTTCGGCTCTTACCTGAAAAAGCCACCGGTGCCGATGAGTCGGCTGGAGCATTATGCCGGTCAGGATGAAATCGGTCTGCGTTACAACAGTCACCGTACAAAACGGGAAGAATACCTGTTGATGAGTGGAGATGAGTTCATGGAAAGGTTCTCCTGGCATGTAGCAGATAAGGGGTTCCGTATGGTGAGGTACTACGGTTTCCTGAGTCCGGTGAAGCGCCGGTTACTGGAAGAAGTTGTGTACGTCATAACGGAGACGGTGAGAAAAACGGCGATGCAAATCAGGTGGAGAGGGATGTATCAGAGGTTACTGAAGGTTGACCCGCTGAAGTGCATTCTGTGCGGAAGTCAGATGCGTTTTACGGGGCTGAAGCGGGGTTACCGACTGGCAGAGCTGGTCCTGATGCATGAGCGACTTGGCACGACAGCAGGTGTGCGGCTGAGAGCCGCAGAGGGGAAGTTGCGTCCATTTTACCGGAAACGGAGCAAAAAACCGCCATTCATACCCTGTATCAATCAGTGTCATCCTGTTTAATAGTCGTTTCCGCTCATATGGTGCACAAGGGGTGTTGAAGAAACATCCGTTTTGTGGTGCTTTTTTAGTCTTTTGGGGATTTAAATTCCTATCGAT

>IS91-V4

CGAGTAGGCAGCCTGGCGGCTGCGGCTTGTCATGGTCTGGAATTACCGTTATAAAAAAAGATAATGTCATTGTCTTTCAGGTAGTTATATGGCCCGTTCAGCTAAACCCCGTAAACGCAAACCCGCACCACAAAGAAGCAAACTTCCCCGCTATGTTGTGAAACTTCATCCGGATGATTTTTTTGACGAAGAAGACGCTGAAGTTCTGCGCTTTGATAATTTTGACGATGCCGTTGAGTGCTGCGCTGACCTGGGTATTCCGTTCTTTCTGGATGCAGGAAACAAAAAGCTGGTCTTCTGGTTTGTTCGTGTCGATGACGAAGGGTATCCGGAAATAGCCCGCTGTACGGAGCGGGAGTTTGCAACCATTCTTGCCGGTATCAGTGCCGGTGGTATGTACTGCCCGGAATGCGGCACAGTTCACTGGCCGGATGGCGTTACCCCCCCGTCTGATGCTTCCCCGTTTTGCCGATATTTTTCAGCAGGGTAACCGCTGGCTTAACTGGCTGGAGAAACAGCCGGAAGGTTCAGTGCGTCCGGTGGTGACTGAGTCAGTGACAAAAATCATGGCATGCGGGACCACGCTGATGGGCTACACGCAATGGTGCTGTTCGTCACCGGACTGTTGCCACACCAAAAAGGTCTGCTTCCGGTGTAAAAGCCGCTCCTGTCCGCACTGCGGGGTGAAGGCTGGCGCACAGTGGATACAGTATCTGCTGAGCCTGGTCCCCGACTGCCCGTGGCAGCATATTGTGTTCACACTTCCCTGCCAGTACTGGTCCCTGGTGTTCCACAACCGGTGGTTACTGGCAGAGATGAGCCGCATTGCAGCGGATGTGATACTGGAAATCTGCCATCAGACAGATGTGGAGCCGGGGATATTCACGGTGATCCACACATGGGGGCGTGACCAGCAGTGGCATCCGCATATCCATTTATCGACAACTGCCGGTGGTGTGACGTCGGGCCACACCTGGAAAAATCTTCATTTTTACGCCCGTAAGGTGATGAGCATGTGGCGTTACCGGATAACGCGGCTACTGTCCCGGAAATACCCGGAGCTGGTAATACCGGATGAACTGGCAGTGGAAGGAAACAGCAAACGGGACTGGAATTGCTTCCTGGACACGCATTACCGCCGCGGCTGGAATGTCAACATATCCAGGGTGATGGATAACGCCACACATGTGGCGGTGTACTTCGGCTCTTACCTGAAAAAGCCACCGGTGCCGATGAGTCGGCTGGAGCATTATGCCGGTCAGGATGAAATCGGTCTGCGTTACAACAGTCACCGTACAAAACGGGAAGAATACCTGTTGATGAGTGGAGATGAGTTCATGGAAAGGTTCTCCTGGCATGTAGCAGATAAGGGGTTCCGTATGGTGAGGTACTACGGTTTCCTGAGTCCGGTGAAGCGCCGCTTACTGGAAGAAGTTGTGTACGTCATAACGGAGACGGTGAGAAAAACGGCGATGCAAATCAGGTGGAGAGGGATGTATCAGCGGTTACTGAAGGTTGACCCGCTGAAGTGCATTCTGTGCGGAAGTCAGATGCGTTTTACGGGGCTGAAGCGGGGTTACCGACTGGCAGAGCTGGTCCTGATGCATGAGCGACTGGCACGACAGCAGGTGTGCGGCTGAGAGCCGCAGAGGGGAAGTTGCGTCCATTTTACCGGAAACGGAGCAAAAAACCGCCATTCATACCCTGTATCAATCAGTGTCATCCTGTTTAATAGTCGTTTCCGCTCATATGGTGCACAAGGGGTGTTGAAGAAACATCCGTTTTGTGGTGCTTTTTTAGTCTTTTGGGGATTTAAATTCCTATCGAT

>IS91-V5

CGAGTAGGCAGCCTGGCGGCTGCGGCTTGTCATGGTCTGGAATTACCGTTATAAAAAAAGATAATGTCATTGTCTTTCAGGTAGTTATATGGCCCGTTCAGCTAAACCCCGTAAACGCAAACCCGCACCACAAAGAAGCAAACTTCCCCGCTATGTTGTGAAACTTCATCCGGATGATTTTTTTGACGAAGAAGACGCTGAAGTTCTGCGCTTTGATAATTTTGACGATGCCGTTGAGTGCTGCGCTGACCTGGGTATTCCGTTCTTTCTGGATGCAGGAAACAAAAAGCTGGTCTTCTGGTTTGTTCGTGTCGATGACGAAGGGTATCCGGAAATAGCCCGCTGTACGGAGCGGGAGTTTGCGACCATTCTTGCCGGTATCAGTGCCGGTGGTATGTACTGCCCGGAATGCGGCACAGTTCACTGGCCGGATGGCGTTACCCCACCCGTCTGATGCTTCCCCGTTTTGCCGATATTTTTCAGCAGGGTAACCGCTGGCTTAACTGGCTGGAGAAACAGCCGGAAGGTTCAGTGCGTCCGGTGGTGACTGAGTCAGTGACAAAAATCATGGCATGCGGGACCACGCTGATGGGCTACACGCAATGGTGCTGTTCGTCACCGGACTGTTGCCACACCAAAAAGGTCTGCTTCCGGTGTAAAAGCCGCTCCTGTCCGCACTGCGGGGTGAAGGCTGGCGCACAGTGGATACAGTATCTGCTGAGCCTGGTCCCCGACTGCCCGTGGCAGCATATTGTGTTCACACTTCCCTGCCAGTACTGGTCCCTGGTGTTCCACAACCGGTGGTTACTGGCAGAGATGAGCCGCATTGCAGCGGATGTGATACTGGAAATCTGCCATCAGACAGATGTGGAGCCGGGGATATTCACGGTGATCCACACATGGGGGCGTGACCAGCAGTGGCATCCGCATATCCATTTATCGACAACTGCCGGTGGTGTGACGTCGGGCCACACCTGGAAAAATCTTCATTTTTACGCCCGTAAGGTGATGAGCATGTGGCGTTACCGGATAACGCGGCTACTGTCCCGGAAATACCCGGAGCTGGTAATACCGGATGAACTGGCAGTGGAAGGAAACAGCAAACGGGACTGGAATTGCTTCCTGGACACGCATTACCGCCGCGGCTGGAATGTCAACATATCCAGGGTGATGGATAACGCCACACATGTGGCGGTGTACTTCGGCTCTTACCTGAAAAAGCCACCGGTGCCGATGAGTCGGCTGGAGCATTATGCCGGTCAGGATGAAATCGGTCTGCGTTACAACAGTCACCGTACAAAACGGGAAGAATACCTGTTGATGAGTGGAGATGAGTTCATGGAAAGGTTCTCCTGGCATGTAGCAGATAAGGGGTTCCGTATGGTGAGGTACTACGGTTTCCTGAGTCCGGTGAAGCGCCGCTTACTGGAAGAAGTTGTGTACGTCATAACGGAGACGGTGAGAAAAACGGCGATGCAAATCAGGTGGAGAGGGATGTATCAGAGGTTACTGAAGGTTGACCCGCTGAAGTGCATTCTGTGCGGAAGTCAGATGCGTTTTACGGGGCTGAAGCGGGGTTACCGACTGGCAGAGCTGGTCCTGATGCATGAGCGACTGGCACGACAGCAGGTGTGCGGCTGAGAGCCGCAGAGGGGAAGTTGCGTCCATTTTACCGGAAACGGAGCAAAAAACCGCCATTCATACCCTGTATCAATCAGTGTCATCCTGTTTAATAGTCGTTTCCGCTCATATGGTGCACAAGGGGTGTTGAAGAAACATCCGTTTTGTGGTGCTTTTTTAGTCTTTTGGGGATTTAAATTCCTATCGAT

>IS91-V6

CGAGTAGGCAGCCTGGCGGCTGCGGCTTGTCATGGTCTGGAATTACCGTTATAAAAAAGATAATGTCATTGTCTTTCAGGTAGTTATATGGCCCGTTCAGCTAAACCCCGTAAACGCAAACCCGCACCACAAAGAAGCAAACTTCCCCGCTATGTTGTGAAACTTCATCCGGATGATTTTTTGACGAAGAAGACGCTGAAGTTCTGCGCTTTGATAATTTTGACGATGCCGTTGAGTGCTGCGCTGACCTGGGTATTCCGTTCTTTCTGGATGCAGGAAACAAAAGCTGGTCTTCTGGTTTGTTCGTGTCGATGACGAAGGGTATCCGGAAATAGCCCGCTGCACGGAGCGGGAGTTTGCAACCATTCTTGCCGGTATCAGCGCCGGCGGCATGTACTGCCCGGAATGCGGCACAGTTCACTGGCCGGACGGAGTACCCCACCCGTCTGATGCTTCCCCGTTTTGCCGATATTTTTCAGCAGGGTAACCGCTGGCTTAACTGGCTGGAGAAACAGCCGGAAGGTTCAGTGCGTCCGGTGGTGACTGAGTCAGTGACAAAAATCATGGCATGCGGGACCACGCTGATGGGCTACACGCAATGGTGCTGTTCGTCACCGGACTGTTGCCACACCAAAAAGGTCTGCTTCCGGTGTAAAAGCCGCTCCTGTCCGCACTGCGGGGTGAAGGCTGGCGCACAGTGGATACAGTATCTGCTGAGCCTGGTCCCCGACTGCCCGTGGCAGCATATTGTGTTCACACTTCCCTGCCAGTACTGGTCCCTGGTGTTCCACAACCGGTGGTTACTGGCAGAGATGAGCCGCATTGCAGCGGATGTGATACTGGAAATCTGCCATCAGACAGATGTGGAGCCGGGGATATTCACGGTGATCCACACATGGGGGCGTGACCAGCAGTGGCATCCGCATATCCATTTATCGACAACTGCCGGTGGTGTGACGTCGGGCCACACCTGGAAAAATCTTCATTTTTACGCCCGTAAGGTGATGAGCATGTGGCGTTACCGGATAACGCGGCTACTGTCCCGGAAATACCCGGAGCTGGTAATACCGGATGAACTGGCAGTGGAAGGAAACAGCAAACGGGACTGGAATCGCTTCCTGGACACGCATTACCGCCGCGGCTGGAATGTCAACATATCCAGGGTGATGGATAACGCCACACATGTGGCGGTGTACTTCGGCTCTTACCTGAAAAAGCCACCGGTGCCGATGAGTCGGCTGGAGCATTATGCCGGTCAGGATGAAATCGGTCTGCGTTACAACAGTCACCGTACAAAACGGGAAGAATACCTGTTGATGAGTGGAGATGAGTTCATGGAAAGGTTCTCCTGGCATGTAGCAGATAAGGGGTTCCGTATGGTGAGGTACTACGGTTTCCTGAGTCCGGTGAAGCGCCGCTTACTGGAAGAAGTTGTGTACGTCATAACGGAGACGGTGAGAAAAACGGCGATGCAAATCAGGTGGAGAGGGATGTATCAGAGGTTACTGAAGGTTGACCCGCTGAAGTGCATTCTGTGCGGAAGTCAGATGCGTTTTACGGGGCTGAAGCGGGGTTACCGACTGGCAGAGCTGGTCCTGATGCATGAGCGACTGGCACGACAGCAGGTGTGCGGCTGAGAGCCGCAGAGGGGAAGTTGCGTCCATTTTACCGGAAACGGAGCAAAAAACCGCCATTCATACCCTGTATCAATCAGTGTCATCCTGTTTAATTGTCGTTTCCGCTCATATGGTGCACAAGGGGTGTTGAAGAAACATCCGTTTTGTGGTGCTTTTTTAGTCTTTTGGGGATTTAAATTCCTATCGAT

>IS91-V7

CGAGTAGGCAGCCTGGCGGCTGCGGCTTGTCATGGTCTGGAATTACCGTTATAAAAAAAGATAATGTCATTGTCTTTCAGGTAGTTATATGGCCCGTTCAGCTAAACCCCGTAAACGCAAACCCGCACCACAAAGAAGCAAACTTCCCCGCTATGTTGTGAAACTTCATCCGGATGATTTTTTTGACGAAGAAGACGCTGAAGTTCTGCGCTTTGATAATTTTGACGATGCCGTTGAGTGCTGCGCTGACCTGGGTATTCCGTTCTTTCTGGATGCAGGAAACAAAAAGCTGGTCTTCTGGTTTGTTCGTGTCGATGACGAAGGGTATCCTGAAATAGCCCGCTGTACGGAGCGGGAGTTTGCAACCATTCTTGCCGGTATCAGTGCCGGTGGTATGTACTGCCCGGAATGCGGCACAGTTCACTGGCCGGATGGCGTTACCCCACCCGTCTGATGCTTCCCCGTTTTGCCGATATTTTTCAGCAGGGTAACCGCTGGCTTAACTGGCTGGAGAAACAGCCGGAAGGTTCAGTGCGTCCGGTGGTGACTGAGTCAGTGACAAAAATCATGGCATGCGGGACCACGCTGATGGGCTACACGCAATGGTGCTGTTCGTCACCGGACTGTTGCCACACCAAAAAGGTCTGCTTCCGGTGTAAAAGCCGCTCCTGTCCGCACTGCGGGGTGAAGGCTGGCGCACAGTGGATACAGTATCTGCTGAGCCTGGTCCCCGACTGCCCGTGGCAGCATATTGTGTTCACACTTCCCTGCCAGTACTGGTCCCTGGTGTTCCACAACCGGTGGTTACTGGCAGAGATGAGCCGCATTGCAGCGGATGTGATACTGGAAATCTGCCATCAGACAGATGTGGAGCCGGGGATATTCACGGTGATCCACACATGGGGGCGTGACCAGCAGTGGCATCCGCATATCCATTTATCGACAACTGCCGGTGGTGTGACGTCGGGCCACACCTGGAAAAATCTTCATTTTTACGCCCGTAAGGTGATGAGCATGTGGCGTTACCGGATAACGCGGCTACTGTCCCGGAAATACCCGGAGCTGGTAATACCGGATGAACTGGCAGTGGAAGGAAACAGCAAACGGGACTGGAATCGCTTCCTGGACACGCATTACCGCCGCGGCTGGAATGTCAACATATCCAGGGTGATGGATAACGCCACACATGTGGCGGTGTACTTCGGCTCTTACCTGAAAAAGCCACCGGTGCCGATGAGTCGGCTGGAGCATTATGCCGGTCAGGATGAAATCGGTCTGCGTTACAACAGTCACCGTACAAAACGGGAAGAATACCTGTTGATGAGTGGAGATGAGTTCATGGAAAGGTTCTCCTGGCATGTAGCAGATAAGGGGTTCCGTATGGTGAGGTACTACGGTTTCCTGAGTCCGGTGAAGCGCCGCTTACTGGAAGAAGTTGTGTACGTCATAACGGAGACGGTGAGAAAAACGGCGATGCAAATCAGGTGGAGAGGGATGTATCAGAGGTTACTGAAGGTTGACCCGCTGAAGTGCATTCTGTGCGGAAGTCAGATGCGTTTTACGGGGCTGAAGCGGGGTTACCGACTGGCAGAGCTGGTCCTGATGCATGAGCGACTGGCACGACAGCAGGTGTGCGGCTGAGAGCCGCAGAGGGGAAGTTGCGTCCATTTTACCGGAAACGGAGCAAAAAACCGCCATTCATACCCTGTATCAATCAGTGTCATCCTGTTTAATAGTCGTTTCCGCTCATATGGTGCACAAGGGGTGTTGAAGAAACATCCGTTTTGTGGTGCTTTTTTAGTCTTTTGGGGATTTAAATTCCTATCGAT

>IS91-V8

CGAGTAGGCAGCCTGGCGGCTGCGGCTTGTCATGGTCTGGAATTACCGTTATAAAAAAAGATAATGTCATTGTCTTTCAGGTAGTTATATGGCCCGTTCAGCTAAACCCCGTAAACGCAAACCCGCACCACAAAGAAGCAAACTTCCCCGCTATGTTGTGAAACTTCATCCGGATGATTTTTTTGACGAAGAAGACGCTGAAGTTCTGCGCTTTGATAATTTTGACGATGCCGTTGAGTGCTGCGCTGACCTGGGTATTCCGTTCTTTCTGGATGCAGGAAACAAAAAGCTGGTCTTCTGGTTTGTTCGTGTCGATGACGAAGGGTATCCGGAAATAGCCCGCTGTACGGAGCGGGAGTTTGCAACCATTCTTGCCGGTATCAGTGCCGGTGGTATGTACTGCCCGGAATGCGGCACAGTTCACTGGCCGGATGGCGTTACCCCCCCGTCTGATGCTTCCCCGTTTTGCCGATATTTTTCAGCAGGGTAACCGCTGGCTTAACTGGCTGGAGAAACAGCCGGAAGGTTCAGTGCGTCCGGTGGTGACTGAGTCAGTGACAAAAATCATGGCATGCGGGACCACGCTGATGGGCTACACGCAATGGTGCTGTTCGTCACCGGACTGTTGCCACACCAAAAAGGTCTGCTTCCGGTGTAAAAGCCGCTCCTGTCCGCACTGCGGGGTGAAGGCTGGCGCACAGTGGATACAGTATCTGCTGAGCCTGGTCCCCGACTGCCCGTGGCAGCATATTGTGTTCACACTTCCCTGCCAGTACTGGTCCCTGGTGTTCCACAACCGGTGGTTACTGGCAGAGATGAGCCGCATTGCAGCGGATGTGATACTGGAAATCTGCCATCAGACAGATGTGGAGCCGGGGATATTCACGGTGATCCACACATGGGGGCGTGACCAGCAGTGGCATCCGCATATCCATTTATCGACAACTGCCGGTGGTGTGACGTCGGGCCACACCTGGAAAAATCTTCATTTTTACGCCCGTAAGGTGATGAGCATGTGGCGTTACCGGATAACGCGGCTACTGTCCCGGAAATACCCGGAGCTGGTAATACCGGATGAACTGGCAGTGGAAGGAAACAGCAAACGGGACTGGAATTGCTTCCTGGACACGCATTACCGCCGCGGCTGGAATGTCAACATATCCAGGGTGATGGATAACGCCACACATGTGGCGGTGTACTTCGGCTCTTACCTGAAAAAGCCACCGGTGCCGATGAGTCGGCTGGAGCATTATGCCGGTCAGGATGAAATCGGTCTGCGTTACAACAGTCACCGTACAAAACGGGAAGAATACCTGTTGATGAGTGGAGATGAGTTCATGGAAAGGTTCTCCTGGCATGTAGCAGATAAGGGGTTCCGTATGGTGAGGTACTACGGTTTCCTGAGTCCGGTGAAGCGCCGCTTACTGGAAGAAGTTGTGTACGTCATAACGGAGACGGTGAGAAAAACGGCGATGCAAATCAGGTGGAGAGGGATGTATCAGAGGTTACTGAAGGTTGACCCGCTGAAGTGCATTCTGTGCGGAAGTCAGATGCGTTTTACGGGGCTGAAGCGGGGTTACCGACTGGCAGAGCTGGTCCTGATGCATGAGCGACTGGCACGACAGCAGGTGTGCGGCTGAGAGCCGCAGAGGGGAAGTTGCGTCCATTTTACCGGAAACGGAGCAAAAAACCGCCATTCATACCCTGTATCAATCAGTGTCATCCTGTTTAATAGTCGTTTCCGCTCATATGGTGCACAAGGGGTGTTGAAGAAACATCCGTTTTGTGGTGCTTTTTTAGTCTTTTGGGGATTTAAATTCCTATCGAT

>IS91-V9

CGAGTAGGCAGCCTGGCGGCTGCGGCTTGTCATGGTCTGGAATTACCGTTATAAAAAAAGATAATGTCATTGTCTTTCAGGTAGTTATATGGCCCGTTCAGCTAAACCCCGTAAACGCAAACCCGCACCACAAAGAAGCAAACTTCCCCGCTATGTTGTGAAACTTCATCCGGATGATTTTTTTGACGAAGAAGACGCTGAAGTTCTGCGCTTTGATAATTTTGACGATGCCGTTGAGTGCTGCGCTGACCTGGGTATTCCGTTCTTTCTGGATGCAGGAAACAAAAAGCTGGTCTTCTGGTTTGTTCGTGTCGATGACGAAGGGTATCCGGAAATAGCCCGCTGTACGGAGCGGGAGTTTGCGACCATTCTTGCCGGTATCAGTGCCGGTGGTATGTACTGCCCGGAATGCGGCACAGTTCACTGGCCGGATGGCGTTACCCCACCCGTCTGATGCTTCCCCGTTTTGCCGATATTTTTCAGCAGGGTAACCGCTGGCTTAACTGGCTGGAGAAACAGCCGGAAGGTTCAGTGCGTCCGGTGGTGACTGAGTCAGTGACAAAAATCATGGCATGCGGGACCACGCTGATGGGCTACACGCAATGGTGCTGTTCGTCACCGGACTGTTGCCACACCAAAAAGGTCTGCTTCCGGTGTAAAAGCCGCTCCTGTCCGCACTGCGGGGTGAAGGCTGGCGCACAGTGGATACAGTATCTGCTGAGCCTGGTCCCCGACTGCCCGTGGCAGCATATTGTGTTCACACTTCCCTGCCAGTACTGGTCCCTGGTGTTCCACAACCGGTGGTTACTGGCAGAGATGAGCCGCATTGCAGCGGATGTGATACTGGAAATCTGCCATCAGACAGATGTGGAGCCGGGGATATTCACGGTGATCCACACATGGGGGCGTGACCAGCAGTGGCATCCGCATATCCATTTATCGACAACTGCCGGTGGTGTGACGTCGGGCCACACCTGGAAAAATCTTCATTTTTACGCGCGTAAGGTGATGAGCATGTGGCGTTACCGGATAACGCGGCTACTGTCCCGGAAATACCCGGAGCTGGTAATACCGGATGAACTGGCAGTGGAAGGAAACAGCAAACGGGACTGGAATCGCTTCCTGGACACGCATTACCGCCGCGGCTGGAATGTCAACATATCCAGGGTGATGGATAACGCCACACATGTGGCGGTGTACTTCGGCTCTTACCTGAAAAAGCCACCGGTGCCGATGAGTCGGCTGGAGCATTATGCCGGTCAGGATGAAATCGGTCTGCGTTACAACAGTCACCGTACAAAACGGGAAGAATACCTGTTGATGAGTGGAGATGAGTTCATGGAAAGGTTCTCCTGGCATGTAGCAGATAAGGGGTTCCGTATGGTGAGGTACTACGGTTTCCTGAGTCCGGTGAAGCGCCGCTTACTGGAAGAAGTTGTGTACGTCATAACGGAGACGGTGAGAAAAACGGCGATGCAAATCAGGTGGAGAGGGATGTATCAGAGGTTACTGAAGGTTGACCCGCTGAAGTGCATTCTGTGCGGAAGTCAGATGCGTTTTACGGGGCTGAAGCGGGGTTACCGACTGGCAGAGCTGGTCCTGATGCATGAGCGACTGGCACGACAGCAGGTGTGCGGCTGAGAGCCGCAGAGGGGAAGTTGCGTCCATTTTACCGGAAACGGAGCAAAAAACCGCCATTCATACCCTGTATCAATCAGTGTCATCCTGTTTAATAGTCGTTTCCGCTCATATGGTGCACAAGGGGTGTTGAAGAAACATCCGTTTTGTGGTGCTTTTTTAGTCTTTTGGGGATTTAAATTCCTATCGAT

>IS91-V10

CGAGTAGGCAGCCTGGCGGCTGCGGCTTGTCATGGTCTGGAATTACCGTTATAAAAAAAGATAATGTCATTGTCTTTCAGGTAGTTATATGGCCCGTTCAGCTAAACCCCGTAAACGCAAACCCGCACCACAAAGAAGCAAACTTCCCCGCTATGTTGTGAAACTTCATCCGGATGATTTTTTTGACGAAGAAGACGCTGAAGTTCTGCGCTTTGATAATTTTGACGATGCCGTTGAGTGCTGCGCTGACCTGGGTATTCCGTTCTTTCTGGATGCAGGAAACAAAAAGCTGGTCTTCTGGTTTGTTCGTGTCGATGACGAAGGGTATCCGGAAATAGCCCGCTGTACGGAGCGGGAGTTTGCAACCATTCTTGCCGGTATCAGTGCCGGTGGTATGTACTGCCCGGAATGCGGCACAGTTCACTGGCCGGATGGCGTTACCCCACCCGTCTGATGCTTCCCCGTTTTGCCGATATTTTTCAGCAGGGTAACCGCTGGCTTAACTGGCTGGAGAAACAGCCGGAAGGTTCAGTGCGTCCGGTGGTGACTGAGTCAGTGACAAAAATCATGGCATGCGGGACCACGCTGATGGGCTACACGCAATGGTGCTGTTCGTCACCGGACTGTTGCCACACCAAAAAGGTCTGCTTCCGGTGTAAAAGCCGCTCCTGTCCGCACTGCGGGGTGAAGGCTGGCGCACAGTGGATACAGTATCTGCTGAGCCTGGTCCCCGACTGCCCGTGGCAGCATATTGTGTTCACACTTCCCTGCCAGTACTGGTCCCTGGTGTTCCACAACCGGTGGTTACTGGCAGAGATGAGCCGCATTGCAGCGGATGTGATACTGGAAATCTGCCATCAGACAGATGTGGAGCCGGGGATATTCACGGTGATCCACACATGGGGGCGTGACCAGCAGTGGCATCCGCATATCCATTTATCGACAACTGCCGGTGGTGTGACGTCGGGCCACACCTGGAAAAATCTTCATTTTTACGCCCGTAAGGTGATGAGCATGTGGCGTTACCGGATAACGCGGCTACTGTCCCGGAAATACCCGGAGCTGGTAATACCGGATGAACTGGCAGTGGAAGGAAACAGCAAACGGGACTGGAATCGCTTCCTGGACACGCATTACCGCCGCGGCTGGAATGTCAACATATCCAGGGTGATGGATAACGCCACACATGTGGCGGTGTACTTCGGCTCTTACCTGAAAAAGCCACCGGTGCCGATGAGTCGGCTGGAGCATTATGCCGGTCAGGATGAAATCGGTCTGCGTTACAACAGTCACCGTACAAAACGGGAAGAATACCTGTTGATGAGTGGAGATGAGTTCATGGAAAGGTTCTCCTGGCATGTAGCAGATAAGGGGTTCCGTATGGTGAGGTACTACGGTTTCCTGAGTCCGGTGAAGCGCCGCTTACTGGAAGAAGTTGTGTACGTCATAACGGAGACGGTGAGAAAAACGGCGATGCAAATCAGGTGGAGAGGGATGTATCAGAGGTTACTGAAGGTTGACCCGCTGAAGTGCATTCTGTGCGGAAGTCAGATGCGTTTTACGGGGCTGAAGCGGGGTTACCGACTGGCAGAGCTGGTCCTGATGCATGAGCGACTGGCACGACAGCAGGTGTGCGGCTGAGAGCCGCAGAGGGGAAGTTGCGTCCATTTTACCGGAAACGGAGCAAAAAACCGCCATTCATACCCTGTATCAATCAGTGTCATCCTGTTTAATAGTCGTTTCCGCTCATATGGTGCACAAGGGGTGTTGAAGAAACATCCGTTTTGTGGTGCTTTTTTAGTCTTTTGGGGATTTAAATTCCTATCGAT

>IS91-V11

CGAGTAGGCAGCCTGGCGGCTGCGGCTTGTCATGGTCTGGAATTACCGTTATAAAAAAAGATAATGTCATTGTCTTTCAGGTAGTTATATGGCCCGTTCAGCTAAACCCCGTAAACGCAAACCCGCACCACAAAGAAGCAAACTTCCCCGCTATGTTGTGAAACTTCATCCGGATGATTTTTTTGACGAAGAAGACGCTGAAGTTCTGCGCTTTGATAATTTTGACGATGCCGTTGAGTGCTGCGCTGACCTGGGTATTCCGTTCTTTCTGGATGCAGGAAACAAAAAGCTGGTCTTCTGGTTTGTTCGTGTCGATGACGAAGGGTATCCGGAAATAGCCCGCTGTACGGAGCGGGAGTTTGCAACCATTCTTGCCGGTATCAGTGCCGGTGGTATGTACTGCCCGGAATGCGGCACAGTTCACTGGCCGGATGGCGTTACCCCACCCGTCTGATGCTTCCCCGTTTTGCCGATATTTTTCAGCAGGGTAACCGCTGGCTTAACTGGCTGGAGAAACAGCCGGAAGGTTCAGTGCGTCCGGTGGTGACTGAGTCAGTGACAAAAATCATGGCATGCGGGACCACGCTGATGGGCTACACGCAATGGTGCTGTTCGTCACCGGACTGTTGCCACACCAAAAAGGTCTGCTTCCGGTGTAAAAGCCGCTCCTGTCCGCACTGCGGGGTGAAGGCTGGCGCACAGTGGATACAGTATCTGCTGAGCCTGGTCCCCGACTGCCCGTGGCAGCATATTGTGTTCACACTTCCCTGCCAGTACTGGTCCCTGGTGTTCCACAACCGGTGGTTACTGGCAGAGATGAGCCGCATTGCAGCGGATGTGATACTGGAAATCTGCCATCAGACAGATGTGGAGCCGGGGATATTCACGGTGATCCACACATGGGGGCGTGACCAGCAGTGGCATCCGCATATCCATTTATCGACAACTGCCGGTGGTGTGACGTCGGGCCACACCTGGAAAAATCTTCATTTTTACGCCCGTAAGGTGATGAGCATGTGGCGTTACCGGATAACGCGGCTACTGTCCCGGAAATACCCGGAGCTGGTAATACCGGATGAACTGGCAGTGGAAGGAAACAGCAAACGGGACTGGAATTGCTTCCTGGACACGCATTACCGCCGCGGCTGGAATGTCAACATATCCAGGGGGATGGATAACGCCACACATGTGGCGGTGTACTTCGGCTCTTACCTGAAAAAGCCACCGGTGCCGATGAGTCGGCTGGAGCATTATGCCGGTCAGGATGAAATCGGTCTGCGTTACAACAGTCACCGTACAAAACGGGAAGAATACCTGTTGATGAGTGGAGATGAGTTCATGGAAAGGTTCTCCTGGCATGTAGCAGATAAGGGGTTCCGTATGGTGAGGTACTACGGTTTCCTGAGTCCGGTGAAGCGCCGCTTACTGGAAGAAGTTGTGTACGTCATAACGGAGACGGTGAGAAAAACGGCGATGCAAATCAGGTGGAGAGGGATGTATCAGAGGTTACTGAAGGTTGACCCGCTGAAGTGCATTCTGTGCGGAAGTCAGATGCGTTTTACGGGGCTGAAGCGGGGTTACCGACTGGCAGAGCTGGTCCTGATGCATGAGCGACTGGCACGACAGCAGGTGTGCGGCTGAGAGCCGCAGAGGGGAAGTTGCGTCCATTTTACCGGAAACGGAGCAAAAAACCGCCATTCATACCCTGTATCAATCAGTGTCATCCTGTTTAATAGTCGTTTCCGCTCATATGGTGCACAAGGGGTGTTGAAGAAACATCCGTTTTGTGGTGCTTTTTTAGTCTTTTGGGGATTTAAATTCCTATCGAT

>IS91-V12

CGAGTAGGCAGCCTGGCGGCTGCGGCTTGTCATGGTCTGGAATTACCGTTATAAAAAAAGATAATGTCATTGTCTTTCAGGTAGTTATATGGCCCGTTCAGCTAAACCCCGTAAACGCAAACCCGCACCACAAAGAAGCAAACTTCCCCGCTATGTTGTGAAACTTCATCCGGATGATTTTTTTGACGAAGAAGACGCTGAAGTTCTGCGCTTTGATAATTTTGACGATGCCGTTGAGTGCTGCGCTGACCTGGGTATTCCGTTCTTTCTGGATGCAGGAAACAAAAAGCTGGTCTTCTGGTTTGTTCGTGTCGATGACGAAGGGTATCCGGAAATAGCCCGCTGTACGGAGCGGGAGTTTGCAACCATTCTTGCCGGTATCAGTGCCGGTGGTATGTACTGCCCGGAATGCGGCACAGTTCACTGGCCGGATGGCGTTACCCCACCCGTCTGATGCTTCCCCGTTTTGCCGACATTTTTCAGCAGGGTAACCGCTGGCTTAACTGGCTGGAGAAACAGCCGGAAGGTTCAGTGCGTCCGGTGGTGACTGAGTCAGTGACAAAAATCATGGCATGCGGGACCACGCTGATGGGCTACACGCAATGGTGCTGTTCGTCACCGGACTGTTGCCACACCAAAAAGGTCTGCTTCCGGTGTAAAAGCCGCTCCTGTCCGCACTGCGGGGTGAAGGCTGGCGCACAGTGGATACAGTATCTGCTGAGCCTGGTCCCCGACTGCCCGTGGCAGCATATTGTGTTCACACTTCCCTGCCAGTACTGGTCCCTGGTGTTCCACAACCGGTGGTTACTGGCAGAGATGAGCCGCATTGCAGCGGATGTGATACTGGAAATCTGCCATCAGACAGATGTGGAGCCGGGGATATTCACGGTGATCCACACATGGGGGCGTGACCAGCAGTGGCATCCGCATATCCATTTATCGACAACTGCCGGTGGTGTGACGTCGGGCCACACCTGGAAAAATCTTCATTTTTACGCGCGTAAGGTGATGAGCATGTGGCGTTACCGGATAACGCGGCTACTGTCCCGGAAATACCCGGAGCTGGTAATACCGGATGAACTGGCAGTGGAAGGAAACAGCAAACGGGACTGGAATCGCTTCCTGGACACGCATTACCGCCGCGGCTGGAATGTCAACATATCCAGGGTGATGGATAACGCCACACATGTGGCGGTGTACTTCGGCTCTTACCTGAAAAAGCCACCGGTGCCGATGAGTCGGCTGGAGCATTATGCCGGTCAGGATGAAATCGGTCTGCGTTACAACAGTCACCGTACAAAACGGGAAGAATACCTGTTGATGAGTGGAGATGAGTTCATGGAAAGGTTCTCCTGGCATGTAGCAGATAAGGGGTTCCGTATGGTGAGGTACTACGGTTTCCTGAGTCCGGTGAAGCGCCGCTTACTGGAAGAAGTTGTGTACGTCATAACGGAGACGGTGAGAAAAACGGCGATGCAAATCAGGTGGAGAGGGATGTATCAGAGGTTACTGAAGGTTGACCCGCTGAAGTGCATTCTGTGCGGAAGTCAGATGCGTTTTACGGGGCTGAAGCGGGGTTACCGACTGGCAGAGCTGGTCCTGATGCATGAGCGACTGGCACGACAGCAGGTGTGCGGCTGAGAGCCGCAGAGGGGAAGTTGCGTCCATTTTACCGGAAACGGAGCAAAAAACCGCCATTCATACCCTGTATCAATCAGTGTCATCCTGTTTAATAGTCGTTTCCGCTCATATGGTGCACAAGGGGTGTTGAAGAAACATCCGTTTTGTGGTGCTTTTTTAGTCTTTTGGGGATTTAAATTCCTATCGAT

>IS91-V13

CGAGTAGGCAGCCTGGCGGCTGCGGCTTGTCATGGTCTGGAATTACCGTTATAAAAAAAGATAATGTCATTGTCTTTCAGGTAGTTATATGGCCCGTTCAGCTAAACCCCGTAAACGCAAACCCGCACCACAAAGAAGCAAACTTCCCCGCTATGTTGTGAAACTTCATCCGGATGATTTTTTTGACGAAGAAGACGCTGAAGTTCTGCGCTTTGATAATTTTGACGATGCCGTTGAGTGCTGCGCTGACCTGGGTATTCCGTTCTTTCTGGATGCAGGAAACAAAAAGCTGGTCTTCTGGTTTGTTCGTGTCGATGACGAAGGGTATCCGGAAATAGCCCGCTGTACGGAGCGGGAGTTTGCAACCATTCTTGCCGGTATCAGTGCCGGTGGTATGTACTGCCCGGAATGCGGCACAGTTCACTGGCCGGATGGCGTTACCCCCCCGTCTGATGCTTCCCCGTTTTGCCGATATTTTTCAGCAGGGTAACCGCTGGCTTAACTGGCTGGAGAAACAGCCGGAAGGTTCAGTGCGTCCGGTGGTGACTGAGTCAGTGACAAAAATCATGGCATGCGGGACCACGCTGATGGGCTACACGCAATGGTGCTGTTCGTCACCGGACTGTTGCCACACCAAAAAGGTCTGCTTCCGGTGTAAAAGCCGCTCCTGTCCGCACTGCGGGGTGAAGGCTGGCGCACAGTGGATACAGTATCTGCTGAGCCTGGTCCCCGACTGCCCGTGGCAGCATATTGTGTTCACACTTCCCTGCCAGTACTGGTCCCTGGTGTTCCACAACCGGTGGTTACTGGCAGAGATGAGCCGCATTGCAGCGGATGTGATACTGGAAATCTGCCATCAGACAGATGTGGAGCCGGGGATATTCACGGTGATCCACACATGGGGGCGTGACCAGCAGTGGCATCCGCATATCCATTTATCGACAACTGCCGGTGGTGTGACGTCGGGCCACACCTGGAAAAATCTTCATTTTTACGCCCGTAAGGTGATGAGCATGTGGCGTTACCGGATAACGCGGCTACTGTCCCGGAAATACCCGGAGCTGGTAATACCGGATGAACTGGCAGTGGAAGGAAACAGCAAACGGGACTGGAATTGCTTCCTGGACACGCATTACCGCCGCGGCTGGAATGTCAACATATCCAGGGTGATGGATAACGCCACACATGTGGCGGTGTACTTCGGCTCTTACCTGAAAAAGCCACCGGTGCCGATGAGTCGGCTGGAGCATTATGCCGGTCAGGATGAAATCGGTCTGCGTTACAACAGTCACCGTACAAAACGGGAAGAATACCTGTTGATGAGTGGAGATGAGTTCATGGAAAGGTTCTCCTGGCATGTAGCAGATAAGGGGTTCCGTATGGTGAGGTACTACGGTTTCCTGAGTCCGGTGAAGCGCCGCTTACTGGAAGAAGTTGTGTACGTCATAACGGAGACGGTGAGAAAAACGGCGATGCAAATCAGGTGGAGAGGGATGTATCAGAGGTTACTGAAGGTTGACCCGCTGAAGTGCATTCTGTGCGGAAGTCAGATGCGTTTTACGGGGCTGAAGCGGGGTTACCGACTGGCAGAGCTGGTCCTGATGCATGAGCGACTGGCACGACAGCAGGTGTGCGGCTGAGAGCCGCAGAGGGGAAGTTGCGTCCATTTTACCGGAAACGGAGCAAAAAACCGCCATTCATACCCTGTATCAATCAGTGTCATCCTGTTTAATAGTCGTTTCCGCTCATATGGTGCACAAGGGGTGTTGAAGAAACATCCGTTTTGTGGTGCTTTTTTAGCCTTTTGGGGATTTAAATTCCTATCGAT

>IS91-V14

CGAGTAGGCAGCCTGGCGGCTGCGGCTTGTCATGGTCTGGAATTACCGTTATAAAAAAAGATAATGTCATTGTCTTTCAGGTAGTTATATGGCCCGTTCAGCTAAACCCCGTAAACGCAAACCCGCACCACAAAGAAGCAAACTTCCCCGCTATGTTGTGAAACTTCATCCGGATGATTTTTTTGACGAAGAAGACGCTGAAGTTCTGCGCTTTGATAATTTTGACGATGCCGTTGAGTGCTGCGCTGACCTGGGTATTCCGTTCTTTCTGGATGCAGGAAACAAAAAGCTGGTCTTCTGGTTTGTTCGTGTCGATGACGAAGGGTATCCGGAAATAGCCCGCTGTACGGAGCGGGAGTTTGCAACCATTCTTGCCGGTATCAGTGCCGGTGGTATGTACTGCCCGGAATGCGGCACAGTTCACTGGCCGGATGGCGTTACCCCCCCGTCTGATGCTTCCCCGTTTTGCCGATATTTTTCAGCAGGGTAACCGCTGGCTTAACTGGCTGGAGAAACAGCCGGAAGGTTCAGTGCGTCCGGTGGTGACTGAGTCAGTGACAAAAATCATGGCATGCGGGACCACGCTGATGGGCTACACGCAATGGTGCTGTTCGTCACCGGACTGTTGCCACACCAAAAAGGTCTGCTTCCGGTGTAAAAGCCGCTCCTGTCCGCACTGCGGGGTGAAGGCTGGCGCACAGTGGATACAGTATCTGCTGAGCCTGGTCCCCGACTGCCCGTGGCAGCATATTGTGTTCACACTTCCCTGCCAGTACTGGTCCCTGGTGTTCCACAACCGGTGGTTACTGGCAGAGATGAGCCGCATTGCAGCGGATGTGATACTGGAAATCTGCCATCAGACAGATGTGGAGCCGGGGATATTCACGGTGATCCACACATGGGGGCGTGACCAGCAGTGGCATCCGCATATCCATTTATCGACAACTGCCGGTGGTGTGACGTCGGGCCACACCTGGAAAAATCTTCATTTTTACGCCCGTAAGGTGATGAGCATGTGGCGTTACCGGATAACGCGGCTACTGTCCCGGAAATACCCGGAGCTGGTAATACCGGATGAACTGGCAGTGGAAGGAAACAGCAAACGGGACTGGAATTGCTTCCTGGACACGCATTACCGCCGCGGCTGGAATGTCAACATATCCAGGGTGATGGATAACGCCACACATGTGGCGGTGTACTTCGGCTCTTACCTGAAAAAGCCACCGGTGCCGATGAGTCGGCTGGAGCATTATGCCGGTCAGGATGAAATCGGTCTGCGTTACAACAGTCACCGTACAAAACGGGAAGAATACCTGTTGATGAGTGGAGATGAGTTCATGGAAAGGTTCTCCTGGCATGTAGCAGATAAGGGGTTCCGTATGGTGAGGTACTACGGTTTCCTGAGTCCGGTGAAGCGCCGCTTACTGGAAGAAGTTGTGTACGTCATAACGGAGACGGTGAGAAAAACGGCGATGCAAATCAGGTGGAGAGGGATGTATCAGAGGTTACTGAAGGTTGACCCGCTGAAGTGCATTCTGTGCGGAAGTCAGATGCGTTTTACGGGGCTGAAGCGGGGTTACCGACTGGCAGAGCTGGTCCTGATGCATGAGCGACTGGCACGACAGCAGGTGTGCGGCTGAGAGCCGCAGAGGGGAAGTTGCGTCCATTTTACCGGAAACGGAGCAAAAAACCGCTATTCATACCCTGTATCAATCAGTGTCATCCTGTTTAATAGTCGTTTCCGCTCATATGGTGCACAAGGGGTGTTGAAGAAACATCCGTTTTGTGGTGCTTTTTTAGTCTTTTGGGGATTTAAATTCCTATCGAT

>IS91-V15

CGAGTAGGCAGCCTGGCGGCTGCGGCTTGTCATGGTCTGGAATTACCGTTATAAAAAAAGATAATGTCATTGTCTTTCAGGTAGTTATATGGCCCGTTCAGCTAAACCCCGTAAACGCAAACCCGCACCACAAAGAAGCAAACTTCCCCGCTATGTTGTGAAACTTCATCCGGATGATTTTTTTGACGAAGAAGACGCTGAAGTTCTGCGCTTTGATAATTTTGACGATGCCGTTGAGTGCTGCGCTGACCTGGGTATTCCGTTCTTTCTGGATGCAGGAAACAAAAAGCTGGTCTTCTGGTTTGTTCGTGTCGATGACGAAGGGTATCCGGAAATAGCCCGCTGTACGGAGCGGGAGTTTGCAACCATTCTTGCCGGTATCAGTGCCGGTGGTATGTACTGCCCGGAATGCGGCACAGTTCACTGGCCGGATGGCGTTACCCCCCCGTCTGATGCTTCCCCGTTTTGCCGATATTTTTCAGCAGGGTAACCGCTGGCTTAACTGGCTGGAGAAACAGCCGGAAGGTTCAGTGCGTCCGGTGGTGACTGAGTCAGTGACAAAAATCATGGCATGCGGGACCACGCTGATGGGCTACACGCAATGGTGCTGTTCGTCACCGGACTGTTGCCACACCAAAAAGGTCTGCTTCCGGTGTAAAAGCCGCTCCTGTCCGCACTGCGGGGTGAAGGCTGGCGCACAGTGGATACAGTATCTGCTGAGCCTGGTCCCCGACTGCCCGTGGCAGCATATTGTGTTCACACTTCCCTGCCAGTACTGGTCCCTGGTGTTCCACAACCGGTGGTTACAGGCAGAGATGAGCCGCATTGCAGCGGATGTGATACTGGAAATCTGCCATCAGACAGATGTGGAGCCGGGGATATTCACGGTGATCCACACATGGGGGCGTGACCAGCAGTGGCATCCGCATATCCATTTATCGACAACTGCCGGTGGTGTGACGTCGGGCCACACCTGGAAAAATCTTCATTTTTACGCCCGTAAGGTGATGAGCATGTGGCGTTACCGGATAACGCGGCTACTGTCCCGGAAATACCCGGAGCTGGTAATACCGGATGAACTGGCAGTGGAAGGAAACAGCAAACGGGACTGGAATTGCTTCCTGGACACGCATTACCGCCGCGGCTGGAATGTCAACATATCCAGGGTGATGGATAACGCCACACATGTGGCGGTGTACTTCGGCTCTTACCTGAAAAAGCCACCGGTGCCGATGAGTCGGCTGGAGCATTATGCCGGTCAGGATGAAATCGGTCTGCGTTACAACAGTCACCGTACAAAACGGGAAGAATACCTGTTGATGAGTGGAGATGAGTTCATGGAAAGGTTCTCCTGGCATGTAGCAGATAAGGGGTTCCGTATGGTGAGGTACTACGGTTTCCTGAGTCCGGTGAAGCGCCGCTTACTGGAAGAAGTTGTGTACGTCATAACGGAGACGGTGAGAAAAACGGCGATGCAAATCAGGTGGAGAGGGATGTATCAGAGGTTACTGAAGGTTGACCCGCTGAAGTGCATTCTGTGCGGAAGTCAGATGCGTTTTACGGGGCTGAAGCGGGGTTACCGACTGGCAGAGCTGGTCCTGATGCATGAGCGACTGGCACGACAGCAGGTGTGCGGCTGAGAGCCGCAGAGGGGAAGTTGCGTCCATTTTACCGGAAACGGAGCAAAAAACCGCCATTCATACCCTGTATCAATCAGTGTCATCCTGTTTAATAGTCGTTTCCGCTCATATGGTGCACAAGGGGTGTTGAAGAAACATCCGTTTTGTGGTGCTTTTTTAGTCTTTTGGGGATTTAAATTCCTATCGAT

>IS91-V16

CGAGTAGGCAGCCTGGCGGCTGCGGCTTGTCATGGTCTGGAATTACCGTTATAAAAAAAGATAATGTCATTGTCTTTCAGGTAGTTATATGGCCCGTTCAGCTAAACCCCGTAAACGCAAACCCGCACCACAAAGAAGCAAACTTCCCCGCTATGTTGTGAAACTTCATCCGGATGATTTTTTTGACGAAGAAGACGCTGAAGTTCTGCGCTTTGATAATTTTGACGATGCCGTTGAGTGCTGCGCTGACCTGGGTATTCCGTTCTTTCTGGATGCAGGAAACAAAAAGCTGGTCTTCTGGTTTGTTCGTGTCGATGACGAAGGGTATCCGGAAATAGCCCGCTGTACGGAGCGGGAGTTTGCAACCATTCTTGCCGGTATCAGTGCCGGTGGTATGTACTGCCCGGAATGCGGCACAGTTCACTGGCCGGATGGCGTTACCCCCCCGTCTGATGCTTCCCCGTTTTGCCGATATTTTTCAGCAGGGTAACCGCTGGCTTAACTGGCTGGAGAAACAGCCGGAAGGTTCAGTGCGTCCGGTGGTGACTGAGTCAGTGACAAAAATCATGGCATGCGGGACCACGCTGATGGGCTACACGCAATGGTGCTGTTCGTCACCGGACTGTTGCCACACCAAAAAGGTCTGCTTCCGGTGTAAAAGCCGCTCCTGTCCGCACTGCGGGGTGAAGGCTGGCGCACAGTGGATACAGTATCTGCTGAGCCTGGTCCCCGACTGCCCGTGGCAGCATATTGTGTTCACACTTCCCTGCCAGTACTGGTCCCTGGTGTTCCACAACCGGTGGTTACTGGCAGAGATGAGCCGCATTGCAGCGGATGTGATACTGGAAATCTGCCATCAGACAGATGTGGAGCCGGGGATATTCACGGTGATCCACACATGGGGGCGTGACCAGCAGTGGCATCCGCATATCCATTTATCGACAACTGCCGGTGGTGTGACGTCGGGCCACACCTGGAAAAATCTTCATTTTTACGCCCGTAAGGTGATGAGCATGTGGCGTTACCGGATAACGCGGCTACTGTCCCGGAAATACCCGGAGCTGGTAATACCGGATGAACTGGCAGTGGAAGGAAACAGCAAACGGGACTGGAATTGCTTCCTGGACACGCATTACCGCCGCGGCTGGAATGTCAACATATCCAGGGTGATGGATAACGCCACACATGTGGCGGTGTACTTCGGCTCTTACCTGAAAAAGCCACCGGTGCCGATGAGTCGGCTGGAGCATTATGCCGGTCAGGATGAAATCGGTCTGCGTTACAACAGTCACCGTACAAAACGGGAAGAATACCTGTTGATGAGTGGAGATGAGTTCATGGAAAGGTTCTCCTGGCATGTAGCAGATAAGGGGTTCCGTATGGTGAGGTACTACGGTTTCCTGAGTCCGGTGAAGCGCCGCTTACTGGAAGAAGTTGTGTACGTCATAACGGAGACGGTGAGAAAAACGGCGATGCAAATCAGGTGGAGAGGGATGTATCAGAGGTTACTGAAGGTTGACCCGCTGAAGTGCATTCTGTGCGGAAGTCAGATGCGTTTTACGGGGCTGAAGCGGGGTTACCGACTGGCAGAGCTGGTCCTGATGCATGAGCGACTGGCACGACAGCAGGTGTGCGGCTGAGAGCCGCAGAGGGGAAGTTGCGTCCATTTTACCGGAAACGGAGCAAAAAACCGCCATTCATACCCTGTATCAATCAGTGTCATCCTGTTTAATAGTCGTTTCCTCTCATATGGTGCACAAGGGGTGTTGAAGAAACATCCGTTTTGTGGTGCTTTTTTAGTCTTTTGGGGATTTAAATTCCTATCGAT

>IS91-V17

CGAGTAGGCAGCCTGGCGGCTGCGGCTTGTCATGGTCTGGAATTACCGTTATAAAAAAAGATAATGTCATTGTCTTTCAGGTAGTTATATGGCCCGTTCAGCTAAACCCCGTAAACGCAAACCCGCACCACAAAGAAGCAAACTTCCCCGCTATGTTGTGAAACATCATCCGGATGATTTTTTTGACGAAGAAGACGCTGAAGTTCTGCGCTTTGATAATTTTGACGATGCCGTTGAGTGCTGCGCTGACCTGGGTATTCCGTTCTTTCTGGATGCAGGAAACAAAAAGCTGGTCTTCTGGTTTGTTCGTGTCGATGACGAAGGGTATCCGGAAATAGCCCGCTGTACGGAGCGGGAGTTTGCAACCATTCTTGCCGGTATCAGTGCCGGTGGTATGTACTGCCCGGAATGCGGCACAGTTCACTGGCCGGATGGCGTTACCCCCCCGTCTGATGCTTCCCCGTTTTGCCGATATTTTTCAGCAGGGTAACCGCTGGCTTAACTGGCTGGAGAAACAGCCGGAAGGTTCAGTGCGTCCGGTGGTGACTGAGTCAGTGACAAAAATCATGGCATGCGGGACCACGCTGATGGGCTACACGCAATGGTGCTGTTCGTCACCGGACTGTTGCCACACCAAAAAGGTCTGCTTCCGGTGTAAAAGCCGCTCCTGTCCGCACTGCGGGGTGAAGGCTGGCGCACAGTGGATACAGTATCTGCTGAGCCTGGTCCCCGACTGCCCGTGGCAGCATATTGTGTTCACACTTCCCTGCCAGTACTGGTCCCTGGTGTTCCACAACCGGTGGTTACTGGCAGAGATGAGCCGCATTGCAGCGGATGTGATACTGGAAATCTGCCATCAGACAGATGTGGAGCCGGGGATATTCACGGTGATCCACACATGGGGGCGTGACCAGCAGTGGCATCCGCATATCCATTTATCGACAACTGCCGGTGGTGTGACGTCGGGCCACACCTGGAAAAATCTTCATTTTTACGCCCGTAAGGTGATGAGCATGTGGCGTTACCGGATAACGCGGCTACTGTCCCGGAAATACCCGGAGCTGGTAATACCGGATGAACTGGCAGTGGAAGGAAACAGCAAACGGGACTGGAATTGCTTCCTGGACACGCATTACCGCCGCGGCTGGAATGTCAACATATCCAGGGTGATGGATAACGCCACACATGTGGCGGTGTACTTCGGCTCTTACCTGAAAAAGCCACCGGTGCCGATGAGTCGGCTGGAGCATTATGCCGGTCAGGATGAAATCGGTCTGCGTTACAACAGTCACCGTACAAAACGGGAAGAATACCTGTTGATGAGTGGAGATGAGTTCATGGAAAGGTTCTCCTGGCATGTAGCAGATAAGGGGTTCCGTATGGTGAGGTACTACGGTTTCCTGAGTCCGGTGAAGCGCCGCTTACTGGAAGAAGTTGTGTACGTCATAACGGAGACGGTGAGAAAAACGGCGATGCAAATCAGGTGGAGAGGGATGTATCAGAGGTTACTGAAGGTTGACCCGCTGAAGTGCATTCTGTGCGGAAGTCAGATGCGTTTTACGGGGCTGAAGCGGGGTTACCGACTGGCAGAGCTGGTCCTGATGCATGAGCGACTGGCACGACAGCAGGTGTGCGGCTGAGAGCCGCAGAGGGGAAGTTGCGTCCATTTTACCGGAAACGGAGCAAAAAACCGCCATTCATACCCTGTATCAATCAGTGTCATCCTGTTTAATAGTCGTTTCCGCTCATATGGTGCACAAGGGGTGTTGAAGAAACATCCGTTTTGTGGTGCTTTTTTAGTCTTTTGGGGATTTAAATTCCTATCGAT

>IS91-V18

CGAGTAGGCAGCCTGGCGGCTGCGGCTTGTCATGGTCTGGAATTACCGTTATAAAAAAAGATAATGTCATTGTCTTTCAGGTAGTTATATGGCCCGTTCAGCTAAACCCCGTAAACGCAAACCCGCACCACAAAGAAGCAAACTTCCCCGCTATGTTGTGAAACTTCATCCGGATGATTTTTTTGACGAAGAAGACGCTGAAGTTCTGCGCTTTGATAATTTTGACGATGCCGTTGAGTGCTGCGCTGACCTGGGTATTCCGTTCTTTCTGGATGCAGGAAACAAAAAGCTGGTCTTCTGGTTTGTTCGTGTCGATGACGAAGGGTATCCGGAAATAGCCCGCTGTACGGAGCGGGAGTTTGCAACCATTCTTGCCGGTATCAGTGCCGGTGGTATGTACTGCCCGGAATGCGGCACAGTTCACTGGCCGGATGGCGTTACCCCCCGTCTGATGCTTCCCCGTTTTGCCGATATTTTTCAGCAGGGTAACCGCTGGCTTAACTGGCTGGAGAAACAGCCGGAAGGTTCAGTGCGTCCGGTGGTGACTGAGTCAGTGACAAAAATCATGGCATGCGGGACCACGCTGATGGGCTACACGCAATGGTGCTGTTCGTCACCGGACTGTTGCCACACCAAAAAGGTCTGCTTCCGGTGTAAAAGCCGCTCCTGTCCGCACTGCGGGGTGAAGGCTGGCGCACAGTGGATACAGTATCTGCTGAGCCTGGTCCCCGACTGCCCGTGGCAGCATATTGTGTTCACACTTCCCTGCCAGTACTGGTCCCTGGTGTTCCACAACCGGTGGTTACTGGCAGAGATGAGCCGCATTGCAGCGGATGTGATACTGGAAATCTGCCATCAGACAGATGTGGAGCCGGGGATATTCACGGTGATCCACACATGGGGGCGTGACCAGCAGTGGCATCCGCATATCCATTTATCGACAACTGCCGGTGGTGTGACGTCGGGCCACACCTGGAAAAATCTTCATTTTTACGCCCGTAAGGTGATGAGCATGTGGCGTTACCGGATAACGCGGCTACTGTCCCGGAAATACCCGGAGCTGGTAATACCGGATGAACTGGCAGTGGAAGGAAACAGCAAACGGGACTGGAATTGCTTCCTGGACACGCATTACCGCCGCGGCTGGAATGTCAACATATCCAGGGTGATGGATAACGCCACACATGTGGCGGTGTACTTCGGCTCTTACCTGAAAAAGCCACCGGTGCCGATGAGTCGGCTGGAGCATTATGCCGGTCAGGATGAAATCGGTCTGCGTTACAACAGTCACCGTACAAAACGGGAAGAATACCTGTTGATGAGTGGAGATGAGTTCATGGAAAGGTTCTCCTGGCATGTAGCAGATAAGGGGTTCCGTATGGTGAGGTACTACGGTTTCCTGAGTCCGGTGAAGCGCCGCTTACTGGAAGAAGTTGTGTACGTCATAACGGAGACGGTGAGAAAAACGGCGATGCAAATCAGGTGGAGAGGGATGTATCAGAGGTTACTGAAGGTTGACCCGCTGAAGTGCATTCTGTGCGGAAGTCAGATGCGTTTTACGGGGCTGAAGCGGGGTTACCGACTGGCAGAGCTGGTCCTGATGCATGAGCGACTGGCACGACAGCAGGTGTGCGGCTGAGAGCCGCAGAGGGGAAGTTGCGTCCATTTTACCGGAAACGGAGCAAAAAACCGCCATTCATACCCTGTATCAATCAGTGTCATCCTGTTTAATAGTCGTTTCCGCTCATATGGTGCACAAGGGGTGTTGAAGAAACATCCGTTTTGTGGTGCTTTTTTAGTCTTTTGGGGATTTAAATTCCTATCGAT

>IS91-V19

CGAGTAGGCAGCCTGGCGGCTGCGGCTTGTCATGGTCTGGAATTACCGTTATAAAAAAGATAATGTCATTGTCTTTCAGGTAGTTATATGGCCCGTTCAGCTAAACCCCGTAAACGCAAACCCGCACCACAAAGAAGCAAACTTCCCCGCTATGTTGTGAAACTTCATCCGGATGATTTTTTTGACGAAGAAGACGCTGAAGTTCTGCGCTTTGATAATTTTGACGATGCCGTTGAGTGCTGCGCTGACCTGGGTATTCCGTTCTTTCTGGATGCAGGAAACAAAAAGCTGGTCTTCTGGTTTGTTCGTGTCGATGACGAAGGGTATCCGGAAATAGCCCGCTGTACGGAGCGGGAGTTTGCAACCATTCTTGCCGGTATCAGTGCCGGTGGTATGTACTGCCCGGAATGCGGCACAGTTCACTGGCCGGATGGCGTTACCCCCCCGTCTGATGCTTCCCCGTTTTGCCGATATTTTTCAGCAGGGTAACCGCTGGCTTAACTGGCTGGAGAAACAGCCGGAAGGTTCAGTGCGTCCGGTGGTGACTGAGTCAGTGACAAAAATCATGGCATGCGGGACCACGCTGATGGGCTACACGCAATGGTGCTGTTCGTCACCGGACTGTTGCCACACCAAAAAGGTCTGCTTCCGGTGTAAAAGCCGCTCCTGTCCGCACTGCGGGGTGAAGGCTGGCGCACAGTGGATACAGTATCTGCTGAGCCTGGTCCCCGACTGCCCGTGGCAGCATATTGTGTTCACACTTCCCTGCCAGTACTGGTCCCTGGTGTTCCACAACCGGTGGTTACTGGCAGAGATGAGCCGCATTGCAGCGGATGTGATACTGGAAATCTGCCATCAGACAGATGTGGAGCCGGGGATATTCACGGTGATCCACACATGGGGGCGTGACCAGCAGTGGCATCCGCATATCCATTTATCGACAACTGCCGGTGGTGTGACGTCGGGCCACACCTGGAAAAATCTTCATTTTTACGCCCGTAAGGTGATGAGCATGTGGCGTTACCGGATAACGCGGCTACTGTCCCGGAAATACCCGGAGCTGGTAATACCGGATGAACTGGCAGTGGAAGGAAACAGCAAACGGGACTGGAATTGCTTCCTGGACACGCATTACCGCCGCGGCTGGAATGTCAACATATCCAGGGTGATGGATAACGCCACACATGTGGCGGTGTACTTCGGCTCTTACCTGAAAAAGCCACCGGTGCCGATGAGTCGGCTGGAGCATTATGCCGGTCAGGATGAAATCGGTCTGCGTTACAACAGTCACCGTACAAAACGGGAAGAATACCTGTTGATGAGTGGAGATGAGTTCATGGAAAGGTTCTCCTGGCATGTAGCAGATAAGGGGTTCCGTATGGTGAGGTACTACGGTTTCCTGAGTCCGGTGAAGCGCCGCTTACTGGAAGAAGTTGTGTACGTCATAACGGAGACGGTGAGAAAAACGGCGATGCAAATCAGGTGGAGAGGGATGTATCAGAGGTTACTGAAGGTTGACCCGCTGAAGTGCATTCTGTGCGGAAGTCAGATGCGTTTTACGGGGCTGAAGCGGGGTTACCGACTGGCAGAGCTGGTCCTGATGCATGAGCGACTGGCACGACAGCAGGTGTGCGGCTGAGAGCCGCAGAGGGGAAGTTGCGTCCATTTTACCGGAAACGGAGCAAAAAACCGCCATTCATACCCTGTATCAATCAGTGTCATCCTGTTTAATAGTCGTTTCCGCTCATATGGTGCACAAGGGGTGTTGAAGAAACATCCGTTTTGTGGTGCTTTTTTAGTCTTTTGGGGATTTAAATTCCTATCGAT

>IS91-V20

CGAGTAGGCAGCCTGGCGGCTGCGGCTTGTCATGGTCTGGAATTACCGTTATAAAAAAAGATAATGTCATTGTCTTTCAGGTAGTTATATGGCCCGTTCAGCTAAACCCCGTAAACGCAAACCCGCACCACAAAGAAGCAAACTTCCCCGCTATGTTGTGAAACTTCATCCGGATGATTTTTTTGACGAAGAAGACGCTGAAGTTCTGCGCTTTGATAATTTTGACGATGCCGTTGAGTGCTGCGCTGACCTGGGTATTCCGTTCTTTCTGGATGCAGGAAACAAAAAGCTGGTCTTCTGGTTTGTTCGTGTCGATGACGAAGGGTATCCGGAAATAGCCCGCTGTACGGAGCGGGAGTTTGCAACCATTCTTGCCGGTATCAGTGCCGGTGGTATGTACTGCCCGGAATGCGGCACAGTTCACTGGCCGGATGGCGTTACCCCCCCGTCTGATGCTTCCCCGTTTTGCCGATATTTTTCAGCAGGGTAACCGCTGGCTTAACTGGCTGGAGAAACAGCCGGAAGGTTCAGTGCGTCCGGAGGTGACTGAGTCAGTGACAAAAATCATGGCATGCGGGACCACGCTGATGGGCTACACGCAATGGTGCTGTTCGTCACCGGACTGTTGCCACACCAAAAAGGTCTGCTTCCGGTGTAAAAGCCGCTCCTGTCCGCACTGCGGGGTGAAGGCTGGCGCACAGTGGATACAGTATCTGCTGAGCCTGGTCCCCGACTGCCCGTGGCAGCATATTGTGTTCACACTTCCCTGCCAGTACTGGTCCCTGGTGTTCCACAACCGGTGGTTACTGGCAGAGATGAGCCGCATTGCAGCGGATGTGATACTGGAAATCTGCCATCAGACAGATGTGGAGCCGGGGATATTCACGGTGATCCACACATGGGGGCGTGACCAGCAGTGGCATCCGCATATCCATTTATCGACAACTGCCGGTGGTGTGACGTCGGGCCACACCTGGAAAAATCTTCATTTTTACGCCCGTAAGGTGATGAGCATGTGGCGTTACCGGATAACGCGGCTACTGTCCCGGAAATACCCGGAGCTGGTAATACCGGATGAACTGGCAGTGGAAGGAAACAGCAAACGGGACTGGAATTGCTTCCTGGACACGCATTACCGCCGCGGCTGGAATGTCAACATATCCAGGGTGATGGATAACGCCACACATGTGGCGGTGTACTTCGGCTCTTACCTGAAAAAGCCACCGGTGCCGATGAGTCGGCTGGAGCATTATGCCGGTCAGGATGAAATCGGTCTGCGTTACAACAGTCACCGTACAAAACGGGAAGAATACCTGTTGATGAGTGGAGATGAGTTCATGGAAAGGTTCTCCTGGCATGTAGCAGATAAGGGGTTCCGTATGGTGAGGTACTACGGTTTCCTGAGTCCGGTGAAGCGCCGCTTACTGGAAGAAGTTGTGTACGTCATAACGGAGACGGTGAGAAAAACGGCGATGCAAATCAGGTGGAGAGGGATGTATCAGAGGTTACTGAAGGTTGACCCGCTGAAGTGCATTCTGTGCGGAAGTCAGATGCGTTTTACGGGGCTGAAGCGGGGTTACCGACTGGCAGAGCTGGTCCTGATGCATGAGCGACTGGCACGACAGCAGGTGTGCGGCTGAGAGCCGCAGAGGGGAAGTTGCGTCCATTTTACCGGAAACGGAGCAAAAAACCGCCATTCATACCCTGTATCAATCAGTGTCATCCTGTTTAATAGTCGTTTCCGCTCATATGGTGCACAAGGGGTGTTGAAGAAACATCCGTTTTGTGGTGCTTTTTTAGTCTTTTGGGGATTTAAATTCCTATCGAT

>IS91-V21

CGAGTAGGCAGCCTGGCGGCTGCGGCTTGTCATGGTCTGGAATTACCGTTATAAAAAAAGATAATGTCATTGTCTTTCAGGTAGTTATATGGCCCGTTCAGCTAAACCCCGTAAACGCAAACCCGCACCACAAAGAAGCAAACTTCCCCGCTATGTTGTGAAACTTCATCCGGATGATTTTTTTGACGAAGAAGACGCTGAAGTTCTGCGCTTTGATAATTTTGACGATGCCGTTGAGTGCTGCGCTGACCTGGGTATTCCGTTCTTTCTGGATGCAGGAAACAAAAAGCTGGTCTTCTGGTTTGTTCGTGTCGATGACGAAGGGTATCCGGAAATAGCCCGCTGTACGGAGCGGGAGTTTGCAACCATTCTTGCCGGTATCAGTGCCGGTGGTATGTACTGCCCGGAATGCGGCACAGTTCACTGGCCGGATGGCGTTACCCCCCCGTCTGATGCTTCCCCGTTTTGCCGATATTTTTCAGCAGGGTAACCGCTGGCTTAACTGGCTGGAGAAACAGCCGGAAGGTTCAGTGCGTCCGGTGGTGACTGAGTCAGTGACAAAAATCATGGCATGCGGGACCACGCTGATGGGCTACACGCAATGGTGCTGTTCGTCACCGGACTGTTGCCACACCAAAAAGGTCTGCTTCCGGTGTAAAAGCCGCTCCTGTCCGCACTGCGGGGTGAAGGCTGGCGCACAGTGGATACAGTATCTGCTGAGCCTGGTCCCCGACTGCCCGTGGCAGCATATTGTGTTCACACTTCCCTGCCAGTACTGGTCCCTGGTGTTCCACAACCGGTGGTTACTGGCAGAGATGAGCCGCATTGCAGCGGATGTGATACTGGAAATCTGCCATCAGACAGATGTGGAGCCGGGGATATTCACGGTGATCCACACATGGGGGCGTGACCAGCAGTGGCATCCGCATATCCATTTATCGACAACTGCCGGTGGTGTGACGTCGGGCCACACCTGGAAAAATCTTCATTTTTACGCCCGTAAGGTGATGAGCATGTGGCGTTACCGGATAACGCGGCTACTGTCCCGGAAATACCCGGAGCTGGTAATACCGGATGAACTGGCAGTGGAAGGAAACAGCAAACGGGACTGGAATTGCTTCCTGGACACGCATTACCGCCGCGGCTGGAATGTCAACATATCCAGGGTGATGGATAACGCCACACATGTGGCGGTGTACTTCGGCTCTTACCTGAAAAAGCCACCGGTGCCGATGAGTCGGCTGGAGCATTATGCCGGTCAGGATGAAATCGGTCTGCGTTACAACAGTCACCGTACAAAACGGGAAGAATACCTGTTGATGAGTGGAGATGAGTTCATGGAAAGGTTCTCCTGGCATGTAGCAGATAAGGGGTTCCGTATGGTGAGGTACTACGGTTTCCTGAGTCCGGTGAAGCGCCGGTTACTGGAAGATGTTGTGTACGTCATAACGGAGACGGTGAGAAAGACGGCGATGCAAATCAGGTGGAGAGGGATGTATCAGAGGTTACTGAAGGTTGACCCGCTGAAGTGCATTCTGTGCGGAAGTCAGATGCGTTTTACGGGGCTGAAGCGGGGTTACCGACTGGCAGAGCTGGTCCTGATGCATGAGCGACTGGCACGACAGCAGGTGTGCGGCTGAGAGCCGCAGAGGGGAAGTTGCGTCCATTTTACCGGAAACGGAGCAAAAAACCGCCATTCATACCCTGTATCAATCAGTGTCATCCTGTTTAATAGTCGTTTCCGCTCATATGGTGCACAAGGGGTGTTGAAGAAACATCCGTTTTGTGGTGCTTTTTTAGTCTTTTGGGGATTTAAATTCCTATCGAT

>IS91-V22

CGAGTAGGCAGCCTGGCGGCTGCGGCTTGTCATGGTCTGGAATTACCGTTAATAAAAAAAGATAATGTCATTGTCTTTCAGGTAGTTATATGGCCCGTTCAGCTAAACCCCGTAAACGCAAACCCGCACCACAAAGAAGCAAACTTCCCCGCTATGTTGTGAAACTTCATCCGGATGATTTTTTGACGAAGAAGACGCTGAAGTTCTGCGCTTTGATAATTTTGACGATGCCGTTGAGTGCTGCGCTGACCTGGGTATTCCGTTCTTTCTGGATGCAGGAAACAAAAAGCTGGTCTTCTGGTTTGTTCGTGTCGATGACGAAGGGTATCCGGAAATAGCCCGCTGTACGGAGCGGGAGTTTGCAACCATTCTTGCCGGTATCAGTGCCGGTGGTATGTACTGCCCGGAATGCGGCACAGTTCACTGGCCGGATGGCGTTACCCCCCCGTCTGATGCTTCCCCGTTTTGCCGATATTTTTCAGCAGGGTAACCGCTGGCTTAACTGGCTGGAGAAACAGCCGGAAGGTTCAGTGCGTCCGGTGGTGACTGAGTCAGTGACAAAAATCATGGCATGCGGGACCACGCTGATGGGCTACACGCAATGGTGCTGTTCGTCACCGGACTGTTGCCACACCAAAAAGGTCTGCTTCCGGTGTAAAAGCCGCTCCTGTCCGCACTGCGGGGTGAAGGCTGGCGCACAGTGGATACAGTATCTGCTGAGCCTGGTCCCCGACTGCCCGTGGCAGCATATTGTGTTCACACTTCCCTGCCAGTACTGGTCCCTGGTGTTCCACAACCGGTGGTTACTGGCAGAGATGAGCCGCATTGCAGCGGATGTGATACTGGAAATCTGCCATCAGACAGATGTGGAGCCGGGGATATTCACGGTGATCCACACATGGGGGCGTGACCAGCAGTGGCATCCGCATATCCATTTATCGACAACTGCCGGTGGTGTGACGTCGGGCCACACCTGGAAAAATCTTCATTTTTACGCCCGTAAGGTGATGAGCATGTGGCGTTACCGGATAACGCGGCTACTGTCCCGGAAATACCCGGAGCTGGTAATACCGGATGAACTGGCAGTGGAAGGAAACAGCAAACGGGACTGGAATTGCTTCCTGGACACGCATTACCGCCGCGGCTGGAATGTCAACATATCCAGGGTGATGGATAACGCCACACATGTGGCGGTGTACTTCGGCTCTTACCTGAAAAAGCCACCGGTGCCGATGAGTCGGCTGGAGCATTATGCCGGTCAGGATGAAATCGGTCTGCGTTACAACAGTCACCGTACAAAACGGGAAGAATACCTGTTGATGAGTGGAGATGAGTTCATGGAAAGGTTCTCCTGGCATGTAGCAGATAAGGGGTTCCGTATGGTGAGGTACTACGGTTTCCTGAGTCCGGTGAAGCGCCGCTTACTGGAAGAAGTTGTGTACGTCATAACGGAGACGGTGAGAAAAACGGCGATGCAAATCAGGTGGAGAGGGATGTATCAGAGGTTACTGAAGGTTGACCCGCTGAAGTGCATTCTGTGCGGAAGTCAGATGCGTTTTACGGGGCTGAAGCGGGGTTACCGACTGGCAGAGCTGGTCCTGATGCATGAGCGACTGGCACGACAGCAGGTGTGCGGCTGAGAGCCGCAGAGGGGAAGTTGCGTCCATTTTACCGGAAACGGAGCAAAAAACCGCCATTCATACCCTGTATCAATCAGTGTCATCCTGTTTAATAGTCGTTTCCGCTCATATGGTGCACAAGGGGTGTTGAAGAAACATCCGTTTTGTGGTGCTTTTTTAGTCTTTTGGGGATTTAAATTCCTATCGAT

>IS91-V23

CGAGTAGGCAGCCTGGCGGCTGCGGCTTGTCATGGTCTGGAATTACCGTTATAAAAAAAGATAATGTCATTGTCTTTCAGGTAGTTATATGGCCCGTTCAGCTAAACCCCGTAAACGCAAACCCGCACCACAAAGAAGCAAACTTCCCCGCTATGTTGTGAAACTTCATCCGGATGATTTTTTTGACGAAGAAGACGCTGAAGTTCTGCGCTTTGATAATTTTGACGATGCCGTTGAGTGCTGCGCTGACCTGGGTATTCCGTTCTTTCTGGATGCAGGAAACAAAAAGCTGGTCTTCTGGTTTGTTCGTGTCGATGACGAAGGGTATCCGGAAATAGCCCGCTGTACGGAGCGGGAGTTTGCAACCATTCTTGCCGGTATCAGTGCCGGTGGTATGTACTGCCCGGAATGCGGCACAGTTCACTGGCCGGATGGCGTTACCCCACCCGTCTGATGCTTCCCCGTTTTGCCGATATTTTTCAGCAGGGTAACCGCTGGCTTAACTGGCTGGAGAAACAGCCGGAAGGTTCAGTGCGTCCGGTGGTGACTGAGTCAGTGACAAAAATCATGGCATGCGGGACCACGCTGATGGGCTACACGCAATGGTGCTGTTCGTCACCGGACTGTTGCCACACCAAAAAGGTCTGCTTCCGGTGTAAAAGCCGCTCCTGTCCGCACTGCGGGGTGAAGGCTGGCGCACAGTGGATACAGTATCTGCTGAGCCTGGTCCCCGACTGCCCGTGGCAGCATATTGTGTTCACACTTCCCTGCCAGTACTGGTCCCTGGTGTTCCACAACCGGTGGTTACTGGCAGAGATGAGCCGCATTGCAGCGGATGTGATACTGGAAATCTGCCATCAGACAGATGTGGAGCCGGGGATATTCACGGTGATCCACACATGGGGGCGTGACCAGCAGTGGCATCCGCATATCCATTTATCGACAACTGCCGGTGGTGTGACGTCGGGCCACACCTGGAAAAATCTTCATTTTTACGCCCGTAAGGTGATGAGCATGTGGCGTTACCGGATAACGCGGCTACTGTCCCGGAAATACCCGGAGCTGGTAATACCGGATGAACTGGCAGTGGAAGGAAACAGCAAACGGGACTGGAATTGCTTCCTGGACACGCATTACCGCCGCGGCTGGAATGTCAACATATCCAGGGGGATGGATAACGCCACACATGTGGCGGTGTACTTCGGCTCTTACCTGAAAAAGCCACCGGTGCCGATGAGTCGGCTGGAGCATTATGCCGGTCAGGATGAAATCGGTCTGCGTTACAACAGTCACCGTACAAAACGGGAAGAATACCTGTTGATGAGTGGAGATGAGTTCATGGAAAGGTTCTCCTGGCATGTAGCAGATAAGGGGTTCCGTATGGTGAGGTACTACGGTTTCCTGAGTCCGGTGAAGCGCCGCTTACTGGAAGAAGTTGTGTACGTCATAACGGAGACGGTGAGAAAAACGGCGATGCAAATCAGGTGGAGAGGGATGTATCAGAGGTTACTGAAGGTTGACCCGCTGAAGTGCATTCTGTGCGGAAGTCAGATGCGTTTTACGGGGCTGAAGCGGGGTTACCGACTGGCAGAGCTGGTCCTGATGCATGAGCGACTGGCACGACAGCAGGTGTGCGGCTGAGAGCCGCAGAGGGGAAGTTGCGTCCATTTTACCGGAAACGGAGCAAAAAAACCGCCATTCATACCCTGTATCAATCAGTGTCATCCTGTTTAATAGTCGTTTCCGCTCATATGGTGCACAAGGGGTGTTGAAGAAACATCCGTTTTGTGGTGCTTTTTTAGTCTTTTGGGGATTTAAATTCCTATCGAT

>IS91-V24

CGAGTAGGCAGCCTGGCGGCTGCGGCTTGTCATGGTCTGGAATTACCGTTATAAAAAAAGATAATGTCATTGTCTTTCAGGTAGTTATATGGCCCGTTCAGCTAAACCCCGTAAACGCAAACCCGCACCACAAAGAAGCAAACTTCCCCGCTATGTTGTGAAACTTCATCCGGATGATTTTTTTGACGAAGAAGACGCTGAAGTTCTGCGCTTTGATAATTTTGACGATGCCGTTGAGTGCTGCGCTGACCTGGGTATTCCGTTCTTTCTGGATGCAGGAAACAAAAAGCTGGTCTTCTGGTTTGTTCGTGTCGATGACGAAGGGTATCCGGAAATAGCCCGCTGTACGGAGCGGGAGTTTGCAACCATTCTTGCCGGTATCAGTGCCGGTGGTATGTACTGCCCGGAATGCGGCACAGTTCACTGGCCGGATGGCGTTACCCCACCCGTCTGATGCTTCCCCGTTTTGCCGATATTTTTCAGCAGGGTAACCGCTGGCTTAACTGGCTGGAGAAACAGCCGGAAGGTTCAGTGCGTCCGGTGGTGACTGAGTCAGTGACAAAAATCATGGCATGCGGGACCACGCTGATGGGCTACACGCAATGGTGCTGTTCGTCACCGGACTGTTGCCACACCAAAAAGGTCTGCTTCCGGTGTAAAAGCCGCTCCTGTCCGCACTGCGGGGTGAAGGCTGGCGCACAGTGGATACAGTATCTGCTGAGCCTGGTCCCCGACTGCCCGTGGCAGCATATTGTGTTCACACTTCCCTGCCAGTACTGGTCCCTGGTGTTCCACAACCGGTGGTTACTGGCAGAGATGAGCCGCATTGCAGCGGATGTGATACTGGAAATCTGCCATCAGACAGATGTGGAGCCGGGGATATTCACGGTGATCCACACATGGGGGCGTGACCAGCAGTGGCATCCGCATATCCATTTATCGACAACTGCCGGTGGTGTGACGTCGGGCCACACCTGGAAAAATCTTCATTTTTACGCGCGTAAGGTGATGAGCATGTGGCGTTACCGGATAACGCGGCTACTGTCCCGGAAATACCCGGAGCTGGTAATACCGGATGAACTGGCAGTGGAAGGAAACAGCAAACGGGACTGGAATCGCTTCCTGGACACGCATTACCGCCGCGGCTGGAATGTCAACATATCCAGGGTGATGGATAACGCCACACATGTGGCGGTGTACTTCGGCTCTTACCTGAAAAAGCCACCGGTGCCGATGAGTCGGCTGGAGCATTATGCCGGTCAGGATGAAATCGGTCTGCGTTACAACAGTCACCGTACAAAACGGGAAGAATACCTGTTGATGAGTGGAGATGAGTTCATGGAAAGGTTCTCCTGGCATGTAGCAGATAAGGGGTTCCGTATGGTGAGGTACTACGGTTTCCTGAGTCCGGTGAAGCGCCGCTTACTGGAAGAAGTTGTGTACGTCATAACGGAGACGGTGAGAAAAACGGCGATGCAAATCAGGTGGAGAGGGATGTATCAGAGGTTACTGAAGGTTGACCCGCTGAAGTGCATTCTGTGCGGAAGTCAGATGCGTTTTACGGGGCTGAAGCGGGGTTACCGACTGGCAGAGCTGGTCCTGATGCATGAGCGACTGGCACGACAGCAGGTGTGCGGCTGAGAGCCGCAGAGGGGAAGTTGCGTCCATTTTACCGGAAACGGAGCAAAAAACCGCCATTCATACCCTGTATCAATCAGTGTCATCCTGTTTAATAGTCGTTTCCGCTCATATGGTGCACAAGGGGTGTTGAAGAAACATCCGTTTTGTGGTGCTTTTTTAGTCTTTTGGGGATTTAAATTCCTATCGAT

>IS91-V25

CGAGTAGGCAGCCTGGCGGCTGCGGCTTGTCATGGTCTGGAATTACCGTTATAAAAAAAGATAATGTCATTGTCTTTCAGGTAGTTATATGGCCCGTTCAGCTAAACCCCGTAAACGCAAACCCGCACCACAAAGAAGCAAACTTCCCCGCTATGTTGTGAAACTTCATCCGGATGATTTTTTTGACGAAGAAGACGCTGAAGTTCTGCGCTTTGATAATTTTGACGATGCCGTTGAGTGCTGCGCTGACCTGGGTATTCCGTTCTTTCTGGATGCAGGAAACAAAAAGCTGGTCTTCTGGTTTGTTCGTGTCGATGACGAAGGGTATCCGGAAATAGCCCGCTGTACGGAGCGGGAGTTTGCGACCATTCTTGCCGGTATCAGTGCCGGTGGTATGTACTGCCCGGAATGCGGCACAGTTCACTGGCCGGATGGCGTTACCCCACCCGTCTGATGCTTCCCCGTTTTGCCGATATTTTTCAGCAGGGTAACCGCTGGCTTAACTGGCTGGAGAAACAGCCGGAAGGTTCAGTGCGTCCGGTGGTGACTGAGTCAGTGACAAAAATCATGGCATGCGGGACCACGCTGATGGGCTACACGCAATGGTGCTGTTCGTCACCGGACTGTTGCCACACCAAAAAGGTCTGCTTCCGGTGTAAAAGCCGCTCCTGTCCGCACTGCGGGGTGAAGGCTGGCGCACAGTGGATACAGTATCTGCTGAGCCTGGTCCCCGACTGCCCGTGGCAGCATATTGTGTTCACACTTCCCTGCCAGTACTGGTCCCTGGTGTTCCACAACCGGTGGTTACTGGCAGAGATGAGCCGCATTGCAGCGGATGTGATACTGGAAATCTGCCATCAGACAGATGTGGAGCCGGGGATATTCACGGTGATCCACACATGGGGGCGTGACCAGCAGTGGCATCCGCATATCCATTTATCGACAACTGCCGGTGGTGTGACGTCGGGCCACACCTGGAAAAATCTTCATTTTTACGCGCGTAAGGTGATGAGCATGTGGCGTTACCGGATAACGCGGCTACTGTCCCGGAAATACCCGGAGCTGGTAATACCGGATGAACTGGCAGTGGAAGGAAACAGCAAACGGGACTGGAATCGCTTCCTGGACACGCATTACCGCCGCGGCTGGAATGTCAACATATCCAGGGTGATGGATAACGCCACACATGTGGCGGTGTACTTCGGCTCTTACCTGAAAAAGCCACCGGTGCCGATGAGTCGGCTGGAGCATTATGCCGGTCAGGATGAAATCGGTCTGCGTTACAACAGTCACCGTACAAAACGGGAAGAATACCTGTTGATGAGTGGAGATGAGTTCATGGAAAGGTTCTCCTGGCATGTAGCAGATAAGGGGTTCCGTATGGTGAGGTACTACGGTTTCCTGAGTCCGGTGAAGCGCCGCTTACTGGAAGAAGTTGTGTACGTCATAACGGAGACGGTGAGAAAAACGGCGATGCAAATCAGGTGGAGAGGGATGTATCAGAGGTTACTGAAGGTTGACCCGCTGAAGTGCATTCTGTGCGGAAGTCAGATGCGTTTTACGGGGCTGAAGCGGGGTTACCGACTGGCAGAGCTGGTCCTGATGCATGAGCGACTGGCACGACAGCAGGTGTGCGGCTGAGAGTCGCAGAGGGGAAGTTGCGTCCATTTTACCGGAAACGGAGCAAAAAACCGCCATTCATACCCTGTATCAATCAGTGTCATCCTGTTTAATAGTCGTTTCCGCTCATATGGTGCACAAGGGGTGTTGAAGAAACATCCGTTTTGTGGTGCTTTTTTAGTCTTTTGGGGATTTAAATTCCTATCGAT

>IS91-V26

CGAGTAGGCAGCCTGGCGGCTGCGGCTTGTCATGGTCTGGAATTACCGTTATAAAAAAAGATAATGTCATTGTCTTTCAGGTAGTTATATGGCCCGTTCAGCTAAACCCCGTAAACGCAAACCCGCACCACAAAGAAGCAAACTTCCCCGCTATGTTGTGAAACTTCATCCGGATGATTTTTTTGACGAAGAAGACGCTGAAGTTCTGCGCTTTGATAATTTTGACGATGCCGTTGAGTGCTGCGCTGACCTGGGTATTCCGTTCTTTCTGGATGCAGGAAACAAAAAGCTGGTCTTCTGGTTTGTTCGTGTCGATGACGAAGGGTATCCGGAAATAGCCCGCTGTACGGAGCGGGAGTTTGCGACCATTCTTGCCGGTATCAGTGCCGGTGGTATGTACTGCCCGGAATGCGGCACAGTTCACTGGCCGGATGGCGTTACCCCACCCGTCTGATGCTTCCCCGTTTTGCCGATATTTTTCAGCAGGGTAACCGCTGGCTTAACTGGCTGGAGAAACAGCCGGAAGGTTCAGTGCGTCCGGTGGTGACTGAGTCAGTGACAAAAATCATGGCATGCGGGACCACGCTGATGGGCTACACGCAATGGTGCTGTTCGTCACCGGACTGTTGCCACACCAAAAAGGTCTGCTTCCGGTGTAAAAGCCGCTCCTGTCCGCACTGCGGGGTGAAGGCTGGCGCACAGTGGATACAGTATCTGCTGAGCCTGGTCCCCGACTGCCCGTGGCAGCATATTGTGTTCACACTTCCCTGCCAGTACTGGTCCCTGGTGTTCCACAACCGGTGGTTACTGGCAGAGATGAGCCGCATTGCAGCGGATGTGATACTGGAAATCTGCCATCAGACAGATGTGGAGCCGGGGATATTCACGGTGATCCACACATGGGGGCGTGACCAGCAGTGGCATCCGCATATCCATTTATCGACAACTGCCGGTGGTGTGACGTCGGGCCACACCTGGAAAAATCTTCATTTTTACGCGCGTAAGGTGATGAGCATGTGGCGTTACCGGATAACGCGGCTACTGTCCCGGAAATACCCGGAGCTGGTAATACCGGATGAACTGGCAGTGGAAGGAAACAGCAAACGGGACTGGAATCGCTTCCTGGACACGCATTACCGCCGCGGCTGGAATGTCAACATATCCAGGGTGATGGATAACGCCACACATGTGGCGGTGTACTTCGGCTCTTACCTGAAAAAGCCACCGGTGCCGATGAGTCGGCTGGAGCATTATGCCGGTCAGGATGAAATCGGTCTGCGTTACAACAGTCACCGTACAAAACGGGAAGAATACCTGTTGATGAGTGGAGATGAGTTCATGGAAAGGTTCTCCTGGCATGTAGCAGATAAGGGGTTCCGTATGGTGAGGTACTACGGTTTCCTGAGTCCGGTGAAGCGCCGCTTACTGGAAGAAGTTGTGTACGTCATAACGGAGACGGTGAGAAAAACGGCGATGCAAATCAGGTGGAGAGGGATGTATCAGAGGTTACTGAAGGTTGACCCGCTGAAGTGCATTCTGTGCGGAAGTCAGATGCGTTTTACGGGGCTGAAGCGGGGTTACCGACTGGCAGAGCTGGTCCTGATGCATGAGCGACTGGCACGACAGCAGGTGTGCGGCTGAGAGTCGCAGAGGGGAAGTTGCGTCCATTTTACCGGAAACGGAGCAAAAAACCGCCATTCATACCCTGTATCAATCAGTGTCATCCTGTTTAATAGTCGTTTCCGCTCATATGGTGCACAAGGGGTGTTGAAGAAACATCCGTTTTGTGGTGCTTTTTTAGTCTTTTGGGGATTTAAATTCCTATCGAT

>IS91-V27

CGAGTAGGCAGCCTGGCGGCTGCGGCTTGTCATGGTCTGGAATTACCGTTATAAAAAAAGATAATGTCATTGTCTTTCAGGTAGTTATATGGCCCGTTCAGCTAAACCCCGTAAACGCAAACCCGCACCACAAAGAAGCAAACTTCCCCGCTATGTTGTGAAACTTCATCCGGATGATTTTTTTGACGAAGAAGACGCTGAAGTTCTGCGCTTTGATAATTTTGACGATGCCGTTGAGTGCTGCGCTGACCTGGGTATTCCGTTCTTTCTGGATGCAGGAAACAAAAAGCTGGTCTTCTGGTTTGTTCGTGTCGATGACGAAGGGTATCCGGAAATAGCCCGCTGTACGGAGCGGGAGTTTGCAACCATTCTTGCCGGTATCAGTGCCGGTGGTATGTACTGCCCGGAATGCGGCACAGTTCACTGGCCGGATGGCGTTACCCCACCCGTCTGATGCTTCCCCGTTTTGCCGATATTTTTCAGCAGGGTAACCGCTGGCTTAACTGGCTGGAGAAACAGCCGGAAGGTTCAGTGCGTCCGGTGGTGACTGAGTCAGTGACAAAAATCATGGCATGCGGGACCACGCTGATGGGCTACACGCAATGGTGCTGTTCGTCACCGGACTGTTGCCACACCAAAAAGGTCTGCTTCCGGTGTAAAAGCCGCTCCTGTCCGCACTGCGGGGTGAAGGCTGGCGCACAGTGGATACAGTATCTGCTGAGCCTGGTCCCCGACTGCCCGTGGCAGCATATTGTGTTCACACTTCCCTGCCAGTACTGGTCCCTGGTGTTCCACAACCGGTGGTTACTGGCAGAGATGAGCCGCATTGCAGCGGATGTGATACTGGAAATCTGCCATCAGACAGATGTGGAGCCGGGGATATTCACGGTGATCCACACATGGGGGCGTGACCAGCAGTGGCATCCGCATATCCATTTATCGACAACTGCCGGTGGTGTGACGTCGGGCCACACCTGGAAAAATCTTCATTTTTACGCGCGTAAGGTGATGAGCATGTGGCGTTACCGGATAACGCGGCTACTGTCCCGGAAATACCCGGAGCTGGTAATACCGGATGAACTGGCAGTGGAAGGAAACAGCAAACGGGACTGGAATCGCTTCCTGGACACGCATTACCGCCGCGGCTGGAATGTCAACATATCCAGGGTGATGGATAACGCCACACTTGTGGCGGTGTACTTCGGCTCTTACCTGAAAAAGCCACCGGTGCCGATGAGTCGGCTGGAGCATTATGCCGGTCAGGATGAAATCGGTCTGCGTTACAACAGTCACCGTACAAAACGGGAAGAATACCTGTTGATGAGTGGAGATGAGTTCATGGAAAGGTTCTCCTGGCATGTAGCAGATAAGGGGTTCCGTATGGTGAGGTACTACGGTTTCCTGAGTCCGGTGAAGCGCCGCTTACTGGAAGAAGTTGTGTACGTCATAACGGAGACGGTGAGAAAAACGGCGATGCAAATCAGGTGGAGAGGGATGTATCAGAGGTTACTGAAGGTTGACCCGCTGAAGTGCATTCTGTGCGGAAGTCAGATGCGTTTTACGGGGCTGAAGCGGGGTTACCGACTGGCAGAGCTGGTCCTGATGCATGAGCGACTGGCACGACAGCAGGTGTGCGGCTGAGAGCCGCAGAGGGGAAGTTGCGTCCATTTTACCGGAAACGGAGCAAAAAACCGCCATTCATACCCTGTATCAATCAGTGTCATCCTGTTTAATAGTCGTTTCCGCTCATATGGTGCACAAGGGGTGTTGAAGAAACATCCGTTTTGTGGTGCTTTTTTAGTCTTTTGGGGATTTAAATTCCTATCGAT

>IS91-V28

CGAGTAGGCAGCCTGGCGGCTGCGGCTTGTCATGGCCTGGAATTACCGTTATAAAAAAAGATAATGTCATTGTCTTTCAGGTAGTTATATGGCCCGTTCAGCTAAACCCCGTAAACGCAAACCCGCACCACAAAGAAGCAAACTTCCCCGCTATGTTGTGAAACTTCATCCGGATGATTTTTTTGACGAAGAAGACGCTGAAGTTCTGCGCTTTGATAATTTTGACGATGCCGTTGAGTGCTGCGCTGACCTGGGTATTCCGTTCTTTCTGGATGCAGGAAACAAAAAGCTGGTCTTCTGGTTTGTTCGTGTCGATGACGAAGGGTATCCGGAAATAGCCCGCTGTACGGAGCGGGAGTTTGCAACCATTCTTGCCGGTATCAGTGCCGGTGGTATGTACTGCCCGGAATGCGGCACTGTTCACTGGCCGGATGGCGTTACCCCACCCGTCTGATGCTTCCCCGTTTTGCCGATATTTTTCAGCAGGGTAACCGCTGGCTTAACTGGCTGGAGAAACAGCCGGAAGGTTCAGTGCGTCCGGTGGTGACTGAGTCAGTGACAAAAATCATGGCATGCGGGACCACGCTGATGGGCTACACGCAATGGTGCTGTTCGTCACCGGACTGTTGCCACACCAAAAAGGTCTGCTTCCGGTGTAAAAGCCGCTCCTGTCCGCACTGCGGGGTGAAGGCTGGCGCACAGTGGATACAGTATCTGCTGAGCCTGGTCCCCGACTGCCCGTGGCAGCATATTGTGTTCACACTTCCCTGCCAGTACTGGTCCCTGGTGTTCCACAACCGGTGGTTACTGGCAGAGATGAGCCGCATTGCAGCGGATGTGATACTGGAAATCTGCCATCAGACAGATGTGGAGCCGGGGATATTCACGGTGATCCACACATGGGGGCGTGACCAGCAGTGGCATCCGCATATCCATTTATCGACAACTGCCGGTGGTGTGACGTCGGGCCACACCTGGAAAAATCTTCATTTTTACGCCCGTAAGGTGATGAGCATGTGGCGTTACCGGATAACGCGGCTACTGTCCCGGAAATACCCGGAGCTGGTAATACTGGATGAACTGGCAGTGGAAGGAAACAGCAAACGGGACTGGAATCGCTTCCTGGACACGCATTACCGCCGCGGCTGGAATGTCAACATATCCAGGGTGATGGATAACGCCACACATGTGGCGGTGTACTTCGGCTCTTACCTGAAAAAGCCACCGGTGCCGATGAGTCGGCTGGAGCATTATGCCGGTCAGGATGAAATCGGTCTGCGTTACAACAGTCACCGTACAAAACGGGAAGAATACCTGTTGATGAGTGGAGATGAGTTCATGGAAAGGTTCTCCTGGCATGTAGCAGATAAGGGGTTCCGTATGGTGAGGTACTACGGTTTCCTGAGTCCGGTGAAGCGCCGCTTACTGGAAGAAGTTGTGTACGTCATAACGGAGACGGTGAGAAAAACGGCGATGCAAATCAGGTGGAGAGGGATGTATCAGAGGTTACTGAAGGTTGACCCGCTGAAGTGCATTCTGTGCGGAAGTCAGATGCGTTTTACGGGGCTGAAGCGGGGTTACCGACTGGCAGAGCTGGTCCTGATGCATGAGCGACTGGCACGACAGCAGGTGTGCGGCTGAGAGCCGCAGAGGGGAAGTTGCGTCCATTTTACCGGAAACGGAGCAAAAAACCGCCATTCATACCCTGTATCAATCAGTGTCATCCTGTTTAATAGTCGTTTCCGCTCATATGGTGCACAAGGGGTGTTGAAGAAACATCCGTTTTGTGGTGCTTTTTTAGTCTTTTGGGGATTTAAATTCCTATCGAT

>IS91-V29

CGAGTAGGCAGCCTGGCGGCTGCGGCTTGTCATGGTCTGGAATTACCGTTATAAAAAAAGATAATGTCATTGTCTTTCAGGTAGTTATATGGCCCGTTCAGCTAAACCCCGTAAACGCAAACCCGCACCACAAAGAAGCAAACTTCCCCGCTATGTTGTGAAACTTCATCCGGATGATTTTTTTGACGAAGAAGACGCTGAAGTTCTGCGCTTTGATAATTTTGACGATGCCGTTGAGTGCTGCGCTGACCTGGGTATTCCGTTCTTTCTGGATGCAGGAAACAAAAAGCTGGTCTTCTGGTTTGTTCGTGTCGATGACGAAGGGTATCCGGAAATAGCCCGCTGTACGGAGCGGGAGTTTGCAACCATTCTTGCCGGTATCAGTGCCGGTGGTATGTACTGCCCGGAATGCGGCACAGTTCACTGGCCGGATGGCGTTACCCCACCCGTCTGATGCTTCCCCGTTTTGCCGATATTTTTCAGCAGGGTAACCGCTGGCTTAACTGGCTGGAGAAACAGCCGGAAGGTTCAGTGCGTCCGGTGGTGACTGAGTCAGTGACAAAAATCATGGCATGCGGGACCACGCTGATGGGCTACACGCAATGGTGCTGTTCGTCACCGGACTGTTGCCACACCAAAAAGGTCTGCTTCCGGTGTAAAAGCCGCTCCTGTCCGCACTGCGGGGTGAAGGCTGGCGCACAGTGGATACAGTATCTGCTGAGCCTGGTCCCCGACTGCCCGTGGCAGCATATTGTGTTCACACTTCCCTGCCAGTACTGGTCCCTGGTGTTCCACAACCGGTGGTTACTGGCAGAGATGAGCCGCATTGCAGCGGATGTGATACTGGAAATCTGCCATCAGACAGATGTGGAGCCGGGGATATTCACGGTGATCCACACATGGGGGCGTGACCAGCAGTGGCATCCGCATATCCATTTATCGACAACAGCCGGTGGTGTGACGTCGGGCCACACCTGGAAAAATCTTCATTTTTACGCGCGTAAGGTGATGAGCATGTGGCGTTACCGGATAACGCGGCTACTGTCCCGGAAATACCCGGAGCTGGTAATACCGGATGAACTGGCAGTGGAAGGAAACAGCAAACGGGACTGGAATCGCTTCCTGGACACGCATTACCGCCGCGGCTGGAATGTCAACATATCCAGGGTGATGGATAACGCCACACATGTGGCGGTGTACTTCGGCTCTTACCTGAAAAAGCCACCGGTGCCGATGAGTCGGCTGGAGCATTATGCCGGTCAGGATGAAATCGGTCTGCGTTACAACAGTCACCGTACAAAACGGGAAGAATACCTGTTGATGAGTGGAGATGAGTTCATGGAAAGGTTCTCCTGGCATGTAGCAGATAAGGGGTTCCGTATGGTGAGGTACTACGGTTTCCTGAGTCCGGTGAAGCGCCGCTTACTGGAAGAAGTTGTGTACGTCATAACGGAGACGGTGAGAAAAACGGCGATGCAAATCAGGTGGAGAGGGATGTATCAGAGGTTACTGAAGGTTGACCCGCTGAAGTGCATTCTGTGCGGAAGTCAGATGCGTTTTACGGGGCTGAAGCGGGGTTACCGACTGGCAGAGCTGGTCCTGATGCATGAGCGACTGGCACGACAGCAGGTGTGCGGCTGAGAGCCGCAGAGGGGAAGTTGCGTCCATTTTACCGGAAACGGAGCAAAAAACCGCCATTCATACCCTGTATCAATCAGTGTCATCCTGTTTAATAGTCGTTTCCGCTCATATGGTGCACAAGGGGTGTTGAAGAAACATCCGTTTTGTGGTGCTTTTTTAGTCTTTTGGGGATTTAAATTCCTATCGAT

>IS91-V30

CGAGTAGGCAGCCTGGCGGCTGCGGCTTGTCATGGTCTGGAATTACCGTTATAAAAAAAGATAATGTCATTGTCTTTCAGGTAGTTATATGGCCCGTTCAGCTAAACCCCGTAAACGCAAACCCGCACCACAAAGAAGCAAACTTCCCCGCTATGTTGTGAAACTTCATCCGGATGATTTTTTTGACGAAGAAGACGCTGAAGTTCTGCGCTTTGATAATTTTGACGATGCCGTTGAGTGCTGCGCTGACCTGGGTATTCCGTTCTTTCTGGATGCAGGAAACAAAAAGCTGGTCTTCTGGTTTGTTCGTGTCGATGACGAAGGGTATCCGGAAATAGCCCGCTGTACGGAGCGGGAGTTTGCAACCATTCTTGCCGGTATCAGTGCCGGTGGTATGTACTGCCCGGAATGCGGCACAGTTCACTGGCCGGATGGCGTTACCCCACCCGTCTGATGCTTCCCCGTTTTGCCGATATTTTTCAGCAGGGTAACCGCTGGCTTAACTGGCTGGAGAAACAGCCGGAAGGTTCAGTGCGTCCGGTGGTGACTGAGTCAGTGACAAAAATCATGGCATGCGGGACCACGCTGATGGGCTACACGCAATGGTGCTGTTCGTCACCGGACTGTTGCCACACCAAAAAGGTCTGCTTCCGGTGTAAAAGCCGCTCCTGTCCGCACTGCGGGGTGAAGGCTGGCGCACAGTGGATACAGTATCTGCTGAGCCTGGTCCCCGACTGCCCGTGGCAGCATATTGTGTTCACACTTCCCTGCCAGTACTGGTCCCTGGTGTTCCACAACCGGTGGTTACTGGCAGAGATGAGCCGCATTGCAGCGGATGTGATACTGGAAATCTGCCATCAGACAGATGTGGAGCCGGGGATATTCACGGTGATCCACACATGGGGGCGTGACCAGCAGTGGCATCCGCATATCCATTTATCGACAACTGCCGGTGGTGTGACGTCGGGCCACACCTGGAAAAATCTTCATTTTTACGCGCGTAAGGTGATGAGCATGTGGCGTTACCGGATAACGCGGCTACTGTCCCGGAAATACCCGGAGCTGGTAATACCGGATGAACTGGCAGTGGAAGGAAACAGCAAACGGGACTGGAATCGCTTCCTGGACACGCATTACCGCCGCGGCTGGAATGTCAACATATCCAGGGTGATGGATAACGCCACACATGTGGCGGTGTACTTCGGCTCTTACCTGAAAAAGCCACCGGTGCCGATGAGTCGGCTGGAGCATTATGCCGGTCAGGATGAAATCGGTCTGCGTTACAACAGTCACCGTACAAAACGGGAAGAATACCTGTTGATGAGTGGAGATGAGTTCATGGAAAGGTTCTCCTGGCATGTAGCAGATAAGGGGTTCCGTATGGTGAGGTACTACGGTTTCCTGAGTCCGGTGAAGCGCCGCTTACTGGAAGAAGTTGTGTACGTCATAACGGAGACGGTGAGAAAAACGGCGATGCAAATCAGGTGGAGAGGGATGTATCAGAGGTTACTGAAGGTTGACCCGCTGAAGTGCATTCTGTGCGGAAGTCAGATGCGTTTTACGGGGCTGAAGCGGGGTTACCGACTGGCAGAGCTGGTCCTGATGCATGAGCGACTGGCACGACAGCAGGTGTGCGGCTGAGAGCCGCAGAGGGGAAGTTGCGTCCATTTTACCGGAAACGGAGCAAAAAACCGCCATTCATACCCTGTATCAATCAGTGTCATCCTGTTTAATAGTCGTTTCCGCTCATATGGTGCACAAGGGGTGTTGAAGAAACATTCGTTTTGTGGTGCTTTTTTAGTCTTTTGGGGATTTAAATTCCTATCGAT

>IS91-V31

CGAGTAGGCAGCCTGGCGGCTGCGGCTTGTCATGGTCTGGAATTACCGTTATAAAAAAAGATAATGTCATTGTCTTTCAGGTAGTTATATGGCCCGTTCAGCTAAACCCCGTAAACGCAAACCCGCACCACAAAGAAGCAAACTTCCCCGCTATGTTGTGAAACTTCATCCGGATGATTTTTTTGACGAAGAAGACGCTGAAGTTCTGCGCTTTGATAATTTTGACGATGCCGTTGAGTGCTGCGCTGACCTGGGTATTCCGTTCTTTCTGGATGCAGGAAACAAAAAGCTGGTCTTCTGGTTTGTTCGTGTCGATGACGAAGGGTATCCGGAAATAGCCCGCTGTACGGAGCGGGAGTTTGCAACCATTCTTGCCGGTATCAGTGCCGGTGGTATGTACTGCCCGGAATGCGGCACAGTTCACTGGCCGGATGGCGTTACCCCACCCGTCTGATGCTTCCCCGTTTTGCCGATATTTTTCAGCAGGGTAACCGCTGGCTTAACTGGCTGGAGAAACAGCCGGAAGGTTCAGTGCGTCCGGTGGTGACTGAGTCAGTGACAAAAATCATGGCATGCGGGACCACGCTGATGGGCTACACGCAATGGTGCTGTTCGTCACCGGACTGTTGCCACACCAAAAAGGTCTGCTTCCGGTGTAAAAGCCGCTCCTGTCCGCACTGCGGGGTGAAGGCTGGCGCACAGTGGATACAGTATCTGCTGAGCCTGGTCCCCGACTGCCCGTGGCAGCATATTGTGTTCACACTTCCCTGCCAGTACTGGTCCCTGGTGTTCCACAACCGGTGGTTACTGGCAGAGATGAGCCGCATTGCAGCGGATGTGATACTGGAAATCTGCCATCAGACAGATGTGGAGCCGGGGATATTCACGGTGATCCACACATGGGGGCGTGACCAGCAGTGGCATCCGCATATCCATTTATCGACAACTGCCGGTGGTGTGACGTCGGGCCACACCTGGAAAAATCTTCATTTTTACGCCCGTAAGGTGATGAGCATGTGGCGTTACCGGATAACGCGGCTACTGTCCCGGAAATACCCGGAGCTGGTAATACCGGATGAACTGGCAGTGGAAGGAAACAGCAAACGGGACTGGAATCGCTTCCTGGACACGCATTACCGCCGCGGCTGGAATGTCAACATATCCAGGGTGATGGATAACGCCACACATGTGGCGGTGTACTTCGGCTCTTACCTGAAAAAGCCACCGGTGCCGATGAGTCGGCTGGAGCATTATGCCGGTCAGGATGAAATCGGTCTGCGTTACAACAGTCACCGTACAAAACGGGAAGAATACCTGTTGATGAGTGGAGATGAGTTCATGGAAAGGTTCTCCTGGCATGTAGCAGATAAGGGGTTCCGTATGGTGAGGTACTACGGTTTCCTGAGTCCGGTGAAGCGCCGCTTACTGGAAGAAGTTGTGTACGTCATAACGGAGACGGTGAGAAAAACGGCGATGCAAATCAGGTGGAGAGGGATGTATCAGAGGTTACTGAAGGTTGACCCGCTGAAGTGCATTCTGTGCGGAAGTCAGATGCGTTTTACGGGGCTGAAGCGGGGTTACCGACTGGCAGAGCTGGTCCTGATGCATGAGCGACTGGCACGACAGCAGGTGTGCGGCTGAGAGCCGCAGAGGGGAAGTTGCGTCCATTTTACCGGAAACGGAGCAAAAAACCGCCATTCATACCCTGTATCAATCAGTGTCATCCTGTTTAATAGTCGTTTCCGCTCATATGGTGCACAAGGGGTGTTGAAGAAATATCCGTTTTGTGGTGCTTTTTTAGTCTTTTGGGGGTTTAAATTCCTATCGAT

>IS91-V32

CGAGTAGGCAGCCTGGCGGCTGCGGCTTGTCATGGTCTGGAATTACCGTTATAAAAAAAGATAATGTCATTGTCTTTCAGGTAGTTATATGGCCCGTTCAGCTAAACCCCGTAAACGCAAACCCGCACCACAAAGAAGCAAACTTCCCCGCTATGTTGTGAAACTTCATCCGGATGATTTTTTTGACGAAGAAGACGCTGAAGTTCTGCGCTTTGATAATTTTGACGATGCCGTTGAGTGCTGCGCTGACCTGGGTATTCCGTTCTTTCTGGATGCAGGAAACAAAAAGCTGGTCTTCTGGTTTGTTCGTGTCGATGACGAAGGGTATCCGGAAATAGCCCGCTGTACGGAGCGGGAGTTTGCAACCATTCTTGCCGGTATCAGCGCCGGCGGCATGTACTGCCCGGAGTGTGGCACGGTTCACTGGCCGGACGGAGTCCCCCCGCCCTTCTGATGCTTCCCCGTTTTGCCGACATTTTTCAGCAGGGAAACCGCTGGCTTAACTGGCTGGAGAAACAACCGGAAGGTTCAGTGCGTCCGGTAGTCATTGAGTCGGTGACAAAAATCATGGCGTGCGGAACCACGCTGATGGGGTACACACAGTGGTGCTGTTCATCTCCGGACTGTTGCCACACAAAAAAGGTCTGCTTCCGGTGTAAAAGTCGCTCCTGCCCGCACTGCGGAGTGAAGGCTGGCGCACAGTGGATACAGTATCTGCTGAGCCTGGTCCCCGACTGCCCGTGGCAGCATATTGTGTTCACACTTCCCTGCCAGTACTGGTCCCTGGTGTTCCACAACCGGTGGTTACTGGCAGAGATGAGCCGCATTGCAGCGGATGTGATACTGGAAATCTGCCATCAGACAGATGTGGAGCCGGGGATATTCACGGTGATCCACACATGGGGGCGTGACCAGCAGTGGCATCCGCATATCCATTTATCGACAACTGCCGGTGGTGTGACGTCGGGCCACACCTGGAAAAATCTTCATTTTTACGCCCGTAAGGTGATGAGCATGTGGCGTTACCGGATAACGCGGCTACTGTCCCGGAAATACCCGGAGCTGGTAATACCGGATGAACTGGCAGTGGAAGGAAACAGCAAACGGGACTGGAATTGCTTCCTGGACACGCATTACCGCCGCGGCTGGAATGTCAACATATCCAGGGTGATGGATAACGCCACACATGTGGCGGTGTACTTCGGCTCTTACCTGAAAAAGCCACCGGTGCCGATGAGTCGGCTGGAGCATTATGCCGGTCAGGATGAAATCGGTCTGCGTTACAACAGTCACCGTACAAAACGGGAAGAATACCTGTTGATGAGTGGAGATGAGTTCATGGAAAGGTTCTCCTGGCATGTAGCAGATAAGGGGTTCCGTATGGTGAGGTACTACGGTTTCCTGAGTCCGGTGAAGCGCCGCTTACTGGAAGAAGTTGTGTACGTCATAACGGAGACGGTGAGAAAGACGGCGATGCAAATCAGGTGGAGAGGGATGTATCAGAGGTTACTGAAGGTTGACCCGCTGAAGTGCATTCTGTGCGGAAGTCAGATGCGTTTTACGGGGCTGAAGCGGGGTTACCGACTGGCAGAGCTGGTCCTGATGCATGAGCGACTGGCACGACAGCAGGTGTGCGGCTGAGAGCCGCAGAGGGGAAGTTGCGTCCATTTTACCGGAAACGGAGCAAAAAACCGCCATTCATACCCTGTATCAATCAGTGTCATCCTGTTTAATAGTCGTTTCCGCTCATATGGTGCACAAGGGGTGTTGAAGAAATATCCGTTTTGTGGTGCTTTTTTAGTCTTTTGGGGATTTAAATTCCTATCGAT

>IS91-V33

CGAGTAGGCAGCCTGGCGGCTGCGGCTTGTCATGGTCTGGAATTACCGTTATAAAAAAAGATAATGTCATTGTCTTTCAGGTAGTTATATGGCCCGTTCAGCTAAACCCCGTAAACGAAAACCCGCCCCACAAAGAAGCAAACTTCCCCGCTATGTTGTGAAACTTCATCCGGATGATTTTTTTGACGAAGAAGACGCTGAAGTTCTGCGCTTTGATAATTTTGACGATGCCGTTGAGTGCTGCGCTGACCTGGGTATTCCGTTCTTTCTGGATGCAGGAAACAAAAAGCTGGTCTTCTGGTTTGTTCGTGTCGATGACGAAGGGTATCCGGAAATAGCCCGCTGTACGGAGCGGGAGTTTGCAACCATTCTTGCCGGTATCAGTGCCGGTGGTATGTACTGCCCGGAATGCGGCACAGTTCACTGGCCGGATGGCGTTACCCCACCCGTCTGATGCTTCCCCGTTTTGCCGATATTTTTCAGCAGGGTAACCGCTGGCTTAACTGGCTGGAGAAACAGCCGGAAGGTTCAGTGCGTCCGGTGGTGACTGAGTCAGTGACAAAAATCATGGCATGCGGGACCACGCTGATGGGCTACACGCAATGGTGCTGTTCGTCACCGGACTGTTGCCACACCAAAAAGGTCTGCTTCCGGTGTAAAAGCCGCTCCTGTCCGCACTGCGGGGTGAAGGCTGGCGCACAGTGGATACAGTATCTGCTGAGCCTGGTCCCCGACTGCCCGTGGCAGCATATTGTGTTCACACTTCCCTGCCAGTACTGGTCCCTGGTGTTCCACAACCGGTGGTTACTGGCAGAGATGAGCCGCATTGCAGCGGATGTGATACTGGAAATCTGCCATCAGACAGATGTGGAGCCGGGGATATTCACGGTGATCCACACATGGGGGCGTGACCAGCAGTGGCATCCGCATATCCATTTATCGACAACAGCCGGTGGTGTGACGTCGGGCCACACCTGGAAAAATCTTCATTTTTACGCGCGTAAGGTGATGAGCATGTGGCGTTACCGGATAACGCGGCTACTGTCCCGGAAATACCCGGAGCTGGTAATACCGGATGAACTGGCAGTGGAAGGAAACAGCAAACGGGACTGGAATCGCTTCCTGGACACGCATTACCGCCGCGGCTGGAATGTCAACATATCCAGGGTGATGGATAACGCCACACATGTGGCGGTGTACTTCGGCTCTTACCTGAAAAAGCCACCGGTGCCGATGAGTCGGCTGGAGCATTATGCCGGTCAGGATGAAATCGGTCTGCGTTACAACAGTCACCGTACAAAACGGGAAGAATACCTGTTGATGAGTGGAGATGAGTTCATGGAAAGGTTCTCCTGGCATGTAGCAGATAAGGGGTTCCGTATGGTGAGGTACTACGGTTTCCTGAGTCCGGTGAAGCGCCGCTTACTGGAAGAAGTTGTGTACGTCATAACGGAGACGGTGAGAAAAACGGCGATGCAAATCAGGTGGAGAGGGATGTATCAGAGGTTACTGAAGGTTGACCCGCTGAAGTGCATTCTGTGCGGAAGTCAGATGCGTTTTACGGGGCTGAAGCGGGGTTACCGACTGGCAGAGCTGGTCCTGATGCATGAGCGACTGGCACGACAGCAGGTGTGCGGCTGAGAGCCGCAGAGGGGAAGTTGCGTCCATTTTACCGGAAACGGAGCAAAAAACCGCCATTCATACCCTGTATCAATCAGTGTCATCCTGTTTAATAGTCGTTTCCGCTCATATGGTGCACAAGGGGTGTTGAAGAAATATCCGTTTTGTGGTGCTTTTTTAGTCTTTTGGGGATTTAAATTCCTATCGAT

>IS91-V34

CGAGTAGGCAGCCTGGCGGCTGCGGCTTGTCATGGTCTGGAATTACCGTTATAAAAAAAGATAATGTCATTGTCTTTCAGGTAGTTATATGGCCCGTTCAGCTAAACCCCGTAAACGCAAACCCGCACCACAAAGAAGCAAACTTCCCCGCTATGTTGTGAAACTTCATCCGGATGATTTTTTTGACGAAGAAGACGCTGAAGTTCTGCGCTTTGATAATTTTGACGATGCCGTTGAGTGCTGCGCTGACCTGGGTATTCCGTTCTTTCTGGATGCAGGAAACAAAAAGCTGGTCTTCTGGTTTGTTCGTGTCGATGACGAAGGGTATCCGGAAATAGCCCGCTGTACGGAGCGGGAGTTTGCAACCATTCTTGCCGGTATCAGTGCCGGCGGCATGTACTGCCCGGAGTGTGGCACGGTTCACTGGCCGGACGGAGTCCCCCCGCCCTTCTGATGCTTCCCCGTTTTGCCGACATTTTTCAGCAGGGAAACCGCTGGCTTAACTGGCTGGAGAAACAACCGGAAGGTTCAGTGCGTCCGGTAGTCATTGAGTCGGTGACAAAAATCATGGCGTGCGGAACCACGCTGATGGGGTACACACAGTGGTGCTGTTCATCTCCGGACTGTTGCCACACAAAAAAGGTCTGCTTCCGGTGTAAAAGTCGCTCCTGCCCGCACTGCGGAGTGAAGGCTGGCGCACAGTGGATACAGTATCTGCTGAGCCTGGTCCCCGACTGCCCGTGGCAGCATATTGTGTTCACACTTCCCTGCCAGTACTGGTCCCTGGTGTTCCACAACCGGTGGTTACTGGCAGAGATGAGCCGCATTGCAGCGGATGTGATACTGGAAATCTGCCATCAGACAGATGTGGAGCCGGGGATATTCACGGTGATCCACACATGGGGGCGTGACCAGCAGTGGCATCCGCATATCCATTTATCGACAACTGCCGGTGGTGTGACGTCGGGCCACACCTGGAAAAATCTTCATTTTTACGCCCGTAAGGTGATGAGCATGTGGCGTTACCGGATAACGCGGCTACTGTCCCGGAAATACCCGGAGCTGGTAATACCGGATGAACTGGCAGTGGAAGGAAACAGCAAACGGGACTGGAATTGCTTCCTGGACACGCATTACCGCCGCGGCTGGAATGTCAACATATCCAGGGTGATGGATAACGCCACACATGTGGCGGTGTACTTCGGCTCTTACCTGAAAAAGCCACCGGTGCCGATGAGTCGGCTGGAGCATTATGCCGGTCAGGATGAAATCGGTCTGCGTTACAACAGTCACCGTACAAAACGGGAAGAATACCTGTTGATGAGTGGAGATGAGTTCATGGAAAGGTTCTCCTGGCATGTAGCAGATAAGGGGTTCCGTATGGTGAGGTACTACGGTTTCCTGAGTCCGGTGAAGCGCCGCTTACTGGAAGAAGTTGTGTACGTCATAACGGAGACGGTGAGAAAGACGGCGATGCAAATCAGGTGGAGAGGGATGTATCAGAGGTTACTGAAGGTTGACCCGCTGAAGTGCATTCTGTGCGGAAGTCAGATGCGTTTTACGGGGCTGAAGCGGGGTTACCGACTGGCAGAGCTGGTCCTGATGCATGAGCGACTGGCACGACAGCAGGTGTGCGGCTGAGAGCCGCAGAGGGGAAGTTGCGTCCATTTTACCGGAAACGGAGCAAAAAACCGCCATTCATACCCTGTATCAATCAGTGTCATCCTGTTTAATAGTCGTTTCCGCTCATATGGTGCACAAGGGGTGTTGAAGAAATATCCGTTTTGTGGTGCTTTTTTAGTCTTTTGGGGATTTAAATTCCTATCGAT

>IS91-V35

CGAGTAGGCAGCCTGGCGGCTGCGGCTTGTCATGGTCTGGAATTACCGTTATAAAAAAGATAATGTCATTGTCTTTCAGGTAGTTATATGGCCCGTTCAGCTAAACCCCGTAAACGCAAACCCGCACCACAAAGAAGCAAACTTCCCCGCTATGTTGTGAAACTTCATCCGGATGATTTTTTTGACGAAGAAGACGCTGAAGTTCTGCGCTTTGATAATTTTGACGATGCCGTTGAGTGCTGCGCTGACCTGGGTATTCCGTTCTTTCTGGATGCAGGAAACAAAAAGCTGGTCTTCTGGTTTGTTCGTGTCGATGACGAAGGGTATCCGGAAATAGCCCGCTGTACGGAGCGGGAGTTTGCAACCATTCTTGCCGGTATCAGTGCCGGTGGTATGTACTGCCCGGAATGCGGCACAGTTCACTGGCCGGATGGCGTTACCCCCCCGTCTGATGCTTCCCCGTTTTGCCGATATTTTTCAGCAGGGTAACCGCTGGCTTAACTGGCTGGAGAAACAGCCGGAAGGTTCAGTGCGTCCGGTGGTGACTGAGTCAGTGACAAAAATCATGGCATGCGGGACCACGCTGATGGGCTACACGCAATGGTGCTGTTCGTCACCGGACTGTTGCCACACCAAAAAGGTCTGCTTCCGGTGTAAAAGCCGCTCCTGTCCGCACTGCGGGGTGAAGGCTGGCGCACAGTGGATACAGTATCTGCTGAGCCTGGTCCCCGACTGCCCGTGGCAGCATATTGTGTTCACACTTCCCTGCCAGTACTGGTCCCTGGTGTTCCACAACCGGTGGTTACTGGCAGAGATGAGCCGCATTGCAGCGGATGTGATACTGGAAATCTGCCATCAGACAGATGTGGAGCCGGGGATATTCACGGTGATCCACACATGGGGGCGTGACCAGCAGTGGCATCCGCATATCCATTTATCGACAACTGCCGGTGGTGTGACGTCGGGCCACACCTGGAAAAACCTTCATTTTTACGCCCGTAAGGTGATGAGCATGTGGCGTTACAGGATAACGTGGTTACTGTCACGAAAATACCCGGAGCTGGTAATACCGGATGAACTGGCAGTGGAAGGAAACAGCAAACGGGACTGGAATTGCTTCCTGGACACGCATTACCGCCGCGGCTGGAATGTCAACATATCCAGGGTGATGGATAACGCCACACATGTGGCGGTGTACTTCGGCTCTTACCTGAAAAAGCCACCGGTGCCGATGAGTCGGCTGGAGCATTATGCCGGTCAGGATGAAATCGGTCTGCGTTACAACAGTCACCGTACAAAACGGGAAGAATACCTGTTGATGAGTGGAGATGAGTTCATGGAAAGGTTCTCCTGGCATGTAGCAGATAAGGGGTTCCGTATGGTGAGGTACTACGGTTTCCTGAGTCCGGTGAAGCGCCGCTTACTGGAAGAAGTTGTGTACGTCATAACGGAGACGGTGAGAAAAACGGCGATGCAAATCAGGTGGAGAGGGATGTATCAGAGGTTACTGAAGGTTGACCCGCTGAAGTGCATTCTGTGCGGAAGTCAGATGCGTTTTACGGGGCTGAAGCGGGGTTACCGACTGGCAGAGCTGGTCCTGATGCATGAGCGACTGGCACGACAGCAGGTGTGCGGCTGAGAGCCGCAGAGGGGAAGTTGCGTCCATTTTACCGGAAACGGAGCAAAAAACCGCCATTCATACCCTGTATCAATCAGTGTCATCCTGTTTAATAGTCGTTTCCGCTCATATGGTGCACAAGGGGTGTTGAAGAAACATCCGTTTTGTGGTGCTTTTTTAGTCTTTTGGGGATTTAAATTCCTATCGAT

>IS91-V36

CGAGTAGGCAGCCTGGCGGCTGCGGCTTGTCATGGTCTGGAATTACCGTTATAAAAAAAGATAATGTCATTGTCTTTCAGGTAGTTATATGGCCCGTTCAGCTAAACCCCGTAAACGCAAACCCGCACCACAAAGAAGCAAACTTCCCCGCTATGTTGTGAAACTTCATCCGGATGATTTTTTTGACGAAGAAGACGCTGAAGTTCTGCGCTTTGATAATTTTGACGATGCCGTTGAGTGCTGCGCTGACCTGGGTATTCCGTTCTTTCTGGATGCAGGAAACAAAAAGCTGGTCTTCTGGTTTGTTCGTGTCGATGACGAAGGGTATCCGGAAATAGCCCGCTGTACGGAGCGGGAGTTTGCAACCATTCTTGCCGGTATCAGTGCCGGTGGTATGTACTGCCCGGAATGCGGCACAGTTCACTGGCCGGATGGCGTTACCCCACCCGTCTGATGCTTCCCCGTTTTGCCGATATTTTTCAGCAGGGTAACCGCTGGCTTAACTGGCTGGAGAAACAGCCGGAAGGTTCAGTGCGTCCGGTGGTGACTGAGTCAGTGACAAAAATCATGGCATGCGGGACCACGCTGATGGGCTACACGCAATGGTGCTGTTCGTCACCGGACTGTTGCCACACCAAAAAGGTCTGCTTCCGGTGTAAAAGCCGCTCCTGTCCGCACTGCGGGGTGAAGGCTGGCGCACAGTGGATACAGTATCTGCTGAGCCTGGTCCCCGACTGCCCGTGGCAGCATATTGTGTTCACACTTCCCTGCCAGTACTGGTCCCTGGTGTTCCACAACCGGTGGTTACTGGCAGAGATGAGCCGCATTGCTGCGGATGTGATACTGGAAATCTGCCATCAGACAGATGTGGAGCCGGGGATATTCACGGTGATCCACACATGGGGGCGTGACCAGCAGTGGCATCCGCATATCCATTTATCGACAACTGCCGGTGGTGTGACGTCGGGCCACACCTGGAAAAATCTTCATTTTTACGCCCGTAAGGTGATGAGCATGTGGCGTTACCGGATAACGCGGCTACTGTCCCGGAAATACCCGGAGCTGGTAATACCGGATGAACTGGCAGTGGAAGGAAACAGCAAACGGGACTGGAATCGCTTCCTGGACACGCATTACCGGCGGGGCTGGAATGTCAACATATCCAGGGTGATGGATAACGCCACACATGTGGCGGTGTACTTCGGCTCTTACCTGAAAAAGCCACCGGTGCCGATGAGTCGGCTGGAGCATTATGCCGGTCAGGATGAAATCGGTCTGCGTTACAACAGTCACCGTACAAAACGGGAAGAATACCTGTTGATGAGTGGAGATGAGTTCATGGAAAGGTTCTCCTGGCATGTAGCAGATAAGGGGTTCCGTATGGTGAGGTACTACGGTTTTCCTGAGTCCGGTGAACGCCGCTTACTGGAAGAAGTTGTGTACGTCATAACGGAGACGGTGAGAAAAACGGCGATGCAAATCAGGTGGAGAGGGATGTATCAGAGGTTACTGAAGGTTGACCCGCTGAAGTGCATTCTGTGCGGAAGTCAGATGCGTTTTACGGGGCTGAAGCGGGGTTACCGACTGGCAGAGCTGGTCCTGATGCATGAGCGACTGGCACGACAGCAGGTGTGCGGCTGAGAGCCGCATAGGGGAAGTTGCGTCCATTTTACCGGAAACGGAGCAAAAAACCGTCATTCATACCCTGTATCAATCAGTGTCATCCTGTTTAATAGTCGTTTCCGCTCATATGGTGCACAAGGGGTGTTGAAGAAACATCCGTTTTGTGGTGCTTTTTTAGTCTTTTGGGGATTTAAATTCCTATCGAT

>IS91-V37

CGAGTAGGCAGCCTGGCGGCTGCGGCTTGTCATGGCCTGGAATTACCGTTATAAAAAAAGATAATGTCATTGTCTTTCAGGTAGTTATATGGCCCGTTCAGCTAAACCCCGTAAACGCAAACCCGCACCACAAAGAAGCAAACTTCCCCGCTATGTTGTGAAACTTCATCCGGATGATTTTTTTGACGAAGAAGACGCTGAAGTTCTGCGCTTTGATAATTTTGACGATGCCGTTGAGTGCTGCGCTGACCTGGGTATTCCGTTCTTTCTGGATGCAGGAAACAAAAAGCTGGTCTTCTGGTTTGTTCGTGTCGATGACGAAGGGTATCCGGAAATAGCCCGCTGTACGGAGCGGGAGTTTGCAACCATTCTTGCCGGTATCAGTGCCGGTGGTATGTACTGCCCGGAATGCGGCACAGTTCACTGGCCGGATGGCGTTACCCCACCCGTCTGATGCTTCCCCGTTTTGCCGATATTTTTCAGCAGGGTAACCGCTGGCTTAACTGGCTGGAGAAACAGCCGGAAGGTTCAGTGCGTCCGGTGGTGACTGAGTCAGTGACAAAAATCATGGCATGCGGGACCACGCTGATGGGCTACACGCAATGGTGCTGTTCGTCACCGGACTGTTGCCACACCAAAAAGATCTGCTTCCGGTGTAAAAGCCGCTCCTGTCCGAACTGCGGGGTGAAGGCTGGCGCACAGTGGATACAGTATCTGCTGAGCCTGGTCCCCGACTGCCCGTGGCAGCATATTGTGTTCACACTTCCCTGCCAGTACTGGTCCCTGGTGTTCCACAACCGGTGGTTACTGGCAGAGATGAGCCGCATTGCAGCGGATGTGATACTGGAAATCTGCCATCAGGCAGATGTGGAGCCGGGGATATTCACGGTGATCCACACATGGGGGCGTGACCAGCAGTGGCATCCGCATATCCATTTATCGACAACTGCCGGTGGTGTGACGTCGGGCCACACCTGGAAAAATCTTCATTTATACGCCCGTAAGGTGATGAGCATGTGGCGCTACCGGATAACGCGGCTACTGTCCCGGAAATACCCGGAGCTGGTAATACCGGATGAACTGGCAGTGGAAGGAAACAGCAAACGGGACTGGAATCGCTTCCTGGACACGCATTACCGCCGCGGCTGGAATGTCAACATATCCAGGGTGATGGATAACGCCACACATGTGGCGGTGTACTTCGGCTCTTACCTGAAAAAGCCACCGGTGCCGATGAGTCGGCTGGAGCATTATGCCGGTCAGGATGAAATCGGTCTGCGTTACAACAGTCACCGTACAAAACGGGAAGAATACCTGTTGATGAGTGGAGATGAGTTCATGGAAAGGTTCTCCTGGCATGTAGCAGATAAGGGGTTCCGTATGGTGAGGTACTACGGTTTCCTGAGTCCGGTGAAGCGCCGCTTACTGGAAGAAGTTGTGTACGTCATAACGGAGACGGTGAGAAAAACGGCGATGCAAATCAGGTGGAGAGGGATGTATCAGAGGTTACTGAAGGTTGACCCGCTGAAGTGCATTCTGTGCGGAAGTCAGATGCGTTTTACGGGGCTGAAGCGGGGTTACCGACTGGCAGAGCTGGTCCTGATGCATGAGCGACTGGCACGACAGCAGGTGTGCGGCTGAGAGTCGCAGAGGGGAAGTTGCGTCCATTTTACCGGAAACGGAGCAAAAAACCGCCATTCATACCCTGTATCAATCAGTGTCATCCTGTTTAATAGTCGTTTCCGCTCATATGGTGCACAAGGGGTGTTGAAGAAATATCCGTTTTGTGGTGCTTTTTTAGTCTTTTGGGGATTTAAATTCCTATCGAT

>IS91-V38

CGAGTAGGCAGCCTGGCGGCTGCGGCTTGTCATGGCCTGGAATTACCATTATAAAAAAAGATAATGTCATTGTCTTTCAGGTAGTTATATGGCCCGTTCAGCTAAACCCCGTAAACGCAAACCCGCACCACAAAGAAGCAAACTTCCCCGCTATGTTGTGAAACTTCATCCGGATGATTTTTTTGACGAAGAAGACGCTGAAGTTCTGCGCTTTGATAATTTTGACGATGCCGTTGAGTGCTGCGCTGACCTGGGTATTCCGTTCTTTCTGGATGCAGGAAACAAAAAGCTGGTCTTCTGGTTTGTTCGTGTCGATGACGAAGGGTATCCGGAAATAGCCCGCTGTACGGAGCGGGAGTTTGCAACCATTCTTGCCGGTATCAGTGCCGGTGGTATGTACTGCCCGGAATGCGGCACAGTTCACTGGCCGGATGGCGTTCCCCCACCCGTCTGATGCTTCCCCGTTTTGCCGATATTTTTCAGCAGGGTAACCGCTGGCTTAACTGGCTGGAGAAACAGCCGGAAGGTTCAGTGCGTCCGGTGGTGACTGAGTCAGTGACAAAAATCATGGCATGCGGGACCACGCTGATGGGCTACACGCAATGGTGCTGTTCGTCACCGGACTGTTGCCACACCAAAAAGGTCTGCTTCCGGTGTAAAAGCCGCTCCTGTCCGCACTGCGGGGTGAAGGCTGGCGCACAGTGGATACAGTATCTGCTGAGCCTGGTCCCCGACTGCCCGTGGCAGCATATTGTGTTCACACTTCCCTGCCAGTACTGGTCCCTGGTGTTCCACAACCGGTGGTTACTGGCAGAGATGAGCCGCATTGCAGCTGATGTGATACTGGAAATCTGCCATCAGGCAGATGTTGAGCCGGGGATATTCACGGTGATCCACACATGGGGGCGTGACCAGCAGTGGCATCCGCATATCCATTTATCGACAACTGCCGGTGGTGTGACGTCGGGCCACACCTGGAAAAACCTTCATTTTTACGCCCGTAAGGTGATGAGCATGTGGCGTTACCGGATAACGCGGCTACTGTCCCGGAAATACCCGGACCTGGTAATACCGGATGAACTGGCAGTGGAAGGAAACAGCAAACGGGACTGGAATCGCTTCCTGGACACGCATTACCGCCGCGGCTGGAATGTCAACATATCCAGGGTGATGGATAACGCCACACATGTGGCGGTGTACTTCGGCTCTTACCTGAAAAAGCCACCGGTGCCGATGAGTCGGCTGGAGCATTATGCCGGTCAGGATGAAATCGGTCTGCGTTACAACAGTCACCGTACAAAACGGGAAGAATACCTGTTGATGAGTGGAGATGAGTTCATGGAAAGGTTCTCCTGGCATGTAGCAGATAAGGGGTTCCGTATGGTGAGGTACTACGGTTTCCTGAGTCCGGTGAAGCGCCGCTTACTGGAAGAAGTTGTGTACGTCATAACGGAGACGGTGAGAAAAACGGCGATGCAAATCAGGTGGAGAGGGATGTATCAGAGGTTACTGAAGGTTGACCCGCTGAAGTGCATTCTGTGCGGAAGTCAGATGCGTTTTACGGGGCTGAAACGGGGTTACCGACTGGCAGAGCTGGTCCTGATGCATGAGCGACTGGCACGACAGCAGGTGTGCGGCTGAGAGCCGCAGAGGGGAAGTTGCGTCCATTTTACCGGAAACGGAGCAAAAAACCGCCATTCATACCCTGTATCAATCAGTGTCATCCTGTTTAATAGTCGTTTCCGCTCATATGGTGCACAAGGGGTGTTGAAGAAACATCCGTTTCGTGGTGCTTTTTTAGTCTTTTGGGGATTTAAATTCCTATCGAT

>IS91-V39

CGAGTAGGCAGCCTGGCGGCTGCGGCTTGTCATGGCCTGGAATTACCGTTATAAAAAAAGATAATGTCATTGTCTTTCAGGTAGTTATATGGCCCGTTCAGCTAAACCCCGTAAACGCAAACCCGCACCACAAAGAAGCAAACTTCCCCGCTATGTTGTGAAACTTCATCCGGATGATTTTTTTGACGAAGAAGACGCTGAAGTTCTGCGCTTTGATAATTTTGACGATGCCGTTGAGTGCTGCGCTGACCTGGGTATTCCGTTCTTTCTGGATGCAGGAAACAAAAAGCTGGTCTTCTGGTTTGTTCGTGTCGATGACGAAGGGTATCCGGAAATAGCCCGCTGTACGGAGCGGGAGTTTGCAACCATTCTTGCCGGTATCAGTGCCGGTGGTATGTACTGCCCGGAATGCGGCACAGTTCACTGGCCGGATGGCGTTACCCCACCCGTCTGATGCTTCCCCGTTTTGCCGATATTTTTCAGCAGGGTAACCGCTGGCTTAACTGGCTGGAGAAACAGCCGGAAGGTTCAGTGCGTCCGGTGGTGACTGAGTCAGTGACAAAAATCATGGCATGCGGGACCACGCTGATGGGCTACACGCAATGGTGCTGTTCGTCACCGGACTGTTGCCACACCAAAAAGATCTGCTTCCGGTGTAAAAGCCGCTCCTGTCCGAACTGCGGGGTGAAGGCTGGCGCACAGTGGATACAGTATCTGCTGAGCCTGGTCCCCGACTGCCCGTGGCAGCATATTGTGTTCACACTTCCCTGCCAGTACTGGTCCCTGGTGTTCCACAACCGGTGGTTACTGGCAGAGATGAGCCGCATTGCAGCGGATGTGATACTGGAAATCTGCCATCAGGCAGATGTGGAGCCGGGGATATTCACGGTGATCCACACATGGGGGCGTGACCAGCAGTGGCATCCGCATATCCATTTATCGACAACTGCCGGTGGTGTGACGTCGGGCCACACCTGGAAAAATCTTCATTTATACGCCCGTAAGGTGATGAGCATGTGGCGCTACCGGATAACGCGGCTACTGTCCCGGAAATACCCGGAGCTGGTAATACCGGATGAACTGGCAGTGGAAGGAAACAGCAAACGGGACTGGAATCGCTTCCTGGACACGCATTACCGCCGCGGCTGGAATGTCAACATATCCAGGGTGATGGATAACGCCACACATGTGGCGGTGTACTTCGGCTCTTACCTGAAAAAGCCACCGGTGCCGATGAGTCGGCTGGAGCATTATGCCGGTCAGGATGAAATCGGTCTGCGTTACAACAGTCACCGTACAAAACGGGAAGAATACCTGTTGATGAGTGGAGATGAGTTCATGGAAAGGTTCTCCTGGCATGTAGCAGATAAGGGGTTCCGTATGGTGAGGTACTACGGTTTCCTGAGTCCGGTGAAGCGCCGCTTACTGGAAGAAGTTGTGTACGTCATAACGGAGACGGTGAGAAAAACGGCGATGCAAATCAGGTGGAGAGGGATGTATCAGAGGTTACTGAAGCTTGACCCGCTGAAGTGCATTCTGTGCGGAAGTCAGATGCGTTTTACGGGGCTGAAGCGGGGTTACCGACTGGCAGAGCTGGTCCTGATGCATGAGCGACTGGCACGACAGCAGGTGTGCGGCTGAGAGTCGCAGAGGGGAAGTTGCGTCCATTTTACCGGAAACGGAGCAAAAAACCGCCATTCATACCCTGTATCAATCAGTGTCATCCTGTTTAATAGTCGTTTCCGCTCATATGGTGCACAAGGGGTGTTGAAGAAATATCCGTTTTGTGGTGCTTTTTTAGTCTTTTGGGGATTTAAATTCCTATCGAT

>IS91-V40

CGAGTAGGCAGCCTGGCGGCTGCGGCTTGTCATGGCCTGGAATTACCGTTATAAAAAAAGATAATGTCATTGTCTTTCAGGTAGTTATATGGCCCGTTCAGCTAAACCCCGTAAACGCAAACCCGCACCACAAAGAAGCAAACTTCCCCGCTATGTTGTGAAACTTCATCCGGATGATTTTTTTGACGAAGAAGACGCTGAAGTTCTGCGCTTTGATAATTTTGACGATGCCGTTGAGTGCTGCGCTGACCTGGGTATTCCGTTCTTTCTGGATGCAGGAAACAAAAAGCTGGTCTTCTGGTTTGTTCGTGTCGATGACGAAGGGTATCCGGAAATAGCCCGCTGTACGGAGCGGGAGTTTGCAACCATTCTTGCCGGTATCAGTGCCGGTGGTATGTACTGCCCGGAATGCGGCACAGTTCACTGGCCGGATGGCGTTACCCCACCCGTCTGATGCTTCCCCGTTTTGCCGATATTTTTCAGCAGGGTAACCGCTGGCTTAACTGGCTGGAGAAACAGCCGGAAGGTTCAGTGCGTCCGGTGGTGACTGAGTCAGTGACAAAAATCATGGCATGCGGGACCACGCTGATGGGCTACACGCAATGGTGCTGTTCGTCACCGGACTGTTGCCACACCAAAAAGATCTGCTTCCGGTGTAAAAGCCGCTCCTGTCCGAACTGCGGGGTGAAGGCTGGCGCACAGTGGATACAGTATCTGCTGAGCCTGGTCCCCGACTGCCCGTGGCAGCATATTGTGTTCACACTTCCCTGCCAGTACTGGTCCCTGGTGTTCCACAACCGGTGGTTACTGGCAGAGATGAGCCGCATTGCAGCGGATGTGATACTGGAAATCTGCCATCAGGCAGATGTGGAGCCGGGGATATTCACGGTGATCCACACATGGGGGCGTGACCAGCAGTGGCATCCGCATATCCATTTATCGACAACTGCCGGTGGTGTGACGTCGGGCCACACCTGGAAAAATCTTCATTTATACGCCCGTAAGGTGATGAGCATGTGGCGCTACCGGATAACGCGGCTACTGTCCCGGAAATACCCGGAGCTGGTAATACCGGATGAACTGGCAGTGGAAGGAAACAGCAAACGGGACTGGAATCGCTTCCTGGACACGCATTACCGCCGCGGCTGGAATGTCAACATATCCAGGGTGATGGATAACGCCACACATGTGGCGGTGTACTTCGGCTCTTACCTGAAAAAGCCACCGGTGCCGATGAGTCGGCTGGAGCATTATGCCGGTCAGGATGAAATCGGTCTGCGTTACAACAGTCACCGTACAAAACGGGAAGAATACCTGTTGATGAGTGGAGATGAGTTCATGGAAAGGTTCTCCTGGCATGTAGCAGATAAGGGGTTCCGTATGGTGAGGTACTACGGTTTCCTGAGTCCGGTGAAGCGCCGCTTACTGGAAGAAGTTGTGTACGTCATAACGGAGACGGTGAGAAAAACGGCGATGCAAATCAGGTGGAGAGGGATGTATCAGAGGTTACTGAAGGTTGACCCGCTGAAGTGCATTCTGTGCGGAAGTCAGATGCGTTTTACGGGGCTGAAGCGGGGTTACCGACTGGCAGAGCTGGTCCTGATGCATGAGCGACTGGCACGACAGCAGGTGTGCGGCTGAGAGTCGCAGAGGGGAAGTTGCGTCCATTTTACCGGAAACGGAGCAAAAAACCGCCATTCATACCCTGTATCAATCAGTGTCATCCTGTTTAATAGTCGTTTCTGCTCATATGGTGCACAAGGGGTGTTGAAGAAATATCCGTTTTGTGGTGCTTTTTTAGTCTTTTGGGGATTTAAATTCCTATCGAT

>IS91-V41

CGAGTAGGCAGCCTGGCGGCTGCGGCTTGTCATGGTCTGGAATTACCGTTATAAAAAAAGATAATGTCATTGTCTTTCAGGTAGTTATATGGCCCGTTCAGCTAAACCCCGTAAACGCAAACCCGCACCACAAAGAAGCAAACTTCCCCGCTATGTTGTGAAACTTCATCCGGATGATTTTTTTGACGAAGAAGACGCTGAAGTTCTGCGCTTTGATAATTTTGACGATGCCGTTGAGTGCTGCGCTGACCTGGGTATTCCGTTCTTTCTGGATGCAGGAAACAAAAAGCTGGTCTTCTGGTTTGTTCGTGTCGATGACGAAGGGTATCCGGAAATAGCCCGCTGTACGGAGCGGGAGTTTGCAACCATTCTTGCCGGTATCAGTGCCGGTGGTATGTACTGCCCGGAATGCGGCACAGTTCACTGGCCGGATGGCGTTACCCCACCCGTCTGATGCTTCCCCGTTTTGCCGATATTTTTCAGCAGGGTAACCGCTGGCTTAACTGGCTGGAGAAACAGCCGGAAGGTTCAGTGCGTCCGGTGGTGACTGAGTCAGTGACAAAAATCATGGCATGCGGGACCACGCTGATGGGCTACACGCAATGGTGCTGTTCGTCACCGGACTGTTGCCACACCAAAAAGGTCTGCTTCCGGTGTAAAAGCCGCTCCTGTCCGCACTGCGGGGTGAAGGCTGGCGCACAGTGGATACAGTATCTGCTGAGCCTGGTCCCCGACTGCCCGTGGCAGCATATTGTGTTCACACTTCCCTGCCAGTACTGGTCCCTGGTGTTCCACAACCGGTGGTTACTGGCAGAGATGAGCCGCATTGCAGCGGATGTGATACTGGAAATCTGCCATCAGACAGATGTGGAGCCGGGGATATTCACGGTGATCCACACATGGGGGCGTGACCAGCAGTGGCATCCGCATATCCATTTATCGACAACTGCCGGTGGTGTGACGTCGGGCCACACCTGGAAAAATCTTCATTTTTACGCCCGTAAGGTGATGAGCATGTGGCGTTACCGGATAACGCGGCTACTGTCCCGGAAATACCCGGAGCTGGTAATACCGGATGAACTGGCAGTGGAAGGAAACAGCAAACGGGACTGGAATCGCTTCCTGGACACGCATTACCGCCGCGGCTGGAATGTCAACATATCCAGGGTGATGGATAACGCCACACATGTGGCGGTGTACTTCGGCTCTTACCTGAAAAAGCCACCGGTGCCGATGAGTCGGCTGGAGCATTATGCCGGTCAGGATGAAATCGGTCTGCGTTACAACAGTCACCGTACAAAACGGGAAGAATACCTGNTGATGAGTGGNGATGAGTTNATGGAAAGGTTCTCCTGGCATGTNGCNGATAAGGGGTTCCGTATGGTGAGGTACTACGGTTTCCTGAGTCCGGTNAAGCGCCGNTTACTGGAAGANGTTGTGTACGTCATAACGGAGACGGTGAGAAANACGGCGATGCAAATCAGGTGGAGAGGGATGTATCAGNGGTTACTGAAGGTTGACCCGCTNAAGTGCATTCTGTGCGGAAGTCAGATGCGTTTTACGGGGCTGAAGCGGGGTTACCGACTGGCAGAGCTGGTCCTGATGCATGAGCGACTGGCACGACAGCAGGTGTGCGGCTGAGAGCCGCAGAGGGGAAGTTGCGTCCATTTTACCGGAAACGGAGCAAAAAACCGCCATTCATACCCTGTATCAATCAGTGTCATCCTGTTTAATAGTCGTTTCCGCTCATATGGTGCACAAGGGGTGTTGAAGAAACATCCGTTTTGTGGTGCTTTTTTAGTCTTTTGGGGATTTAAATTCCTATCGAT

>IS91-V42

CGAGTAGGCAGCCTGGCGGCTGCGGCTTGTCATGGCCTGGAATTACCGTTATAAAAAATGATAATGTCATTGTCTTTCAGGTAGTTATATGGCCCGTTCAGCTAAACCCCGTAAACGCAAACCCGCACAACAAAGAAGCAAACTTCCCCGCTATGTTGTGAAACTTCATCCGGATGATTTTTTTGACGAAGAAGACGCTGAAGTTCTGCGCTTTGATAATTTTGACGATGCCGTTGAGTGCTGCGCTGACCTGGGTATTCCGTTCTTTCTGGATGCTGGAAACAAAAAGCTGGTCTTCTGGTTTGTTCGTGTCGATGACGAAGGGTATCCGGAAATAGCCCGCTGTACGGAGCGGGAGTTTGCAACCATTCTTGCCGGTATCAGTGCCGGTGGTATGTACTGCCCGGAATGCGGCACAGTTCACTGGCCGGATGACGTTACCCCACCCGTCTGATGCTTCCCCGTTTTGCCGATATTTTTCAGCAGGGTAACCGCTGGCTTAACTGGCTGGAGAAACAGCCGGAAGGTTCAGTGCGTCCGGTGGTGACTGAGTCAGTGACAAAAATCATGGCATGCGGGACCACGCTGATGGGCTACACACAATGGTGCTGTTCGTCACCGGACTGTTGCCACACCAAAAAGGTCTGCTTCCGGTGTAAAAGCCGCTCCTGTCCGCCCTGCGGGGTGAAGGCTGGCGCACAGTGGATACAGTATCTGCTGAGCCTGGTCCCCGACTGCCCGTGGCAGCATATTGTGTTCACACTTCCCTGCCAGTACTGGTCCCTGGTGTTCCACAACCGGTGGTTACTGGCAGAGATGAGCCGCATTGCAGCTGATGTGATACTGGAAATCTGCCATCAGGCAGATGTTGAGCCGGGGATATTCACGGTGATCCACACATGGGGGCGTGACCAGCAGTGGCATCCGCATATCCATTTATCGACAACTGCCGGTGGTGTGACGTCGGGCCACACCTGGAAAAATCTTCATTTTTACGCCCGTAAGGTGATGAGCATGTGGCGTTACCGGATAACGCGGCTACTGTCCCGGAAATACCCGGAGCTGGTAATACCGGATGAACTGGCAGTGGAAGGAAACAGCAAACGGGACTGGAATCGCTTCCTGGACACGCATTACCGCCGCGGCTGGAATGTCAACATATCCAGGGTGATGGATAACGCCACACATGTGGCGGTGTACTTCGGCTCTTACCTGAAAAAGCCACCGGTGCCGATGAGTCGGCTGGAGCATTATGCCGGTCAGGATGAAATCGGTCTGCGTTACAACAGTCACCGTACAAAACGGGAAGAATACCTGTTGATGAGTGGAGATGAGTTCATGGAAAGGTTCTCCTGGCATGTAGCAGATAAGGGGTTCCGTATGGTGAGGTACTACGGTTTCCTGAGTCCGGTGAAGCGCCGCTTACTGGAAGAAGTTGTGTACGTCATAACGGAGACGGTGAGAAAGATGGCGATGCAAATCAGGTGGAGAGGGATGTATCAGAGGTTACTGAAGGTTGACCCGCTGAAGTGCATTCTGTGCGGAAGTCAGATGCGTTTTACGGGGCTGAAGCGGGGTTACCGACTGGCAGAGCTGGTCCTGATGCATGAGCGACTGGCACGACAGCAGGTGTGCGGCTGAGAGCCGCAGAGGGGAAGTTGCGTCCATTTTACCGGAAACGGAGCAAAAAACCGGCATTCATACCCTGTATCAATCAGTGTCATCCTGTTTAATAGTCGTTTCCGCTCATATGGTGCACAAGGGGTGTTGAAGAAACATCCGTTTTGTGGTGCTTTTTTAGTCTTTTGGGGATTTAAATTCCTATCGAT

>IS91-V43

CGAGTAGGCAGCCTGGCGGCTGCGGCTTGTCATGGTCTGAGATTACCGTTATAAAAACAGGTAATATCATTGTCTTTCAGGTGGTTATATGGCCCGTTCAGCTAAACCCCGTAAACGAAAACCCGCACCACAAAGAAGCAAACTTCCCCGCTATGTTGTGAAACTTCATCCGGATGATTTTTTTGACGAAGAAGACGCTGAAGTTCTGCGCTTTGATAATTTTGACGATGCCGTTGAGTGCTGCGCTGACCTGGGTATTCCGTTCTTTCTGGATGCAGGAAACAAAAAGCTGGTCTTCTGGTTTGTTCGTGTCGATGACGAAGGGTATCCGGAAATAGCCCGCTGTACGGAGCGGGAGTTTGCAACCATTCTTGCCGGTATCAGTGCCGGTGGTATGTACTGCCCGGAATGCGGCACAGTTCACTGGCCGGATGGCGTTACCCCACCCGTCTGATGCTTCCCCGTTTTGCCGATATTTTTCAGCAGGGTAACCGCTGGCTTAACTGGCTGGAGAAACAGCCGGAAGGTTCAGTGCGTCCGGTGGTGATTGAGTCAGTGACAAAAATCATGGCATGCGGGACCACGCTGATGGGCTACACGCAATGGTGCTGTTCGTCACCGGACTGTTGCCACACCAAAAAGGTCTGCTTCCGGTGTAAAAGCCGCTCCTGTCCGCACTGCGGGGTGAAGGCTGGCGCACAGTGGATACAGTATCTGCTGAGCCTGGTCCCCGACTGCCCGTGGCAGCATATTGTGTTCACACTTCCCTGCCAGTACTGGTCCCTGGTGTTCCACAACCGGTGGTTACTGGCAGAGATGAGCCGCATTGCAGCGGATGTGATACTGGAAATCTGCCATCAGACAGATGTGGAGCCGGGGATATTCACGGTGATCCACACATGGGGGCGTGACCAGCAGTGGCATCCGCATATCCATTTATCGACAACTGCCGGTGGTGTGACGTCGGGCCACACCTGGAAAAATCTTCATTTTTACGCCCGTAAGGTGATGAGCATGTGGCGTTACCGGATAACGCGGCTACTGTCCCGGAAATACCCGGAGCTGGTAATACCGGATGAACTGGCAGTGGAAGGAAACAGCAAACGGGACTGGAATCGCTTCCTGGACACGCATTACCGCCGCGGCTGGAATGTCAACATATCCAGGGTGATGGATAACGCCACACATGTGGCGGTGTACTTCGGCTCTTACCTGAAAAAGCCACCGGTGCCGGTGAGTCGGCTGGAGCATTATGCCGGTCAGGATGAAATCGGTCTGCGTTACAACAGTCACCGTACAAAACGGGAAGAATATCTGTTGATGAGTGGAGATGAGTTCATGGAAAGGTTCTCCTGGCATGTAGCAGATAAGGGGTTCCGTATGGTGAGGTACTACGGTTTCCTGAGTCCGGTGAAGCGCCGCTTACTGGAAGAAGTGGTGTACGTCATAACGGAGACGGTGAGAAAAACGGCGATGCAAATCAGGTGGAGAGGGATGTATCAGAGGTTACTGAAGGTTGACCCGCTGAAGTGCATTCTGTGCGGAAGTCAGATGCGTTTTACGGGGCTGAAGCGGGGTTACCGACTGGCAGAGCTGGTCCTGATGCATGAGCGACTGGCACGACAGCAGGTGTGCGGCTGAGAGCCGCAGAGGGGAAGTTGCGTCCATTTTACCGGAAACGGAGCAAAAAACCGGCATTCATACCCTGTATCAATCAGTGTCATCCTGTTTAATAGTCGTTTCCGCTCATATGGTGCACAAGGGGTGTTGAAGAAACATCCGTTTTGTGGTGCTTTTTTAGTCTTTTGGGGATTTAAATTCCTATCGAT

>IS91-V44

CGAGTAGGCAGCCTGGCGGCTGCGGCTTGTCATGGTCTGAGATTACCGTTATAAAAACAGGTAATATCATTGTCTTTCAGGTGGTTATATGGCCCGTTCAGCTAAACCCCGTAAACGAAAACCCGCACCACAAAGAAGCAAACTTCCCCGCTATGTTGTGAAACTTCATCCGGATGATTTTTTTGACGAAGAAGACGCTGAAGTTCTGCGCTTTGATAATTTTGACGATGCCGTTGAGTGCTGCGCTGACCTGGGGATTCCGTTCTTTCTGGATGCAGGAAACAAAAAGCTGGTCTTCTGGTTTGTACGTGTCGATGACGAAGGGTATCCGGAAATAGCCCGCTGTACGGAGCGGGAGTTTGCAACCATTCTTGCCGGTATCAGTGCCGGTGGTATGTACTGCCCGGAATGCGGCACAGTTCACTGGCCGGATGGCGTTACCCCACCCGTCTGATGCTTCCCCGTTTTGCCGATATTTTTCAGCAGGGTAACCGCTGGCTTAACTGGCTGGAGAAACAGCCGGAAGGTTCAGTGCGTCCGGTGGTGACTGAGTCAGTGACAAAAATCATGGCATGCGGGACCACGCTGATGGGCTACACGCAATGGTGCTGTTCGTCACCGGACTGTTGCCACACCAAAAAGGTCTGCTTCCGGTGTAAAAGCCGCTCCTGTCCGCACTGCGGGGTGAAGGCTGGCGCACAGTGGATACAGTATCTGCTGAGCCTGGTCCCCGACTGCCCGTGGCAGCATATTGTGTTCACACTTCCCTGCCAGTACTGGTCCCTGGTGTTCCACAACCGGTGGTTACTGGCAGAGATGAGCCGCATTGCAGCGGATGTGATACTGGAAATCTGCCATCAGACAGATGTGGAGCCGGGGATATTCACGGTGATCCACACATGGGGGCGTGACCAGCAGTGGCATCCGCATATCCATTTATCGACAACTGCCGGTGGTGTGACGTCGGGCCACACCTGGAAAAATCTTCATTTTTACGCCCGTAAGGTGATGAGCATGTGGCGTTACCGGATAACGCGGCTACTGTCCCGGAAATACCCGGAGCTGGTAATACCGGATGAACTGGCAGTGGAAGGAAACAGCAAACGGGACTGGAATCGCTTCCTGGACACGCATTACCGCCGCGGCTGGAATGTCAACATATCCAGGGTGATGGATAACGCCACACATGTGGCGGTGTACTTCGGCTCTTACCTGAAAAAGCCACCGGTGCCGGTGAGTCGGCTGGAGCATTATGCCGGTCAGGATGAAATCGGTCTGCGTTACAACAGTCACCGTACAAAACGGGAAGAATATCTGTTGATGAGTGGCGATGAGTTCATGGAAAGGTTCTCGTGGCATGTAGCAGATAAGGGGTTCCGTATGGTGAGGTACTACGGTTTCCTGAGTCCGGTGAAGCGCCGCTTACTGGAAGAAGTGGTGTACGTCATAACGGAGACGGTGAGAAAAACGGCGATGCAAATCAGGTGGAGAGGGATGTATCAGAGGTTACTGAAGGTTGACCCGCTGAAGTGCATTCTGTGCGGAAGTCAGATGCGTTTTACGGGGCTGAAGCGGGGTTACCGACTGGCAGAGCTGGTCCTGATGCATGAGCGACTGGCACGACAGCAGGTGTGCGGCTGAGAGCCGCAGAGGGGAAGTTGCGTCCATTTTACCGGAAACGGAGCAAAAAACCGCCATTCATACCCTGTATCAATCAGTGTCATCCTGTTTAATAGTCGTTTCCGCTCATATGGTGCACAAGGGGTGTTGAAGAAACATCCGTTTTGTGGTGCTTTTTTAGTCTTTTGGGGATTTAAATTCCTATCGAT

>IS91-V45

CGAGTAGGCAGCCTGGCGGCTGCGGCTTGTCATGGCCTGGAATTACCGTTATAAAAAAGATAATGTCATTGTCTTTCAGGTAGTTATATGGCCCGTTCAGCTAAACCCCGTAAACGCAAACCCGCACAACAAAGAAGCAAACTTCCCCGCTATGTTGTGAAACTTCATCCGGATGATTTTTTTGACGAAGAAGACGCTGAAGTTCTGCGCTTTGATAATTTTGACGATGCCGTTGAGTGCTGCGCTGACCTGGGTATTCCGTTCTTTCTGGATGCAGGAAACAAAAAGCTGGTCTTCTGGTTTGTTCGTGTCGATGACGAAGGGTATCCGGAAATAGCCCGCTGTACGGAGCGGGAGTTTGCAACCATTCTTGCCGGTATCAGTGTCGGTGGTATGTACTGCCCGGAATGCGGCACAGTTCACTGGCCGGATGGCGTTACCCCACCCGTCTGATGCTTCCCCGTTTTGCCGATATTTTTCAGCAGGGTAACCGCTGGCTTAACTGGCTGGATAAACAGCCGGAAGGTTCAGTGCGTCCGGTGGTGACTGAGTCAGTGACAAAAATCATGGCATGCGGGACCACGCTGATGGGCTACACACAATGGTGCTGTTCGTCACCGGACTGTTGCCACACCAAAAAGGTCTGCTTCCGGTGTAAAAGCCGCTCCTGTCCGCACTGCGGGGTGAAGGCTGGCGCACAGTGGATACAGTATCTGCTGAGCCTGGTCCCCGACTGCCCGTGGCAGCATATTGTGTTCACACTTCCCTGCCAGTACTGGTCCCTGGTGTTCCACAACCGGTGGTTACTGGCAGAGATGAGCCGCATTGCAGCGGATGTGATACTGGAAATCTGCCGTCAGGCAGATGTGGAGCCGGGGATATTCACGGTGATCCACACATGGGGGCGTGACCAGCAGTGGCATCCGCATATCCATTTATCGACAACTGCCGGTGGTGTGACGTCGGGCCACACCTGGAAAAATCTTCATTTTTACGCCCGTAAGGTGATGAGCATGTGGCGTTACCGGATAACGCGGCTACTGTCCCGGAAATACCCGGAGCTGGTAATACCGGATGAACTGGCAGTGGAAGGAAACAGCAAACGGGACTGGAATCGCTTCCTGGACACGCATTACCGCCGCGGCTGGAATGTCAACATATCCAGGGTGATGGATAACGCCACACATGTGGCGGTGTACTTCGGCTCTTACCTGAAAAAGCCACCGGTGCCGATGAGTCGGCTGGAGCATTATGCCGGTCAGGATGAAATCGGTCTGCGTTACAACAGTCACCGTACAAAACGGGAAGAATACCTGTTGATGAGTGGAGATGAGTTCATGGAAAGGTTCTCCTGGCATGTAGCAGATAAGGGGTTCCGTATGGTGAGGTACTACGGTTTCCTGAGTCCGGTGAAGCGCCGCTTACTGGAAGAAGTTGTGTACGTCATAACGGAGACGGTGAGAAAAACGGCGATGCAAATCAGGTGGAGAGGGATGTATCAGAGGTTACTGAAGGTTGACCCGCTGAAGTGCATTCTGTGCGGAAGTCAGATGCGCTTTACGGGGCTGAAACGGGGTTACCGACTGGCAGAGCTGGTCCTGATGCATGAGCGACTGGCACGACAGCAGGTGTGCGGCTGAGAGCCGCAGAGGGGAAGTTGCGTCCATTTTACCGGAAACGGAGCAAAAAACCGCCATTCATACCCTGTATCAATCAGTGTCATCCTGTTTAATAGTCGTTTTCGCTCATATGGTGCACAAGGGGTGTTGAAGAAACATCCGTTTTGTGGTGCTTTTTTAGTCTTATGGGGATTTAAATTCCTATCGAT

>IS91-V46

CGAGTAGGCAGCCTGGCGGCTGCGGCTTGTCATGGTCTGAGATTACCGTTATAAAAACAGGTAATATCATTGTCTTTCAGGTGGTTATATGGCCCGTTCAGCTAAACCCCGTAAACGAAAACCCGCACAACAAAGAAGCAAACTTCCCCGCTATGTTGTGAAACTTCATCCGGATGATTTTTTTGACGAAGAAGACGCTGAAGTTCTGCGCTTTGATAATTTTGACGATGCCGTTGAGTGCTGCGCTGACCTGGGGATTCCGTTCTTTCTGGATGCAGGAAACAAAAAGCTGGTCTTCTGGTTTGTACGTGTCGATGACGAAGGGTATCCGGAAATAGCCCGCTGTACGGAGCGGGAGTTTGCGAACATTCTTGCCGGTATCAGCGCCGGCGGTATGTACTGTCCGGAATGCGGCATGGTTCACTGGCCGGATGGCGTTACCCCACCCGTCTGATGCTTCCCCGTTTTGCCGATATTTTTCAGCAGGGTAACCGCTGGCTTAACTGGCTGGAGAAACAGCCGGAAGGTTCAGTGCGTCCGGTGGTGACTGAGTCAGTGACAAAAATCATGGCATGCGGGACCACGCTGATGGGCTACACGCAATGGTGCTGTTCGTCACCGGACTGTTGCCACACCAAAAAGGTCTGCTTCCGGTGTAAAAGCCGCTCCTGTCCGCACTGCGGGGTGAAGGCTGGCGCACAGTGGATACAGTATCTGCTGAGCCTGGTCCCCGACTGCCCGTGGCAGCATATTGTGTTCACACTTCCCTGCCAGTACTGGTCCCTGGTGTTCCACAACCGGTGGCTACTGGCAGAGATGAGCCGCATTGCAGCGGATGTGATACTGGAAATCTGCCATCAGACAGATGTGGAGCCGGGGATATTCACGGTGATCCACACATGGGGGCGTGACCAGCAGTGGCATCCGCATATCCATTTATCGACAACTGCCGGTGGTGTGACGTCGGGCCACACCTGGAAAAATCTTCATTTTTACGCCCGTAAGGTGATGAGCATGTGGCGTTACCGGATAACGCGGCTACTGTCCCGGAAATACCCGGAGCTGGTAATACCGGATGAACTGGCAGTGGAAGGAAACAGCAAACGGGACTGGAATCGCTTCCTGGACACGCATTACCGCCGCGGCTGGAATGTCAACATATCCAGGGTGATGGATAACGCCACACATGTGGCGGTGTACTTCGGCTCTTACCTGAAAAAGCCACCGGTGCCGGTGAGTCGGCTGGAGCATTATGCCGGTCAGGATGAAATCGGTCTGCGTTACAACAGTCACCGTACAAAACGGGAAGAATATCTGTTGATGAGTGGAGATGAGTTCATGGAAAGGTTCTCCTGGCATGTAGCAGATAAGGGGTTCCGTATGGTGAGGTACTACGGTTTCCTGAGTCCGGTGAAGCGCCGCTTACTGGAAGAAGTGGTGTACGTCATAACGGAGACGGTGAGAAAAACGGCGATGCAAATCAGGTGGAGAGGGATGTATCAGAGGTTACTGAAGGTTGACCCGCTGAAGTGCATTCTGTGCGGAAGTCAGATGCGTTTTACGGGGCTGAAGCGGGGTTACCGACTGGCAGAGCTGATCCTGATGCATGAGCGACTGGCACGACAGCAGGTGTGCGGCTGAGAGCCGCAGAGGGGAAGTTGCGTCCATTTTACCGGAAACGGAGCAAAAAACCGCCATTCATACCCTGTATCAATCAGTGTCATCCTGTTTAATAGTCGTTTCCGCTCATATGGTGCACAAGGGGTGTTGAAGAAACATCCGTTTTGTGGTGCTTTTTTAGTCTTTTGGGGATTTAAATTCCTATCGAT

>IS91-V47

CGAGTAGGCAGCCTGGCGGCTGCGGCTTGTCATGGTCTGAGATTACCGTTATAAAAACAGGTAATATCATTGTCTTTCAGGTGGTTATATGGCCCGTTCAGCTAAACCCCGTAAACGAAAACCCGCACAACAAAGAAGCAAACTTCCCCGCTATGTTGTGAAACTTCATCCGGATGATTTTTTTGACGAAGAAGACGCTGAAGTTCTGCGCTTTGATAATTTTGACGATGCCGTTGAGTGCTGCGCTGACCTGGGGATTCCGTTCTTTCTGGATGCAGGAAACAAAAAGCTGGTCTTCTGGTTTGTACGTGTCGATGACGAAGGGTATCCGGAAATAGCCCGCTGTACGGAGCGGGAGTTTGCGAACATTCTTGCCGGTATCAGCGCCGGCGGTATGTACTGTCCGGAATGCGGCATGGTTCACTGGCCGGATGGCGTTACCCCACCCGTCTGATGCTTCCCCGTTTTGCCGATATTTTTCAGCAGGGTAACCGCTGGCTTAACTGGCTGGAGAAACAGCCGGAAGGTTCAGTGCGTCCGGTGGTGACTGAGTCAGTGACAAAAATCATGGCATGCGGGACCACGCTGATGGGCTACACGCAATGGTGCTGTTCGTCACCGGACTGTTGCCACACCAAAAAGGTCTGCTTCCGGTGTAAAAGCCGCTCCTGTCCGCACTGCGGGGTGAAGGCTGGCGCACAGTGGATACAGTATCTGCTGAGCCTGGTCCCCGACTGCCCGTGGCAGCATATTGTGTTCACACTTCCCTGCCAGTACTGGTCCCTGGTGTTCCACAACCGGTGGTTACTGGCAGAGATGAGCCGCATTGCAGCGGATGTGATACTGGAAATCTGCCATCAGACAGATGTGGAGCCGGGGATATTCACGGTGATCCACACATGGGGGCGTGACCAGCAGTGGCATCCGCATATCCATTTATCGACAACTGCCGGTGGTGTGACGTCGGGCCACACCTGGAAAAATCTTCATTTTTACGCCCGTAAGGTGATGAGCATGTGGCGTTACCGGATAACGCGGCTACTGTCCCGGAAATACCCGGAGCTGGTAATACCGGATGAACTGGCAGTGGAAGGAAACAGCAAACGGGACTGGAATCGCTTCCTGGACACGCATTACCGCCGCGGCTGGAATGTCAACATATCCAGGGTGATGGATAACGCCACACATGTGGCGGTGTACTTCGGCTCTTACCTGAAAAAGCCACCGGTGCCGGTGAGTCGGCTGGAGCATTATGCCGGTCAGGATGAAATCGGTCTGCGTTACAACAGTCACCGTACAAAACGGGAAGAATATCTGTTGATGAGTGGAGATGAGTTCATGGAAAGGTTCTCCTGGCATGTAGCAGATAAGGGGTTCCGTATGGTGAGGTACTACGGTTTCCTGAGTCCGGTGAAGCGCCGCTTACTGGAAGAAGTGGTGTACGTCATAACGGAGACGGTGAGAAAAACGGCGATGCAAATCAGGTGGAGAGGGATGTATCAGAGGTTACTGAAGGTTGACCCGCTGAAGTGCATTCTGTGCGGAAGTCAGATGCGTTTTACGGGGCTGAAGCGGGGTTACCGACTGGCAGAGCTGATCCTGATGCATGAGCGACTGGCACGACAGCAGGTGTGCGGCTGAGAGCCGCAGAGGGGAAGTTGCGTCCATTTTACCGGAAACGGAGCAAAAAACCGCATTCATACCCTGTATCAATCAGTGTCATCCTGTTTAATAGTCGTTTCCGCTCATATGGTGCACAAGGGGTGTTGAAGAAACATCCGTTTTGTGGTGCTTTTTTAGTCTTTTGGGGATTTAAATTCCTATCGAT

>IS91-V48

CGAGTAGGCAGCCTGGCGGCTGCGGCTTGTCATGGCCTGGAATTACCGTTATAAAAAAAGATAATGTCATTGTCTTTCAGGTAGTTATATGGCCCGTTCAGCTAAACCCCGTAAACGCAAACCCGCACAACAAAGAAGCAAACTTCCCCGCTATGTTGTGAAACTTCATCCGGATGATTTTTTTGACGAAGAAGACGCTGAAGTTCTGCGCTTTGATAATTTTGACGATGCCGTTGAGTGCTGCGCTGACCTGGGTATTCCGTTCTTTCTGGATGCTGGAAACAAAAAGCTGGTCTTCTGGTTTGTTCGTGTCGATGACGAAGGGTATCCGGAAATAGCCCGCTGTACGGAGCGGGAGTTTGCAACCATTCTTGCCGGTATCAGTGCCGGTGGTATGTACTGCCCGGAATGCGGCACGGTTCACTGGCCGGATGGCGTTACCCCACCCGTCTGATGCTTCCCCGTTTTGCCGATATTTTTCAGCAGGGTAACCGCTGGCTTAACTGGCTGGAGAAACAGCCGGAAGGTTCAGTGCGTCCGGTGGTGACTGAGTCAGTGACAAAAATCATGGCATGCGGGACCACGCTGATGGGCTACACACAATGGTGCTGTTCGTCACCGGACTGTTGCCACACCAAAAAGGTCTGCTTCCGGTGTAAAAGCCGCTCCTGTCCGCACTGCGGGGTGAAGGCTGGCGCACAGTGGATACAGTATCTGCTGAGCCTGGTCCCCGACAGCCCGTGGCAGCATATTGTGTTCACACTTCCCTGCCAGTACTGGTCCCTGGTGTTCCACAACCGGTGGTTACTGGCAGAGATGAGCCGCATTGCAGCTGATGTGATACTGGAAATCTGCCATCAGGCAGATGTTGAGCCGGGGATATTCACGGTGATCCACACATGGGGGCGTGACCAGCAGTGGCATCCGCATATCCATTTATCGACAACTGCCGGTGGTGTGACGTCGGGCCACACCTGGAAAAATCTTCATTTTTACGCCCGTAAGGTGATGAGCATGTGGCGTTACCGGATAACACGGCTACTGTCCCGGAAATACCCGGAGCTGGTAATACAGGATGAACTGGCAGTGGAAGGAAACAGCAAACGGGACTGGAATCGCTTCCCGGACACGCATTACCGCCGCGGCTGGAATGTCAACATATCCAGGGTGATGGATAATGCCACACATGTGGCGGTGTACTTCGGCTCTTACCTGAAAAAGCCACCGGTGCCGATGAGTCGGCTGGAGCATTATGCCGGTCAGGATGAAATCGGTCTGCGTTACAACAGTCACCGTACAAAACGGGAAGAATACCTGTTGATGAGTGGAGATGAGTTCATGGAAAGGTTCTCCTGGCATGTAGCAGATAAGGGGTTCCGTATGGTGAGGTACTACGGTTTCCTGAGTCCGGTGAAGCGCCGCTTACTGGAAGAAGTTGTGTACGTCATAACGGAGACGGTGAGAAAAACGGCGATGCAAATCAGGTGGAGAGGGATGTATCAGAGGTTACTGAAGGTTGACCCGCTGAAGTGCATTCTGTGCGGAAGTCAGATGCGTTTTACGGGGCTGAAACGGGGTTACCGACTGGCAGAGCTGGTCCTGATGCATGAGCGACTGGCACGACAGCAGGTGTGCGGCTGAGAGCCGCAGAGGGGAAGTTGCGTCCATTTTACCGGAAACGGAGCAAAAAACCGCCATTCATACCCTGTATCAATCAGTGTCATCCTGTTTAATAGTCGTTTCCGCTCATATGGTGCACAAGGGGTGTTGAAGAAACATCCGTTTCGTGGTGCTTTTTTAGTCTTTTGGGGATTTTAATTCCTATCGAT

>IS91-V49

CGAGTAGGCAGCCTGGCGGCTGCGGCTTGTCATGGTCTGAGATTACCGTTATAAAAACAGGTAATATCATTGTCTTTCAGGTGGTTATATGGCCCGTTCAGCTAAACCCCGTAAACGAAAACCCGCACCACAAAGAAGCAAACTTCCCCGCTATGTTGTGAAACTTCATCCGGATGATTTTTTTGACGAAGAAGACGCTGAAGTTCTGCGCTTTGATAATTTTGACGATGCCGTTGAGTGCTGCGCTGACCTGGGGATTCCGTTCTTTCTGGATGCAGGAAACAAAAAGCTGGTCTTCTGGTTTGTACGTGTCGATGACGAAGGGTATCCGGAAATAGCCCGCTGTACGGAGCGGGAGTTTGCAACCATTCTTGCCGGTATCAGTGCCGGTGGTATGTACTGCCCGGAATGCGGCACAGTTCACTGGCCGGATGGCGTTACCCCACCCGTCTGATGCTTCCCCGTTTTGCCGATATTTTTCAGCAGGGTAACCGCTGGCTTAACTGGCTGGAGAAACAGCCGGAAGGTTCAGTGCGTCCGGTGGTGACTGAGTCAGTGACAAAAATCATGGCATGCGGGACCACGCTGATGGGCTACACGCAATGGTGCTGTTCGTCACCGGACTGTTGCCACACCAAAAAGGTCTGCTTCCGGTGTAAAAGCCGCTCCTGTCCGCACTGCGGGGTGAAGGCTGGCGCACAGTGGATACAGTATCTGCTGAGCCTGGTCCCCGACTGCCCGTGGCAGCATATTGTGTTCACACTTCCCTGCCAGTACTGGTCCCTGGTGTTCCACAACCGGTGGTTACTGGCAGAGATGAGCCGCATTGCAGCGGATGTGATACTGGAAATCTGCCATCAGACAGATGTGGAGCCGGGGATATTCACGGTGATCCACACATGGGGGCGTGACCAGCAGTGGCATCCGCATATCCATTTATCGACAACTGCCGGTGGTGTGACGTCGGGCCACACCTGGAAAAATCTTCATTTTTACGCCCGTAAGGTGATGAGCATGTGGCGTTACCGGATAACGCGGCTACTGTCCCGGAAATACCCGGAGCTGGTAATACTGGATGAACTGGCAGTGGAAGGAAACAGCAAACGGGACTGGAATCGCTTCCTGGACACGCATTACCGCCGCGGCTGGAATGTCAACATATCCAGGGTGATGGATAACGCCACACATGTGGCGGTGTACTTCGGCTCTTACCTGAAAAAGCCACCGGTGCCGGTGAGTCGGCTGGAGCATTATGCCGGTCAGGATGAAATCGGTCTGCGTTACAACAGTCACCGTACAAAACGGGAAGAATATCTGTTGATGAGTGGAGATGAGTTCATGGAAAGGTTCTCCTGGCATGTAGCAGATAAGGGGTTCCGTATGGTGAGGTACTACGGTTTCCTGAGTCCGGTGAAGCGCCGCTTACTGGAAGAAGTGGTGTACGTCATAACGGAGACGGTGAGAAAAACGGCGATGCAAATCAGGTGGAGAGGGATGTATCAGAGGTTACTGAAGGTTGACCCGCTGAAGTGCATTCTGTGCGGAAGTCAGATGCGTTTTACGGGACTGAAGCGGGGTTACCGACTGGCAGAGCTGGTCCTGATGCATGAGCGACTGGCACGACAGCAGGTGTGCGGCTGAGAGCCGCAGAGGGGAAGTTGCGTCCATTTTACCGGAAACGGAGCAAAAAACCGGCATTCATACCCTGTATCAATCAGTGTCATCCTGTTTAATAGTCGTTTCCGCTCATATGGTGCACAAGGGGTGTTGAAGAAACATCCGTTTTGTGGTGCTTTTTTAGTCTTTTGGGGATTTAAATTCCTATCGAT

>IS91-V50

CGAGTAGGCAGCCTGGCGGCTGCGGCTTGTCATGGTCTGAGATTACCGTTATAAAAACAGGTAATATCATTGTCTTTCAGGTGGTTATATGGCCCGTTCAGCTAAACCCCGTAAACGAAAACCCGCACCACAAAGAAGCAAACTTCCCCGCTATGTTGTGAAACTTCATCCGGATGATTTTTTTGACGAAGAAGACGCTGAAGTTCTGCGCTTTGATAATTTTGACGATGCCGTTGAGTGCTGCGCTGACCTGGGTATTCCGTTCTTTCTGGATGCAGGAAACAAAAAGCTGGTCTTCTGGTTTGTTCGTGTCGATGACGAAGGGTATCCGGAAATAGCCCGCTGTACGGAGCGGGAGTTTGCAACCATTCTTGCCGGTATCAGTGCCGGTGGTATGTACTGCCCGGAATGCGGCACAGTTCACTGGCCGGATGGCGTTACCCCACCCGTCTGATGCTTCCCCGTTTTGCCGATATTTTTCAGCAGGGTAACCGCTGGCTTAACTGGCTGGAGAAACAGCCGGAAGGTTCAGTGCGTCCGGTGGTGATTGAGTCAGTGACAAAAATCATGGCATGCGGGACCACGCTGATGGGCTACACGCAATGGTGCTGTTCGTCACCGGACTGTTGCCACACCAAAAAGGTCTGCTTCCGGTGTAAAAGCCGCTCCTGTCCGCACTGCGGGGTGAAGGCTGGCGCACAGTGGATACAGTATCTGCTGAGCCTGGTCCCCGACTGCCCGTGGCAGCATATTGTGTTCACACTTCCCTGCCAGTACTGGTCCCTGGTGTTCCACAACCGGTGGTTACTGGCAGAGATGAGCCGCATTGCAGCGGATGTGATACTGGAAATCTGCCATCAGACAGATGTGGAGCCGGGGATATTCACGGTGATCCACACATGGGGGCGTGACCAGCAGTGGCATCCGCATATCCATTTATCGACAACTGCCGGTGGTGTGACGTCGGGCCACACCTGGAAAAATCTTCATTTTTACGCCCGTAAGGTGATGAGCATGTGGCGTTACCGGATAACGCGGCTACTGTCCCAGAAATACCCGGAGCTGGTAATACCGGATGAACTGGCAGTGGAAGGAAACAGCAAACGGGACTGGAATCGCTTCCTGGACACGCATTACCGCCGCGGCTGGAATGTCAACATATCCAGGGTGATGGATAACGCCACACATGTGGCGGTGTACTTCGGCTCTTACCTGAAAAAGCCACCGGTGCCGGTGAGTCGGCTGGAGCATTATGCCGGTCAGGATGAAATCGGTCTGCGTTACAACAGTCACCGTACAAAACGGGAAGAATATCTGTTGATGAGTGGAGATGAGTTCATGGAAAGGTTCTCCTGGCATGTAGCAGATAAGGGGTTCCGTATGGTGAGGTACTACGGTTTCCTGAGTCCGGTGAAGCGCCGCTTACTGGAAGAAGTGGTGTACGTCATAACGGAGACGGTGAGAAAAACGGCGATGCAAATCAGGTGGAGAGGGATGTATCAGAGGTTACTGAAGGTTGACCCGCTGAAGTGCATTCTGTGCGGAAGTCAGATGCGTTTTACGGGGCTGAAGCGGGGTTACCGACTGGCAGAGCTGGTCCTGATGCATGAGCGACTGGCACGACAGCAGGTGTGCGGCTGAGAGCCGCAGAGGGGAAGTTGCGTCCATTTTACCGGAAACGGAGCAAAAAACCGGCATTCATACCCTGTATCAATCAGTGTCATCCTGTTTAATAGTCGTTTCCGCTCATATGGTGCACAAGGGGTGTTGAAGAAACATCCGTTTTGTGGTGCTTTTTTAGTCTTTTGGGGATTTAAATTCCTATCGAT

>IS91-V51

CGAGTAGGCAGCCTGGCGGCTGCGGCTTGTCATGGCCTGGAATTACCGTTATAAAAAATGATAATGTCATTGTCTTTCAGGTAGTTATATGGCCCGTTCAGCTAAACCCCGTAAACGCAAACCCGCACAACAAAGAAGCAAACTTCCCCGCTATGTTGTGAAACTTCATCCGGATGATTTTTTTGACGAAGAAGACGCTGAAGTTCTGCGCTTTGATAATTTTGACGATGCCGTTGAGTGCTGCGCTGACCTGGGTATTCCGTTCTTTCTGGATGCTGGAAACAAAAAGCTGGTCTTCTGGTTTGTTCGTGTCGATGACGAAGGGTATCCGGAAATAGCCCGCTGTACGGAGCGGGAGTTTGCAACCATTCTTGCCGGTATCAGTGCCGGTGGTATGTACTGCCCGGAATGCAGCACAGTTCACTGGCCGGATGGCGTTACCCCACCCGTCTGATGCTTCCCCGTTTTGCCGATATTTTTCAGCAGGGTAACCGCTGGCTTAACTGGCTGGAGAAACAGCCGGAAGGTTCAGTGCGTCCGGTGGTGACTGAGTCAGTGACAAAAATCATGGCATGCGGGACCACGCTGATGGGCTACACACAATGGTGCTGTTCGTCACCGGACTGTTGCCACACCAAAAAGGTCTGCTTCCGGTGTAAAAGCCGCTCCTGTCCGCACTGCGGGGTGAAGGCTGGCGCACAGTGGATACAGTATCTGCTGAGCCTGGTCCCCGACTGCCCGTGGCAGCATATTGTGTTCACACTTCCCTGCCAGTACTGGTCCCTGGTGTTCCACAACCGGTGGTTACTGGCAGAGATGAGCCGCATTGCAGCTGATGTGATACTGGAAATCTGCCATCAGGCAGATGTTGAGCCGGGGATATTCACGGTGATCCACACATGGGGGCGTGACCAGCAGTGGCATCCGCATATCCATTTATCGACAACTGCCGGTGGTGTGACGTCGGGCCACACCTGGAAAAATCTTCATTTTTACGCCCGTAAGGTGATGAGCATGTGGCGTTACCGGATAACGCGGCTACTGTCCCGGAAATACCCGGAGCTGGTAATACCGGATGAACTGGCAGTGGAAGGAAACAGCAAACGGGACTGGAATCGCTTCCCGGACACGCATTACCGCCGCGGCTGGAATGTCAACATATCCAGGGTGATGGATAACGCCACACATGTGGCGGTGTACTTCGGCTCTTACCTGAAAAAGCCACCGGTGCCGATGAGTCGGCTGGAGCATTATGCCGGTCAGGATGAAATCGGTCTGCGTTACAACAGTCACCGTACAAAACGGGAAGAATACCTGTTGATGAGTGGAGATGAGTTCATGGAAAGGTTCTCCTGGCATGTAGCAGATAAGGGGTTCCGTATGGTGAGGTACTACGGTTTCCTGAGTCCGGTGAAGCGCCGCTTACTGGAAGAAGTTGTGTACGTCATAACGGAGACGGTGAGAAAAACGGCGATGCAAATCAGGTGGAGAGGGATGTATCAGAGGTTACTGAAGGTTGACCCGCTGAAGTGCATTCTGTACGGAAGTCAGATGCGTTTTACGGGGCTGAAGCGGGGTTACCGACTGGCAGAGCTGGCCCTGATGCATGAGCGACTGGCACGACAGCAGGTGTGCGGCTGAGAGCCGCAGAGGGGAAGTTGCGTCCATTTTACCGGAAACGGAGCAAAAAACCGCCATTCATACCCTGTATCAATCAGTGTCATCCTGTTTAATAGTCGTTTCCGCTCATATGGTGCACAAGGGGTGTTGAAGAAACATCCGTTTTGTGGTGCTTTTTTAGCCTCATGGGGATTTAAATTCCTATCGAT

>IS91-V52

CGAGTAGGCAGCCTGGCGGCTGCGGCTTGTCATGGTCTGAGATTACCGTTATAAAAACAGGTAATATCATTGTCTTTCAGGTGGTTATATGGCCCGTTCAGCTAAACCCCGTAAACGAAAACCCGCACCACAAAGAAGCAAACTTCCCCGCTATGTTGTGAAACTTCATCCGGATGATTTTTTTGACGAAGAAGACGCAGAAGTTCTGCGCTTTGATAGTTTTGACGATGCCGTTGAATGCTGCGCAGACCTGAATATTCCGTTCTTTCTGGATGCAGGAAACAAAAAGCTGGTCTTCTGGTTTGTACGTGTCGATGACGAAGGGTATCCGGAAATAGCCCGCTGTACGGAGCGGGAGTTTGCAACCATTCTTGCCGGTATCAGTGCCGGTGGTATGTACTGCCCGGAATGCGGCACAGTTCACTGGCCGGATGGCGTTACCCCACCCGTCTGATGCTTCCCCGTTTTGCCGATATTTTTCAGCAGGGTAACCGCTGGCTTAACTGGCTGGAGAAACAGCCGGAAGGTTCAGTGCGTCCGGTGGTGACTGAGTCAGTGACAAAAATCATGGCATGCGGGACCACGCTGATGGGCTACACGCAATGGTGCTGTTCGTCACCGGACTGTTGCCACACCAAAAAGGTCTGCTTCCGGTGTAAAAGCCGCTCCTGTCCGCACTGCGGGGTGAAGGCTGGCGCACAGTGGATACAGTATCTGCTGAGCCTGGTCCCCGACTGCCCGTGGCAGCATATTGTGTTCACACTTCCCTGCCAGTACTGGTCCCTGGTGTTCCACAACCGGTGGTTACTGGCAGAGATGAGCCGCATTGCAGCGGATGTGATACTGGAAATCTGCCATCAGACAGATGTGGAGCCGGGGATATTCACGGTGATCCACACATGGGGGCGTGACCAGCAGTGGCATCCGCATATCCATTTATCGACAACTGCCGGTGGTGTGACGTCGGGCCACACCTGGAAAAATCTTCATTTTTACGCCCGTAAGGTGATGAGCATGTGGCGTTACCGGATAACGCGGCTACTGTCCCGGAAATACCCGGAGCTGGTAATACCGGATGAACTGGCAGTGGAAGGAAACAGCAAACGGGACTGGAATCGCTTCCTGGACACGCATTACCGCCGCGGCTGGAATGTCAACATATCCAGGGTGATGGATAACGCCACACATGTGGCGGTGTACTTCGGCTCTTACCTGAAAAAGCCACCGGTGCCGGTGAGTCGGCTGGAGCATTATGCCGGTCAGGATGAAATCGGTCTGCGTTACAACAGTCACCGTACAAAACGGGAAGAATATCTGTTGATGAGTGGCGATGAGTTCATGGAAAGGTTCTCGTGGCATGTGGCGGATAAGGGGTTCCGTATGGTGAGGTACTACGGTTTCCTGAGTCCGGTGAAGCGCCGCTTACTGGAAGAAGTGGTGTACGTCATAACGGAGACGGTGAGAAAAACGGCGATGCAAATCAGGTGGAGAGGGATGTATCAGAGGTTACTGAAGGTTGACCCGCTGAAGTGCATTCTGTGCGGAAGTCAGATGCGTTTTACGGGGCTGAAGCGGGGTTACCGACTGGCAGAGCTGATCCTGATGCATGAGCGACTGGCACGACAGCAGGTGTGCGGCTGAGAGCCGCAGAGGGGAAGTTGCGTCCATTTTACCGGAAACGGAGCAAAAAACCGCCATTCATACCCTGTATCAATCAGTGTCATCCTGTTTAATAGTCGTTTCCGCTCATATGGTGCACAAGGGGTGTTGAAGAAACATCCGTTTTGTGGTGCTTTTTTAGTCTTTTGGGGATTTAAATTCCTATCGAT

>IS91-V53

CGAGTAGGCAGCCTGGCGGCTGCGGCTTGTCATGGCCTGGAATTACCGTTATAAAAAAAGATAATGTCATTGTCTTTCAGGTAGTTATATGGCCCGTTCAGCTAAACCCCGTAAACGCAAACCCGCACAACAAAGAAGCAAACTTCCCCGCTATGTTGTGAAACTTCATCCGGATGATTTTTTTGACGAAGAAGACGCAGAAGTTCTGCGCTTTGATAATTTTGACGATGCCGTTGAGTGCTGCGCTGACCTGGGTATTCCGTTCTTTCTGGATGCTGGAAACAAAAAGCTGGTCTTCTGGTTTGTTCGTGTCGATGACGAAGGGTATCCGGAAATAGCTCGCTGTACGGAGCGGGAGTTTGCAACCATTCTTGCCGGTATCAGTGCCGGTGGTATGTACTGCCCGGAATGCGGCACAGTTCACTGGCCGGATGGCGTTACCCCACCCGTCTGATGCTTCCCCGTTTTGCCGATATTTTTCAGCAGGGTAACCGCTGGCTTAACTGGCTGGAGAAACAGCCGAAAGGTTCAGTGCGTCCGGTGGTGACTGAGTCAGTGGCAAAAATCATGGCATGCGGGACCACGCCGATGGGCTACACACAATGGTGCTGTTCGTCACCGGACTGTTGCCACACCAAAAAGGTCTGCTTCCGGTGTAAAAGCCGCTCCTGTCCGCACTGCGGGGTGAAGGCTGGCGCACAGTGGATACAGTATCTGCTGAGCCTGGTCCCCGACAGCCCGTGGCAGCATATTGTGTTCACACTTCCCTGCCAGTACTGGTCCCTGGTGTTCCACAACCGGTGGTTACTGGCAGAGATGAGCCGCATTGCAGCTGATGTGATACTGGAAATCTGCCATCAGGCAGATGTTGAGCCGGGGATATTCACGGTGATCCACACATGGGGGCGTGACCAGCAGTGGCATCCGCATATCCATTTATCGACAACTGCCGGTGGTGTGACGTCGGGCCACACCTGGAAAAATCTTTATTTTTACGCCCGTAAGGTGATGAGCATGTGGCGTTACCGGATAACGCGGCTACTGTCCCGGAAATACCCGGAGCTGGTAATACCGGATGAACTGGCAGTGGAAGGAAACAGCAAACGGGACTGGAATCGCTTCCCGGACACGCATTACCGCCGCGGCTGGAATGTCAACATATCCAGGGTGATGGATAACGCCACACATGTGGCGGTGTACTTCGGCTCTTACCTGAAAAAGCCACCGGTGCCGATGAGTCGGCTGGAGCATTATGCCGGTCAGGATGAAATCGGTCTGCGTTACAACAGTCACCGTACAAAACGGGAAGAATACCTGTTGATGAGTGGAGATGAGTTCATGGAAAGGTTCTCCTGGCATGTAGCAGATAAGGGGTTCCGTATGGTGAGGTACTACGGTTTCCTGAGTCCGGTGAAGCGCCGCTTACTGGAAGAAGTTGTGTACGTCATAACGGAGACGGTGAGAAAAACGGCGATGCAAATCAGGTGGAGAGGGATGTATCAGAGGTTACTGAAGGTTGACCCGCTGAAGTGCATTCTGTGCGGAAGTCAGATGCGTTTTACGGGGCTGAAGCGGGGTTACCGACTGGCAGAGCTGGTCCTGATGCATGAGCGACTGGCACGACAGCAGGTGTGCGGCTGAGAGCCGCAGAGGGGAAGTTGCGTCCATTTTACCGGAAACGGAGCAAAAAACCGCCATTCATACCCTGTATCAATCAGTGTCATCCTGTTTAATAGTCGTTTCCGCTCATATGGTGCACAAGGGGGTTGAAGAAACATCCGTTTCGTGGTGCTTTTTTAGTCTTTTGGGGATTTAAATTCCTATCGAT

>IS91-V54

CGAGTAGGCAGCCTGGCGGCTGCGGCTTGTCATGGCCTGGAATTACCGTTATAAAAAAAGATAATGTCATTGTCTTTCAGGTAGTTATATGGCCCGTTCAGCTAAACCCCGTAAACGCAAACCCGCACAACAAAGAAGCAAACTTCCCCGCTATGTTGTGAAACTTCATCCGGATGATTTTTTTGACGAAGAAGACGCAGAAGTTCTGCGCTTTGATAATTTTGACGATGCCGTTGAGTGCTGCGCTGACCTGGGTATTCCGTTCTTTCTGGATGCTGGAAACAAAAAGCTGGTCTTCTGGTTTGTTCGTGTCGATGACGAAGGGTATCCGGAAATAGCTCGCTGTACGGAGCGGGAGTTTGCAACCATTCTTGCCGGTATCAGTGCCGGTGGTATGTACTGCCCGGAATGCGGCACAGTTCACTGGCCGGATGGCGTTACCCCACCCGTCTGATGCTTCCCCGTTTTGCCGATATTTTTCAGCAGGGTAACCGCTGGCTTAACTGGCTGGAGAAACAGCCGAAAGGTTCAGTGCGTCCGGTGGTGACTGAGTCAGTGACAAAAATCATGGCATGCGGGACCACGCCGATGGGCTACACACAATGGTGCTGTTCGTCACCGGACTGTTGCCACACCAAAAAGGTCTGCTTCCGGTGTAAAAGCCGCTCCTGTCCGCACTGCGGGGTGAAGGCTGGCGCACAGTGGATACAGTATCTGCTGAGCCTGGTCCCCGACAGCCCGTGGCAGCATATTGTGTTCACACTTCCCTGCCAGTACTGGTCCCTGGTGTTCCACAACCGGTGGTTACTGGCAGAGATGAGCCGCATTGCAGCTGATGTGATACTGGAAATCTGCCATCAGGCAGATGTTGAGCCGGGGATATTCACGGTGATCCACACATGGGGGCGTGACCAGCAGTGGCATCCGCATATCCATTTATCGACAACTGCCGGTGGTGTGACGTCGGGCCACACCTGGAAAAATCTTTATTTTTACGCCCGTAAGGTGATGAGCATGTGGCGTTACCGGATAACGCGGCTACTGTCCCGGAAATACCCGGAGCTGGTAATACCGGATGAACTGGCAGTGGAAGGAAACAGCAAACGGGACTGGAATCGCTTCCCGGACACGCATTACCGCCGCGGCTGGAATGTCAACATATCCAGGGTGATGGATAACGCCACACATGTGGCGGTGTACTTCGGCTCTTACCTGAAAAAGCCACCGGTGCCGATGAGTCGGCTGGAGCATTATGCCGGTCAGGATGAAATCGGTCTGCGTTACAACAGTCACCGTACAAAACGGGAAGAATACCTGTTGATGAGTGGAGATGAGTTCATGGAAAGGTTCTCCTGGCATGTAGCAGATAAGGGGTTCCGTATGGTGAGGTACTACGGTTTCCTGAGTCCGGTGAAGCGCCGCTTACTGGAAGAAGTTGTGTACGTCATAACGGAGACGGTGAGAAAAACGGCGATGCAAATCAGGTGGAGAGGGATGTATCAGAGGTTACTGAAGGTTGACCCGCTGAAGTGCATTCTGTGCGGAAGTCAGATGCGTTTTACGGGGCTGAAGCGGGGTTACCGACTGGCAGAGCTGGCCCTGATGCATGAGCGACTGGCACGACAGCAGGTGTGCGGCTGAGAGCCGCAGAGGGGAAGTTGCGTCCATTTTACCGGAAACGGAGCAAAAAACCGCCATTCATACCCTGTATCAATCAGTGTCATCCTGTTTAATAGTCGTTTCCGCTCATATGGTGCACAAGGGGGTTGAAGAAACATCCGTTTCGTGGTGCTTTTTTAGTCTTTTGGGGATTTAAATTCCTATCGAT

>IS91-V55

CGAGTAGGCAGCCTGGCGGCTGCGGCTTGTCATGGTCTGCAATTACCGTTATAAAAACAGGCAATATCATTGTCTTTCTGGTGGTTATATGGCCCGTTCAGCTAAACCCCGTAAACGAAAACCTTCCCCACAACGAAGCAAACTTCCCCGCTATGTTGTGAAACTTCATCCGGATGATTTTTTTGACGAAGAAGACGCTGAAGTTCTGCGCTTTGATAATTTTGACGATGCCGTTGAGTGCTGCGCTGACCTGGGTATTCCGTTCTTTCTGGATGCAGGAAACAAAAAGCTGGTCTTCTGGTTTGTTCGTGTCGATGACGAAGGGTATCCGGAAATAGCCCGCTGTACGGAGCGGGAGTTTGCAACCATTCTTGCCGGTATCAGTGCCGGTGGTATGTACTGCCCGGAATGCGGCACAGTTCACTGGCCGGATGGCGTTACCCCACCCGTCTGATGCTTCCCCGTTTTGCCGATATTTTTCAGCAGGGTAACCGCTGGCTTAACTGGCTGGAGAAACAGCCGGAAGGTTCAGTGCGTCCGGTGGTGATTGAGTCAGTGACAAAAATCATGGCATGCGGGACCACGCTGATGGGCTACACGCAATGGTGCTGTTCGTCACCGGACTGTTGCCACACCAAAAAGGTCTGCTTCCGGTGTAAAAGCCGCTCCTGTCCGCACTGCGGGGTGAAGGCTGGCGCACAGTGGATACAGTATCTGCTGAGCCTGGTCCCCGACTGCCCGTGGCAGCATATTGTGTTCACACTTCCCTGCCAGTACTGGTCCCTGGTGTTCCACAACCGGTGGTTACTGGCAGAGATGAGCCGCATTGCAGCGGATGTGATACTGGAAATCTGCCATCAGACAGATGTGGAGCCGGGGATATTCACGGTGATCCACACATGGGGGCGTGACCAGCAGTGGCATCCGCATATCCATTTATCGACAACTGCCGGTGGTGTGACGTCGGGCCACACCTGGAAAAATCTTCATTTTTACGCCCGTAAGGTGATGAGCATGTGGCGTTACCGGATAACGCGGCTACTGTCCCGGAAATACCCGGAGCTGGTAATACCGGATGAACTGGCAGTGGAAGGAAACAGCAAACGGGACTGGAATCGCTTCCTGGACACGCATTACCGCCGCGGCTGGAATGTCAACATATCCAGGGTGATGGATAACGCCACACATGTGGCGGTGTACTTCGGCTCTTACCTGAAAAAGCCACCGGTGCCGGTGAGTCGGCTGGAGCATTATGCCGGTCAGGATGAAATCGGTCTGCGTTACAACAGTCACCGTACAAAACGGGAAGAATATCTGTTGATGAGTGGAGATGAGTTCATGGAAAGGTTCTCCTGGCATGTAGCAGATAAGGGGTTCCGTATGGTGAGGTACTACGGTTTCCTGAGTCCGGTGAAGCGCCGCTTACTGGAAGAAGTGGTGTACGTCATAACGGAGACGGTGAGAAAAACGGCGATGCAAATCAGGTGGAGAGGGATGTATCAGAGGTTACTGAAGGTTGACCCGCTGAAGTGCATTCTGTGCGGAAGTCAGATGCGTTTTACGGGGCTGAAGCGGGGTTACCGACTGGCAGAGCTGGTCCTGATGCATGAGCGACTGGCACGACAGCAGGTGTGCGGCTGAGAGCCGTAGAGGGGAAGTTGCGTCCATTTTACCGGAAACGGAGCAAAAAACCGGCATTCATACCCTGTATCAATCAGTGTCATCCTGTTTAATAGTCGTTTCCGCTCATATGGTGCACAAGGGGTGTTGAAGAAACATCCGTTTTGTGGTGCTTTTTTAGTCTTTTGGGGATTTAAATTCCTATCGAT

>IS91-V56

CGAGTAGGCAGCCTGGCGGCTGCGGCTTGTCATAGCCTGGAATTACCGTTATAAAAAAAGATAATGTCATTGTCTTTCAGGTAGTTATATGGCCCGTTCAGCTAAACCCCGTAAACGCAAACCCGCACCACAAAGAAGCAAACTTCCCCGCTATGTTGTGAAACTTCATCCGGATGATTTTTTTGACGAAGAAGACGCTGAAGTTCTGCACTTTGATAATTTTGACGATGCCGTTGAGTGCTGCGCTGACCTGGGTATTCCGTTCTTTCTGGATGCAGGAAACAAAAAGCTGGTCTTCTGGTTTGTTCGTGTCGATGACGAAGGGTATCCGGAAATAGCCCGCTGTACGGAGCGGGAGTTTGCAACCATTCTTGCCGGTATCAGTGCCGGTGGTATGTACTGCCCGGAATGCGGCACAGTTCACTGGCCGGATGGCGTTACCCCACCCGTCTGATACTTCCCCGTTTTGCCGATATTTTTCAGCAGGGTAACCGCTGGCTTAACTGGCTGGAGAAACAGCCGGAAGGTTCAGTGCGTCCGGTGGTGACTGAGTCAGTGACAAAAATCATGGCATGCGGGACCACGCTGATGGGCTACACGCAATGGTGCTGTTCATCACCGGACTGTTGCCACACAAAAAAGGTCTGCTTCCGGTGTAAAAGTCGCTCCTGCCCGCACTGCGGAGTGAAGGCTGGCGCACAGTGGATACAGTATCTGCTGAGTCTGGTCCCCGACTGCCCGTGGCAGCATATTGTGTTCACACTTCCCTGCCAGTACTGGCCCCTGGTGTTCCACAACAGGTGGTTACTGGCAGAGATGAGCCGCATTGCTGCGGATGTGATACTGGAAATCTGCCGCCAGGCAGATGTGGAGCCGGGGATATTCACGGTGATCCACACATGGGGGCGTGACCAGCAGTGGCATCCGCATATTCATTTATCGACAACTGCCGGTGGTGTGACGTCGGGCCACACCTGGAAAAATCTTCATTTTTACGCCCGTAAGGTGATGAGCATGTGGCGTTACCGGATAACGCGGCTACTGTCCCGGAAATACCCGGAGCTGGTAATACCGGATGAACTGGCAGTGGAAGGAAACAGCAAACGGGACTGGAATCGCTTCCTGGACACGCATTACCGCCGCGGCTGGAATGTCAACATATCCAGGGTGATGGATAACGCCACACATGTGGCGGTGTACTTCGGCTCTTACCTGAAAAAGCCACCGGTGCCGATGAGTCGGCTGGAACATTATGCCGGTCAGGATGAAATCGGTCTGCGTTACAACAGTCACCGTACAAAACGGGAAGAATACCTGTTGATGAGTGGAGATGAGTTCATGGAAAGGTTCTCCTGGCATGTAGCAGATAAGGGGTTCCGTATGGTGAGGTACTACGGTTTCCTGAGTCCGGTGAAGCGCCGCTTACTGGAAGAAGTTGTGTACGTCATAACGGAGACGGTGAGAAAAACGGCGATGCAAATCAGGTGGAGAGGGATGTATCAGAGGTTACTGAAGGTTGACCCGCTGAAGTGCATTCTGTGCGGAAGTCAGATGCGTTTTACGGGGCTGAAGCGGGGTTACCGACTGGCAGAGCTGGTCCTGATGCATGAGCGACTGGCACGACAGCAGGTGTGCGGCTGAGAGCCTCAGAGGGGAAGTTGCGTCCATTTTACCGGAAACGGAGCAAAAAACCGCCATTCATACCCTGTATCAATCAGTGTCATCCTGTTTAATAGTCGTTTCCGCTCATATGGTGCACAAGGGGTGTTGAAGAAACATCCGTTTTGTGGTGCTTTTTTAGTCTTTTGGGGATTTAAATTCCTATCGAT

>IS91-V57

CGAGTAGGCAGCCTGGCGGCTGCGGCTTGTCATGGCCTGGAATTACCGTTATAAAAAAGATAATGTCATTGTCTTTCAGGTAGTTATATGGCCCGTTCAGCTAAACCCCGTAAACGCAAACCCGCACAACAAAGAAGCAAACTTCCCCGCTATGTTGTGAAACTTCATCCGGATGATTTTTTTGACGAAGAAGACGCTGAAGTTCTGCGCTTTGATAATTTTGACGATGCCGTTGAGTGCTGCGCTGACCTGGGTATTCCGTTCTTTCTGGATGCAGGAAACAAAAAGCTGGTCTTCTGGTTTGTTCGTGTCGATGACGAAGGGTATCCGGAAATAGCCCGCTGTACGGAGCGGGAGTTTGCAACCATTCTTGCCGGTATCAGTGTCGGTGGTATGTACTGCCCGGAATGCGGCACAGTTCACTGGCCGGATGGCGTTACCCCACCCGTCTGATGCTTCCCCGTTTTGCCGATATTTTTCAGCAGGGTAACCGCTGGCTTAACTGGCTGGATAAACAGCCGGAAGGTTCAGTGCGTCCGGTGGTGACTGAGTCAGTGACAAAAATCATGGCATGCGGGACCACGCTGATGGGCTACACACAATGGTGCTGTTCGTCACCGGACTGTTGCCACACCAAAAAGGTCTGCTTCCGGTGTAAAAGCCGCTCCTGTCCGCACTGCGGGGTGAAGGCTGGCGCACAGTGGATACAGTATCTGCTGAGCCTGGTCCCCGACTGCCCGTGGCAGCATATTGTGTTCACACTTCCCTGCCAGTACTGGTCCCTGGTGTTCCACAACCGGTGGTTACTGGCAGAGATGAGCCGCATTGCAGCGGATGTGATACTGGAAATCTGCCGTCAGGCAGATGTGGAGCCGGGGATATTCACGGTGATCCACACATGGGGGCGTGACCAGCAGTGGCATCCGCATATCCATTTATCGACAACTGCCGGTGGTGTGACGTCGGGCCACACCTGGAAAAATCTTCATTTTTACGCCCGTAAGGTGATGAGCATGTGGCGTTACCGGATAACGCGGCTACTGTCCCGGAAATACCCGGAGCTGGTAATACCGGATGAACTGGCAGTGGAAGGAAACAGCAAACGGGACTGGAATCGCTTCCTGGACACGCATTACCGCCGCGGCTGGAATGTCAACATATCCAGGGTGATGGATAACGCCACACATGTGGCGGTGTACTTCGGCTCTTACCTGAAAAAGCCACCGGTGCCGATGAGTCGGCTGGAGCATTATGCCGGTCAGGATGAAATCGGTCTGCGTTACAACAGTCACCGTACAAAACGGGAAGAATACCTGTTGATGAGTGGAGATGAGTTCATGGAAAGGTTCTCCTGGCATGTAGCAGATAAGGGGTTCCGTATGGTGAGGTACTACGGTTTCCTGAGTCCGGTGAAGCGCCGCTTACTGGAAGAAGTTGTGTACGTCATAACGGAGACGGTGAGAAAAACGGCGATGCAAATCAGGTGGAGAGGGATGTATCAGAGGTTACTGAAGGTTGACCCGCTGAAGTGCATTCTGTGCGGAAGTCAGATGCGCTTTACGGGGCTGAAACGGGGTTACCGACTGGCAGAGCTGGTCCTGATGCATGAGCGACTGGCACGACAGCAGGTGTGCGGCTGAGAGCCGCAGTTGCGTCCATTTTACCGGAAACGGAGCAAAAAACCGCCATTCATACCCTGTATCAATCAGTGTCATCCTGTTTAATAGTCGTTTTCGCTCATATGGTGCACAAGGGGTGTTGAAGAAACATCCGTTTTGTGGTGCTTTTTTAGTCTTATGGGGATTTAAATTCCTATCGAT

>IS91-V58

CGAGTAGGCAGCCTGGCGGCTGCGGCTTGTCATGGCCTGGAATTACCGTTATAAAAAAAGATAATGTCATTGTCTTTCAGGTAGTTATATGGCCCGTTCAGCTAAACCCCGTAAACGCAAACCCGCACCACAAAGAAGCAAACTTCCCCGCTATGTTGTGAAACTTCATCCGGATGATTTTTTTGACGAAGAAGACGCTGAAGTTCTGCACTTTGATAATTTTGACGATGCCGTTGAGTGCTGCGCTGACCTGGGTATTCCGTTCTTTCTGGATGCAGGAAACAAAAAGCTGGTCTTCTGGTTTGTTCGTGTCGATGACGAAGGGTATCCGGAAATAGCCCGCTGTACGGAGCGGAAGTTTGCAACCATTCTTGCCGGTATCAGTGCCGGTGGTATGTACTGCCCGGAATGCGGCACAGTTCACTGGCCGGATGGCGTTACCCCACCCGTCTGATGCTTCCCCGTTTTGCCGATATTTTTCAGCAGGGTAACCGCTGGCTTAACTGGCTGGAGAAACAGCCGGAAGGTTCAGTGCGTCCGGTGGTGACTGAGTCAGTGACAAAAATCATGGCATGCGGGACCACGCTGATGGGCTACACGCAATGGTGCTGTTCATCACCGGACTGTTGCCACACAAAAAAGGTCTGCTTCCGGTGTAAAAGTCGCTCCTGCCCGCACTGCGGAGTGAAGGCTGGCGCACAGTGGATACAGTATCTGCTGAGTCTGGTCCCCGACTGCCCGTGGCAGCATATTGTGTTCACACTTCCCTGCCAGTACTGGCCCCTGGTGTTCCACAACAGGTGGTTACTGGCAGAGATGAGCCGCATTGCTGCGGATGTGATACTGGAAATCTGCCGCCAGGCAGATGTGGAGCCGGGGATATTCACGGTGATCCACACATGGGGGCGTGACCAGCAGTGGCATCCGCATATTCATTTATCGACAACTGCCGGTGGTGTGACGTCGGGCCACACCTGGAAAAATCTTCATTTTTACGCCCGTAAGGTGATGAGCATGTGGCGTTACCGGATAACGCGGCTACTGTCCCGGAAATACCCGGAGCTGGTAATACCGGATGAACTGGCAGTGGAAGGAAACAGCAAACGGGACTGGAATCGCTTCCTGGACACGCATTACCGCCGCGGCTGGAATGTCAACATATCCAGGGTGATGGATAACGCCACACATGTGGCGGTGTACTTCGGCTCTTACCTGAAAAAGCCACCGGTGCCGATGAGTCGGCTGGAACATTATGCCGGTCAGGATGAAATCGGTCTGCGTTACAACAGTCACCGTACAAAACGGGAAGAATACCTGTTGATGAGTGGAGATAAGTTCATGGAAAGGTTCTCCTGGCATGTAGCAGATAAGGGGTTCCGTATGGTGAGGTACTACGGTTTCCTGAGTCCGGTGAAGCGCCGCTTACTGGAAGAAGTTGTGTACGTCATAACGGAGACGGTGAGAAAAACGGCGATGCAAATCAGGTGGAGAGGGATGTATCAGAGGTTACTGAAGGTTGACCCGCTGAAGTGCATTCTGTGCGGAAGTCAGATGCGTTTTACGGGGCTGAAGCGGGGTTACCGACTGGCAGAGCTGGTCCTGATGCATGAGCGACTGGCACGACAGCAGGTGTGCGGCTGAGAGCCTCAGAGGGGAAGTTGCGTCCATTTTACCGGAAACGGAGCAAAAAACCGCCATTCATACCCTGTATCAATCAGTGTCATCCTGTTTAATAGTCGTTTCCGCTCATATGGTGCACAAGGGGTGTTGAAGAAACATCCGCTTTGTGGTGCTTTTTTAGTCTTTTGGGGATTTAAATTCCTATCGAT

>IS91-V59

CGAGTAGGCAGCCTGGCGGCTGCGGCTTGTCATGGCCTGGAATTACCGTTATAAAAAAAGATAATGTCATTGTCTTTCAGGTAGTTATATGGCCCGTTCAGCTAAACCCCGTAAACGCAAACCCGCACCACAAAGAAGCAAACTTCCCCGCTATGTTGTGAAACTTCATCCGGATGATTTTTTTGACGAAGAAGACGCTGAAGTTCTGCACTTTGATAATTTTGACGATGCCGTTGAGTGCTGCGCTGACCTGGGTATTCCGTTCTTTCTGGATGCAGGAAACAAAAAGCTGGTCTTCTGGTTTGTTCGTGTCGATGACGAAGGGTATCCGGAAATAGCCCGCTGTACGGAGCGGAAGTTTGCAACCATTCTTGCCGGTATCAGTGCCGGTGGTATGTACTGCCCGGAATGCGGCACAGTTCACTGGCCGGATGGCGTTACCCCACCCGTCTGATGCTTCCCCGTTTTGCCGATATTTTTCAGCAGGGTAACCGCTGGCTTAACTGGCTGGAGAAACAGCCGGAAGGTTCAGTGCGTCCGGTGGTGACTGAGTCAGTGACAAAAATCATGGCATGCGGGACCACGCTGATGGGCTACACGCAATGGTGCTGTTCATCACCGGACTGTTGCCACACAAAAAAGGTCTGCTTCCGGTGTAAAAGTCGCTCCTGCCCGCACTGCGGAGTGAAGGCTGGCGCACAGTGGATACAGTATCTGCTGAGTCTGGTCCCCGACTGCCCGTGGCAGCATATTGTGTTCACACTTCCCTGCCAGTACTGGCCCCTGGTGTTCCACAACAGGTGGTTACTGGCAGAGATGAGCCGCATTGCTGCGGATGTGATACTGGAAATCTGCCGCCAGGCAGATGTGGAGCCGGGGATATTCACGGTGATCCACACATGGGGGCGTGACCAGCAGTGGCATCCGCATATTCATTTATCGACAACTGCCGGTGGTGTGACGTCGGGCCACACCTGGAAAAATCTTCATTTTTACGCCCGTAAGGTGATGAGCATGTGGCGTTACCGGATAACGCGGCTACTGTCCCGGAAATACCCGGAGCTGGTAATACCGGATGAACTGGCAGTGGAAGGAAACAGCAAACGGGACTGGAATCGCTTCCTGGACACGCATTACCGCCGCGGCTGGAATGTCAACATATCCAGGGTGATGGATAACGCCACACATGTGGCGGTGTACTTCGGCTCTTACCTGAAAAAGCCACCGGTGCCGATGAGTCGGCTGGAACATTATGCCGGTCAGGATGAAATCGGTCTGCGTTACAACAGTCACCGTACAAAACGGGAAGAATACCTGTTGATGAGTGGAGATAAGTTCATGGAAAGGTTCTCCTGGCATGTAGCAGATAAGGGGTTCCGTATGGTGAGGTACTACGGTTTCCTGAGTCCGGTGAAGCGCCGCTTACTGGAAGAAGTTGTGTACGTCATAACGGAGACGGTGAGAAAAACGGCGATGCAAATCAGGTGGAGAGGGATGTATCAGAGGTTACTGAAGGTTGACCCGCTGAAGTGCATTCTGTGCGGAAGTCAGATGCGTTTTACGGGGCTGAAGCGGGGTTACCGACTGGCAGAGCTGGTCCTGATGCATGAGCGACTGGCACGACAGCAGGTGTGCGGCTGAGAGCCTCAGAGGGGAAGTTGCGTCCATTTTACCGGAAACGGAGCAAAAAACCGCCATTCATACCCTGTATCAATCAGTGTCATCCTGTTTAATAGTCGTTTCCGCTCATATGGTGCACAAGGGATGTTGAAGAAACATCCGCTTTGTGGTGCTTTTTTAGTCTTTTGGGGATTTAAATTCCTATCGAT

>IS91-V60

CGAGTAGGCAGCCTGGCGGCTGCGGCTTGTCATGGCCTGGAATTACCGTTATAAAAAAAGATAATGTCATTGTCTTTCAGGTAGTTATATGGCCCGTTCAGCTAAACCCCGTAAACGCAAACCCGCACCACAAAGAAGCAAACTTCCCCGCTATGTTGTGAAACTTCATCCGGATGATTTTTTTGACGAAGAAGACGCTGAAGTTCTGCACTTTGATAATTTTGACGATGCCGTTGAGTGCTGCGCTGACCTGGGTATTCCGTTCTTTCTGGATGCAGGAAACAAAAAGCTGGTCTTCTGGTTTGTTCGTGTCGATGACGAAGGGTATCCGGAAATAGCCCGCTGTACGGAGCGGAAGTTTGCAACCATTCTTGCCGGTATCAGTGCCGGTGGTATGTACTGCCCGGAATGCGGCACAGTTCACTGGCCGGATGGCGTTACCCCACCCGTCTGATGCTTCCCCGTTTTGCCGATATTTTTCAGCAGGGTAACCGCTGGCTTAACTGGCTGGAGAAACAGCCGGAAGGTTCAGTGCGTCCGGTGGTGACTGAGTCAGTGACAAAAATCATGGCATGCGGGACCACGCGGATGGGCTACACGCAATGGTGCTGTTCATCACCGGACTGTTGCCACACAAAAAAGGTCTGCTTCCGGTGTAAAAGTCGCTCCTGCCCGCACTGCGGAGTGAAGGCTGGCGCACAGTGGATACAGTATCTGCTGAGTCTGGTCCCCGACTGCCCGTGGCAGCATATTGTGTTCACACTTCCCTGCCAGTACTGGCCCCTGGTGTTCCACAACAGGTGGTTACTGGCGGAGATGAGCCGCATTGCTGCGGATGTGATACTGGAAATCTGCCGTCAGGCAGATGTGGAGCCGGGGATATTCACGGTGATCCACACATGGGGGCGTGACCAGCAGTGGCATCCGCATATTCATTTATCGACAACTGCCGGTGGTGTGACGTCGGGCCACACCTGGAAAAATCTTCATTTTTACGCCCGTAAGGTGATGAGCATGTGGCGTTACCGGATAACGCGGCTACTGTCCCGGAAATACCCGGAGCTGGTAATACCGGATGAACTGGCAGTGGAAGGAAACAGCAAACGGGACTGGAATCGCTTCCTGGACACGCATTACCGCCGCGGCTGGAATGTCAACATATCCAGGGTGATGGATAACGCCACACATGTGGCGGTGTACTTCGGCTCTTACCTGAAAAAGCCACCGGTGCCGATGAGTCGGCTGGAACATTATGCCGGTCAGGATGAAATCGGTCTGCGTTACAACAGTCACCGTACAAAACGGGAAGAATACCTGTTGATGAGTGGAGATAAGTTCATGGAAAGGTTCTCCTGGCATGTAGCAGATAAGGGGTTCCGTATGGTGAGGTACTACGGTTTCCTGAGTCCGGTGAAGCGCCGCTTACTGGAAGAAGTTGTGTACGTCATAACGGAGACGGTGAGAAAAACGGCGATGCAAATCAGGTGGAGAGGGATGTATCAGAGGTTACTGAAGGTTGACCCGCTGAAGTGCATTCTGTGCGGAAGTCAGATGCGTTTTACGGGGCTGAAGCGGGGTTACCGACTGGCAGAGCTGGTCCTGATGCATGAGCGACTGGCACGACAGCAGGTGTGCGGCTGAGAGCCTCAGAGGGGAAGTTGCGTCCATTTTACCGGAAACGGAGCAAAAAACCGCCATTCATACCCTGTATCAATCAGTGTCATCCTGTTTAATAGTCGTTTCCGCTCATATGGTGCACAAGGGGTGTTGAAGAAACATCCGCTTTGTGGTGCTTTTTTAGTCTTTTGGGGATTTAAATTCCTATCGAT

>IS91-V61

CGAGTAGGCAGCCTGGCGGCTGCGGCTTGTCATGGTCTGGAATTACCGTTATAAAAAAAGATAATGTCATTGTCTTTCAGGTAGTTATATGGCCCGTTCAGCTAAACCCCGTAAACGCAAACCCGCACCACAAAGAAGCAAACTTCCCCGCTATGTTGTGAAACTTCATCCGGATGATTTTTTTGACGAAGAAGACGCTGAAGTTCTGCGCTTTGATAATTTTGACGATGCCGTTGAGTGCTGCGCTGACCTGGGTATTCCGTTCTTTCTGGATGCAGGAAACAAAAAGCTGGTCTTCTGGTTTGTTCGTGTCGATGACGAAGGGTATCCGGAAATAGCCCGCTGTACGGAGCGGGAGTTTGCAACCATTCTTGCCGGTATCAGTGCCGGCGGCATGTACTGCCCGGAGTGTGGCACGGTTCACTGGCCGGACGGAGTCCCCCCCGCCCTTCTGATGCTTCCCCGTTTTGCCGACATTTTTCAGCAGGGAAACCGCTGGCTTAACTGGCTGGAGAAACAACCGGAAGGTTCAGTGCGTCCGGTAGTCATTGAGTCGGTGACAAAAATCATGGCGTGCGGAACCACGCTGATGGGGTACACACAGTGGTGCTGTTCATCTCCGGACTGTTGCCACACAAAAAAGGTCTGCTTCCGGTGTAAAAGTCGCTCCTGCCCGCACTGCGGAGTGAAGGCTGGCGCACAGTGGATACAGTATCTGCTGAGCCTGGTCCCCGACTGCCCGTGGCAGCATATTGTGTTCACACTTCCCTGCCAGTACTGGTCCCTGGTGTTCCACAACCGGTGGTTACTGGCAGAGATGAGCCGCATTGCAGCGGATGTGATACTGGAAATCTGCCATCAGACAGATGTGGAGCCGGGGATATTCACGGTGATCCACACATGGGGGCGTGACCAGCAGTGGCATCCGCATATCCATTTATCGACAACTGCCGGTGGTGTGACGTCGGGCCACACCTGGAAAAATCTTCATTTTTACGCCCGTAAGGTGATGAGCATGTGGCGTTACCGGATAACGCGGCTACTGTCCCGGAAATACCCGGAGCTGGTAATACCGGATGAACTGGCAGTGGAAGGAAACAGCAAACGGGACTGGAATTGCTTCCTGGACACGCATTACCGCCGCGGCTGGAATGTCAACATATCCAGGGTGATGGATAACGCCACACATGTGGCGGTGTACTTCGGCTCTTACCTGAAAAAACCGCCGGTGCCGATGAGCCGTCTGGAGCACTATGCTGGTCAGGATGAAATTGGTCTGCGTTACAACAGTCACCGGACAAAACGGGAAGAATACCTGGTGATGAGTGGTGATGAGTTTATGGAAAGGTTCTCCTGGCATGTGGCGGATAAGGGGTTCCGTATGGTGAGGTACTACGGTTTCCTGAGTCCGGTAAAGCGCCGGTTACTGGAAGATGTTGTGTACGTCATAACGGAGACGGTGAGAAAGACGGCGATGCAAATCAGGTGGAGAGGGATGTATCAGAGGTTACTGAAGGTTGACCCGCTGAAGTGCATTCTGTGCGGAAGTCAGATGCGTTTTACGGGGCTGAAGCGGGGTTACCGACTGGCAGAGCTGGTCCTGATGCATGAGCGACTGGCACGACAGCAGGTGTGCGGCTGAGAGCCGCAGAGGGGAAGTTGCGTCCATTTTACCGGAAACGGAGCAAAAAACCGCCATTCATACCCTGTATCAATCAGTGTCATCCTGTTTAATAGTCGTTTCCGCTCATATGGTGCACAAGGGGTGTTGAAGAAATATCCGTTTTGTGGTGCTTTTTTAGTCTTTTGGGGATTTAAATTCCTATCGAT

>IS91-V62

CGAGTAGGCAGCCTGGCGGCTGCGGCTTGTCATGGCCTGGAATTACCGTTATAAAAAAAGATAATGTCATTGTCTTTCAGGTAGTTATATGGCCCGTTCAGCTAAACCCCGTAAACGCAAACCCGCACCACAAAGAAGCAAACTTCCCCGCTATGTTGTGAAACTTCATCCGGATGATTTTTTTGACGAAGAAGACGCTGAAGTTCTGCACTTTGATAATTTTGACGATGCCGTTGAGTGCTGCGCTGACCTGGGTATTCCGTTCTTTCTGGATGCTGGAAACAAAAAGCTGGTCTTCTGGTTTGTTCGTGTCGATGACGAAGGGTATCCGGAAATAGCCCGCTGTACGGAGCGGAAGTTTGCAACCATTCTTGCCGGTATCAGTGCCGGTGGTATGTACTGCCCGGAATGCGGCACAGTTCACTGGCCGGATGGCGTTACCCCACCCGTCTGATACTTCCCCGTTTTGCCGATATTTTTCAGCAGGGTAACCGCTGGCTTAACTGGCTGGAGAAACAGCCGGAAGGTTCAGTGCGTCCGGTGGTGACTGAGTCAGTGACAAAAATCATGGCATGCGGGACCACGCTGATGGGCTACACGCAATGGTGCTGTTCATCACCGGACTGTTGCCACACAAAAAAGGTCTGCTTCCGGTGTAAAAGTCGCTCCTGCCCGCACTGCGGAGTGAAGGCTGGCGCACAGTGGATACAGTATCTGCTGAGTCTGGTCCCCGACTGCCCGTGGCAGCATATTGTGTTCACACTTCCCTGCCAGTACTGGCCCCTGGTGTTCCACAACAGGTGGTTACTGGCAGAGATGAGCCGCATTGCTGCGGATGTGATACTGGAAATCTGCCGCCAGGCAGATGTGGAGCCGGGGATATTCACGGTGATCCACACATGGGGGCGTGACCAGCAGTGGCATCCGCATATTCATTTATCGACAACTGCCGGTGGTGTGACGTCGGGCCACACCTGGAAAAATCTTCATTTTTACGCCCGTAAGGTGATGAGCATGTGGCGTTACCGGATAACGCGGCTACTGTCCCGGAAATACCCAGAGCTGGTAATACCGGATGAACTGGCAGTGGAAGGAAACAGCAAACGGGACTGGAATCGCTTCCTGGACACGCATTACCGCCGCGGCTGGAATGTCAACATATCCAGGGTGATGGATAACGCCACACATGTGGCGGTGTACTTCGGCTCTTACCTGAAAAAGCCACCGGTGCCGATGAGTCGGCTGGAACATTATGCCGGTCAGGATGAAATCGGTCTGCGTTACAACAGTCACCGTACAAAACGGGAAGAATACCTGTTGATGAGTGGAGATAAGTTCATGGAAAGGTTCTCCTGGCATGTAGCAGATAAGGGGTTCCGTATGGTGAGGTACTACGGTTTCCTGAGTCCGGTGAAGCGCCGCTTACTGGAAGAAGTTGTGTACGTCATAACGGAGACGGTGAGAAAAACGGCGATGCAAATCAGGTGGAGAGGGATGTATCAGAGGTTACTGAAGGTTGACCCGCTGAAGTGCATTCTGTGCGGAAGTCAGATGCGTTTTACGGGGCTGAAGCGGGGTTACCGACTGGCAGAGCTGGTCCTGATGCATGAGCGACTGGCACGACAGCAGGTGTGCGGCTGAGAGCCTCAGAGGGGAAGTTGCGTCCATTTTACCGGAAACGGAGCAAAAAACCGCCATTCATACCCTGTATCAATCAGTGTCATCCTGTTTAATAGTCGTTTCCGCTCATATGGTGCACAAGGGGTGTTGAAGAAACATCCGCTTTGTGGTGCTTTTTTAGTCTTTTGGGGATTTAAATTCCTATCGAT

>IS91-V63

CGAGTAGGCAGCCTGGCGGCTGCGGCTTGTCATGGCCTGGAATTACCGTTATAAAAAAAGATAATGTCATTGTCTTTCAGGTAGTTATATGGCCCGTTCAGCTAAACCCCGTAAACGCAAACCCGCACCACAAAGAAGCAAACTTCCCCGCTATGTTGTGAAACTTCATCCGGATGATTTTTTTGACGAAGAAGACGCTGAAGTTCTGCACTTTGATAATTTTGACGATGCCGTTGAGTGCTGCGCTGACCTGGGTATTCCGTTCTTTCTGGATGCAGGAAACAAAAAGCTGGTCTTCTGGTTTGTTCGTGTCGATGACGAAGGGTATCCGGAAATAGCCCGCTGTACGGAGCGGAAGTTTGCAACCATTCTTGCCGGTATCAGTGCCGGTGGTATGTACTGCCCGGAATGCGGCACAGTTCACTGGCCGGATGGCGTTACCCCACCCGTCTGATGCTTCCCCGTTTTGCCGATATTTTTCAGCAGGGTAACCGCTGGCTTAACTGGCTGGAGAAACAGCCGGAAGGTTCAGTGCGTCCGGTGGTGACTGAGTCAGTGACAAAAATCATGGCATGCGGGACCACGCTGATGGGCTACACGCAATGGTGCTGTTCATCACCGGACTGTTGCCACACAAAAAAGGTCTGCTTCCGGTGTAAAAGTCGCTCCTGCCCGCACTGCGGAGTGAAGGCTGGCGCACAGTGGATACAGTATCTGCTGAGTCTGGTCCCCGACTGCCCGTGGCAGCATATTGTGTTCACACTTCCCTGCCAGTACTGGCCCCTGGTGTTCCACAACAGGTGGTTACTGGCAGAGATGAGCCGCATTGCTGCGGATGTGATACTGGAAATCTGCCGCCAGGCAGATGTGGAGCCGGGGATATTCACGGTGATCCACACATGGGGGCGTGACCAGCAGTGGCATCCGCATATTCATTTATCGACAACTGCCGGTGGTGTGACGTCGGGCCACACCTGGAAAAATCTTCATTTTTACGCCCGTAAGGTGATGAGCATGTGGCGTTACCGGATAACGCGGCTACTGTCCCGGAAATACCCGGAGCTGGTAATACCGGATGAACTGGCAGTGGAAGGAAACAGCAAACGGGACTGGAATCGCTTCCTGGACACGCATTACCGCCGCGGCTGGAATGTCAACATATCCAGGGTGATGGATAACGCCACACATGTGGCGGTGTACTTCGGCTCTTACCTGAAAAAGCCACCGGTGCCGATGAGTCGGCTGGAACATTATGCCGGTCAGGATGAAATCGATCTGCGTTACAACAGTCACCGTACAAAACGGGAAGAATACCTGTTGATGAGTGGAGATAAGTTCATGGAAAGGTTCTCCTGGCATGTAGCAGATAAGGGGTTCCGTATGGTGAGGTACTACGGTTTCCTGAGTCCGGTGAAGCGCCGCTTACTGGAAGAAGTTGTGTACGTCATAACGGAGACGGTGAGAAAAACGGCGATGCAAATCAGGTGGAGAGGGATGTATCAGAGGTTACTGAAGGTTGACCCGCTGAAGTGCATTCTGTGCGGAAGTCAGATGCGTTTTACGGGGCTGAAGCGGGGTTACCGACTGGCAGAGCTGGTCCTGATGCATGAGCGACTGGCACGACAGCAGGTGTGCGGCTGAGAGCCTCAGAGGGGAAGTTGCGTCCATTTTACCGGAAACGGAGCAAAAAACCGCCATTCATACCCTGTACCAATCAGTGTCATCCTGTTTAATAGTCGTTTCCGCTCATATGGTGCACAAGGGGTGTTGAAGAAACATCCGCTTTGTGGTGCTTTTTTAGTCTTTTGGGGATTTAAATTCCTATCGAT

>IS91-V64

CGAGTAGGCAGCCTGGCGGCTGCGGCTTGTCATAGCCTGGAATTACCGTTATAAAAAAAGATAATGTCATTGTCTTTCAGGTAGTTATATGGCCCGTTCAGCTAAACCCCGTAAACGCAAACCCGCACCACAAAGAAGCAAACTTCCCCGCTATGTTGTGAAACTTCATCCGGATGATTTTTTTGACGAAGAAGACGCTGAAGTTCTGCACTTTGATAATTTTGACGATGCCGTTGAGTGCTGCGCTGACCTGGGTATTCCGTTCTTTCTGGATGCAGGAAACAAAAAGCTGGTCTTCTGGTTTGTTCGTGTCGATGACGAAGGGTATCCGGGAATAGCCCGCTGTACGGAGCGGGAGTTTGCAACCATTCTTGCCGGTATCAGTGCCGGTGGTATGTACTGCCCGGAATGCGGCACAGTTCACTGGCCGGATGGCGTTACCCCACCCGTCTGATGCTTCCCCGTTTTGCCGATATTTTTCAGCAGGGTAACCGCTGGCTTAACTGGCTGGAGAAACAGCCGGAAGGTTCAGTGCGTCCGGTGGTGACTGAGTCAGTGACAAAAATCATGGCATGCGGGACCACGCTGATGGGCTACACGCAATGGTGCTGTTCATCACCGGACTGTTGCCACACAAAAAAGGTCTGCTTCCGGTGTAAAAGTCGCTCCTGCCCGCACTGCGGAGTGAAGGCTGGCGCACAGTGGATACAGTATCTGCTGAGTCTGGTCCCCGACTGCCCGTGGCAGCATATTGTGTTCACACTTCCCTGCCAGTACTGGCCCCTGGTGTTCCACAACAGGTGGTTACTGGCAGAGATGAGCCGCATTGCTGCGGATGTGATACTGGAAATCTGCCGCCATGCAGATGTGGAGCCGGGGATATTCACGGTGATCCACACATGGGGGCGTGACCAGCAGTGGCATCCGCATATTCATTTATCGACAACTGCCGGTGGTGTGACGTCGGGCCACACCTGGAAAAATCTTCATTTTTACGCCCGTAAGGTGATGAGCATGTGGCGTTACCGGATAACGCGGCTACTGTCCCGGAAATACCCGGAGCTGGTAATACCGGATGAACTGGCAGTGGAAGGAAACAGCAAACGGGACTGGAATCGCTTCCTGGACACGCATTACCGCCGCGGCTGGAATGTCAACATATCCAGGGTGATGGATAACACCACACATGTGGCGGTGTACTTCGGCTCTTACCTGAAAAAGCCACCGGTGCCGATGAGTAGGCTGGAACATTATGCCGGTCAGGATGAAATCGGTCTGCGTTACAACAGTCACCGTACAAAACGGGAAGAATACCTGTTGATGAGTGGAGATGAGTTCATGGAAAGGTTCTCCTGGCATGTAGCAGATAAGGGGTTCCGTATGGTGAGGTACTACGGTTTCCTGAGTCCGGTGAAGCGCCGCTTACTGGAAGAAGTTGTGTACGTCATAACGGAGACGGTGAGAAAAACGGCGATGCAAATCAGGTGGAGAGGGATGTATCAGAGGTTACTGAAGGTTGACCCGCTGAAGTGCATTCTGTGCGGAAGTCAGATGCGTTTTACGGGGCTGAAGCGGGGTTACCGACTGGCAGAGCTGGTCCTGATGCATGAGCGACTGGCACGACAGCAGGTGTGCGGCTGAGAGCCTCAGAGGGGAAGTTGCGTCCATTTTACCGGAAACGGAGCAAAAAACCGCCATTCATACCCTGTATCAATCAGTGTCATCCTGTTTAATAGTCGTTTCCGCTCATATGGTGCACAAGGGGTGTTGAAGAAACATCCGTTTTGTGGTGCTTTTTTAGTCTTTTGGGGATTTAAATTCCTATCGAT

>IS91-V65

CGAGTAGGCAGCCTGGCGGCTGCGGCTTGTCATGGCCTGGAATTACCGTTATAAAAAAAGATAATGTCATTGTCTTTCAGGTAGTTATATGGCCCGTTCAGCTAAACCCCGTAAACGCAAACCCGCACCACAAAGAAGCAAACTTCCCCGCTATGTTGTGAAACTTCATCCGGATGATTTTTTTGACGAAGAAGACGCTGAAGTTCTGCACTTTGATAATTTTGACGATGCCGTTGAGTGCTGCGCTGACCTGGGTATTCCGTTCTTTCTGGATGCAGGAAACAAAAAGCTGGTCTTCTGGTTTGTTCGTGTCGATGACGAAGGGTATCCGGAAATAGCCCGCTGTACGGAGCGGAAGTTTGCAACCATTCTTGCCGGTATCAGTGCCGGTGGTATGTACTGCCCGGAATGCGGCACAGTTCACTGGCCGGATGGCGTTACCCCACCCGTCTGATGCTTCCCCGTTTTGCCGATATTTTTCAGCAGGGTAACCGCTGGCTTAACTGGCTGGAGAAACAACCGGAAGGTTCAGTGCGTCCGGTGGTGACTGAGTCAGTGACAAAAATCATGGCATGCGGGACCACGCTGATGGGCTACACGCAATGGTGCTGTTCATCACCGGACTGTTGCCACACAAAAAAGGTCTGCTTCCGGTGTAAAAGTCGCTCCTGCCCGCACTGCGGAGTGAAGGCTGGCGCACAGTGGATACAGTATCTGCTGAGTCTGCTCCCCGACTGCCCGTGGCAGCATATTGTGTTCACACTTCCCTGCCAGTACTGGCCCCTGGTGTTCCACAACAGGTGGTTACTGGCAGAGATGAGCCGCATTGCTGCGGATGTGATACTGGAAATCTGCCGCCAGGCAGATGTGGAGCCGGGGATATTCACGGTGATCCACACATGGGGGCGTGACCAGCAGTGGCATCCGCATATTCATTTATCGACAACTGCCGGTGGTGTGACGTCGGGCCACACCTGGAAAAATCTTCATTTTTACGCCCGTAAGGTGATGAGCATGTGGCGTTACCGGATAACGCGGCTACTGTCCCGGAAATACCCGGAGCTGGTAATACCGGATGAACTGGCAGTGGAAGGAAACAGCAAACGGGACTGGAATCGCTTCCTGGACACGCATTACCGCCGCGGCTGGAATGTCAACATATCCAGGGTGATGGATAACGCCACACATGTGGCGGTGTACTTCGGCTCTTACCTGAAAAAGCCACCGGTGCCGATGAGTCGGCTGGAACATTATGCCGGTCAGGATGAAATCGGTCTGCGTTACAACAGTCACCGTACAAAACGGGAAGAATACCTGTTGATGAGTGGAGATAAGTTCATGGAAAGGTTCTCCTGGCATGTAGCAGATAAGGGGTTCCGTATGGTGAGGTACTACGGTTTCCTGAGTCCGGTGAAGCGCCGCTTACTGGAAGAAGTTGTGTACGTCATAACGGAGACGGTGAGAAAAACGGCGATGCAAATCAGGTGGAGAGGGATGTATCAGAGGTTACTGAAGGTTGACCCGCTGAAGTGCATTCTGTGCGGAAGTCAGATGCGTTTTACGGGGCTGAAGCGGGGTTACCGACTGGCAGAGCTGGTCCTGATGCATGAGCGACTGGCACGACAGCAGGTGTGCGGCTGAGAGCCTCAGAGGGGAAGTTGCGTCCATTTTACCGGAAACGGAGCAAAAAACCGCCATTCATACCCTGTATCAATCAGTGTCATCCTGTTTAATAGTCGTTTCCGCTCATATGGTGCACAAGGGGTGTTGAAGAAACATCCGCTTTGTGGTGCTTTTTTAGTCTTTTGGGGATTTAAATTCCTATCGCT

>IS91-V66

CGAGTAGGCAGCCTGGCGGCTGCGGCTTGTCATGGCCTGGAATTACCGTTATAAAAAAAGATAATGTCATTGTCTTTCAGGTAGTTATATGGCCCGTTCAGCTAAACCCCGTAAACGCAAACCCGCACCACAAAGAAGCAAACTTCCCCGCGTATGTTGTGAAACTTCATCCGGATGATTTTTTTGACGAAGAAGACGCTGAAGTTCTGCACTTTGATAATTTTGACGATGCCGTTGAGTGCTGCGCTGACCTGGGTATTCCGTTCTTTCTGGATGCAGGAAACAAAAAGCTGGTCTTCTGGTTTGTTCGTGTCGATGACGAAGGGTATCCGGAAATAGCCCGCTGTACGGAGCGGAAGTTTGCAACCATTCTTGCCGGTATCAGTGCCGGTGGTATGTACTGCCCGGAATGCGGCACAGTTCACTGGCCGGATGGCCTTACCCCACCCGTCTGATGCTTCCCCGTTTTGCCGATATTTTTCAGCAGGGTAACCGCTGGCTTAACTGGCTGGAGAAACAGCCGGAAGGTTCAGTGCGTCCGGTGGTGACTGAGTCAGTGACAAAAATCATGGCATGCGGGACCACGCTGATGGGCTACACGCAATGGTGCTGTTCATCACCGGACTGTTGCCACACAAAAAAGGTCTGCTTCCGGTGTAAAAGTCGCTCCTGCCCGCACTGCGGAGTGAAGGCTGGCGCACAGTGGATACAGTATCTGCTGAGTCTGGTCCCCGACTGCCCGTGGCAGCATATTGTGTTCACACTTCCCTGCCAGTACTGGCCCCTGGTGTTCCACAACAGGTGGTTACTGGCAGAGATGAGCCGCATTGCTGCGGATGTGATACTGGAAATCTGCCGCCAGGCAGATGTGGAGCCGGGGATATTCACGGTAATCCACACATGGGGGCGTGACCAGCAGTGGCATCCGCATATTCATTTATCGACAACTGCCGGTGGTGTGACGTCGGGCCACACCTGGAAAAATCTTCATTTTTACGCCCGTAAGGTGATGAGCATGTGGCGTTACCGGATAACGCGGCTACTGTCCCGGAAATACCCGGAGCTGGTAATACCGGATGAACTGGCAGTGGAAGGAAACAGCAAACGGGACTGGAATCGCTTCCTGGACACGCATTACCGCCGCGGCTGGAATGTCAACATATCCAGGGTGATGGATAACGCCACACATGTGGCGGTGTACTTCGGCTCTTACCTGAAAAAGCCACCGGTGCCGATGAGTCGGCTGGAACATTATGCCGGTCAGGATGAAATCGGTCTGCGTTACAACAGTCACCGTACAAAACGGGAAGAATACCTGTTGATGAGTGGAGATAAGTTCATGGAAAGGTTCTCCTGGCATGTAGCAGATAAGGGGTTCCGTATGGTGAGGTACTACGGTTTCCTGAGTCCGGTGAAGCGCCGCTTACTGGAAGAAGTTGTGTACGTCATAACGGAGACGGTGAGAAAAACGGCGATGCAAATCAGGTGGAGAGGGATGTATCAGAGGTTACTGAAGGTTGACCCGCTGAAGTGCATTCTGTGCGGAAGTCAGATGCGTTTTACGGGGCTGAAGCGGGGTTACCGACTGGCAGAGCTGGTCCTGATGCATGAGCGACTGGCACGACAGCAGGTGTGCGGCTGAGAGCCTCAGAGGGGAAGTTGCGTCCATTTTACCGGAAACGGAGCAAAAAACCGCCATTCATACCCTGTATCAATCAGTGTCATCCTGTTTAATAGTCGTTTCCGCTCATATGGTGCACAAGGGGTGTTGAAGAAACATCCGCTTTGTGGTGCTTTTTTAGTCTTTTGGGGATTTAAATTCCTATCGAT

>IS91-V67

CGAGTAGGCAGCCTGGCGGCTGCGGCTTGTCATGGCCTGGAATTACCGTTATAAAAAAAGATAATGTCATTGTCTTTCAGGTAGTTATATGGCCCGTTCAGCTAAACCCCGTAAACGCAAACCCGCACCACAAAGAAGCAAACTTCCCCGCTATGTTGTGAAACTTCATCCGGATGATTTTTTTGACGAAGAAGACGCTGAAGTTCTGCACTTTGATAATTTTGACGATGCCGTTGAGTGCTGCGCTGACCTGGGTATTCCGTTCTTTCTGGATGCAGGAAACAAAAAGCTGGTCTTCTGGTTTGTTCGTGTCGATGACGAAGGGTATCCGGAAATAGCCCGCTGTACGAAGCGGAAGTTTGCAACCATTCTTGCCGGTATCAGTGCCGGTGGTATGTACTGCCCGGAATGCGGCACAGTTCACTGGCCGGATGGCGTTACCCCACCCGTCTGATGCTTCCCCGTTTTGCCGATATTTTTCAGCAGGGTAACCGCTGGCTTAACTGGCTGGAGAAACAGCCGGAAGGTTCAGTGCGTCCGGTGGTGACTGAGTCAGTGACAAAAATCATGGCATGCGGGACCACGCTGATGGGCTACACGCAATGGTGCTGTTCATCACCGGACTGTTGCCACACAAAAAAGGTCTGCTTCCGGTGTAAAAGTCGCTCCTGCCCGCACTGCGGAGTGAAGGCTGGCGCACAGTGGATACAGTATCTGCTGAGTCTGGTCCCCGACTGCCCGTGGCAGCATATTGTGTTCACACTTCCCTGCCAGTACTGGCCCCTGGTGTTCCACAACAGGTGGTTACTGGCAGAGATGAGCCGCATTGCTGCGGATGTGATACTGGAAATCTGCCGCCAGGCAGATGTGGAGCCGGGGATATTCACGGTGATCCACACATGGGGGCGTGACCAGCAGTGGCATCCGCATATTCATTTATCGACAACTGCCGGTGGTGTGACGTCGGGCCACACCTGGAAAAATCTTCATTTTTACGCCCGTAAGGTGATGAGCATGTGGCGTTACCGGATAACGCGGCTACTGTCCCGGAAATACCCGGAGCTGGTAATACCGGATGAACTGGCAGTGGAAGGAAACAGCAAACGGGACTGGAATCGCTTCCTGGACACGCATTACCGCCGCGGCTGGAATGTCAACATATCCAGGGTGATGGATAACGCCACACATGTGGCGGTGTACTTCGGCTCTTACCTGAAAAAGCCACCGGTGCCGATGAGTCGGCTGGAACATTATGCCGGTCAGGATGAAATCGGTCTGCGTTACAACAGTCACCGTACAAAACGGGAAGAATACCTGTTGATGAGTGGAGATAAGTTCATGGAAAGGTTCTCCTGGCATGTAGCAGATAAGGGGTTCCTTATGGTGAGGTACTACGGTTTCCTGAGTCCGGTGAAGCGCCGCTTACTGGAAGAAGTTGTGTACGTCATAACGGAGACGGTGAGAAAAACGGCGATGCAAATCAGGTGGAGAGGGATGTATCAGAGGTTACTGAAGGTTGACCCGCTGAAGTGCATTCTGTGCGGAAGTCAGATGCGTTTTACGGGGCTGAAGCGGGGTTACCGACTGGCAGAGCTGGTCCTGATGCATGAGCGACTGGCACGACAGCAGGTGTGCGGCTGAGAGCCTCAGAGGGGAAGTTGCGTCCATTTTACCGGAAACGGAGCAAAAAACCGCCATTCATACCCTGTATCAATCAGTGTCATCCTGTTTAATAGTCGTTTCCGCTCATATGGTGCACAAGAGGTGTTGAAGAAACATCCGCTTTGTGGTGCTTTTTTAGTCTTTTGGGGATTTAAATTCCTATCGAT

>IS91-V68

CGAGTAGGCAGCCTGGCGGCTGCGGCTTGTCATGGCCTGGAATTACCGTTATAAAAAAAGATAATGTCATTGTCTTTCAGGTAGTTATATGGCCCGTTCAGCTAAACCCCGTAAACGCAAACCCGCACCACAAAGAAGCAAACTTCCCCGCTATGTTGTGAAACTTCATCCGGATGATTTTTTTGACGAAGAAGACGCTGAAGTTCTGCACTTTGATAATTTTGACGATGCCGTTGAGTGCTGCGCTGACCTGGGTATTCCGTTCTTTCTGGATGCTGGAAACAAAAAGCTGGTCTTCTGGTTTGTTCGTGTCGATGACGAAGGGTATCCGGAAATAGCCCGCTGTACGGAGCGGAAGTTTGCAACCATTCTTGCCGGTATCAGTGCCGGTGGTATGTACTGCCCGGAATGCGGCACAGTTCACTGGCCGGATGGCGTTACCCCACCCGTCTGATGCTTCCCCGTTTTGCCGATATTTTTCAGCAGGGTAACCGCTGGCTTAACTGGCTGGAGAAACAGCCGGAAGGTTCAGTGCGTCCGGTGGTGACTGAGTCAGTGACAAAAATCATGGCATGCGGGACCACGCTGATGGGCTACACGCAATGGTGCTGTTCATCACCGGACTGTTGCCACACAAAAAAGGTCTGCTTCCGGTGTAAAAGTCGCTCCTGCCCGCACTGCGGAGTGAAGGCTGGCGCACAGTGGATACAGTATCTGCTGAGTCTGGTCCCCGACTGCCCGTGGCAGCATATTGTGTTCACACTTCCCTGCCAGTACTGGCCCCTGGTGTTCCACAACAGGTGGTTACTGGCAGAGATGAGCCGCATTGCTGCGGATGTGATACTGGAAATCTGCCGCCAGGCAGATGTGGAGCCGGGGATATTCACGGTGATCCACACATGGGGGCGTGACCAGCAGTGGCATCCGCATATTCATTTATCGACAACTGCCGGTGGTGTGACGTCGGGCCACACCTGGAAAAATCTTCATTTTTACGCCCGTAAGGTGATGAGCATGTGGCGTTACCGGATAACGCGGCTACTGTCCCGGAAATACCCGGAGCTGGTAATACCGGATGAACTGGCAGTGGAAGGAAACAGCAAACGGGACTGGAATCGCTTCCTGGACACGCATTACCGCCGCGGCTGGAATGTCAACATATCCAGGGTGATGGATAACGCCACACATGTGGCGGTGTACTTCGGCTCTTACCTGAAAAAGCCACCGGTGCCGATGAGTCGGCTGGAACATTATGCCGGTCAGGATGAAATCGGTCTGCGTTACAACAGTCACCGTACAAAACGGGAAGAATACCTGTTGATGAGTGGAGATAAGTTCATGGAAAGGTTCTCCTGGCATGTAGCAGATAAGGGGTTCCGTATGGTGAGGTACTACGGTTTCCTGAGTCCGGTGAAGCGCCGCTTACTGGAAGAAGTTGTGTACGTCATAACGGAGACGGTGAGAAAAACGGCGATGCAAATCAGGTGGAAAGGGATGTATCAGAGGTTACTGAAGGTTGACCCGCTGAAGTGCATTCTGTGCGGAAGTCAGATGCGTTTTACGGGGCTGAAGCGGGGTTACCGACTGGCAGAGCTGGTCCTGATGCATGAGCGACTGGCACGACAGCAGGTGTGCGGCTGAGAGCCTCAGAGGGGAAGTTGCGTCCATTTTACCGGAAACGGAGCAAAAAACCGCCATTCATACCCTGTATCAATCAGGGTCATCCTGTTTAATAGTCGTTTCCGCTCATATGGTGCACAAGGGGTGTTGAAGAAACATCCGCCTTGTGGTGCTTTTTTAGTCTTTTGGGGATTTAAATTCCTATCGAT

>IS91-V69

CGAGTAGGCAGCCTGGCGGCTGCGGCTTGTCATGGCCTGGAATTACCGTTATAAAAAAAGATAATGTCATTGTCTTTCAGGTAGTTATATGGCCCGTTCAGCTAAACCCCGTAAACGCAAACCCGCACCACAAAGAAGCAAACTTCCCCGCTATGTTGTGAAACTTCATCCGGATGATTTTTTTGACGAAGAAGACGCTGAAGTTCTGCACTTTGATAATTTTGACGATGCCGTTGAGTGCTGCTCTGACCTGGGTATTCCGTTCTTTCTGGATGCAGGAAACAAAAAGCTGGTCTTCTGGTTTGTTCGTGTCGATGACGAAGGGTATCCGGAAATAGCCCGCTGTACGGAGCGGAAGTTTGCAACCATTCTTGCCGGTATCAGTGCCGGTGGTATGTACTGCCCGGAATGCGGCACAGTTCACTGGCCGGATGGCGTTACCCCACCCGTCTGATGCTTCCCCGTTTTGCCGATATTTTTCAGCAGGGTAACCGCTGGCTTAACTGGCTGGAGAAACAGCCGGAAGGTTCAGTGCGTCCGGTGGTGACTGAGTCAGTGACAAAAATCATGGCATGCGGGACCACGCTGATGGGCTACACGCAATGGTGCTGTTCATCACCGGACTGTTGCCACACAAAAAAGGTCTGCTTCCGGTGTAAAAGTCGCTCCTGCCCGCACTGCGGAGTGAAGGCTGGCGCACAGTGGATACAGTATCTGCTGAGTCTGGTCCCCGACTGCCCGTGGCAGCATATTGTGTTCACACTTCCCTGCCAGTACTGGCCCCTGGTGTTCCACAACAGGTGGTTACTGGCAGAGATGAGCCGCATTGCTGCGGATGTGATACTGGAAATCTGCCGCCAGGCAGATGTGGAGCCGGGGATATTCACGGTGATCCACACATGGGGGCGTGACCAGCAGTGGCATCCGCATATTCATTTATCGACAACTGCCGGTGGTGTGACGTCGGGCCACACCTGGAAAAATCTTCATTTTTACGCCCGTAAGGTGATGAGCATGTGGCGTTACCGGATAACGCGGCTACTGTCCCGGAAATACCCGGAGCTGGTAATACCGGATGAACTGGCAGTGGAAGGAAACAGCAAACGGGACTGGAATCGCTTCCTGGACACGCATTACCGCCGCGGCTGGAATGTCAACATATCCAGGGTGATGGATAACGCCACACATGTGGTGGTGTACTTCGGCTCTTACCTGAAAAAGCCACCGGTGCCGATGAGTCGGCTGGAACATTATGCCGGTCAGGATGAAATCGGTCTGCGTTACAACAGTCACCGTACAAAACGGGAAGAATACCTGTTGATGAGTGGAGATAAGTTCATGGAAAGGTTCTCCTGGCATGTAGCAGATAAGGGGTTCCGTATGGTGAGGTACTACGGTTTCCTGAGTCCGGTGAAGCGCCGCTTACTGGAAGAAGTTGTGTACGTCATAACGGAGACGGTGAGAAAAACGGCGATGCAAATCAGGTGGAGAGGGATGTATCAGAGGTTACTGAAGGTTGACCCGCTGAAGTGCATTCTGTGCGGAAGTCAGATGCGTTTTACGGGGCTGAAGCGGGGTTACCGACTGGCAGAGCTGCTCCTGATGCATGAGCGACTGGCACGACAGCAGGTGTGCGGCTGAGAGCCTCAGAGGGGAAGTTGCGTCCATTTTACCGGAAACGGAGCAAAAAACCGCCATTCATACCCTGTATCAATCAGTGTCATCCTGTTTAATAGTCGTTTCCGCTCATATGGTGCACAAGGGGTGTTGAAGAAACATCCGCTTTGTGGTGCTTTTTTAGTCTTTTGGGGATTTAAATTCCTATCGAT

>IS91-V70

CGAGTAGGCAGCCTGGCGGCTGCGGCTTGTCATGGCCTGGAATTACCGTTATAAAAAAAGATAATGTCATTGTCTTTCAGGTAGTTATATGGCCCGTTCAGCTAAACCCCGTAAACGCAAACCCGCACCACAAAGAAGCAAACTTCCCCGCGTATGTTGTGAAACTTCATCCGGATGATTTTTTTGACGAAGAAGACGCTGAAGTTCTGCACTTTGATAATTTTGACGATGCCGTTGAGTGCTGCGCTGACCTGGGTATTCCGTTCTTTCTGGATGCAGGAAACAAAAAGCTGGTCTTCTGGTTTGTTCGTGTCGATGACGAAGGGTATCCGGAAATAGCCCGCTGTACGGAGCGGAAGTTTGCAACCATTCTTGCCGGTATCAGTGCCGGTGGTATGTACTGCCCGGAATGCGGCACAGTTCACTGGCCGGATGGCCTTACCCCACCCGTCTGATGCTTCCCCGTTTTGCCGATATTTTTCAGCAGGGTAACCGCTGGCTTAACTGGCTGGAGAAACAGCCGGAAGGTTCAGTGCGTCCGGTGGTGACTGAGTCAGTGACAAAAATCATGGCATGCGGGACCACGCTGATGGGCTACACGCAATGGTGCTGTTCATCACCGGACTGTTGCCACACAAAAAAGGTCTGCTTCCGGTGTAAAAGTCGCTCCTGCCCGCACTGCGGAGTGAAGGCTGGCGCACAGTGGATACAGTATCTGCTGAGTCTGGTCCCCGACTGCCCGTGGCAGCATATTGTGTTCACACTTCCCTGCCAGTACTGGCCCCTGGTGTTCCACAACAGGTGGTTACTGGCAGAGATGAGCCGCATTGCTGCGGATGTGATACTGGAAATCTGCCGCCAGGCAGATGTGGAGCCGGGGATATTCACGGTGATCCACACATGGGGGCGTGACCAGCAGTGGCATCCGCATATTCATTTATCGACAACTGCCGGTGGTGTGACGTCGGGCCACACCTGGAAAAATCTTCATTTTTACGCCCGTAAGGTGATGAGCATGTGGCGTTACCGGATAACGCGGCTACTGTCCCGGAAATACCCGGAGCTGGTAATACCGGATGAACTGGCAGTGGAAGGAAACAGCAAACGGGACTGGAATCGCTTCCTGGACATGCATTACCGCCGCGGCTGGAATGTCAACATATCCAGGGTGATGGATAACGCCACACATGTGGCGGTGTACTTCGGCTCTTACCTGAAAAAGCCACCGGTGCCGATGAGTCGGCTGGAACATTATGCCGGTCAGGATGAAATCGGTCTGCGTTACAACAGTCACCGTACAAAACGGGAAGAATACCTGTTGATGAGTGGAGATAAGTTCATGGAAAGGTTCTCCTGGCATGTAGCAGATAAGGGGTTCCGTATGGTGAGGTACTACGGTTTCCTGAGTCCGGTGAAGCGCCGCTTACTGGAAGAAGTTGTGTACGTCATAACGGAGACGGTGAGAAAAACGGCGATGCAAATCAGGTGGAGAGGGATGTATCAGAGGTTACTGAAGGTTGACCCGCTGAAGTGCATTCTGTGCGGAAGTCAGATGCGTTTTACGGGGCTGAAGCGGGGTTACCGACTGGCAGAGCTGGTCCTGATGCATGAGCGACTGGCACGACAGCAGGTGTGCGGCTGAGAGCCTCAGAGGGGAAGTTGCGTCCATTTTACCGGAAACGGAGCAAAAAACCGCCATTCATACCCTGTATCAATCAGTGTCATCCTGTTTAATAGTCGTTTCCGCTCATATGGTGCACAAGGGGTGTTGAAGAAACATCCGCTTTGTGGTGCTTTTTTAGTCTTTTGGGGATTTAAATTCCTATCGAT

>IS91-V71

CGAGTAGGCAGCCTGGCGGCTGCGGCTTGTCATGGCCTGGAATTACCGTTATAAAAAAAGATAATGTCATTGTCTTTCAGGTAGTTATATGGCCCGTTCAGCTAAACCCCGTAAACGCAAACCCGCACCACAAAGAAGCAAACTTCCCCGCTATGTTGTGAAACTTCATCCGGATGATTTTTTTGACGAAGAAGACGCTGAAGTTCTGCACTTTGATAATTTTGACGATGCCGTTGAGTGCTGCGCTGACCTGGGTATTCCGTTCTTTCTGGATGCAGGAAACAAAAAGCTGGTCTTCTGGTTTGTTCGTGTCGATGACGAAGGGTATCCGGAAATAGCCCGCTGTACGGAGCGGAAGTTTGCAACCATTCTTGCCGGTATCAGTGCCGGTGGTATGTACTGCCCGGAATGCGGCACAGTTCACTGGCCGGATGGCGTTACCCCACCCGTCTGATGCTTCCCCGTTTTGCCGATATTTTTCAGCAGGGTAACCGCTGGCTTAACTGGCTGGAGAAACAACCGGAAGGTTCAGTGCGTCCGGTGGTGACTGAGTCAGTGACAAAAATCATGGCATGCGGGACCACGCTGATGGGCTACACGCAATGGTGCTGTTCATCACCGGACTGTTGCCACACAAAAAAGGTCTGCTTCCGGTGTAAAAGTCGCTCCTGCCCGCACTGCGGAGTGAAGGCTGGCGCACAGTGGATACAGTATCTGCTGAGTCTGCTCCCCGACTGCCCGTGGCAGCATATTGTGTTCACACTTCCCTGCCAGTACTGGCCCCTGGTGTTCCACAACAGGTGGTTACTGGCAGAGATGAGCCGCATTGCTGCGGATGTGATACTGGAAATCTGCCGCCAGGCAGATGTGGAGCCGGGGATATTCACGGTGATCCACACATGGGGGCGTGACCAGCAGTGGCATCCGCATATTCATTTATCGACAACTGCCGGTGGTGTGACGTCGGGCCACACCTGGAAAAATCTTCATTTTTACGCCCGTAAGGTGATGAGCATGTGGCGTTACCGGATAACGCGGCTACTGTCCCGGAAATACCCGGAGCTGGTAATACCGGATGAACTGGCAGTGGAAGGAAACAGCAAACGGGACTGGAATCGCTTCCTGGACACGCATTACCGCCGCGGCTGGAATGTCAACATATCCAGGGTGATGGATAACGCCACACATGTGGCGGTGTACTTCGGCTCTTACCTGAAAAAGCCACCGGTGCCGATGAGTCGGCTGGAACATTATGCCGGTCAGGATGAAATCGGTCTGCGTTACAACAGTCACCGTACAAAACGGGAAGAATACCTGTTGATGAGTGGAGATAAGTTCATGGAAAGGTTCTCCTGGCATGTAGCAGATAAGGGGTTCCGTATGGTGAGGTACTACGGTTTCCTGAGTCCGGTGAAGCGCCGCTTACTGGAAGAAGTTGTGTACGTCATAACGGAGACGGTGAGAAAAACGGCGATGCAAATCAGGTGGAGAGGGATGTATCAGAGGTTACTGAAGGTTGACCCGCTGAAGTGCATTCTGTGCGGAAGTCAGATGCGTTTTACGGGGCTGAAGCGGGGTTACCGACTGGCAGAGCTGGTCCTGATGCATGAGCGACTGGCACGACAGCAGGTGTGCGGCTGAGAGCTTCAGAGGGGAAGTTGCGTCCATTTTACCGGAAACGGAGCAAAAAACCGCCATTCATACCCTGTATCAATCAGTGTCATCCTGTTTAATAGTCGTTTCCGCTCATATGGTGCACAAGGGGTGTTGAAGAAACATCCGCTTTGTGGTGCTTTTTTAGTCTTTTGGGGATTTAAATTCCTATCGCT

>IS91-V72

CGAGTAGGCAGCCTGGCGGCTGCGGCTTGTCATGGCCTGGAATTACCGTTATAAAAAAAGATAATGTCATTGTCTTTCAGGTAGTTATATGGCCCGTTCAGCTAAACCCCGTAAACGCAAACCCGCACCACAAAGAAGCAAACTTCCCCGCTATGTTGTGAAACTTCATCCGGATGATTTTTTTGACGAAGAAGACGCTGAAGTTCTGCGCTTTGATAATTTTGACGATGCCGTTGAGTGCTGCGCTGACCTGGGTATTCCGTTCTTTCTGGATGCAGGAAACAAAAAGCTGGTCTTCTGGTTTGTTCGTGTCGATGACGAAGGGTATCCGGAAATAGCCCGCTGTACGGAGCGGGAGTTTGCAACCATTCTTGCCGGTATCAGTGCCGGTGGTATGTACTGCCCGGAATGCGGCACAGTTCACTGGCCGGATGGCGTTACCCCACCCGTCTGATGCTTCCCCGTTTTGCCGATATTTTTCAGCAGGGTAACCGCTGGCTTAACTGGCTGGAGAAACAGCCGGAAGGTTCAGTGCGTCCGGTGGTGACTGAGTCAGTGACAAAAATCATGGCATGCGGGACCACGCTGATGGGCTACACGCAATGGTGCTGTTCGTCACCGGACTGTTGCCACACCAAAAAGGTCTGCTTCCGGTGTAAAAGCCGCTCCTGTCCGCACTGCGGGGTGAAGGCTGGCGCACAGTGGATACAGTATCTGCTGAGCCTGGTCCCCGACTGCCCGTGGCAGCATATTGTGTTCACACTTCCCTGCCAGTACTGGTCCCTGGTGTTCCACAACCGGTGGTTACTGGCAGAGATGAGCCGCATTGCAGCGGATGTGATACTGGAAATCTGCCATCAGACAGATGTGGAGCCGGGGATATTCACGGTGAACCACACATGGGGGCGTGACCAGCAGTGGCATCCACATATTCATCTGTCGACAACTGCCGGTGGTGTGACGTCGGGCCACACCTGGAAAAATCTTCATTTTTACGCCCGTAAGGTGATGAGCATGTGGCGTTACCGGATAACGCGGCTACTGTCCCGGAAATACCCGGAGCTGGTAATACTGGATGAACTGGCAGTGGAAGGAAACAGCAAACGGGACTGGAATCGCTTCCTGGACACGCATTACCGCCGCGGCTGGAATGTCAACATATCCAGGGTGATGGATAACGCCACACATGTGGCGGTGTACTTTGGCTCTTACCTGAAAAAGCCGCCGGTGCCAATGAGCCGTCTGGAACACTATGCCGGTCAGGATGAAACTGGTCTGCGTTACAACAGCCACCGCACAAAACGGGAAGAATACCTGGTGATGAGTGGCGATGAGTTCATGGAAAGGTTCTCGTGGCATGTGGCAGATAAGGGGTTCCGTATGGTGAGGTACTACGGTTTCCTGAGTCCGGTGAAGCGCCGCTTACTGGAAGAAGTTGTGTACGTCATAACGGAGACGGTGAGAAAAACGGCGATGCAAATCAGGTGGAGAGGGATGTATCAGAGGTTACTGAAGGTTGACCCGCTGAAGTGCATTCTGTGCGGAAGTCAGATGCGTTTTACGGGGCTGAAGCGGGGTTACCGACTGGCAGAGCTGGTCCTGATGCATGAGCGACTGGCACGACAGCAGGTGTGCGGCTGAGAGCCGCAGAGGGGAAGTTGCATCCATTTTACCGGAAACGGAGCAAAAAACCGCCATTCATACTCTGTATCAATCAGTGTCATCCTGTTTAATAGTCGTTTCCGCTCATATGGTGCACAAGGGGTGTTGAAGAAACATCCGTTTTGTGGTGCTTTTTTAGTCTTTTGGGGATTTAAATTCCTATCGAT

>IS91-V73

CGAGTAGGCAGCCTGGCGGCTGCGGCTTGTCATGGCCTGGAATTACCGTTATAAAAAAGATAATGTCATTGTCTTTCAGGTAGTTATATGGCCCGTTCAGCTAAACCCCGTAAACGCAAACCCGCACAACAAAGAAGCAAACTTCCCCGCTATGTTGTGAAACTTCATCCGGATGATTTTTTTGACGAAGAAGACGCTGAAGTTCTGCGCTTTGATAATTTTGACGATGCCGTTGAGTGCTGCGCTGACCTGGGTATTCCGTTCTTTCTGGATGCAGGAAACAAAAAGCTGGTCTTCTGGTTTGTTCGTGTCGATGACGAAGGGTATCCGGAAATAGCCCGCTGTACGGAGCGGGAGTTTGCAACCATTCTTGCCGGTATCAGTGTCGGTGGTATGTACTGCCCGGAATGCGGCACAGTTCACTGGCCGGATGGCGTTACCCCACCCGTCTGATGCTTCCCCGTTTTGCCGATATTTTTCAGCAGGGTAACCGCTGGCTTAACTGGCTGGATAAACAGCCGGAAGGTTCAGTGCGTCCGGTGGTGACTGAGTCAGTGACAAAAATCATGGCATGCGGGACCACGCTGATGGGCTACACACAATGGTGCTGTTCGTCACCGGACTGTTGCCACACCAAAAAGGTCTGCTTCCGGTGTAAAAGCCGCTCCTGTCCGCACTGCGGGGTGAAGGCTGGCGCACAGTGGATACAGTATCTGCTGAGCCTGGTCCCCGACTGCCCGTGGCAGCATATTGTGTTCACACTTCCCTGCCAGTACTGGTCCCTGGTGTTCCACAACCGGTGGTTACTGGCAGAGATGAGCCGCATTGCAGCGGATGTGATACTGGAAATCTGCCGTCAGGCAGATGTGGAGCCGGGGATATTCACGGTGATCCACACATGGGGGCGTGACCAGCAGTGGCATCCGCACATCCATTTATCGACAACTGCCGGTGGTGTGACATCAGGTCACACCTGGAAAAATCTTCATTTTTACGCCCGTAAGGTGATGAGCATGTGGCGTTACCGGATAACGCGGCTACTGTCCCGGAAATACCCGGAGCTGGTAATACCGGATGAACTGGCAGTGGAAGGAAACAGCAAACGGGACTGGAATCGCTTCCTGGACACGCATTACCGCCGCGGCTGGAATGTCAACATATCCAGGGTGATGGATAACGCCACACATGTGGCGGTGTACTTCGGCTCTTACCTGAAAAAGCCACCGGTGCCGATGAGTCGGCTGGAGCATTATGCCGGTCAGGATGAAATCGGTCTGCGTTACAACAGTCACCGTACAAAACGGGAAGAATACCTGTTGATGAGTGGAGATGAGTTCATGGAAAGGTTCTCCTGGCATGTAGCAGATAAGGGGTTCCGTATGGTGAGGTACTACGGTTTCCTGAGTCCGGTGAAGCGCCGCTTACTGGAAGAAGTTGTGTACGTCATAACGGAGACGGTGAGAAAAACGGCGATGCAAATCAGGTGGAGAGGGATGTATCAGAGGTTACTGAAGGTTGACCCGCTGAAGTGCATTCTGTGCGGAAGTCAGATGCGCTTTACGGGGCTGAAACGGGGTTACCGACTGGCAGAGCTGGTCCTGATGCATGAGCGACTGGCACGACAGCAGGTGTGCGGCTGAGAGCCGCAGTTGCGTCCATTTTACCGGAAACGGAGCAAAAAACCGCCATTCATACCCTGTATCAATCAGTGTCATCCTGTTTAATAGTCGTTTTCGCTCATATGGTGCACAAGGGGTGTTGAAGAAACATCCGTTTTGTGGTGCTTTTTTAGTCTTATGGGGATTTAAATTCCTATCGAT

>IS91-V74

CGAGTAGGCAGCCTGGCGGCTGCGGCTTGTCATGGCCTGGAATTACCGTTATAAAAAAAGATAATGTCATTGTCTTTCAGGTAGTTATATGGCCCGTTCAGCTAAACCCCGTAAACGCAAACCCGCACCACAAAGAAGCAAACTTCCCCGCGTATGTTGTGAAACTTCATCCGGATGATTTTTTTGACGAAGAAGACGCTGAAGTTCTGCACTTTGATAATTTTGACGATGCCGTTGAGTGCTGCGCTGACCTGGGTATTCCGTTCTTTCTGGATGCAGGAAACAAAAAGCTGGTCTTCTGGTTTGTTCGTGTCGATGACGAAGGGTATCCGGAAATAGCCCGCTGTACGGAGCGGAAGTTTGCAACCATTCTTGCCGGTATCAGTGCCGGTGGTATGTACTGCCCGGAATGCGGCACAGTTCACTGGCCGGATGGCCTTACCCCACCCGTCTGATGCTTCCCCGTTTTGCCGATATTTTTCAGCAGGGTAACCGCTGGCTTAACTGGCTGGAGAAACAGCCGGAAGGTTCAGTGCGTCCGGTGGTGACTGAGTCAGTGACAAAAATCATGGCATGCGGGACCACGCTGATGGGCTACACGCAATGGTGCTGTTCATCACCGGACTGTTGCCACACAAAAAAGGTCTGCTTCCGGTGTAAAAGTCGCTCCTGCCCGCACTGCGGAGTGAAGGCTGGCGCACAGTGGATACAGTATCTGCTGAGTCTGGTCCCCGACTGCCCGTGGCAGCATATTGTGTTCACACTTCCCTGCCAGTACTGGCCCCTGGTGTTCCACAACAGGTGGTTACTGGCAGAGATGAGCCGCATTGCTGCGGATGTGATACTGGAAATCTGCCGCCAGGCAGATGTGGAGCCGGGGATATTCACGGTGATCCACACATGGGGGCGTGACCAGCAGTGGCATCCGCATATTCATTTATCGACAACTGCCGGTGGTGTGACGTCGGGCCACACCTGGAAAAATCTTCATTTTTACGCCCGTAAGGTGATGAGCATGTGGCGTTACCGGATAACGCGGCTACTGTCCCGGAAATACCCGGAGCTGGTAATACCGGATGAACTGGCAGTGGAAGGAAACAGCAAACGGGACTGGAATCGCTTCCTGGACATGCATTACCGCCGCGGCTGGAATGTCAACATATCCAGGGTGATGGATAACGCCACACATGTGGCGGTGTACTTCGGCTCTTACCTGAAAAAGCCACCGGTGCCGATGAGTCGGCTGGAACATTATGCCGGTCAGGATGAAATCGGTCTGCGTTACAACAGTCACCGTACAAAACGGGAAGAATACCTGTTGATGAGTGGAGATAAGTTCATGGAAAGGTTCTCCTGGCATGTAGCAGATAAGGGGTTCCGTATGGTGAGGTACTACGGTTTCCTGAGTCCGGTGAAGCGCCGCTTACTGGAAGAAGTTGTGTACGTCATAACGGAGACGGTGAGAAAAACGGCGATGCAAATCAGGTGGAGAGGGATGTATCAGAGGTTACTGAAGGTTGACCCGCTGAAGTGCATTCTATGCGGAAGTCAGATGCGTTTTACGGGGCTGAAGCGGGGTTACCGACTGGCAGAGCTGGTCCTGATGCATGAGCGACTGGCACGACAGCAGGTGTGCGGCTGAGAGCCTCAGAGGGGAAGTTGCGTCCATTTTACCGGAAACGGAGCAAAAAACCGCCATTCATACCCTGTATCAATCAGTGTCATCCTGTTTAATAGTCGTTTCCGCTCATATGGTGCACAAGGGGTGTTGAAGAAACATCCGCTTTGTGGTGCTTTTTTAGTCTTTTGGGGATTTAAATTCCTATCGAT

>IS91-V75

CGAGTAGGCAGCCTGGCGGCTGCGGCTTGTCATGGCCTGGAATTACCGTTATAAAAAAAGATAATGTCATTGTCTTTCAGGTAGTTATATGGCCCGTTCAGCTAAACCCCGTAAACGCAAACCCGCACCACAAAGAAGCAAACTTCCCCGCTATGTTGTGAAACTTCATCCGGATGATTTTTTTGACGAAGAAGACGCTGAAGTTCTGCACTTTGATAATTTTGACGATGCCGTTGAGTGCTGCTCTGACCTGGGTATTCCGTTCTTTCTGGATGCAGGAAACAAAAAGCTGGTCTTCTGGTTTGTTCGTGTCGATGACGAAGGGTATCCGGAAATAGCCCGCTGTACGGAGCGGAAGTTTGCAACCATTCTTGCCGGTATCAGTGCCGGTGGTATGTACTGCCCGGAATGCGGCACAGTTCACTGGCCGGATGGCGTTACCCCACCCGTCTGATGCTTCCCCGTTTTGCCGATATTTTTCAGCAGGGTAACCGCTGGCTTAACTGGCTGGAGAAACAGCCGGAAGGTTCAGTGCGTCCGGTGGTGACTGAGTCAGTGACAAAAATCATGGCATGCGGGACCACGCTGATGGGCTACACGCAATGGTGCTGTTCATCACCGGACTGTTGCCACACAAAAAAGGTCTGCTTCCGGTGTAAAAGTCGCTCCTGCCCGCACTGCGGAGTGAAGGCTGGCGCACAGTGGATACAGTATCTGCTGAGTCTGCTCCCCGACTGCCCGTGGCAGCATATTGTGTTCACACTTCCCTGCCAGTACTGGCCCCTGGTGTTCCACAACAGGTGGTTACTGGCAGAGATGAGCCGCATTGCTGCGGATGTGATACTGGAAATCTGCCGCCAGGCAGATGTGGAGCCGGGGATATTCACGGTGATCCACACATGGGGGCGTGACCAGCAGTGGCATCCGCATATTCATTTATCGACAACTGCCGGTGGTGTGACGTCGAGCCACACCTGGAAAAATCTTCATTTTTACGCCCGTAAGGTGATGAGCATGTGGCGTTACCGGATAACGCGGCTACTGTCCCGGAAATACCCGGAGCTGGTAATACCGGATGAACTGGCAGTGGAAGGAAACAGCAAACGGGACTGGAATCGCTTCCTGGACACGCATTACCGCCGCGGCTGGAATGTCAACATATCCAGGGTGATGGATAACGCCACACATGTGGCGGTGTACTTCGGCTCTTACCTGAAAAAGCCACCGGTGCCGATGAGTCGGCTGGAACATTATGCCGGTCAGGATGAAATCGGTCTGCGTTACAACAGTCACCGTACAAAACGGGAAGAATACCTGTTGATGAGTGGAGATAAGTTCATGGAAAGGTTCTCCTGGCATGTAGCAGATAAGGGGTTCCGTATGGTGAGGTACTACGGTTTCCTGAGTCCGGTGAAGCGCCGCTTACTGGAAGAAGTTGTGTACGTCATAACGGAGACGGTGAGAAAAACGGCGATGCAAATCAGGTGGAGAGGGATGTATCAGAGGTTACTGAAGGTTGACCCGCTGAAGTGCATTCTGTGCGGAAGTCAGATGCGTTTTACGGGGCTGAAGCGGGGTTACCGACTGGCAGAGCTGCTCCTGATGCATGAGCGACTGGCACGACAGCAGGTGTGCGGCTGAGAGCCTCAGAGGGGAAGTTGCGTCCATTTTACCGGAAACGGAGCAAAAAACCGCCATTCATACCCTGTATCAATCAGTGTCATCCTGTTTAATAGTCGTTTCCGCTCATATGGTGCACAAGGGGTGTTGAAGAAACATCCGCTTTGTGGTGCTTTTTTAGTCTTTTGGGGATTTAAATTCCTATCGAT

>IS91-V76

CGAGTAGGCAGCCTGGCGGCTGCGGCTTGTCATGGCCTGGAATTACCGTTATAAAAAAAGATAATGTCATTGTCTTTCAGGTAGTTATATGGCCCGTTCAGCTAAACCCCGTAAACGCAAACCCGCACCACAAAGAAGCAAACTTCCCCGCTATGTTGTGAAACTTCATCCGGATGATTTTTTTGACGAAGAAGACGCTGAAGTTCTGCACTTTGATAATTTTGACGATGCCGTTGAGTGCTGCGCTGACCTGGGTATTCCGTTCTTTCTGGATGCAGGAAACAAAAAGCTGGTCTTCTGGTTTGTTCGTGTCGATGACGAAGGGTATCCGGAAATAGCCCGCTGTACGAAGCGGAAGTTTGCAACCATTCTTGCCGGTATCAGTGCCGGTGGTATGTACTGCCCGGAATGCGGCACAGTTCACTGGCCGGATGGCGTTACCCCACCCGTCTGATGCTTCCCCGTTTTGCCGATATTTTTCAGCAGGGTAACCGCTGGCTTAACTGGCTGGAGAAACAGCCGGAAGGTTCAGTGCGTCCGGTGGTGACTGAGTCAGTGACAAAAATCATGGCATGCGGGACCACGCTGATGGGCTACACGCAATGGTGCTGTTCATCACCGGACTGTTGCCACACAAAAAAGGTCTGCTTCCGGTGTAAAAGTCGCTCCTGCCCGCACTGCGGAGTGAAGGCTGGCGCACAGTGGATACAGTATCTGCTGAGTCTGGTCCCCGACTGCCCGTGGCAGCATATTGTGTTCACACTTCCCTGCCAGTACTGGCCCCTGGTGTTCCACAACAGGTGGTTACTGGCAGAGATGAGCCGCATTGCTGCGGATGTGATACTGGAAATCTGCCGCCAGGCAGATGTGGAGTCGGGGATATTCACGGTGATCCACACATGGGGGCGTGACCAGCAGTGGCATCCGCATATTCATTTATCGACAACTGCCGGTGGTGTGACGTCGGGCCACACCTGGAAAAATCTTCATTTTTACGCCCGTAAGGTGATGAGCATGTGGCGTTACCGGATAACGCGGCTACTGTCCCGGAAATACCCGGAGCTGGTAATACCGGATGAACTGGCAGTGGAAGGAAACAGCAAACGGGACTGGAATCGCTTCCTGGACACGCATTACCGCCGCGGCTGGAATGTCAACATATCCAGGGTGATGGATAACGCCACACATGTGGCGGTGTACTTCGGCTCTTACCTGAAAAAGCCACCGGTGCCGATGAGTCGGCTGGAACATTATGCCGGTCAGGATGAAATCGGTCTGCGTTACAACAGTCACCGTACAAAACGGGAAGAATACCTGTTGATGAGTGGAGATAAGTTCATGGAAAGGTTCTCCTGGCATGTAGCAGATAAGGGGTTCCTTATGGTGAGGTACTACGGTTTCCTGAGTCCGGTGAAGCGCCGCTTACTGGAAGAAGTTGTGTACGTCATAACGGAGACGGTGAGAAAAACGGCGATGCAAATCAGGTGGAGAGGGATGTATCAGAGGTTACTGAAGGTTGACCCGCTGAAGTGCATTCTGTGCGGAAGTCAGATGCGTTTTACGGGGCTGAAGCGGGGTTACCGACTGGCAGAGCTGGTCCTGATGCATGAGCGACTGGCACGACAGCAGGTGTGCGGCTGAGAGCCTCAGAGGGGAAGTTGCGTCCATTTTACCGGAAACGGAGCAAAAAACCGCCATTCATACCCTGTATCAATCAGTGTCATCCTGTTTAATAGTCGTTTCCGCTCATATGGTGCACAAGAGGTGTTGAAGAAACATCCGCTTTGTGGTGCTTTTTTAGTCTTTTGGGGATTTAAATTCCTATCGAT

>IS91-V77

CGAGTAGGCAGCCTGGCGGCTGCGGCTTGTCATGGTCTGGAATTACCGTTATAAAAAAAGATAATGTCATTGTCTTTCAGGTAGTTATATGGCCCGTTCAGCTAAACCCCGTAAACGCAAACCCGCACCACAAAGAAGCAAACTTCCCCGCTATGTTGTGAAACTTCATCCGGATGATTTTTTTGACGAAGAAGACGCTGAAGTTCTGCGCTTTGATAATTTTGACGATGCCGTTGAGTGCTGCGCTGACCTGGGTATTCCGTTCTTTCTGGATGCAGGAAACAAAAAGCTGGTCTTCTGGTTTGTTCGTGTCGATGACGAAGGGTATCCGGAAATAGCCCGCTGTACGGAGCGGGAGTTTGCAACCATTCTTGCCGGTATCAGTGCCGGTGGTATGTACTGCCCGGAATGCGGCACAGTTCACTGGCCGGATGGCGTTACCCCACCCGTCTGATGCTTCCCCGTTTTGCCGATATTTTTCAGCAGGGTAACCGCTGGCTTAACTGGCTGGAGAAACAGCCGGAAGGTTCAGTGCGTCCGGTGGTGACTGAGTCAGTGACAAAAATCATGGCATGCGGGACCACGCTGATGGGCTACACGCAATGGTGCTGTTCGTCACCGGACTGTTGCCACACCAAAAAGGTCTGCTTCCGGTGTAAAAGCCGCTCCTGTCCGCACTGCGGGGTGAAGGCTGGCGCACAGTGGATACAGTATCTGCTGAGCCTGGTCCCCGACTGCCCGTGGCAGCATATTGTGTTCACACTTCCCTGCCAGTACTGGTCCCTGGTGTTCCACAACCGGTGGTTACTGGCAGAGATGAGCCGCATTGCAGCGGATGTGATACTGGAAATCTGCCATCAGACAGATGTGGAGCCGGGGATATTCACGGTGATCCACACATGGGGGCGTGACCAGCAGTGGCATCCGCATATCCATTTATCGACAACTGCCGGTGGTGTGACGTCGGGCCACACCTGGAAAAATCTTCATTTTTACGCGCGTAAGGTGATGAGCATGTGGCGTTACCGGATAACGCGGCTACTGTCCCGGAAATACCCGGAGCTGGTAATACCGGATGAACTGGCAGTGGAAGGAAACAGCAAACGGGACTGGAATCGCTTCCTGGACAGTCATTACCGGCGGGGCTGGAATGTCAACGTATCCCGGGTGATGGATAACGCCACACATGTGGCGGTGTACTTCGGCTCTTACCTGAAAAAACCGCCGGTGCCGATGAGCCGTCTGGAGCACTATGCTGGTCAGGATGAAATTGGTCTGCGTTACAACAGTCACCGGACAAAACGGGAAGAATACCTGGTGATGAGTGGTGATGAGTTTATGGAAAGGTTCTCCTGGCATGTGGCGGATAAGGGGTTCCGTATGGTGAGGTACTACGGTTTCCTGAGTCCGGTAAAGCGCCGGTTACTGGAAGATGTTGTGTACGTCATAACGGAGACGGTGAGAAAGACGGCGATGCAAATCAGGTGGAGAGGGATGTATCAGCGGTTACTGAAGGTTGACCCGCTAAAGTGCATCCTGTGCGGATGTCAGATGCGTTTTACGGGGCTGAAGCGGGGCTACCGACTGGCAGAGCTGGTCCTGATGCATGAGCGACTGGCACGACAGCAGGTGTGCGGCTGAGAGCCGCAGAGGGGAAGTTGCGTCCATTTTACCGGAAACGGAGCAAAAAACCGCCATTCATACCCTGTATCAATCAGTGTCATCCTGTTTAATAGTCGTTTCCGCTCATATGGTGCACAAGGGGTGTTGAAGAAACATCCGTTTTGTGGTGCTTTTTTAGTCTTTTGGGGATTTAAATTCCTATCGAT

>IS91-V78

CGAGTAGGCAGCCTGGCGGCTGCGGCTTGTCATGGCCTGGAATTACCGTTATAAAAAAAGATAATGTCATTGTCTTTCAGGTAGTTATATGGCCCGTTCAGCTAAACCCCGTAAACGCAAACCCGCACCACAAAGAAGCAAACTTCCCCGCGTATGTTGTGAAACTTCATCCGGATGATTTTTTTGACGAAGAAGACGCTGAAGTTCTGCACTTTGATAATTTTGACGATGCCGTTGAGTGCTGCGCTGACCTGGGTATTCCGTTCTTTCTGGATGCAGGAAACAAAAAGCTGGTCTTCTGGTTTGTTCGTGTCGATGACGAAGGGTATCCGGAAATAGCCCGCTGTACGGAGCGGAAGTTTGCAACCATTCTTGCCGGTATCAGTGCCGGTGGTATGTACTGCCCAGAATGCGGCACAGTTCACTGGCCGGATGGCCTTACCCCACCCGTCTGATGCTTCCCCGTTTTGCCGATATTTTTCAGCAGGGTAACCGCTGGCTTAACTGGCTGGAGAAACAGCCGGAAGGTTCAGTGCGTCCGGTGGTGACTGAGTCAGTGACAAAAATCATGGCATGCGGGACCACGCTGATGGGCTACACGCAATGGTGCTGTTCATCACCGGACTGTTGCCACACAAAAAAGGTCTGCTTCCGGTGTAAAAGTCGCTCCTGCCCGCACTGCGGAGTGAAGGCTGGCGCACAGTGGATACAGTATCTGCTGAGTCTGGTCCCCGACTGCCCGTGGCAGCATATTGTGTTCACACTTCCCTGCCAGTACTGGCCCCTGGTGTTCCACAACAGGTGGTTACTGGCAGAGATGAGCCGCATTGCTGCGGATGTGATACTGGAAATCTGCCGCCAGGCAGATGTGGAGCCGGGGATATTCACGGTGATCCACACATGGGGGCGTGACCAGCAGTGGCATCCGCATATTCATTTATCGACAACTGCCGGTGGTGTGACGTCGGGCCACACCTGGAAAAATCTTCATTTTTACGCCCGTAAGGTGATGAGCATGTGGCGTTACCGGATAACGCGGCTACTGTCCCGGAAATACCCGGAGCTGGTAATACCGGATGAACTGGCAGTGGAAGGAAACAGCAAACGGGACTGGAATCGCTTCCTGGACATGCATTACCGCCGCGGCTGGAATGTCAACATATCCAGGGTGATGGATAACGCCACACATGTGGCGGTGTACTTCGGCTCTTACCTGAAAAAGCCACCGGTGCCGATGAGTCGGCTGGAACATTATGCCGGTCAGGATGAAATCGGTCTGCGTTACAACAGTCACCGTACAAAACGGGAAGAATACCTGTTGATGAGTGGAGATAAGTTCATGGAAAGGTTCTCCTGGCATGTAGCAGATAAGGGGTTCCGTATGGTGAGGTACTACGGTTTCCTGAGTCCGGTGAAGCGCCGCTTACTGGAAGAAGTTGTGTACGTCATAACGGAGACGGTGAGAAAAACGGCGATGCAAATCAGGTGGAGAGGGATGTATCAGAGGTTACTGAAGGTTGACCCGCTGAAGTGCATTCTGTGCGGAAGTCAGATGCGTTTTACGGGGCTGAAGCGGGGTTACCGACTGGCAGAGCTGGTCCTGATGCATGAGCGACTGGCACGACAGCAGGTGTGCGGCTGAGAGCCTCAGAGGGGAAGTTGCGTCCATTTTACCGGAAACGGAGCAAAAAACCGCCATTCATACCCTGTATCAATCAGTGTCATCCTGTTTAATAGTCGTTTCCGCTCATATGGTGCACAAGGGGTGTTGAAGAAACATCCGCTTTGTGGTGCTTTTTTAGTCTTTTAGGGATTTAAATTCCTATCGAT

>IS91-V79

CGAGTAGGCAGCCTGGCGGCTGCGGCTTGTCATGGCCTGGAATTACCGTTATAAAAAAAGATAATGTCATTGTCTTTCAGGTAGTTATATGGCCCGTTCAGCTAAACCCCGTAAACGCAAACCCGCACCACAAAGAAGCAAACTTCCCCGCTATGTTGTGAAACTTCATCCGGATGATTTTTTTGACGAAGAAGACGCTGAAGTTCTGCACTTTGATAATTTTGACGATGCCGTTGAGTGCTGCGCTGACCTGGGTATTCCGTTCTTTCTGGATGCAGGAAACAAAAAGCTGGTCTTCTGGTTTGTTCGTGTCGATGACGAAGGGTATCCGGAAATAGCCCGCTGTACGAAGCGGAAGTTTGCAACCATTCTTGCCGGTATCAGTGCCGGTGGTATGTACTGCCCGGAATGCGGCACAGTTCACTGGCCGGATGGCGTTACCCCACCCGTCTGATGCTTCCCCGTTTTGCCGATATTTTTCAGCAGGGTAACCGCTGGCTTAACTGGCTGGAGAAACAGCCGGAAGGTTCAGTGCGTCCGGTGGTGACTGAGTCAGTGACAAAAATCATGGCATGCGGGACCACGCTGATGGGCTACACGCAATGGTGCTGTTCATCACCGGACTGTTGCCACACAAAAAAGGTCTGCTTCCGGTGTAAAAGTCGCTCCTGCCCGCACTGCGGAGTGAAGGCTGGCGCACAGTGGATACAGTATCTGCTGAGTCTGGTCCCCGACTGCCCGTGGCAGCATATTGTGTTCACACTTCCCTGCCAGTACTGGCCCCTGGTGTTCCACAACAGGTGGTTACTGGCAGAGATGAGCCGCATTGCTGCGGATGTGATACTGGAAATCTGCCGCCAGGCAGATGTGGAGCCGGGGATATTCACGGTGATCCACACATGGGGGCGTGACCAGCAGTGGCATCCGCATATTCATTTATCGACAACTGCCGGTGGTGTGACGTCGGGCCACACCTGGAAAAATCTTCATTTTTACGCCCGTAAGGTGATGAGCATGTGGCGTTACCGGATAACGCGGCTACTGTCCCGGAAATACCCGGAGCTGGTAATACCGGATGAACTGGCAGTGGAAGGAAACAGCAAACGGGACTGGAATCGCTTCCTGGACACGCATTACCGCCGCGGCTGGAATGTCAACATATCCAGGGTGATGGATAACGCCACACATGTGGCGGTGTACTTCGGCTCTTACCTGAAAAAGCCACCGGTGCCGATGAGTCGGCTGGAACATTATGCCGGTCAGGATGAAATCGGTCTGCGTTACAACAGTCACCGTACAAAACAGGAAGAATACCTGTTGATGAGTGGAGATAAGTTCATGGAAAGGTTCTCCTGGCATGTAGCAGATAAGGGGTTCCTTATGGTGAGGTACTACGGTTTCCTGAGTCCGGTGAAGCGCCGCTTACTGGAAGAAGTTGTGTACGTCATAACGGAGACGGTGAGAAAAACGGCGATGCAAATCAGGTGGAGAGGGATGTATCAGAGGTTACTGAAGGTTGACCCGCTGAAGTGCATTCTGTGCGGAAGTCAGATGCGTTTTACGGGGCTGAAGCGGGGTTACCGACTGGCAGAGCTGGTCCTGATGCATGAGCGACTGGCACGACAGCAGGTGTGCGGCTGAGAGCCTCAGAGGGGAAGTTGCGTCCATTTTACCGGAAACGGAGCAAAAAACCGCCATTCATACCCTGTATCAATCAGTGTCATCCTGTTTAATAGTCGTTTCCGCTCATATGGTGCACAAGAGGTGTTGAAGAAACATCCGCTTTGTGGTGCTTTTTTGGTCTTTTGGGGATTTAAATTCCTATCGAT

>IS91-V80

CGAGTAGGCAGCCTGGCGGCTGCGGCTTGTCATGTCCTGGAATTACCGTTATAAAAAAAGATAATGTCATTGTCTTTCAGGTAGTTATATGGCCCGTTCAGCTAAACCCCGTAAACGCAAACCCGCACCACAAAGAAGCAAACTTCCCCGCTATGTTGTGAAACTTCATCCGGATGATTTTTTTGACGAAGAAGACGCTGAAGTTCTGCACTTTGATAATTTTGACGATGCCGTTGAGTGCTGCGCTGACCTGGGTATTCCGTTCTTTCTGGATGCAGGAAACAAAAAGCTGGTCTTCTGGTTTGTTCGTGTCGATGACGAAGGGTATCCGGAAATAGCCCGCTGTACGGAGCGGAAGTTTGCAACCATTCTTGCCGGTATCAGTGCCGGTGGTATGTACTGCCCGGAATGCGGCACAGTTCACTGGCCGGATGGCGTTACCCCACCCGTCTGATGCTTCCCCGTTTTGCCGATATTTTTCAGCAGGGTAACCGCTGGCTTAACTGGCTGGAGAAACAGCCGGAAGGTTCAGTGCGTCCGGTGGTGACTGAGTCAGTGACAAAAATCATGGCATGCGGAACCACTCTGATGGGCTACACGCAATGGTGCTGTTCATCACCGGACTGTTGCCACACAAAAAAGGTCTGCTTCCGGTGTAAAAGTCGCTCCTGCCCGCACTGCGGAGTGAAGGCTGGCGCACAGTGGATACAGTATCTGCTGAGTCTGGTCCCCGACTGCCCGTGGCAGCATATTGTGTTCACACTTCCCTGCCAGTACTGGCCCCTGGTGTTCCACAACAGGTGGTTACTGGCAGAGATGAGCCGCATTGCTGCGGATGTGATACTGGAAATCTGCCGTCAGGCAGATGTGGAGCCGGGGATATTCACGGTGATCCACACATGGGGGCGTGACCAGCAGTGGCATCCGCATATTCATTTATCGACAACTGCCGGTGGTGTGACGTCGGGCCACACCTGGGAAAATCTTCATTTTTACGCCCGTAAGGTGATGAGCATGTGGCGTTACCGGATAACGCGGCTACTGTCCCGGAAATACCCGGAGCTGGTAATACCGGATGAACTGGCAGTGGAAGGAAACAGCAAACGGGACAGGAATCGCTTCCTGGACACGCATTACCGCCGCGGCTGGAATGTCAACATATCCAGGGTGATGGATAACGCCACACATGTGGCGGTGTACTTCGGCTCTTACCTGAAAAAGCCACCGGTGCCGATGAGTCGGCTGGAACATTATGCCGGTCAGGATGAAATCAGTCTGCGTTACAACAGTCACCGTACAAAACGGGAAGAATACCTGTTGATGAGTGGAGATAAGTTCATGGAAAGGTTCTCCTGGCATGTAGCAGATAAGGGGTTCCGTATGGTGAGGTACTACGGTTTCCTGAGTCCGGTGAAGCGCCGCTTACTGGAAGAAGTTGTGTACGTCATAACGGAGACGGTGAGAAAAACGGCGATGCAAATCAGGTGGAGAGGGATGTATCAGAGGTTACTGAAGGTTGACCCGCTGAAGTGCATTCTGTGCGGAAGTCAGATGCGTTTTACGGGGCTGAAGCGGGGTTACCGACTGGCAGAGCTGGTCCTGATGCATGAGCGACTGGCACGACAGCAGGTGTGCGGCTGAGAGCCTCAGAGGGGAAGTTGCGTCCATTTTACCGGAAACGGAGCAAAAAACCGCCATTCATACCCTGTATCAATCAGTGTCATCCTGTTTAATAGTCGTTTCCGCTCATATGGTGCACAAGGGGTGTTGAAGAAACATCCGCTTTGTGGTGCTTTTTTAGTCTTTTGGGGATTTAAATTCCTATCGAT

>IS91-V81

CGAGTAGGCAGCCTGGCGGCTGCGGCTTGTCATGGCCTGGAATTACCGTTATAAAAAAGATAATGTCATTGTCTTTCAGGTAGTTATATGGCCCGTTCAGCTAAACCCCGTAAACGCAAACCCGCACCACAAAGAAGCAAACTTCCCCGCTATGTTGTGAAACTTCATCCGGATGATTTTTTTGACGAAGAAGACGCTGAAGTTCTGCACTTTGATAATTTTGACGATGCCGTTGAGTGCTGCGCTGACCTGGGTATTCCGTTCTTTCTGGATGCAGGAAACAAAAAGCTGGTCTTCTGGTTTGTTCGTGTCGATGACGAAGGGTATCCGGAAATAGCCCGCTGTACGAAGCGGAAGTTTGCAACCATTCTTGCCGGTATCAGTGCCGGTGGTATGTACTGCCCGGAATGCGGCACAGTTCACTGGCCGGATGGCGTTACCCCACCCGTCTGATGCTTCCCCGTTTTGCCGATATTTTTCAGCAGGGTAACCGCTGGCTTAACTGGCTGGAGAAACAGCCGGAAGGTTCAGTGCGTCCGGTGGTGACTGAGTCAGTGACAAAAATCATGGCATGCGGGACCACGCTGATGGGCTACACGCAATGGTGCTGTTCATCACCGGACTGTTGCCACACAAAAAAGGTCTGCTTCCGGTGTAAAAGTCGCTCCTGCCCGCACTGCGGAGTGAAGGCTGGCGCACAGTGGATACAGTATCTGCTGAGTCTGGTCCCCGACTGCCCGTGGCAGCATATTGTGTTCACACTTCCCTGCCAGTACTGGCCCCTGGTGTTCCACAACAGGTGGTTACTGGCAGAGATGAGCCGCATTGCTGCGGATGTGATACTGGAAATCTGCCGCCAGGCAGATGTGGAGCCGGGGATATTCACGGTGATCCACACATGGGGGCGTGACCAGCAGTGGCATCCGCATATTCATTTATCGACAACTGCCGGTGGTGTGACGTCGGGCCACACCTGGAAAAATCTTCATTTTTACGCCCGTAAGGTGATGAGCATGTGGCGTTACCGGATAACGCGGCTACTGTCCCGGAAATACCCGGAGCTGGTAATACCGGATGAACTGGCAGTGGAAGGAAACAGCAAACGGGACTGGAATCGCTTCCTGGACACGCATTACCGCCGCGGCTGGAATGTCAACATATCCAGGGTGATGGATAACGCCACACATGTGGCGGTGTACTTCGGCTCTTACCTGAAAAAGCCACCGGTGCCGATGAGTCGGCTGGAACATTATGCCGGTCAGGATGAAATCGGTCTGCGTTACAACAGTCACCGTACAAAACAGGAAGAATACCTGTTGATGAGTGGAGATAAGTTCATGGAAAGGTTCTCCTGGCATGTAGCAGATAAGGGGTTCCTTATGGTGAGGTACTACGGTTTCCTGAGTCCGGTGAAGCGCCGCTTACTGGAAGAAGTTGTGTACGTCATAACGGAGACGGTGAGAAAAACGGCGATGCAAATCAGGTGGAGAGGGATGTATCAGAGGTTACTGAAGGTTGACCCGCTGAAGTGCATTCTGTGCGGAAGTCAGATGCGTTTTACGGGGCTGAAGCGGGGTTACCGACTGGCAGAGCTGGTCCTGATGCATGAGCGACTGGCACGACAGCAGGTGTGCGGCTGAGAGCCTCAGAGGGGAAGTTGCGTCCATTTTACCGGAAACGGAGCAAAAAACCGCCATTCATACCCTGTATCAATCAGTGTCATCCTGTTTAATAGTCGTTTCCGCTCATATGGTGCACAAGAGGTGTTGAAGAAACATCCGCTTTGTGGTGCTTTTTTGGTCTTTTGGGGATTTAAATTCCTATCGAT

>IS91-V82

CGAGTAGGCAGCCTGGCGGCTGCGGCTTGTCATGGCCTGGAATTACCGTTATAAAAAAAAGATAATGTCATTGTCTTTCAGGTAGTTATATGGCCCGTTCAGCTAAACCCCGTAAACGCAAACCCGCACCACAAAGAAGCAAACTTCCCCGCTATGTTGTGAAACTTCATCCGGATGATTTTTTTGACGAAGAAGACGCTGAAGTTCTGCACTTTGATAATTTTGACGATGCCGTTGAGTGCTGCGCTGACCTGGGTATTCCGTTCTTTCTGGATGCAGGAAACAAAAAGCTGGTCTTCTGGTTTGTTCGTGTCGATGACGAAGGGTATCCGGAAATAGCCCGCTGTACGGAGCGGAAGTTTGCAACCATTCTTGCCGGTATCAGTGCCGGTGGTATGTACTGCCCGGAATGCGGCACAGTTCACTGGCCGGATGGCGTTACCCCACCCGTCTGATGCTTCCCCGTTTTGCCGATATTTTTCAGCAGGGTAACCGCTGGCTTAACTGGCTGGAGAAACAGCCGGAAGGTTCAGTGCGTCCGGTGGTGACTGAGTCAGTGACAAAAATCATGGCATGCGGGACCACGCTGATGGGCTACACGCAATGGTGCTGTTCATCACCGGACTGTTGCCACACAAAAAAGGTCTGCTTCCGGTGTAAAAGTCGCTCCTGCCCGCACTGCGGAGTGAAGGCTGGCGCACAGTGGATACAGTATCTGCTGAGTCTGGTCCCCGACTGCCCGTGGCAGCATATTGTGTTCACACTTCCCTGCCAGTACTGGCCCCTGGTGTTCCACAACAGGTGGTTACTGGCAGAGATGAGCCGCATTGCTGCGGATGTGATACTGGAAATCTGCCGTCAGGCAGATGTGGAGCCGGGGATATTCACGGTGATCCACACATGGGGGCGTGACCAGCAGTGGCATCCGCATATTCATTTATCGACAACTGCCGGTGGTGTGACGTCGGGCCACACCTGGAAAAATCTTCATTTTTACGCCCGTAAGGTGATGAGCATGTGGCGTTACCGGATAACGCGGCTACTGTCCCGGAAATACCCGGAGCTGGTAATACCGGATGAACTGGCAGTGGAAGGAAACAGCAAACGGGACTGGAATCGCTTCCTGGACACGCATTACCGCCGCGGCTGGAATGTCAACATATCCAGGGTGATGGATAACGCCACACATGTGGCGGTGTACTTCGGCTCTTACCTGAAAAAGCCACCGGTGCCGATGAGTCGGCTGGAACATTATGCCGGTCAGGATGAAATCGGTCTGCGTTACAACAGTCACCGTACAAAACGGGAAGAATACCTGTTGATGAGTGGAGATAAGTTCATGGAAAGGTTCTCCTGGCATGTAGCAGATAAGGGGTTCCGTATGGTGAGGTACTACGGTTTCCTGAGTCCGGTGAAGCGCCGCTTACTGGAAGAAGTTGTGTACGTCATAACGGAGACGGTGAGAAAAACGGCGATGCAAATCAGGTGGAGAGGGATGTATCAGAGGTTACTGAAGGTTGACCCGCTGAAGTGCATTCTGTGCATTCTGTGCGGAAGTCAGATGCGTTTTACGGGGCTGAAGCGGGGTTACCGACTGGCAGAGCTGGTCCTGATGCATGAGCGACTGGCACGACAGCAGGTGTGCGGCTGAGAGCCTCAGAGGGGAAGTTGCGTCCATTTTACCGGAAACGGAGCAAAAAACCGCCATTCATACCCTGTATCAATCAGTGTCATCCTGTTTAATAGTCGTTTCCGCTCATATGGTGCACAAGGGGTGTTGAAGAAACATCCGCTTTGTGGTGCTTTTTTAGTCTTTTGGGGATTTAAATTCCTATCGTT

>IS91-V83

AGAGCCTTCGGGTAGGCAGCCTGACGGCTGCGGCTTGTCATGGTCTGCAATTACCGTTATAAAAACAGGCAATGTCATTGTCTTTCAGGTGGTTATATGGCCCGTTCAGCTAAACCCCGTAAACGTAAATCCGCCTCTCAAAGAAGCAAACTTCCCCGCTATGTTGTGAAACTTCATCCGGATGATTTTTTTGACGAAGAAGACGCTGAAGTTCTGCGCTTTGATAATTTTGACGATGCCGTTGAGTGCTGCGCTGACCTGGGTATTCCGTTCTTTCTGGATGCTGGAAACAAAAAGCTGGTCTTCTGGTTTGTTCGTGTCGATGACGAAGGGTATCCGGAAATAGCCCGCTGTACGGAGCGGGAGTTTGCAACCATTCTTGCCGGTATCAGTGCCGGTGGTATGTACTGCCCGGAATGCGGCACAGTTCACTGGCCGGATGGCGTTACCCCACCCGTCTGATGCTTCCCCGTTTTGCCGATATTTTTCAGCAGGGTAACCGCTGGCTTAACTGGCTGGAGAAACAGCCGAAAGGTTCAGTGCGTCCGGTGGTGACTGAGTCAGTGACAAAAATCATGGCATGCGGGACCACGCCGATGGGCTACACACAATGGTGCTGTTCGTCACCGGACTGTTGCCACACCAAAAAGGTCTGCTTCCGGTGTAAAAGCCGCTCCTGTCCGCACTGCGGGGTGAAGGCTGGCGCACAGTGGATACAGTATCTGCTGAGCCTGGTCCCCGACAGCCCGTGGCAGCATATTGTGTTCACACTTCCCTGCCAGTACTGGTCCCTGGTGTTCCACAACCGGTGGTTACTGGCAGAGATGAGCCGCATTGCAGCTGATGTGATACTGGAAATCTGCCATCAGGCAGATGTTGAGCCGGGGATATTCACGGTGATCCACACATGGGGGCGTGACCAGCAGTGGCATCCGCATATCCATTTATCGACAACTGCCGGTGGTGTGACGTCGGGCCACACCTGGAAAAATCTTTATTTTTACGCCCGTAAGGTGATGAGCATGTGGCGTTACCGGATAACGCGGCTACTGTCCCGGAAATACCCGGAGCTGGTAATACCGGATGAACTGGCAGTGGAAGGAAACAGCAAACGGGACTGGAATCGCTTCCCGGACACGCATTACCGCCGCGGCTGGAATGTCAACATATCCAGGGTGATGGATAACGCCACACATGTGGCGGTGTACTTCGGCTCTTACCTGAAAAAGCCACCGGTGCCGATGAGTCGGCTGGAGCATTATGCCGGTCAGAATGAAATCGGTCTGCGTTACAACAGTCACCGTACAAAACGGGAAGAATACCTGTTGATGAGTGGAGATGAGTTCATGGAAAGGTTCTCCTGGCATGTAGCAGATAAGGGGTTCCGTATGGTGAGGTACTACGGTTTCCTGAGTCCGGTGAAGCGCCGCTTACTGGAAGAAGTTGTGTACGTCATAACGGAGACGGTGAGAAAAACGGCGATGCAAATCAGGTGGAGAGGGATGTATCAGAGGTTACTGAAGGTTGACCCGCTGAAGTGCATTCTGTGCGGAAGTCAGATGCGTTTTACGGGGCTGAAGCGGGGTTACCGACTGGCAGAGCTGGCCCTGATGCATGAGCGACTGGCACGACAGCAGGTGTGCGGCTGAGAGCCGCAGAGGGGAAGTTGCGTCCATTTTACCGGAAACGGAGCAAAAAACCGCCATTCATACCCTGTATCAATCAGTGTCATCCTGTTTAATAGTCGTTTCCGCTCATATGGTGCACAAGGGGGTTGAAGAAACATCCGTTTTGTGGTGTTTTTTAATCTTTTGGGGGTTTAATTCCTGTCGAT

>IS91-V84

CGAGTAGGCGGCCTGACGGCTGCGGCTTGTCATGGTCTGGAATTACCGTTATAAAAAAAGATAATGTCATTGTCTTTCAGGTAGTTATATGGCCCGTTCAGCTAAACCCCGTAAACGCAAACCCGCACCACAAAGAAGCAAACTTCCCCGCTATGTTGTGAAACTTCATCCGGATGATTTTTTTGACGAAGAAGACGCTGAAGTTCTGCGCTTTGATAATTTTGACGATGCCGTTGAGTGCTGCGCAGACCTGAATATTCCCTTCTTTGTGGATGCCGGAAACAAAAAGCTGGTCTTCTGGTTTGTACGTGTTGATGACGAAGGGTATCCTGAAATAGCCCGCTGCACGGAGCGGGAGTTTGCGACCATTCTTGCCGGTATCAGCGCCGGCGGCATGTACTGCCCGGAGTGTGGCACGGTTCACTGGCCGGACGGAGTCCCCCCGCCTTCTGATGCTTCCCCGTTTTGCCGACATTTTTCAGCAGGGTAACCGCTGGCTTAACTGGCTGGAGAAACAGCCGGAAGGTTCAGTGCGTCCGGTGGTGACTGAGTCAGTGACAAAAATCATGGCATGCGGGACCACGCTGATGGGCTACACGCAATGGTGCTGTTCGTCACCGGACTGTTGCCACACCAAAAAGGTCTGCTTCCGGTGTAAAAGCCGCTCCTGTCCGCACTGCGGGGTGAAGGCTGGCGCACAGTGGATACAGTATCTGCTGAGCCTGGTCCCCGACTGCCCGTGGCAGCATATTGTGTTCACACTTCCCTGCCAGTACTGGTCCCTGGTGTTCCACAACCGGTGGTTACTGGCAGAGATGAGCCGCATTGCAGCGGATGTGATACTGGAAATCTGCCATCAGACAGATGTGGAGCCGGGGATATTCACGGTGATCCACACATGGGGGCGTGACCAGCAGTGGCATCCGCATATCCATTTATCGACAACTGCCGGTGGTGTGACGTCGGGCCACACCTGGAAAAATCTTCATTTTTACGCCCGTAAGGTGATGAGCATGTGGCGTTACCGGATAACGCGGCTACTGTCCCGGAAATACCCGGAGCTGGTGATACCGGATGAACTGGCAGTGGAAGGAAACAGCAAACGGGACTGGAATCGCTTCCTGGACACGCATTACCGCCGNGGCTGGAATGTCAACGTATCCCGGGTAATGGATAATGCCACCCATGTGGCGGTGTACTTTGGCTCTTACCTGAAAAAGCCACCGGTGCCGATGAGTCGTCTGGAGCATTATGCCGGTCAGGATGAAATCGGTCTGCGTTACAACAGTCACCGAACAAAACGGGAAGAATACCTGTTGATGAGTGGAGATGAGTTCATGGAAAGGTTCTCCTGGCATGTGGCGGATAAGGGGTTCCGTATGGTGAGGTACTACGGTTTTTTGAGTCCGGCGAAACGGCGGTTACTGGAAGAAGTGGTGTACATCATAACGGAGACAGTGAGAAAAACGGCGATGCAAATCACCTGGAGAGGGATGTATCAGAGGTTACTGAAGGTTGACCCGCTGAAGTGCGTGCTGTGCGGGAGTCAGATGCGTTTTACGGGGCTGAAGCGGGGCTACCGTCTGGCAGAGCAGGTCCTGATGCATGAGCCGCTGGCCCGGATGCGGTGGTGCGGCTGAGAGCCGCAGAGGGGAAGTTGCGTCCATTTTGAGGGAAACGGAGCAAAAAACCGCCATTCATACCCTGAATCAATCAGAGCCATCCAGTTTAATCGTCGGTTCCGTTCATATGGAGCAAAAGTGGTGTTGAAGAAACATCCGTTTTGTGGTGTTTTTTTAATCTTTTTGGGGTTTTAATTCCTATCGAT

>IS91-V85

CGAGTAGGCAGCCTGGCGGCTGCGGCTTGTCATGGTCTGGAATTACCGTTATAAAAAAAGATAATGTCATTGTCTTTCAGGTAGTTATATGGCCCGTTCAGCTAAACCCCGTAAACGCAAACCCGCACCACAAAGAAGCAAACTTCCCCGCTATGTTGTGAAACTTCATCCGGATGATTTTTTTGACGAAGAAGACGCTGAAGTTCTGCGCTTTGATAATTTTGACGATGCCGTTGAGTGCTGCGCTGACCTGGGTATTCCGTTCTTTCTGGATGCAGGAAACAAAAAGCTGGTCTTCTGGTTTGTTCGTGTCGATGACGAAGGGTATCCGGAAATAGCCCGCTGCACGGAGCGGGAGTTTGCGACCATTCTTGCCGGTATCAGCGCCGGCGGCATGTACTGCCCGGAGTGTGGCACGGTTCACTGGCCGGACGGAGTCCCCCCGCCCTTCTGATGCTTCCCCGTTTTGCCGACATTTTTCAGCAGGGAAACCGCTGGCTTAACTGGCTGGAGAAACAACCGGAAGGTTCAGTGCGTCCGGTAGTCATTGAGTCGGTGACAAAAATCATGGCGTGCGGAACCACGCTGATGGGGTACACACAGTGGTGCTGTTCATCTCCGGACTGTTGCCACACAAAAAAGGTCTGCTTCCGGTGTAAAAGTCGCTCCTGCCCGCACTGCGGAGTGAAGGCTGGCGCACAGTGGATACAGTATCTGCTGAGCCTGGTCCCCGACTGCCCGTGGCAGCATATTGTGTTCACACTTCCCTGCCAGTACTGGTCCCTGGTGTTCCACAACCGGTGGTTACTGGCAGAGATGAGCCGCATTGCTGCGGATGTGATACAGGAAATCTGCCGCCAGGCAGATGTGGTGCCGGGGATATTCACGGTCATCCACACATGGGGACGTGACCAGCAGTGGCATCCGCACATTCACCTGTCGACAACGGCCGGCGGCGTGACACCAGACCACACCTGGAAAAACCTTCATTTTTACGCCCGTAAGGTGATGAGCATGTGGCGTTACAGGATAACGTGGTTACTGTCACGAAAATACCCGGAGCTGGTGATACCGGATGCGCTGGCAGTTGAAGGAAGCAGCAAACGGGACTGGAATCGCTTCCTGGACACGCATTACCGCCGCGGCTGGAATGTCAACATATCCAGGGTGATGGATAACGCCACACATGTGGCGGTGTACTTCGGCTCTTACCTGAAAAAGCCACCGGTGCCGATGAGTCGGCTGGAGCATTATGCCGGTCAGGATGAAATCGGTCTGCGTTACAACAGTCACCGTACAAAACGGGAAGAATACCTGTTGATGAGTGGAGATGAGTTCATGGAAAGGTTCTCCTGGCATGTAGCAGATAAGGGGTTCCGTATGGTGAGGTACTACGGTTTCCTGAGTCCGGTGAAGCGCCGCTTACTGGAAGAAGTTGTGTACGTCATAACGGAGACGGTGAGAAAAACGGCGATGCAAATCAGGTGGAGAGGGATGTATCAGAGGTTACTGAAGGTTGACCCGCTGAAGTGCATTCTGTGCGGAAGTCAGATGCGTTTTACGGGGCTGAAGCGGGGTTACCGACTGGCAGAGCTGGTCCTGATGCATGAGCGACTGGCACGACAGCAGGTGTGCGGCTGAGAGCCGCAGAGGGGAAGTTGCGTCCATTTTACCGGAAACGGAGCAAAAAACCGCCATTCATACCCTGTATCAATCAGTGTCATCCTGTTTAATAGTCGTTTCCGCTCATATGGTGCACAAGGGGTGTTGAAGAAACATCCGTTTTGTGGTGCTTTTTTAGTCTTTTGGGGATTTAAATTCCTATCGAT

>IS91-V86

CGAGTAGGCAGCCTGGCGGCTGCGGCTTGTCATGGTCTGGAATTACCGTTATAAAAAAAGATAATGTCATTGTCTTTCAGGTAGTTATATGGCCCGTTCAGCTAAACCCCGTAAACGCAAACCCGCACCACAAAGAAGCAAACTTCCCCGCTATGTTGTGAAACTTCATCCGGATGATTTTTTTGACGAAGAAGACGCTGAAGTTCTGCGCTTTGATAATTTTGACGATGCCGTTGAGTGCTGCGCTGACCTGGGTATTCCGTTCTTTCTGGATGCAGGAAACAAAAAGCTGGTCTTCTGGTTTGTTCGTGTCGATGACGAAGGGTATCCGGAAATAGCCCGCTGTACGGAGCGGGAGTTTGCGACCATTCTTGCCGGTATCAGCGCCGGCGGCATGTACTGCCCGGAGTGTGGCACGGTTCACTGGCCGGACGGAGTCCCCCCGCCCTTCTGATGCTTCCCCGTTTTGCCGACATTTTTCAGCAGGGAAACCGCTGGCTTAACTGGCTGGAGAAACAACCGGAAGGTTCAGTGCGTCCGGTAGTCATTGAGTCGGTGACAAAAATCATGGCGTGCGGAACCACGCTGATGGGGTACACACAGTGGTGCTGTTCATCTCCGGACTGTTGCCACACAAAAAAGGTCTGCTTCCGGTGTAAAAGTCGCTCCTGCCCGCACTGCGGAGTGAAGGCTGGCGCACAGTGGATACAGTATCTGCTGAGCCTGGTCCCCGACTGCCCGTGGCAGCATATTGTGTTCACACTTCCCTGCCAGTACTGGTCCCTGGTGTTCCACAACCGGTGGTTACTGGCAGAGATGAGCCGCATTGCTGCGGATGTGATACAGGAAATCTGCCGCCAGGCAGATGTGGTGCCGGGGATATTCACGGTCATCCACACATGGGGACGTGACCAGCAGTGGCATCCGCACATTCACCTGTCGACAACGGCCGGCGGCGTGACACCAGACCACACCTGGAAAAACCTTCATTTTTACGCCCGTAAGGTGATGAGCATGTGGCGTTACAGGATAACGTGGTTACTGTCACGAAAATACCCGGAGCTGGTGATACCGGATGCGCTGGCAGTTGAAGGAAGCAGCAAACGGGACTGGAATCGCTTCCTGGACACGCATTACCGCCGCGGCTGGAATGTCAACATATCCAGGGTGATGGATAACGCCACACATGTGGCGGTGTACTTCGGCTCTTACCTGAAAAAGCCACCGGTGCCGATGAGTCGGCTGGAGCATTATGCCGGTCAGGATGAAATCGGTCTGCGTTACAACAGTCACCGTACAAAACGGGAAGAATACCTGTTGATGAGTGGAGATGAGTTCATGGAAAGGTTCTCCTGGCATGTAGCAGATAAGGGGTTCCGTATGGTGAGGTACTACGGTTTCCTGAGTCCGGTGAAGCGCCGCTTACTGGAAGAAGTTGTGTACGTCATAACGGAGACGGTGAGAAAAACGGCGATGCAAATCAGGTGGAGAGGGATGTATCAGAGGTTACTGAAGGTTGACCCGCTGAAGTGCATTCTGTGCGGAAGTCAGATGCGTTTTACGGGGCTGAAGCGGGGTTACCGACTGGCAGAGCTGGTCCTGATGCATGAGCGACTGGCACGACAGCAGGTGTGCGGCTGAGAGCCGCAGAGGGGAAGTTGCGTCCATTTTACCGGAAACGGAGCAAAAAACCGCCATTCATACCCTGTATCAATCAGTGTCATCCTGTTTAATAGTCGTTTCCGCTCATATGGTGCACAAGGGGTGTTGAAGAAACATCCGTTTTGTGGTGCTTTTTTAGTCTTTTGGGGATTTAAATTCCTATCGAT

>IS91-V87

CGAGTAGGCAGCCTGGCGGCTGCGGCTTGTCATGGTCTGGAATTACCGTTATAAAAAAAGATAATGTCATTGTCTTTCAGGTAGTTATATGGCCCGTTCAGCTAAACCCCGTAAACGAAAACCTGCCCCTCAAAGAAGCAAACTTCCCCGCTATGTCGTGAAGCTTCACGACGATGACTTCTTTGACGAAGAAGACGCAGAAGCTCTGCGCTTTGATAATTTTGACGATGCCGTTGAGTGCTGCGCAGACCTGAATATTCCCTTCTTTGTGGATGCCGGAAACAAAAAGCTGGTCTTCTGGTTTGTTCGTGTCGATGACGAAGGGTATCCTGAAATAGCCCGCTGCACGGAGCGGGAGTTTGCAACCATTCTTGCCGGTATCAGCGCCGGCGGCATGTACTGCCCGGAGTGTGGCACGGTTCACTGGCCGGACGGAGTCCCCCCGCCCTTCTGATGCTTCCCCGTTTTGCCGACATTTTTCAGCAGGGAAACCGCTGGCTTAACTGGCTGGAGAAACAACCGGAAGGTTCAGTGCGTCCGGTAGTCATTGAGTCGGTGACAAAAATCATGGCGTGCGGAACCACGCTGATGGGGTACACACAGTGGTGCTGTTCATCTCCGGACTGTTGCCACACAAAAAAGGTCTGCTTCCGGTGTAAAAGTCGCTCCTGCCCGCACTGCGGAGTGAAGGCTGGCGCACAGTGGATACAGTATCTGCTGAGTCTGGTTCCCGACTGTCCGTGGCAGCATATTGTGTTCACACTTCCCTGCCAGTACTGGTCCCTGGTGTTCCACAACCGGTGGTTACTGGCAGAGATGAGCCGCATTGCTGCGGATGTGATACAGGAAATCTGCCGCCAGGCAGATGTGGTGCCGGGGATATTCACGGTCATCCACACATGGGGACGTGACCAGCAGTGGCATCCGCACATTCACCTGTCGACAACGGCCGGCGGCGTGACACCAGACCACACCTGGAAAAACCTTCATTTTTACGCCCGTAAGGTGATGAGCATGTGGCGTTACAGGATAACGTGGTTACTGTCACGAAAATACCCGGAGCTGGTAATACCGGATGAACTGGCAGTGGAAGGAAACAGCAAACGGGACTGGAATTGCTTCCTGGACACGCATTACCGCCGCGGCTGGAATGTCAACATATCCAGGGTGATGGATAACGCCACACATGTGGCGGTGTACTTCGGCTCTTACCTGAAAAAGCCACCGGTGCCGATGAGTCGGCTGGAGCATTATGCCGGTCAGGATGAAATCGGTCTGCGTTACAACAGTCACCGTACAAAACGGGAAGAATACCTGTTGATGAGTGGAGATGAGTTCATGGAAAGGTTCTCCTGGCATGTAGCAGATAAGGGGTTCCGTATGGTGAGGTACTACGGTTTCCTGAGTCCGGTGAAGCGCCGCTTACTGGAAGAAGTTGTGTACGTCATAACGGAGACGGTGAGAAAAACGGCGATGCAAATCAGGTGGAGAGGGATGTATCAGAGGTTACTGAAGGTTGACCCGCTGAAGTGCATTCTGTGCGGAAGTCAGATGCGTTTTACGGGGCTGAAGCGGGGTTACCGACTGGCAGAGCTGGTCCTGATGCATGAGCGACTGGCACGACAGCAGGTGTGCGGCTGAGAGCCGCAGAGGGGAAGTTGCGTCCATTTTACCGGAAACGGAGCAAAAAACCGCCATTCATACCCTGTATCAATCAGTGTCATCCTGTTTAATAGTCGTTTCCGCTCATATGGTGCACAAGGGGTGTTGAAGAAATATCCGTTTTGTGGTGCTTTTTTAGTCTTTTGGGGATTTAAATTCCTATCGAT

>IS91-V88

CGAGTAGGCAGCCTGGCGGCTGCGGCTTGTCATGGCCTGGAATTACCGTTATAAAAAAAGATAATGTCATTGTCTTTCAGGTAGTTATATGGCCCGTTCAGCTAAACCCCGTAAACGCAAACCCGCACCACAAAGAAGCAAACTTCCCCGCTATGTTGTGAAACTTCATCCGGATGATTTTTTTGACGAAGAAGACGCTGAAGTTCTGCGCTTTGATAATTTTGACGATGCCGTTGAGTGCTGCGCTGACCTGGGTATTCCGTTCTTTCTGGATGCAGGAAACAAAAAGCTGGTCTTCTGGTTTGTTCGTGTCGATGACGAAGGGTATCCGGAAATAGCCCGCTGTACGGAGCGGGAGTTTGCGACCATTCTTGCCGGTATCAGCGCCGGCGGTATGTACTGTCCGGAATGCGGCACGGTTCACTGGCCGGACGGAGTCACCCCGCCCTTCTGATGCTTCCCCGTTTTGCCGATATTTTTCAGCAGGGAAACCGCTGGCTTAACTGGCTGGAGAAGCAGCCGGAAGGGTCTGTGCGTCCGGTGGTGACTGAGTCAGTGACAAAAATCATGGCATGCGGGACCACGCTGATGGGCTACACGCAATGGTGCTGTTCGTCACCGGACTGTTGCCACACCAAAAAGGTCTGCTTCCGGTGTAAAAGCCGCTCCTGTCCGCACTGCGGGGTGAAGGCTGGCGCACAGTGGATACAGTATCTGCTGAGCCTGGTCCCCGACTGCCCGTGGCAGCATATTGTGTTCACACTTCCCTGCCAGTACTGGTCCCTGGTGTTCCACAACCGGTGGTTACTGGCAGAGATGAGCCGCATTGCAGCTGATTTGATACTGGAAATCTGCCATCAGGCAGATGTGGAGCCGGGGATATTCACGGTGATCCACACATGGGGGCGTGACCAGCAGTGGCATCCGCATATCCATTTATCGACAACTGCCGGTGGTGTGACGTCGGGCCACACCTGGAAAAATCTTCATTTTTACGCCCGTAAGGTGATGAGCATGTGGCGTTACCGGATAACGCGGCTACTGTCCCGGAAATCCCCGGAGCTGGTGATACCGGCTGAGCTGGCAGCAGAGGGAAGCGGCAGACGGGAATGGAATCGCTTCCTGGCCACCCACTACCGGCGTGGCTGGAATGTCAACGTATCCCGGATGATGGATAACGCCACGCATGTGGCGGTGTACTTTGGCTCTTACCTGAAAAAGCCGCCGGTGCCAATGAGCCGTCTGGAACACTATGCCGGTCAGGATGAAATTGGTCTGCGTTACAACAGCCACCGCACAAAACGGGAAGAATACCTGGTGATGAGTGGCGATGAGTTCATGGAAAGGTTCTCGTGGCATGTGGCGGATAAGGGGTTCCGTATAGTGAGGTACTACGGTTTCCTGAGTCCGTCGAAACGGCGGTTACTGGAAGAGGTGGTGTACGTCATAACGGAGACGGTGAGAAAAACGGCGATGCAAATCAGGTGGAGAGGGATGTATCAGAGGTTACTGAAGGTTGACCCGCTGAAGTGCATTCTGTGCGGAAGTCAGATGCGTTTTACGGGGCTGAAGCGGGGTTACCGACTGGCAGAGCTGGTCCTGATGCATGAGCGACTGGCACGACAGCAGGTGTGCGGCTGAGAGTCGCAGAGGGGAAGTTGCGTCCATTTTACCGGAAACGGAGCAAAAAACCGCCATTCATACCCTGTATCAATCAGTGTCATCCTGTTTAATAGTCGTTTCCGCTCATATGGTGCACAAGAGGTGTTGAAGAAATATCCGTTTTGTGGTGCTTTTTTAGTCTTTTGGGGATTTAAATTCCTATCGAT

>IS91-V89

CGAGGCCCGTTCAGCTAAACCCCGTAAACGAAAACCTGCCCCTCAAAGAAACAAACTTCCCCGCTATGTCGTGAAGCTTCACGACGATGACTTCTTTGACGAAGAAGACGCAGAAGCTCTGCGCTTTGATAATTTTGACGATGCCGTTGAGTGCTGCGCAGACCTGAATATTCCCTTCTTTGTGGATGCCGGAAACAAAAAGCTGGTCTTCTGGTTTGTTCGTGTCGATGACGAAGGGTATCCTGAAATAGCCCGCTGCACGGAGCGGGAGTTTGCGACCATTCTTGCCGGTATCAGCGCCGGCGGCATGTACTGCCCGGAGTGTGGCACGGTTCACTGGCCGGACGGAGTCCCCCCGCCCTTCTGATGCTTCCCCGTTTTGCCGACATTTTTCAGCAGGGAAACCGCTGGCTTAACTGGCTGGAGAAACAACCGGAAGGTTCAGTGCGTCCGGTAGTCATTGAGTCGGTGACAAAAATCATGGCGTGCGGAACCACGCTGATGGGGTACACACAGTGGTGCTGTTCATCTCCGGACTGTTGCCACACAAAAAAGGTCTGCTTCCGGTGTAAAAGTCGCTCCTGCCCGCACTGCGGAGTGAAGGCTGGCGCACAGTGGATACAGTATCTGCTGAGTCTGGTTCCCGACTGTCCGTGGCAGCATATTGTGTTCACACTTCCCTGCCAGTACTGGTCCCTGGTGTTCCACAACCGGTGGTTACTGGCAGAGATGAGCCGCATTGCTGCGGATGTGATACAGGAAATCTGCCGCCAGGCAGATGTGGTGCCGGGGATATTCACGGTCATCCACACATGGGGACGTGACCAGCAGTGGCATCCGCACATTCACCTGTCGACAACGGCCGGCGGCGTGACACCAGACCACACCTGGAAAAACCTTCATTTTTACGCCCGTAAGGTGATGAGCATGTGGCGTTACAGGATAACGTGGTTACTGTCACGAAAATACCCGGAGCTGGTGATACCGGATGAACTGGCAGTGGAAGGAAACAGCAAACGGGACTGGAATTGCTTCCTGGACACGCATTACCGCCGCGGCTGGAATGTCAACATATCCAGGGTGATGGATAACGCCACACATGTGGCGGTGTACTTCGGCTCTTACCTGAAAAAGCCACCGGTGCCGATGAGTCGGCTGGAGCATTATGCCGGTCAGGATGAAATCGGTCTGCGTTACAACAGTCACCGTACAAAACGGGAAGAATACCTGTTGATGAGTGGAGATGAGTTCATGGAAAGGTTCTCCTGGCATGTAGCAGATAAGGGGTTCCGTATGGTGAGGTACTACGGTTTCCTGAGTCCGGTGAAGCGCCGCTTACTGGAAGAAGTTGTGTACGTCATAACGGAGACGGTGAGAAAAACGGCGATGCAAATCAGGTGGAGAGGGATGTATCAGAGGTTACTGAAGGTTGACCCGCTGAAGTGCATTCTGTGCGGAAGTCAGATGCGTTTTACGGGGCTGAAGCGGGGTTACCGACTGGCAGAGCTGGTCCTGATGCATGAGCGACTGGCACGACAGCAGGTGTGCGGCTGAGAGCCGCAGAGGGGAAGTTGCGTCCATTTTACCGGAAACGGAGCAAAAAACCGCCATTCATACCCTGTATCAATCAGTGTCATCCTGTTTAATAGTCGTTTCCGCTCATATGGTGCACAAGGGGTGTTGAAGAATATCCGTTTTGTGGTGCTTTTTTAGTCTTTTGGGGATTTAAATTCCTATCGAT

>IS91-V90

CGAGTAGGCAGCCTGGCGGCTGCGGCTTGTCATGGCCTGAAATTACCGTTATAAAAACAGACAATATCATTGTCTTTCAGGTAGTTATATGTCCCGTTCAGCTAAACCCCGTAAACGAAAAACCTGCCCCTCAAAGAAGCAAACTTCCCCGCTATGTCGTGAAGCTTCACGACGATGACTTCTTTGACGAAGAAGACGCAGAAGCTCTGCGCTTTGATAATTTTGACGATGCCGTTGAGTGCTGCGCAGACCTGAATATTCCCTTCTTTGTGGATGCCGGAAACAAAAAGCTGGTCTTCTGGTTTGTTCGTGTCGATGACGAAGGGTATCCTGAAATAGCCCGCTGCACGGAGCGGGAGTTTGCGACCATTCTTGCCGGTATCAGCGCCGGCGGCATGTACTGCCCGGAGTGTGGCACGGTTCACTGGCCGGACGGAGTCCCCCCGCCCTTCTGATGCTTCCCCGTTTTGCCGACATTTTTCAGCAGGGAAACCGCTGGCTTAACTGGCTGGAGAAACAACCGGAAGGTTCAGTGCGTCCGGTAGTCATTGAGTCGGTGACAAAAATCATGGCGTGCGGAACCACGCTGATGGGGTACACACAGTGGTGCTGTTCATCTCCGGACTGTTGCCACACAAAAAAGGTCTGCTTCCGGTGTAAAAGTCGCTCCTGCCCGCACTGCGGAGTGAAGGCTGGCGCACAGTGGATACAGTATCTGCTGAGTCTGGTTCCCGACTGTCCGTGGCAGCATATTGTGTTCACACTTCCCTGCCAGTACTGGTCCCTGGTGTTCCACAACCGGTGGTTACTGGCAGAGATGAGCCGCATTGCTGCGGATGTGATACAGGAAATCTGCCGCCAGGCAGATGTGGTGCCGGGGATATTCACGGTCATCCACACATGGGGACGTGACCAGCAGTGGCATCCGCACATTCACCTGTCGACAACGGCCGGCGGCGTGACACCAGACCACACCTGGAAAAACCTTCATTTTTACGCCCGTAAGGTGATGAGCATGTGGCGTTACAGGATAACGTGGTTACTGTCACGAAAATACCCGGAGCTGGTGATACCGGATGAACTGGCAGTGGAAGGAAACAGCAAACGGGACTGGAATTGCTTCCTGGACACGCATTACCGCCGCGGCTGGAATGTCAACATATCCAGGGTGATGGATAACGCCACACATGTGGCGGTGTACTTCGGCTCTTACCTGAAAAAGCCACCGGTGCCGATGAGTCGGCTGGAGCATTATGCCGGTCAGGATGAAATCGGTCTGCGTTACAACAGTCACCGTACAAAACGGGAAGAATACCTGTTGATGAGTGGAGATGAGTTCATGGAAAGGTTCTCCTGGCATGTAGCAGATAAGGGGTTCCGTATGGTGAGGTACTACGGTTTCCTGAGTCCGGTGAAGCGCCGCTTACTGGAAGAAGTTGTGTACGTCATAACGGAGACGGTGAGAAAAACGGCGATGCAAATCAGGTGGAGAGGGATGTATCAGAGGTTACTGAAGGTTGACCCGCTGAAGTGCATTCTGTGCGGAAGTCAGATGCGTTTTACGGGGCTGAAGCGGGGTTACCGACTGGCAGAGCTGGTCCTGATGCATGAGCGACTGGCACGACAGCAGGTGTGCGGCTGAGAGCCGCAGAGGGGAAGTTGCGTCCATTTTACCGGAAACGGAGCAAAAAACCGCCATTCATACCCTGTATCAATCAGTGTCATCCTGTTTAATAGTCGTTTCCGCTCATATGGTGCACAAGGGGTGTTGAAGAAACATCCGTTTTGTGGTGCTTTTTTAGTCTTTTGGGGATTTAAATTCCTATCGAT

>IS91-V91

CGAGTAGGCAGCCTGGCGGCTGCGGCTTGTCATGGTCTGAGATTACCGTTATAAAAACAGGCAATATCATTGTCTTTCAGGTGGTTATATGGCCCGTTCAGCTAAACCCCGTAAACGAAAACCTTCCCCACAACACAGCAAACTTCCCCGCTATGTCGTGACGCTTCACGACGATGGCTTCTTTGACGAAGAAGACGCAGAAGTTCTGCGCTTTGATAGTTTTGACGATGCCGTTGAATGCTGCGCAGACCTGAATATTCCGTTCTTTGTGGATGTGGGAAACAAAAAGCTGGTCTTCTGGTTTGTACGTGTTGATGACGAAGGGTATCCGGAAATAGCCCGCTGCACGGAGCGGGAGTTTGCGACCATTCTTGCCGGTATCAGCGCCGGCGGTATGTACTGTCCGGAATGCGGCACGGTTCACTGGCCGGACGGAGTCACCCCGCCCTTCTGATGCTTCCCCGTTTTGCCGATATTTTTCAGCAGGGAAACCGCTGGGTTAACTGGCTGGAGAAGCAGCCGGAAGGGTCTGTGCGTCCGGTGGTGATTGAGTCGGTGACAAAAATCATGGCGTGCGGGACCACGCTGATGGGGTACACACAGTGGTGTTGTTCCTCACCGGACTGTTGCCACACAAAAAAGGTCTGCTTCCGGTGTAAAAGTCGCTCCTGCCCGCACTGCGGGGTGAAGGCTGGCGCACAGTGGATACAGTATCTGCTGAGTCTGGCTCCCGACTGCCCGTGGCAGCATATTGTGTTCACACTTCCCTGCCAGTACTGGTCCCTGATATTCCACAACCGGTGGTTGCTGGCAGAGATGAGCCGTATCGCAGCGAATGTGATACTGGAAATCTGCCGTCAGGCAGATGTGGAGCCGGGGATATTCACGGTGATCCACACATGGGGGCGTGACCAGCAGTGGCATCCGCATATTCATTTATCGACAACTGCCGGTGGTGTGACGTCGGGCCACACCTGGAAAAATCTTCATTTTTACGCCCGTAAGGTGATGAGCATGTGGCGTTACCGGATAACGCGGCTACTGTCCCGGAAATACCCGGAGCTGGTAATACCGGATGAACTGGCAGTGGAAGGAAACAGCAAACGGGACTGGAATCGCTTCCTGGACACGCATTACCGCCGCGGCTGGAATGTCAACATATCCAGGGTGATGGATAACGCCACACATGTGGCGGTGTACTTCGGCTCTTACCTGAAAAAGCCACCGGTGCCGATGAGTCGGCTGGAACATTATGCCGGTCAGGATGAAATCGGTCTGCGTTACAACAGTCACCGTACAAAACGGGAAGAATACCTGTTGATGAGTGGAGATGAGTTCATGGAAAGGTTCTCCTGGCATGTAGCAGATAAGGGGTTCCGTATGGTGAGGTACTACGGTTTCCTGAGTCCGGTGAAGCGCCGCTTACTGGAAGAAGTTGTGTACGTCATAACGGAGACGGTGAGAAAAACGGCGATGCAAATCAGGTGGAGAGGGATGTATCAGAGGTTACTGAAGGTTGACCCGCTGAAGTGCATTCTGTGCGGAAGCCAGATGCGTTTTACGGGGCTGAAGCGGGGTTACCGACTGGCAGAGCTGGTCCTGATGCATGAGCGACTGGCACGACAGCAGGTGTGCGGCTGAGAGCCGCAGAGGGGAAGTTGCGTCCATTTTACCGGAAACGGAGCAAAAAACCGCCATTCATACCCTGTATCAATCAGTGTCATCCTGTTTAATAGTCGTTTCCGCTCATATGGTGCACAAGGGGTGTTGAAGAAACATCCGTTTTGTGGTGCTTTTTTAGTCTTTTGGGGATTTAAATTCCTATCGAT

>IS91-V92

CGAGTAGGCAGCCTGGCGGCTGCGGCTTGTCATGGTCTGAGATTACCGTTATAAAAACAGGCAATATCATTGTCTTTCAGGTGGTTATATGGCCCGTTCAGCTAACCCCCGTAAACGAAAACCTTCCCCACAACACAGCAAACTTCCCCGCTATGTCGTGACGCTTCACGACGATGACTTCTTTGACGAAGAAGACGCAGAAGTTCTGCGCTTTGATAGTTTTGACGATGCCGTTGAATGCTGCGCAGACCTGAATATTCCGTTCTTTGTGGATGTGGGAAACAAAAAGCTGGTCTTCTGGTTTGTACGTGTTGATGACGAAGGGTATCCGGAAATAGCCCGCTGCACGGAGCGGGAGTTTGCGACCATTCTTGCCGGTATCAGCGCCGGCGGTATGTACTGTCCGGAATGCGGCACGGTTCACTGGCCGGACGGAGTCACCCCGCCCTTCTGATGCTTCCCCGTTTTGCCGATATTTTTCAGCAGGGAAACCGCTGGGTTAACTGGCTGGAGAAGCAGCCGGAAGGGTCTGTGCGTCCGGTGGTGATTGAGTCGGTGACAAAAATCATGGCGTGCGGGACCACGCTGATGGGGTACACACAGTGGTGTTGTTCCTCACCGGACTGTTGCCACACAAAAAAGGTCTGCTTCCGGTGTAAAAGTCGCTCCTGCCCGCACTGCGGGGTGAAGGCTGGCGCACAGTGGATACAGTATCTGCTGAGTCTGGCTCCCGACTGCCCGTGGCAGCATATTGTGTTCACACTTCCCTGCCAGTACTGGTCCCTGATATTCCACAACCGGTGGTTGCTGGCAGAGATGAGCCGTATCGCAGCGAATGTGATACTGGAAATCTGCCGTCAGGCAGATGTGGAGCCGGGGATATTCACGGTGATCCACACATGGGGGCGTGACCAGCAGTGGCATCCGCATATTCATTTATCGACAACTGCCGGTGGTGTGACGTCGGGCCACACCTGGAAAAATCTTCATTTTTACGCCCGTAAGGTGATGAGCATGTGGCGTTACCGGATAACGCGGCTACTGTCCCGGAAATACCCGGAGCTGGTAATACCGGATGAACTGGCAGTGGAAGGAAACAGCAAACGGGACTGGAATCGCTTCCTGGACACGCATTACCGCCGCGGCTGGAATGTCAACATATCCAGGGTGATGGATAACGCCACACATGTGGCGGTGTACTTCGGCTCTTACCTGAAAAAGCCACCGGTGCCGATGAGTCGGCTGGAACATTATGCCGGTCAGGATGAAATCGGTCTGCGTTACAACAGTCACCGTACAAAACGGGAAGAATACCTGTTGATGAGTGGAGATGAGTTCATGGAAAGGTTCTCCTGGCATGTAGCAGATAAGGGGTTCCGTATGGTGAGGTACTACGGTTTCCTGAGTCCGGTGAAGCGCCGCTTACTGGAAGAAGTTGTGTACGTCATAACGGAGACGGTGAGAAAAACGGCGATGCAAATCAGGTGGAGAGGGATGTATCAGAGGTTACTGAAGGTTGACCCGCTGAAGTGCATTCTGTGCGGAAGCCAGATGCGTTTTACGGGGCTGAAGCGGGGTTACCGACTGGCAGAGCTGGTCCTGATGCATGAGCGACTGGCACGACAGCAGGTGTGCGGCTGAGAGCCGCAGAGGGGAAGTTGCGTCCATTTTACCGGAAACGGAGCAAAAAACCGCCATTCATACCCTGTATCAATCAGTGTCATCCTGTTTAATAGTCGTTTCCGCTCATATGGTGCACAAGGGGTGTTGAAGAAACATCCGTTTTGTGGTGCTTTTTTAGTCTTTTGGGGATTTAAATTCCTATCGAT

>IS91-V93

CGAGTAGGCAGCCTGGCGGCTGCGGCTTGTCATGGTCTGAGATTACCGTTATAAAAACAGGCAATATCATTGTCTTTCAGGTGGTTATATGGCCCGTTCAGCTAAACCCCGTAAACGAAAACCTTCCCCACAACACAGCAAACTTCCCCGCTATGTCGTGACGCTTCACGACGATGGCTTCTTTGACGAAGAAGACGCAGAAGTTCTGCGCTTTGATAGTTTTGACGATGCCGTTGAATGCTGCGCAGACCTGAATATTCCGTTCTTTGTGGATGTGGGAAACAAAAAGCTGGTCTTCTGGTTTGTACGTGTTGATGACGAAGGGTATCCGGAAATAGCCCGCTGCACGGAGCGGGAGTTTGCGACCATTCTTGCCGGTATCAGCGCCGGCGGTATGTACTGTCCGGAATGCGGCACGGTTCACTGGCCGGACGGAGTCACCCCGCCCTTCTGATGCTTCCCCGTTTTGCCGATATTTTTCAGCAGGGAAACCGCTGGCTTAACTGGCTGGAGAAGCAGCCGGAAGGGTCTGTGCGTCCGGTGGTGATTGAGTCGGTGACAAAAATCATGGCGTGCGGGACCACGCTGATGGGGTACACACAGTGGTGTTGTTCCTCACCGGACTGTTGCCACACAAAAAAGGTCTGCTTCCGGTGTAAAAGTCGCTCCTGCCCGCACTGCGGGGTGAAGGCTGGCGCACAGTGGATACAGTATCTGCTGAGTCTGGCTCCCGACTGCCCGTGGCAGCATATTGTGTTCACACTTCCCTGCCAGTACTGGTCCCTGATATTCCACAACCGGTGGTTGCTGGCAGAGATGAGCCGTATCGCAGCGAATGTGATACTGGAAATCTGCCGTCAGGCAGATGTGGAGCCGGGGATATTCACGGTGATCCACACATGGGGGCGTGACCAGCAGTGGCATCCGCATATTCATTTATCGACAACTGCCGGTGGTGTGACGTCGGGCCACACCTGGAAAAATCTTCATTTTTACGCCCGTAAGGTGATGCGCATGTGGCGTTACCGGATAACGCGGCTACTGTCCCGGAAATACCCGGAGCTGGTAATACCGGATGAACTGGCAGTGGAAGGAAACAGCAAACGGGACTGGAATCGCTTCCTGGACACGCATTACCGCCGCGGCTGGAATGTCAACATATCCAGGGTGATGGATAACGCCACACATGTGGCGGTGTACTTCGGCTCTTACCTGAAAAAGCCACCGGTGCCGATGAGTCGGCTGGAACATTATGCCGGTCAGGATGAAATCGGTCTGCGTTACAACAGTCACCGTACAAAACGGGAAGAATACCTGTTGATGAGTGGAGATGAGTTCATGGAAAGGTTCTCCTGGCATGTAGCAGATAAGGGGTTCCGTATGGTGAGGTACTACGGTTTCCTGAGTCCGGTGAAGCGCCGCTTACTGGAAGAAGTTGTGTACGTCATAACGGAGACGGTGAGAAAAACGGCGATGCAAATCAGGTGGAGAGGGATGTATCAGAGGTTACTGAAGGTTGACCCGCTGAAGTGCATTCTGTGCGGAAGCCAGATGCGTTTTACGGGGCTGAAGCGGGGTTACCGACTGGCAGAGCTGGTCCTGATGCATGAGCGACTGGCACGACAGCAGGTGTGCGGCTGAGAGCCGCAGAGGGGAAGTTGCGTCCATTTTACCGGAAACGGAGCAAAAAACCGCCATTCATACCCTGTATCAATCAGTGTCATCCTGTTTAATAGTCGTTTCCTCTCATATGGTGCACAAGGGGTGTTGAAGAAACATCCGTTTTGTGGTGCTTTTTTAGTCTTTTGGGGATTTAAATTCCTATCGAT

>IS91-V94

CGAGTAGGCAGCCTGGCGGCTGCGGCTTGTCATGGCCTGGAATTACCGTTATAAAAAAAGATAATGTCATTGTCTTTCAGGTAGTTATATGGCCCGTTCAGCTAAACCCCGTAAACGCAAACCCGCACCACAAAGAAGCAAACTTCCCCGCTATGTTGTGAAACTTCATCCGGATGATTTTTTTGACGAAGAAGACGCTGAAGTTCTGCGCTTTGATAATTTTGACGATGCCGTTGAGTGCTGCGCTGACCTGGGTATTCCGTTCTTTCTGGATGCAGGAAACAAAAAGCTGGTCTTCTGGTTTGTTCGTGTCGATGACGAAGGGTATCCGGAAATAGCCCGCTGTACGGAGCGGGAGTTTGCGACCATTCTTGCCGGTATCAGCGCCGGCGGTATGTACTGTCCGGAATGCGGCACGGTTCACTGGCCGGACGGAGTCACCCCGCCCTTCTGATGCTTCCCCGTTTTGCCGATATTTTTCAGCAGGGAAACCGCTGGCTTAACTGGCTGGAGAAGCAGCCGGAAGGGTCTGTGCGTCCGGTGGTGACTGAGTCAGTGACAAAAATCATGGCATGCGGGACCACGCTGATGGGCTACACGCAATGGTGCTGTTCGTCACCGGACTGTTGCCACACCAAAAAGGTCTGCTTCCGGTGTAAAAGCCGCTCCTGTCCGCACTGCGGGGTGAAGGCTGGCGCACAGTGGATACAGTATCTGCTGAGCCTGGTCCCCGACTGCCCGTGGCAGCATATTGTGTTCACACTTCCCTGCCAGTACTGGTCCCTGGTGTTCCACAACCGGTGGTTACTGGCAGAGATGAGCCGCATTGCAGCTGATGTGATACTGGAAATCTGCCATCAGGCAGATGTGGAGCCGGGGATATTCACGGTGATCCACACATGGGGGCGTGACCAGCAGTGGCATCCGCATATCCATTTATCGACAACTGCCGGTGGTGTGACGTCGGGCCACACCTGGAAAAATCTTCATTTTTACGCCCGTAAGGTGATGAGCATGTGGCGTTACCGGATAACGCGGCTACTGTCCCGGAAATCCCCGGAGCTGGTGATACCGGCTGAGCTGGCAGCAGAGGGAAGCGGCAGACGGGAATGGAATCGCTTCCTGGCCACCCACTACCGGCGTGGCTGGAATGTCAACGTATCCCGGGTGATGGATAACGCCACGCATGTGGCGGTGTACTTTGGCTCTTACCTGAAAAAGCCGCCGGTGCCAATGAGCCGTCTGGAACACTATGCCGGCCAGGATGAAATTGGTCTGCGTTACAACAGCCACCGCACAAAACGGGAAGAATACCTGGTGATGAGTGGCGATGAGTTCATGGAAAGGTTCTCGTGGCATGTGGCGGATAAGGGGTTCCGTATAGTGAGGTACTACGGTTTCCTGAGTCCGTCGAAACGGCGGTTACTGGAAGAGGTGGTGTACGTCATAACGGAGACGGTGAGAAAAACGGCGATGCAAATCAGGTGGAGAGGGATGTATCAGAGGTTACTGAAGGTTGACCCGCTGAAGTGCATTCTGTGCGGAAGTCAGATGCGTTTTACGGGGCTGAAGCGGGGTTACCGACTGGCAGAGCTGGTCCTGATGCATGAGCGACTGGCACGACAGCAGGTGTGCGGTTGAGAGTCGCAGAGGGGAAGTTGCGTCCATTTTACCGGAAACGGAGCAAAAAACCACCATTCATACCCTGTATCAATCAATGTCATCCTGTTTAATAGTCGTTTCCGCTCATATGGTGCACAAGGGGTGTTGAAGAAATATCCGTTTTGTGGTGCTTTTTTAGTCTTTCGGGGATTTAAATTCCTATCGAT

>IS91-V95

CGAGTAGGCAGCCTGGCGGCTGCGGCTTGTCATGGCCTGAAATTACCGTTATAAAAACAGACAATATCATTGTCTTTCAGGTAGTTATATGTCCCGTTCAGCTAAACCCCGTAAACGAAAACCTGCCCCTCAAAGAAGCAAACTTCCCCGCTATGTCGTGAAGCTTCACGACGATGACTTCTTTGACGAAGAAGACGCAGAAGCTCTGCGCTTTGATAATTTTGACGATGCCGTTGAGTGCTGCGCAGACCTGAATATTCCCTTCTTTGTGGATGCCGGAAACAAAAAGCTGGTCTTCTGGTTTGTTCGTGTCGATGACGAAGGGTATCCTGAAATAGCCCGCTGCACGGAGCGGGAGTTTGCGACCATTCTTGCCGGTATCAGCGCCGGCGGCATGTACTGCCCGGAGTGTGGCACGGTTCACTGGCCGGACGGAGTCCCCCCGCCCTTCTGATGCTTCCCCGTTTTGCCGACATTTTTCAGCAGGGAAACCGCTGGCTTAACTGGCTGGAGAAACAACCGGAAGGTTCAGTGCGTCCGGTAGTCATTGAGTCGGTGACAAAAATCATGGCGTGCGGAACCACGCTGATGGGGTACACACAGTGGTGCTGTTCATCTCCGGACTGTTGCCACACAAAAAAGGTCTGCTTCCGGTGTAAAAGTCGCTCCTGCCCGCACTGCGGAGTGAAGGCTGGCGCACAGTGGATACAGTATCTGCTGAGTCTGGTTCCCGACTGTCCGTGGCAGCATATTGTGTTCACACTTCCCTGCCAGTACTGGTCCCTGGTGTTCCACAACCGGTGGTTACTGGCAGAGATGAGCCGCATTGCTGCGGATGTGATACAGGAAATCTGCCGCCAGGCAGATGTGGTGCCGGGGATATTCACGGTCATCCACACATGGGGACGTGACCAGCAGTGGCATCCGCACATTCACCTGTCGACAACGGCCGGCGGCGTGACACCAGACCACACCTGGAAAAACCTTCATTTTTACGCCCGTAAGGTGATGAGCATGTGGCGTTACAGGATAACGTGGTTACTGTCACGAAAATACCCGGAGCTGGTAATACCGGATGAACTGGCAGTGGAAGGAAACAGCAAACGGGACTGGAATTGCTTCCTGGACACGCATTACCGCCGCGGCTGGAATGTCAACATATCCCGGGTGATGGATAACGCCACACATGTGGCGGTGTACTTCGGCTCTTACCTGAAAAAGCCACCGGTGCCGATGAGTCGGCTGGAGCATTATGCCGGTCAGGATGAAATCGGTCTGCGTTACAACAGTCACCGTACAAAACGGGAAGAATACCTGTTGATGAGTGGAGATGAGTTCATGGAAAGGTTCTCCTGGCATGTAGCAGATAAGGGGTTCCGTATGGTGAGGTACTACGGTTTCCTGAGTCCGGTGAAGCGCCGCTTACTGGAAGAAGTTGTGTACGTCATAACGGAGACGGTGAGAAAAACGGCGATGCAAATCAGGTGGAGAGGGATGTATCAGAGGTTACTGAAGGTTGACCCGCTGAAGTGCATTCTGTGCGGAAGTCAGATGCGTTTTACGGGGCTGAAGCGGGGTTACCGACTGGCAGAGCTGGTCCTGATGCATGAGCGACTGGCACGACAGCAGGTGTGCGGCTGAGAGCCGCAGAGGGGAAGTTGCGTCCATTTTACCGGAAACGGAGCAAAAAACCGCCATTCATACCCTGTATCAATCAGTGTCATCCTGTTTAATAGTCGTTTCCGCTCATATGGTGCACAAGGGGTGTTGAAGAAACATCCGTTTTGTGGTGCTTTTTTAGTCTTTTGGGGATTTAAATTCCTATCGAT

>IS91-V96

CGAGTAGGCAGCCTGGCGGCTGCGGCTTGTCATGGTCTGAGATTACCGTTATAAAAACAGGCAATATCATTGTCTTTCAGGTGGTTATATGGCCCGTTCAGCTAAACCCCGTAAACGAAAACCTTCCCCACAACACAGCAAACTTCCCCGCTATGTCGTGACGCTTCACGACGATGACTTCTTTGACGAAGAAGACGCAGAAGTTCTGCGCTTTGATAGTTTTGACGATGCCGTTGAATGCTGCGCAGACCTGAATATTCCGTTCTTTGTGGATGTGGGAAACAAAAAGCTGGTCTTCTGGTTTGTACGTGTTGATGACGAAGGGTATCCGGAAATAGCCCGCTGCACGGAGCGGGAGTTTGCGACCATTCTTGCCGGTATCAGCGCCGGCGGTATGTACTGTCCGGAATGCGGCACGGTTCACTGGCCGGACGGAGTCACCCCGCCCTTCTGATGCTTCCCCGTTTTGCCGATATTTTTCAGCAGGGAAACCGCTGGCTTAACTGGCTGGAGAAGCAGCCGGAAGGGTCTGTGCGTCCGGTGGTGATTGAGTCGGTGACAAAAATCATGGCGTGCGGGACCACGCTGATGGGGTACACACAGTGGTGTTGTTCCTCACCGGACTGTTGCCACACAAAAAAGGTCTGCTTCCGGTGTAAAAGTCGCTCCTGCCCGCACTGCGGGGTGAAGGCTGGCGCACAGTGGATACAGTATCTGCTGAGTCTGGCTCCCGACTGCCCGTGGCAGCATATTGTGTTCACACTTCCCTGCCAGTACTGGTCCCTGATATTCCACAACCGGTGGTTGCTGGCAGAGATGAGCCGTATCGCAGCGAATGTGATACTGGAAATCTGCCGTCAGGCAGATGTGGAGCCGGGGATATTCACGGTGATCCACACATGGGGGCGTGACCAGCAGTGGCATCCGCATATTCATTTATCGACAACTGCCGGTGGTGTGACGTCGGGCCACACCTGGAAAAATCTTCATTTTTACGCCCGTAAGGTGATGAGCATGTGGCGTTACCGGATAACGCGGCTACTGTCCCGGAAATACCCGGAGCTGGTAATACCGGATGAACTGGCAGTGGAAGGAAACAGCAAACGGGACTGGAATCGCTTCCTGGACACGCATTACCGCCGCGGCTGGAATGTCAACATATCCAGGGTGATGGATAACGCCACACATGTGGCGGTGTACTTCGGCTCTTACCTGAAAAAGCCACCGGTGCCGATGAGTCGGCTGGAACATTATGCCGGTCAGGATGAAATCGGTCTGCGTTACAACAGTCACCGTACAAAACGGGAAGAATACCTGTTGATGAGTGGAGATGAGTTCATGGAAAGGTTCTCCTGGCATGTAGCAGATAAGGGGTTCCGTATGGTGAGGTACTACGGTTTCCTGAGTCCGGTGAAGCGCCGCTTACTGGAAGAAGTTGTGTACGTCATAACGGAGACGGTGAGAAAAACGGCGATGCAAATCAGGTGGAGAGGGATGTATCAGAGGTTACTGAAGGTTGACCCGCTGAAGTGCATTCTGTGCGGAAGCCAGATGCGTTTTACGGGGCTGAAGCGGGGTTACCGACTGGCAGAGCTGGTCCTGATGCATGAGCGACTGGCACGACAGCAGGTGTGCGGCTGAGAGCCGCAGAGGGGAAGTTGCGTCCATTTTGAGGGAAACGGAGCAAAAAACCGCCATTCATACCCTGTATCAATCAGTGTCATCCTGTTTAATAGTCGTTTCCGCTCATATGGTGCACAAGGGGTGTTGAAGAAACATCCGTTTTGTGGTGCTTTTTTAGTCTTTTGGGGATTTAAATTCCTATCGAT

>IS91-V97

CGAGTAGGCAGCCTGGCGGCTGCGGCTTGTCATGGTCTGAGATTACCGTTATAAAAACAGGCAATATCATTGTCTTTCAGGTGGTTATATGGCCCGTTCAGCTAAACCCCGTAAACGAAAACCTTCCCCACAACACAGCAAACTTCCCCGCTATGTCGTGACGCTTCACGACGATGACTTCTTTGACGAAGAAGACGCAGAAGTTCTGCGCTTTGATAGTTTTGACGATGCCGTTGAATGCTGCGCAGACCTGAATATTCCGTTCTTTGTGGATGTGGGAAACAAAAAGCTGGTCTTCTGGTTTGTACGTGTTGATGACGAAGGGTATCCGGAAATAGCCCGCTGCACGGAGCGGGAGTTTGCGACCATTCTTGCCGGTATCAGCGCCGGCGGTATGTACTGTCCGGAATGCGGCACGGTTCACTGGCCGGACGGAGTCACCCCGCCCTTCTGATGCTTCCCCGTTTTGCCGATATTTTTCAGCAGGGAAACCGCTGGGTTAACTGGCTGGAGAAGCAGCCGGAAGGGTCTGTGCGTCCGGTGGTGATTGAGTCGGTGACAAAAATCATGGCGTGCGGGACCACGCTGATGGGGTACACACAGTGGTGTTGTTCCTCACCGGACTGTTGCCACACAAAAAAGGTCTGCTTCCGGTGTAAAAGTCGCTCCTGCCCGCACTGCGGGGTGAAGGCTGGCGCACAGTGGATACAGTATCTGCTGAGTCTGGCTCCCGACTGCCCGTGGCAGCATATTGTGTTCACACTTCCCTGCCAGTACTGGTCCCTGATATTCCACAACCGGTGGTTGCTGGCAGAGATGAGCCGTATCGCAGCGAATGTGATACTGGAAATCTGCCGTCAGGCAGATGTGGAGCCGGGGATATTCACGGTGATCCACACATGGGGGCGTGACCAGCAGTGGCATCCGCATATTCATTTATCGACAACTGCCGGTGGTGTGACGTCGGGCCACACCTGGAAAAATCTTCATTTTTACGCCCGTAAGGTGATGAGCATGTGGCGTTACCGGATAACGCGGCTACTGTCCCGGAAATACCCGGAGCTGGTAATACCGGATGAACTGGCAGTGGAAGGAAACAGCAAACGGGACTGGAATCGCTTCCTGGACACGCATTACCGCCGCGGCTGGAATGTCAACATATCCAGGGTGATGGATAACGCCACACATGTGGCGGTGTACTTCGGCTCTTACCTGAAAAAGCCACCGGTGCCGATGAGTCGGCTGGAACATTATGCCGGTCAGGATGAAATCGGTCTGCGTTACAACAGTCACCGTACAAAACGGGAAGAATACCTGTTGATGAGTGGAGATGAGTTCATGGAAAGGTTCTCCTGGCATGTAGCAGATAAGGGGTTCCGTATGGTGAGGTACTACGGTTTCCTGAGTCCGGTGAAGCGCCGCTTACTGGAAGAAGTTGTGTACGTCATAACGGAGACGGTGAGAAAAACGGCGATGCAAATCAGGTGGAGAGGGATGTATCAGAGGTTACTGAAGGTTGACCCGCTGAAGTGCATTCTGTGCGGAAGCCAGATGCGTTTTACGGGGCTGAAGCGGGGTTACCGACTGGCAGAGCTGGTCCTGATGCATGAGCGACTGGCACGACAGCAGGTGTGCGGCTGAGAGCCGCAGAGGGGAAGTTGCGTCCATTTTGAGGGAAACGGAGCAAAAAACCGCCATTCATACCCTGTATCAATCAGTGTCATCCTGTTTAATAGTCGTTTCCGCTCATATGGTGCACAAGGGGTGTTGAAGAAACATCCGTTTTGTGGTGCTTTTTTAGTCTTTTGGGGATTTAAATTCCTATCGAT

>IS91-V98

CGAGTAGGCAGCCTGGCGGCTGCGGCTTGTCATGGTCTGAGATTACCGTTATAAAAACAGGTAATATCATTGTCTTTCAGGTGGTTATATGGCCCGTTCAGCTAAACCCCGTAAACGAAAACCCGCACCACAAAGAAGCAAACTTCCCCGCTATGTTGTGAAACTTCATCCGGATGATTTTTTTGACGAAGAAGACGCTGAAGTTCTGCGCTTTGATAATTTTGACGATGCCGTTGAGTGCTGCGCTGACCTGGGTATTCCGTTCTTTCTGGATGCAGGAAACAAAAAGCTGGTCTTCTGGTTTGTTCGTGTCGATGACGAAGGGTATCCGGAAATAGCCCGCTGTACGGAGCGGGAGTTTGCAACCATTCTTGCCGGTATCAGTGCCGGTGGTATGTACTGCCCGGAATGCGGCACAGTTCACTGGCCGGATGGCGTTACCCCACCCGTCTGATGCTTCCCCGTTTTGCCGATATTTTTCAGCAGGGTAACCGCTGGCTTAACTGGCTGGAGAAACAGCCGGAAGGTTCAGTGCGTCCGGTGGTGATTGAGTCAGTGACAAAAATCATGGCATGCGGGACCACGCTGATGGGCTACACGCAATGGTGCTGTTCGTCACCGGACTGTTGCCACACCAAAAAGGTCTGCTTCCGGTGTAAAAGCCGCTCCTGTCCGCACTGCGGGGTGAAGGCTGGCGCACAGTGGATACAGTATCTGCTGAGCCTGGTCCCCGACTGCCCGTGGCAGCATATTGTGTTCACACTTCCCTGCCAGTACTGGTCCCTGGTGTTCCACAACCGGTGGTTACTGGCAGAGATGAGCCGCATTGCAGCGGATGTGATACTGGAAATCTGCCATCAGACAGATGTGGAGCCGGGGATATTCACGGTGATCCACACATGGGGGCGTGACCAGCAGTGGCATCCGCATATCCATTTATCGACAACTGCCGGTGGTGTGACGTCGGGCCACACCTGGAAAAATCTTCATTTTTACGCCCGTAAGGTGATGAGCATGTGGCGTTACCGGATAACGCGGCTACTGTCCCAGAAATACCCGGAGCTGGTAATACCGGATGAACTGGCAGTGGAAGGAAACAGCAAACGGGACTGGAATCGCTTCCTGGACACGCATTACCGCCGCGGCTGGAATGTCAACATATCCAGGGTGATGGATAACGCCACACATGTGGCGGTGTACTTCGGCTCTTACCTGAAAAAGCCACCGGTGCCGGTGAGTCGGCTGGAGCATTATGCCGGTCAGGATGAAATCGGTCTGCGTTACAACAGTCACCGTACAAAACGGGAAGAATATCTGTTGATGAGTGGAGATGAGTTCATGGAAAGGTTCTCCTGGCATGTAGCAGATAAGGGGTTCCGTATGGTGAGGTACTACGGTTTCCTGAGTCCGTCGAAACGGCGGTTACTGGAAGAGGTGGTGTACGTCATAACGGAGACGGTGAGAAAAACGGCGATGCAAATCAGGTGGAGAGGGATGTATCAGCGGTTACTGAAGGTTGACCCGCTGAAATGCATTCTGTGCGGAAGTCAGATGCGGTTTACGGGGCTGAAGCGGGGATACCGTCTGGCAGAGCTGGTTATGATGCATGAGCCGCTGGCCCGAATGCAGTGTTGCGGCTGAGAGCCGCAGAGGGGAAGTTGCGTCCATTTTACGGGGAGCGGAGCAAAAAACCACCACTCATACCCTTTATCAATCCGTGTTATCCTGTTTAATAGTCGTTTCCGTTCATATGGTGCATAAGGAGTGTTGAAGAAATATCCGTTTTGTGATGTTTTTTAATCTTTTGGGGGTTTTAATTCCTATTGAT

>IS91-V99

CGAGTAGGCAGCCTGGCGGCTGCGGCTTGTCATGGTCTGAGATTACCGTTATAAAAACAGGCAATATCATTGTCTTTCAGGTGGTTATATGGCCCGTTCAGCTAAACCCCGTAAACGAAAACCTTCCCCACAACACAGCAAACTTCCCCGCTATGTCGTGACGCTTCACGACGATGGCTTCTTTGACGAAGAAGACGCAGAAGTTCTGCGCTTTGATAGTTTTGACGATGCCGTTGAATGCTGCGCAGACCTGAATATTCCGTTCTTTGTGGATGTGGGAAACAAAAAGCTGGTCTTCTGGTTTGTACGTGTTGATGACGAAGGGTATCCGGAAATAGCCCGCTGCACGGAGCGGGAGTTTGCGACCATTCTTGCCGGTATCAGCGCCGGCGGTATGTACTGTCCGGAATGCGGCACGGTTCACTGGCCGGACGGAGTCACCCCGCCCTTCTGATGCTTCCCCGTTTTGCCGATATTTTTCAGCAGGGAAACCGCTGGGTTAACTGGCTGGAGAAGCAGCCGGAAGGGTCTGTGCGTCCGGTGGTGATTGAGTCGGTGACAAAAATCATGGCGTGCGGGACCACGCTGATGGGGTACACACAGTGGTGTTGTTCCTCACCGGACTGTTGCCACACAAAAAAGGTCTGCTTCCGGTGTAAAAATCGCTCCTGCCCGCACTGCGGGGTGAAGGCTGGCGCACAGTGGATACAGTATCTGCTGAGTCTGGCTCCCGACTGCCCGTGGCAGCATATTGTGTTCACACTTCCCTGCCAGTACTGGTCCCTGATATTCCACAACCGGTGGTTGCTGGCAGAGATGAGCCGTATCGCAGCGAATGTGATACTGGAAATCTGCCGTCAGGCAGATGTGGAGCCGGGGATATTCACGGTGATCCACACATGGGGGCGTGACCAGCAGTGGCATCCGCATATTCATTTATCGACAACTGCCGGTGGTGTGACGTCGGGCCACACTTGGAAAAATCTTCATTTTTACGCCCGTAAGGTGATGAGCATGTGGCGTTACCGGATAACGCGGCTACTGTCCCGGAAATACCCGGAGCTGGTAATACCGGATGAACTGGCAGTGGAAGGAAACAGCAAACGGGACTGGAATCGCTTCCTGGACACGCATTACCGCCACGGCTGGAATGTCAACATATCCAGGGTGATGGATAACGCCACACATGTGGCGGTGTACTTCGGCTCTTACCTGAAAAAGCCACCGGTGCCGATGAGTCGGCTGGAACATTATGCCGGTCAGGATGAAATCGGTCTGCGTTACAACAGTCACCGTACAAAACGGGAAGAATACCTGTTGATGAGTGGAGATGAGTTCATGGAAAGGTTCTCCTGGCATGTAGCAGATAAGGGGTTCCGTATGGTGAGGTACTACGGTTTCCTGAGTCCGGTGAAGCGCCGCTTACTGGAAGAAGTTGTGTACGTCATAACGGAGACGGTGAGAAAAACGGCGATGCAAATCAGGTGGAGAGGGATGTATCAGAGGTTACTGAAGGTTGACTCGCTGAAGTGCATTCTGTGCGGAAGCCAGATGCGTTTTACGGGGCTGAAGCGGGGTTACCGACTGGCAGAGCTGGTCCTGATGCATGAGCGACTGGCACGACAGCAGGTGTGCGGCTGAGAGCCGCAGAGGGGAAGTTGCGTCCATTTTACCGGAAACGGAGCAAAAAACCGCCATTCATACCCTGTATCAATCAGTGTCATCCTGTTTAATAGTCGTTTCCGCTCATATGGTGCACAAGGGGTGTTGAAGAAACATCCGTTTTGTGGTGCTTTTTTAGTCTTTTGGGGATTTAAATTCCTATCGAT

>IS91-V100

CGAGTAGGCAGCCTGGCGGCTGCGGCTTGTCATGGTCTGGAATTACCGTTATAAAAAAAGATAATGTCATTGTCTTTCAGGTAGTTATATGGCCCGTTCAGCTAAACCCCGTAAACGAAAACCTGCCCCTCAAAGAAGCAAACTTCCCCGCTATGTCGTGAAGCTTCACGACGATGACTTCTTTGACGAAGAAGACGCAGAAGCTCTGCGCTTTGATAATTTTGACGATGCCGTTGAGTGCTGCGCAGACCTGAATATTCCCTTCTTTGTGGATGCCGGAAACAAAAAGCTGGTCTTCTGGTTTGTTCGTGTCGATGACGAAGGGTATCCTGAAATAGCCCGCTGCACGGAGCGGGAGTTTGCGACCATTCTTGCCGGTATCAGCGCCGGCGGCATGTACTGCCCGGAGTGTGGCACGGTTCACTGGCCGGACGGAGTCCCCCCGCCCTTCTGATGCTTCCCCGTTTTGCCGACATTTTTCAGCAGGGAAACCGCTGGCTTAACTGGCTGGAGAAACAACCGGAAGGTTCAGTGCGTCCGGTAGTCATTGAGTCGGTGACAAAAATCATGGCGTGCGGAACCACGCTGATGGGCTACACACAGTGGTGCTGTTCATCTCCGGACTGTTGCCACACAAAAAAGGTCTGCTTCCGGTGTAAAAGTCGCTCCTGCCCGCACTGCGGAGTGAAGGCTGGCGCACAGTGGATACAGTATCTGCTGAGTCTGGTTCCCGACTGTCCGTGGCAGCATATTGTGTTCACACTTCCCTGCCAGTACTGGTCCCTGGTGTTCCACAACCGGTGGTTACTGGCAGAGATGAGCCGCATTGCTGCGGATGTGATACAGGAAATCTGCCGCCAGGCAGATGTGGTGCCGGGGATATTCACGGTCATCCACACATGGGGACGTGACCAGCAGTGGCATCCGCACATTCACCTGTCGACAACGGCCGGCGGCGTGACACCAGACCACACCTGGAAAAACCTTCATTTTTACGCCCGTAAGGTGATGAGCATGTGGCGTTACAGGATAACGTGGTTACTGTCACGAAAATACCCGGAGCTGGTGATACCGGATGCGCTGGCAGTTGAAGGAAGCAGCAGACGGGACTGGAATTGCTTCCTGGACACGCATTACCGCCGCGGCTGGAATGTCAACGTATCCCGGGTGATGGATAACGCCACACATGTGGCGGTGTACTTCGGCTCTTACCTGAAAAAACCGCCGGTGCCGATGAGCCGTCTGGAGCATTATGCCGGTCAGGATGAAATCGGTCTGCGTTACAACAGTCACCGTACAAAACGGGAAGAATACCTGTTGATGAGTGGAGATGAGTTCATGGAAAGGTTCTCCTGGCATGTAGCAGATAAGGGGTTCCGTATGGTGAGGTACTACGGTTTCCTGAGTCCGGTGAAGCGCCGCTTACTGGAAGAAGTTGTGTACGTCATAACGGAGACGGTGAGAAAAACGGCGATGCAAATCAGGTGGAGAGGGATGTATCAGAGGTTACTGAAGGTTGACCCGCTGAAGTGCATTCTGTGCGGAAGTCAGATGCGTTTTACGGGGCTGAAGCGGGGTTACCGACTGGCAGAGCTGGTCCTGATGCATGAGCGACTGGCACGACAGCAGGTGTGCGGCTGAGAGCCGCAGAGGGGAAGTTGCGTCCATTTTACCGGAAACGGAGCAAAAAACCGCCATTCATACCCTGTATCAATCAGTGTCATCCTGTTTAATAGTCGTTTCCGCTCATATGGTGCACAAGGGGTGTTGAAGAAATATCCGTTTTGTGGTGCTTTTTTAGTCTTTTGGGGATTTAAATTCCTATCGAT

>IS91-V101

CGAGTAGGCAGCCTGGCGGCTGCGGCTTGTCATGGTCTGAGATTACCGTTATAAAAACAGGCAATATCATTGTCTTTCAGGTGGTTATATGGCCCGTTCAGCTAAACCCCGTAAACGAAAACCTTCCCCACAACGAAGCAAACTTCCCCGCTATGTCGTGACGCTTCACGACGATGACTTCTTTGACGAAGAAGACGCAGAAGTTCTGCGCTTTGATAGTTTTGACGATGCCGTTGAATGCTGCGCAGACCTGAATATTCCGTTCTTTGTGGATGTGGGAAACAAAAAGCTGGTCTTCTGGTTTGTACGTGTTGATGACGAAGGGTATCCGGAAATAGCCCGCTGCACGGAGCGGGAGTTTGCGACCATTCTTGCCGGTATCAGCGCCGGCGGTATGTACTGTCCGGAATGCGGCACGGTTCACTGGCCGGACGGAGTCACCCCGCCCTTCTGATGCTTCCCCGTTTTGCCGATATTTTTCAGCAGGGAAACCGCTGGCTTAACTGGCTGGAGAAGCAGCCGGAAGGGTCTGTGCGTCCGGTGGTGATTGAGTCGGTGACAAAAATCATGGCGTGCGGGACCACGCTGATGGGTTACACACAGTGGTGTTGTTCCTCACCGGACTGTTGCCACACAAAAAAGGTCTGCTTCCGGTGTAAAAGTCGCTCCTGCCCGCACTGCGGGGTGAAGGCTGGCACACAGTGGATACAGTATCTGCTGAGTCTGGTTCCCGACTGCCCGTGGCAGCATATTGTGTTCACACTTCCCTGCCAGTACTGGTCCCTGATATTCCACAACCGGTGGTTGCTGGCAGAGATGAGCCGTATCGCAGCGAATGTGATACTGGAAATCTGCCGTCAGGCGGACGTGGAGCCGGGGATATTCACGGTAATCCACACATGGGGGCGTGACCAGCAGTGGCATCCGCATATCCATTTATCGACAACTGCCGGTGGTGTGACGTCGGGCCACACCTGGAAAAATCTTCATTTTTACGCGCGTAAGGTGATGAGCATGTGGCGTTACCGGATAACGCGGCTACTGTCCCGGAAATACCCGGAGCTGGTAATACCGGATGAACTGGCAGTGGAAGGAAACAGCAAACGGGACTGGAATCGCTTCCTGGACACGCATTACCGCCGCGGCTGGAATGTCAACATATCCAGGGTGATGGATAACGCCACACATGTGGCGGTGTACTTCGGCTCTTACCTGAAAAAGCCACCGGTGCCGATGAGTCGGCTGGAGCATTATGCCGGTCAGGATGAAATCGGTCTGCGTTACAACAGTCACCGTACAAAACGGGAAGAATACCTGTTGATGAGTGGAGATGAGTTCATGGAAAGGTTCTCCTGGCATGTAGCAGATAAGGGGTTCCGTATGGTGAGGTACTACGGTTTCCTGAGTCCGGTGAAGCGCCGCTTACTGGAAGAAGTTGTGTACGTCATAACGGAGACGGTGAGAAAAACGGCGATGCAAATCAGGTGGAGAGGGATGTATCAGAGGTTACTGAAGGTTGACCCGCTGAAGTGCATTCTGTGCGGAAGTCAGATGCGTTTTACGGGGCTGAAGCGGGGTTACCGACTGGCAGAGCTGGTCCTGATGCATGAGCGACTGGCACGACAGCAGGTGTGCGGCTGAGAGCCGCAGAGGGGAAGTTGCGTCCATTTTACCGGAAACGGAGCAAAAAACCGCCATTCATACCCTGTATCAATCAGTGTCATCCTGTTTAATAGTCGTTTCCGTTCATATGGTGCATAAGGAGTGTTGAAGAAATATCCGTTTTGTGGTGTTTTTTAATCTTTTTGGGGTTTTAATTCCTATTGAT

>IS91-V102

CGAGTAGGCAGCCTGGCGGCTGCGGCTTGTCATGGTCTGAGATTACCGTTATAAAAAAAGATAATGTCATTGTCTTTCAGGTAGTTATATGGCCCGTTCAGCTAAACCCCGTAAACGCAAACCCGCACCACAAAGAAGCAAACTTCCCCGCTATGTTGTGAAACTTCATCCGGATGATTTTTTTGACGAAGAAGACGCTGAAGTTCTGCGCTTTGATAATTTTGACGATGCCGTTGAGTGCTGCGCTGACCTGGGTATTCCGTTCTTTCTGGATGCAGGAAACAAAAAGCTGGTCTTCTGGTTTGTTCGTGTCGATGACGAAGGGTATCCGGAAATAGCCCGCTGTACGGAGCGGGAGTTTGCAACCATTCTTGCCGGTATCAGTGCCGGTGGTATGTACTGCCCGGAATGCGGCACAGTTCACTGGCCGGATGGCGTTACCCCACCCGTCTGATGCTTCCCCGTTTTGCCGATATTTTTCAGCAGGGTAACCGCTGGCTTAACTGGCTGGAGAAACAGCCGGAAGGTTCAGTGCGTCCGGTGGTGACTGAGTCAGTGACAAAAATCATGGCATGCGGGACCACGCTGATGGGCTACACGCAATGGTACTGTTCATCACCGGACTGTTGCCACACAAAAAAGGTCTGCTTCCGGTGTAAAAGTCGCTCCTGCCCGCACTGCGGGGTGAAGGCTGGCGCACAGTGGATACAGTATCTGCTGAGTCTGGTCCCCGACTGCCCGTGGCAGCATATTGTGTTCACACTTCCCTGCCAGTACTGGCCCCTGATATTCCACAACAGGTGGTTACTGGCAGAGATGAGCCGCATTGCTGCGAATGTGATACTGGAAATCTGCCGTCAGGCGGACGTGGAGCCGGGGATATTCACGGTAATCCACACATGGGGGCGTGACCAGCAGTGGCATCCGCATATCCATTTATCGACAACTGCCGGTGGTGTGACGTCGGGCCACACCTGGAAAAATATTCATTTTTACGCCCGTAAGGTGATGAGCATGTGGCGTTACCGGATAACGCGGCTACTGTCCCGGAAATACCCGGAGCTGGTAATACCGGATGAACTGGCAGTGGAAGGAAACAGCAAACGGGACTGGAATCGCTTTCTGGACACGCATTACCGCCGCGGCTGGAATGTCAACATATCCAGGGTGATGGATAACGCCACACATGTGGCGGTGTACTTCGGCTCTTACCTGAAAAAGCCACCGGTGCCGATGAGTCGGCTGGAGCATTATGCCGGTCAGGATGAAATCGGTCTGCGTTACAACAGTCACCGTACAAAACGGGAAGAATACCTGTTGATGAGTGGAGATGAGTTCATGGAAAGGTTCTCCTGGCATGTAGCAGATAAGGGGTTCCGTATGGTGAGGTACTACGGTTTCCTGAGTCCGGTGAAGCGCCGCTTACTGGAAGAAGTTGTGTACGTCATAACGGAGACGGTGAGAAAAACGGCGATGCAAATCAGGTGGAGAGGGATGTATCAGCGGTTACTGAAGGTTGACCCGCTGAAATGCGTCCTGTACGGAAGTCAGATGCGGTTTACGGGGCTGAAGCGGGGATACCGTCTGGCAGAGCTGGTTATGATGCATGAGCCGCTGGCCCGAATGCAGTGTTGCGGCTGAGAGCCGCAGAGGGGAAGTTGCGTCCATTTTACGGGGAACGGAGCAAAAAACCACCATTCATACCCTTTATCAATCAGTGTTATCCTGTTTAATAGTCGTTTCCGTTCATATGGTGCATAAGGAGTGTTGAAGAAATATCCGTTTTGTGGTGTTTTTTAATCTTTTGGGGGTTTTAATTCCTATTGAT

>IS91-V103

CGAGTAGGCAGCCTGGCGGCTGCGGCTTGTCATGGCCTAGAATTACCGTTATAAAAAAAGATAATGTCATTGTCTTTCAGGTAGTTATATGGCCCGTTCAGCTAAACCCCGTAAACGCAAACCCGCACCACAAAGAAGCAAACTTCCCCGCTATGTTGTGAAACTTCATCCGGATGATTTTTTTGACGAAGAAGACGCTGAAGTTCTGCGCTTTGATAATTTTGACGATGCCGTTGAGTGCTGCGCTGACCTGGGTATTCCGTTCTTTCTGGATGCAGGAAACAAAAAGCTGGTCTTCTGGTTTGTTCGTGTCGATGACGAAGGGTATCCGGAAATAGCCCGCTGTACGGAGCGGGAGTTTGCAACCATTCTTGCCGGTATCAGTGCCGGTGGTATGTACTGCCCGGAATGCGGCACAGTTCACTGGCCGGATGGCGTTACCCCACCCGTCTGATGCTTCCCCGTTTTGCCGATATTTTTCAGCAGGGTAACCGCTGGCTTAACTGGCTGGAGAAACAGCCGGAAGGTTCAGTGCGTCCGGTGGTGACTGAGTCAGTGACAAAAATCATGGCATGCGGGACCACGCTGATGGGCTACACGCAATGGTACTGTTCATCACCGGACTGTTGCCACACAAAAAAGGTCTGCTTCCGGTGTAAAAGTCGCTCCTGCCCGCACTGCGGGGTGAAGGCTGGCGCACAGTGGATACAGTATCTGCTGAGTCTGGTCCCCGACTGCCCGTGGCAGCATATTGTGTTCACACTTCCCTGCCAGTACTGGCCCCTGATATTCCACAACAGGTGGTTACTGGCAGAGATGAGCCGCATTGCTGCGAATGTGATACTGGAAATCTGCCGTCAGGCGGACGTGGAGCCGGGGATATTCACGGTAATCCACACATGGGGGCGTGACCAGCAGTGGCATCCGCATATCCATTTATCGACAACTGCCGGTGGTGTGACGTCGGGCCACACCTGGAAAAATCTTCATTTTTACGCCCGTAAGGTGATGAGCATGTGGCGTTACCGGATAACGCGGCTACTGTCCCGGAAATACCCGGAGCTGGTAATACCGGATGAACTGGCAGTGGAAGGAAACAGCAAACGGGACTGGAATCGCTTTCTGGACACGCATTACCGCCGCGGCTGGAATGTCAACATATCCAGGGTGATGGATAACGCCACACATGTGGCGGTGTACTTCGGCTCTTACCTGAAAAAGCCACCGGTGCCGATGAGTCGGCTGGAGCATTATGCCGGTCAGGATGAAATCGGTCTGCGTTACAACAGTCACCGTACAAAACGGGAAGAATACCTGTTGATGAGTGGAGATGAGTTCATGGAAAGGTTCTCCTGGCATGTAGCAGATAAGGGGTTCCGTATGGTGAGGTACTACGGTTTCCTGAGTCCGGTGAAGCGCCGCTTACTGGAAGAGGTGGTGTACGTCATAACGGAGACGGTGAGAAAAACGGCGATGCAAATCAGGTGGAGAGGGATGTATCAGCGGTTACTGAAGGTTGACCCGCTGAAATGCGTCCTGTACGGAAGTCAGATGCGGTTTACGGGGCTGAAGCGGGGATACCGTCTGGCAGAGCTGGTTATGATGCATGAGCCGCTGGCCCGAATGCAGTGTTGCGGCTGAGAGCCGCAGAGGGGAAGTTGCGTCCATTTTACGGGGAACGGAGCAAAAAACCACCATTCATACCCTTTATCAATCAGTGTTATCCTGTTTAATAGTCGTTTCCGTTCATATGGTGCATAAGGAGTGTTGAAGAAATATCCGTTTTGTGGTGTTTTTTAATCTTTTGGGGGTTTTAATTCCTATTGAT

>IS91-V104

CGAGTAGGCAGCCTGGCGGCTGCGGCTTGTCATGGTCTGGAATTACCGTTATAAAAAAGATAATATCATTGTCTTTCAGGTAGTTATATGGCCCGTTCAGCTAAACCCCGTAAACGCAAACCCGCACCACAAAGAAGCAAACTTCCCCGCTATGTTGTGAAACTTCATCCGGATGATTTTTTTGACGAAGAAGAAGCTGAAATTCTGCGCTTTGATAATTTTGACGATGCTGTTGAGTGCTGCGCTGACCTGGGTATTCCGTTCTTTCTGGATGCGGGAAACAAAAAGCTGGTCTTCTGGTTTGTTCGTGTTGATGACGAAGGGTATCCGGAAATAGCCCGCTGTACGGAGCGGGAGTTTGCGACCATTCTTGCCGGTATCAGCGCCGGCGGTATGTACTGCCAGGAATGCGGCACTGTTCACTGGCCGGATGGCGTTACCCCGCCCTTCTGATGCTTCCCCGTTTTGCAGACATCTTTCAGCAGGGAAACCGCTGGCTTAACTGGCTGGAGAAGCAGCCGGAAGGGACAGTGCGTCCGGTGGTGACTGAGTCAGTGACAAAAATCATGGCCTGCGGGACCACCATGATGGGGTACACACAGTGGTGCTGTTCATCACCGGACTGTTGCCACACCAAAAAGGTCTGCTTCCGGTGTAAAAGCCGCTCCAGCCCGCACTGCGGAGTGAAGGCCGGCGCACAGTGGATACAGTATCTGCTGAGTCTGGTCCCCGACTGCCCGTGGCAGCATATTGTGTTCACGCTTCCGTGCCAGTACTGGTCCCTGGTGTTCCACAACAGGTGGTTACTGGCAGAGATGAGCCGCATTGCTGCGGATGTGATACTGGAAATCTGCCGCCAGGCAGATGTGGAGCCGGGGATATTCACGGTAATCCACACATGGGGACGAGACCAGCAGTGGCATCCGCACATTCACCTGTCAACAACAGCCGGCGGAGTGACGTCAGGTCACACCTGGAAAAACCTTCATTTTTATGCCCGTAAGGTGATGAGTATGTGGCGTTACCGGATAACGCGGTTACTGTCACGAAAATACCCTGACCTGGTGATACCGGATGAGCTGGCCGCGGAAGGAAACAGCAAACGGGAATGGAATCGCTTCCTGGACACGCATTACCGCCGCGGCTGGAATGTCAACATATCCAGGGTGATGGATAACGCCACACATGTGGCGGTGTACTTCGGCTCTTACCTGAAAAAGCCACCGGTGCCGATGAGTCGGCTGGAGCATTATGCCGGTCAGGATGAAATCGGTCTGCGTTACAACAGTCACCGTACAAAACGGGAAGAATACCTGTTGATGAGTGGAGATGAGTTCATGGAAAGGTTCTCCTGGCATGTAGCAGATAAGGGGTTCCGTATGGTGAGGTACTACGGTTTCCTGAGTCCGGTGAAGCGCCGCTTACTGGAAGAAGTTGTGTACGTCATAACGGAGACGGTGAGAAAAACGGCGATGCAAATCAGGTGGAGAGGGATGTATCAGCGGTTACTGAAGGTTGACCCGCTGAAGTGCATTCTGTGCGGAAGTCAGATGCGTTTTACGGGGCTGAAGCGGGGTTACCGACTGGCAGAGCTGGTCCTGATGCATGAGCGACTGGCACGACAGCAGGTGTGCGGCTGAGAGCCGCAGAGGGGAAGTTGCGTCCATTTTACCGGGAACGGAGCAAAAAACCGCCATTCATACCCTGTATAAATCAGTGTCATCCTGTTTAATAGTCGTTTCCGTTCATATGGTGCACAAGGGGTGTTGAAGAAACATCCGTTTTGTGGTGCTTTTTTAGTCTTTTGGGGATTTAAATTCCTATCGAT

>IS91-V105

CGAGTAGGCAGCCTGGCGGCTGCGGCTTGTCATGGTCTGGAATTACCGTTATAAAAAAGATAATATCATTGTCTTTCAGGTAGTTATATGGCCCGTTCAGCTAAACCCCGTAAACGCAAACCCGCACCACAAAGAAGCAAACTTCCCCGCTATGTTGTGAAACTTCATCCGGATGATTTTTTTGACGAAGAAGAAGCTGAAATTCTGCGCTTTGATAATTTTGACGATGCTGTTGAGTGCTGCGCTGACCTGGGTATTCCGTTCTTTCTGGATGCGGGAAACAAAAAGCTGGTCTTCTGGTTTGTTCGTGTTGATGACGAAGGGTATCCGGAAATAGCCCGCTGTACGGAGCGGGAGTTTGCGACCATTCTTGCCGGTATCAGCGCCGGCGGAATGTACTGCCAGGAATGCGGCACTGTTCACTGGCCGGATGGCGTTACCCCGCCCTTCTGATGCTTCCCCGTTTTGCAGACATCTTTCAGCAGGGAAACCGCTGGCTTAACTGGCTGGAGAAGCAGCCGGAAGGGACAGTGCGTCCGGTGGTGACTGAGTCAGTGACAAAAATCATGGCCTGCGGGACCACCATGATGGGGTACACACAGTGGTGCTGTTCATCACCGGACTGTTGCCACACCAAAAAGGTCTGCTTCCGGTGTAAAAGCCGCTCCAGCCCGCACTGCGGAGTGAAGGCCGGCGCACAGTGGATACAGTATCTGCTGAGTCTGGTCCCCGACTGCCCGTGGCAGCATATTGTGTTCACGCTTCCGTGCCAGTACTGGTCCCTGGTGTTCCACAACAGGTGGTTACTGGCAGAGATGAGCCGCATTGCTGCGGATGTGATACTGGAAATCTGCCGCCAGGCAGATGTGGAGCCGGGGATATTCACGGTAATCCACACATGGGGACGAGACCAGCAGTGGCATCCGCACATTCACCTGTCAACAACAGCCGGCGGAGTGACGTCAGGTCACACCTGGAAAAACCTTCATTTTTATGCCCGTAAGGTGATGAGTATGTGGCGTTACCGGATAACGCGGTTACTGTCACGAAAATACCCTGACCTGGTGATACCGGATGAGCTGGCCGCGGAAGGAAACAGCAAACGGGAATGGAATCGCTTCCTGGACACGCATTACCGCCGCGGCTGGAATGTCAACATATCCAGGGTGATGGATAACGCCACACATGTGGCGGTGTACTTCGGCTCTTACCTGAAAAAGCCACCGGTGCCGATGAGTCGGCTGGAGCATTATGCCGGTCAGGATGAAATCGGTCTGCGTTACAACAGTCACCGTACAAAACGGGAAGAATACCTGTTGATGAGTGGAGATGAGTTCATGGAAAGGTTCTCCTGGCATGTAGCAGATAAGGGGTTCCGTATGGTGAGGTACTACGGTTTCCTGAGTCCGGTGAAGCGCCGCTTACTGGAAGAAGTTGTGTACGTCATAACGGAGACGGTGAGAAAAACGGCGATGCAAATCAGGTGGAGAGGGATGTATCAGCGGTTACTGAAGGTTGACCCGCTGAAGTGCATTCTGTGCGGAAGTCAGATGCGTTTTACGGGGCTGAAGCGGGGTTACCGACTGGCAGAGCTGGTCCTGATGCATGAGCGACTGGCACGACAGCAGGTGTGCGGCTGAGAGCCGCAGAGGGGAAGTTGCGTCCATTTTACCGGGAACGGAGCAAAAAACCGCCATTCATACCCTGTATAAATCAGTGTCATCCTGTTTAATAGTCGTTTCCGTTCATATGGTGCACAAGGGGTGTTGAAGAAACATCCGTTTTGTGGTGCTTTTTTAGTCTTTTGGGGATTTAAATTCCTATCGAT

>IS91-V106

CGAGTAGGCAGCCTGGCGGCTGCGGCTTGTCATGGTCTGGAATTACCGTTATAAAAAAGATAATATCATTGTCTTTCAGGTAGTTATATGGCCCGTTCAGCTAAACCCCGTAAACGCAAACCCGCACCACAAAGAAGCAAACTTCCCCGCTATGTTGTGAAACTTCATCCGGATGATTTTTTTGACGAAGAAGAAGCTGAAATTCTGCGCTTTGATAATTTTGACGATGCTGTTGAGTGCTGCGCTGACCTGGGTATTCCGTTCTTTCTGGATGCGGGAAACAAAAAGCTGGTCTTCTGGTTTGTTCGTGTTGATGACGAAGGGTATCCGGAAATAGCCCGCTGTACGGAGCGGGAGTTTGCGACCATTCTTGCCGGTATCAGCGCCGGCGGTATGTACTGCCAGGAATGCGGCACTGTTCACTGGCCGGATGGCGTTACCCCGCCCTTCTGATGCTTCCCCGTTTTGCAGACATCTTTCAGCAGGGAAACCGCTGGCTTAACTGGCTGGAGAAGCAGCCGGAAGGGACAGTGCGTCCGGTGGTGACTGAGTCAGTGACAAAAATCATGGCCTGCGGGACCACCATGATGGGGTACACACAGTGGTGCTGTTCATCACCGGACTGTTGCCACACCAAAAAGGTCTGCTTCCGGTGTAAAAGCCGCTCCAGCCCGCACTGCGGAGTGAAGGCCGGCGCACAGTGGATACAGTATCTGCTGAGTCTGGTCCCCGACTGCCCGTGGCAGCATATTGTGTTCACGCTTCCGTGCCAGTACTGGTCCCTGGTGTTCCACAACAGGTGGTTACTGGCAGAGATGAGCCGCATTGCTGCGGATGTGATACTGGAAATCTGCCGCCAGGCAGATGTGGAGCCGGGGATATTCACGGTAATCCACACATGGGGACGAGACCAGCAGTGGCATCCGCACATTCACCTGTCAACAACAGCCGGCGGAGTGACGTCAGGTCACACCTGGAAAAACCTTCATTTTTATGCCCGTAAGGTGATGAGTATGTGGCGTTACCGGATAACGCGGTTACTGTCACGAAAATACCCTGACCTGGTGATACCGGATGAGCTGGCCGCGGAAGGAAACAGCAAACGGGAATGGAATCGCTTCCTGGACACGCATTACCGCCGCGGCTGGAATGTCAACATATCCAGGGTGATGGATAACGCCACACATGTGGCGGTGTACTTCGGCTCTTACCTGAAAAAGCCACCGGTGCCGATGAGTCGGCTGGAGCATTATGCCGGTCAGGATGAAATCGGTCTGCGTTACAACAGTCACCGTACAAAACGGGAAGAATACCTGTTGATGAGTGGAGATGAGTTCATGGAAAGGTTCTCCTGGCATGTAGCAGATAAGGGGTTCCGTATGGTGAGGTACTACGGTTTCCTGAGTCCGGTGAAGCGCCGCTTACTGGAAGAAGTTGTGCACGTCATAACGGAGACGGTGAGAAAAACGGCGATGCAAATCAGGTGGAGAGGGATGTATCAGCGGTTACTGAAGGTTGACCCGCTGAAGTGCATTCTGTGCGGAAGTCAGATGCGTTTTACGGGGCTGAAGCGGGGTTACCGACTGGCAGAGCTGGTCCTGATGCATGAGCGACTGGCACGACAGCAGGTGTGCGGCTGAGAGCCGCAGAGGGGAAGTTGCGTCCATTTTACCGGGAACGGAGCAAAAAACCGCCATTCATACCCTGTATAAATCAGTGTCATCCTGTTTAATAGTCGTTTCCGTTCATATGGTGCACAAGGGGTGTTGAAGAAACATCCGTTTTGTGGTGCTTTTTTAGTCTTTTGGGGATTTAAATTCCTATCGAT

>IS91-V107

CGAGTAGGCAGCCTGGCGGCTGCGGCTTGTCATGGTCTGGAATTACCGTTATAAAAAAGATAATATCATTGTCTTTCAGGTAGTTATATGGCCCGTTCAGCTAAACCCCGTAAACGCAAACCCGCACCACAAAGAAGCAAACTTCCCCGCTATGTTGTGAAACTTCATCCGGATGATTTTTTTGACGAAGAAGAAGCTGAAATTCTGCGCTTTGATAATTTTGACGATGCTGTTGAGTGCTGCGCTGACCTGGGTATTCCGTTCTTTCTGGATGCGGGAAACAAAAAGCTGGTCTTCTGGTTTGTTCGTGTTGATGACGAAGGGTATCCGGAAATAGCCCGCTGTACGGAGCGGGAGTTTGCGACCATTCTTGCCGGTATCAGCGCCGGCGGTATGTACTGCCAGGAATGCGGCACTGTTCACTGGCCGGATGGCGTTACCCCGCCCTTCTGATGCTTCCCCGTTTTGCAGACATCTTTCAGCAGGGAAACCGCTGGCTTAACTGGCTGGAGAAGCAGCCGGAAGGGACAGTGCGTCCGGTGGTGACTGAGTCAGTGACAAAAATCATGGCCTGCGGGACCACCATGATGGGGTACACACAGTGGTGCTGTTCATCACCGGACTGTTGCCACACCAAAAAGGTCTGCTTCCGGTGTAAAAGCCGCTCCAGCCCGCACTGCGGAGTGAAGGCCGGCGCACAGTGGATACAGTATCTGCTGAGTCTGGTCCCCGACTGCCCGTGGCAGCATATTGTGTTCACGCTTCCGTGCCAGTACTGGTCCCTGGTGTTCCACAACAGGTGGTTACTGGCAGAGATGAGCCGCATTGCTGCGGATGTGATACTGGAAATCTGCCGCCAGGCAGATGTGGAGCCGGGGATATTCACGGTAATCCACACATGGGGACGAGACCAGCAGTGGCATCCGCACATTCACCTGTCAACAACAGCCGGCGGAGTGACGTCAGGTCACACCTGGAAAAACCTTCATTTTTATGCCCGTAAGGTGATGAGTATGTGGCGTTACCGGATAACGCGGTTACTGTCACGAAAATACCCTGACCTGGTGATACCGGATGAGCTGGCCGCGGAAGGAAACAGCAAACGGGAATGGAATCGCTTCCTGGACACGCATTACCGCCGCGGCTGGAATGTCAACATATCCAGGGTGATGGATAACGCCACACATGTGGCGGTGTACTTCGGCTCTTACCTGAAAAAGCCACCGGTGCCGATGAGTCGGCTGGAGCATTATGCCGGTCAGGATGAAATCGGTCTGCGTTACAACAGTCACCGTACAAAACGGGAAGAATACCTGTTGATGAGAGGAGATGAGTTCATGGAAAGGTTCTCCTGGCATGTAGCAGATAAGGGGTTCCGTATGGTGAGGTACTACGGTTTCCTGAGTCCGGTGAAGCGCCGCTTACTGGAAGAAGTTGTGTACGTCATAACGGAGACGGTGAGAAAAACGGCGATGCAAATCAGGTGGAGAGGGATGTATCAGCGGTTACTGAAGGTTGACCCGCTGAAGTGCATTCTGTGCGGAAGTCAGATGCGTTTTACGGGGCTGAAGCGGGGTTACCGACTGGCAGAGCTGGTCCTGATGCATGAGCGACTGGCACGACAGCAGGTGTGCGGCTGAGAGCCGCAGAGGGGAAGTTGCGTCCATTTTACCGGGAACGGAGCAAAAAACCGCCATTCATACCCTGTATAAATCAGTGTCATCCTGTTTAATAGTCGTTTCCGTTCATATGGTGCACAAGGGGTGTTGAAGAAACATCCGTTTTGTGGTGCTTTTTTAGTCTTTTGGGGATTTAAATTCCTATCGAT

>IS91-V108

CGAGTAGGCAGCCTGGCGGCTGCGGCTTGTCATGGCCTGAAATTACCGTTATAAAAACAGACAATATCATTGTCTTTCAGGTAGTTATATGTCCCGTTCAGCTAAACCCCGTAAACGAAAACCTGCCCCTCAAAGAAGCAAACTTCCCCGCTATGTCGTGAAGCTTCACGACGATGACTTCTTTGACGAAGAAGACGCAGAAGCTCTGCGCTTTGATAATTTTGACGATGCCGTTGAGTGCTGCGCAGACCTGAATATTCCCTTCTTTGTGGATGCCGGAAACAAAAAGCTGGTCTTCTGGTTTGTTCGTGTCGATGACGAAGGGTATCCTGAAATAGCCCGCTGCACGGAGCGGGAGTTTGCGACCATTCTTGCCGGTATCAGCGCCGGCGGCATGTACTGCCCGGAGTGTGGCACGGTTCACTGGCCGGACGGAGTCCCCCCGCCCTTCTGATGCTTCCCCGTTTTGCCGACATTTTTCAGCAGGGAAACCGCTGGCTTAACTGGCTGGAGAAACAGCCGGAAGGTTCAGTGCGTCCGGTGGTGACTGAGTCAGTGACAAAAATCATGGCATGCGGGACCACGCTGATGGGCTACACGCAATGGTGCTGTTCGTCACCGGACTGTTGCCACACCAAAAAGGTCTGCTTCCGGTGTAAAAGCCGCTCCTGTCCGCACTGCGGGGTGAAGGCTGGCGCACAGTGGATACAGTATCTGCTGAGCCTGGTCCCCGACTGCCCGTGGCAGCATATTGTGTTCACACTTCCCTGCCAGTACTGGTCCCTGGTGTTCCACAACCGGTGGTTACTGGCAGAGATGAGCCGCATTGCAGCGGATGTGATACTGGAAATCTGCCATCAGACAGATGTGGAGCCGGGGATATTCACGGTGATCCACACATGGGGGCGTGACCAGCAGTGGCATCCGCATATCCATTTATCGACAACTGCCGGTGGTGTGACGTCGGGCCACACCTGGAAAAATCTTCATTTTTACGCCCGTAAGGTGATGAGCATGTGGCGTTACCGGATAACGCGGCTACTGTCCCGGAAATACCCGGAGCTGGTGATACCGGATGAACTGGCAGTGGAAGGAAACAGCAAACGGGACTGGAATTGCTTCCTGGACACGCATTACCGCCGCGGCTGGAATGTCAACATATCCAGGGTGATGGATAACGCCACACATGTGGCGGTGTACTTCGGCTCTTACCTGAAAAAGCCACCGGTGCCGATGAGTCGGCTGGAGCATTATGCCGGTCAGGATGAAATCGGTCTGCGTTACAACAGTCACCGTACAAAACGGGAAGAATACCTGTTGATGAGTGGAGATGAGTTCATGGAAAGGTTCTCCTGGCATGTAGCAGATAAGGGGTTCCGTATGGTGAGGTACTACGGTTTCCTGAGTCCGGTGAAGCGCCGGTTACTGGAAGATGTTGTGTACGTCATAACGGAGACGGTGAGAAAGACGGCGATGCAAATCAGGTGGAGAGGGATGTATCAGCGGTTACTGAAGGTTGACCCGCTAAAGTGCATCCTGTGCGGATGTCAGATGCGTTTTACGGGGCTGAAGCGGGGCTACCGTCTGACAGAGCTGGTCCTGATGCATGAGCCACTGGCGCAACAGCGGGTGTGCGGCTGAGAGCCGCATCGGGGAAGTTGCGTCCATTTTCAGGGGAATGGAGTAAAAAAACCATCAGTGATATACAGTATCACTCGATAAGATCCATTTAATTGATGGCGGTGCACTCATGGCACGCAGGCAGTGTTGAATAAACATCCGTTTTTGGGTGTTTTTTAATCTTTTTGGGATTTAAATTCCTATCGAT

>IS91-V109

CGAGTAGGCAGCCTGGCGGCTGCGGCTTGTCATGGCCTGGAATTACCGTTATAAAAAAAGATAATGTCATTGTCTTTCAGGTAGTTATATGGCCCGTTCAGCTAAACCCCGTAAACGCAAACCCGCACCACAAAGAAGCAAACTTCCCCGCTATGTTGTGAAACTTCATCCGGATGATTTTTTTGACGAAGAAGACGCTGAAGTTCTGCGCTTTGATAATTTTGACGATGCCGTTGAGTGCTGCGCTGACCTGGGTATTCCGTTCTTTCTGGATGCAGGAAACAAAAAGCTGGTCTTCTGGTTTGTTCGTGTCGATGACGAAGGGTATCCGGAAATAGCCCGCTGTACGGAGCGGGAGTTTGCGACCATTCTTGCCGGTATCAGCGCCGGCGGTATGTACTGTCCGGAATGCGGCACGGTTCACTGGCCGGACGGAGTCACCCCGCCCTTCTGATGCTTCCCCGTTTTGCCGATATTTTTCAGCAGGGAAACCGCTGGCTTAACTGGCTGGAGAAGCAGCCGGAAGGGTCTGTGCGTCCGGTGGTGACTGAGTCAGTGACAAAAATCATGGCATGCGGGACCACGCTGATGGGCTACACGCAATGGTGCTGTTCGTCACCGGACTGTTGCCACACCAAAAAGGTCTGCTTCCGGTGTAAAAGCCGCTCCTGTCCGCACTGCGGGGTGAAGGCTGGCGCACAGTGGATACAGTATCTGCTGAGTCTGGTTCCCGACTGCCCGTGGCAGCATATTGTGTTCACACTTCCCTGCCAGTACTGGTCCCTGATATTCCACAACCGGTGGTTGCTGGCAGAGATGAGCCGTATCGCAGCGGATGTGATACTGGAAATCTGCCGTCAGGCGGACGTGGAGCCGGGGATATTCACGGTAATCCACACATGGGGACGAGACCAGCAGTGGCATCCGCATATCCATTTATCGACAACTGCCGGTGGTGTGACGTCGGGCCACACCTGGAAAAATCTTCATTTTTACGCCCGTAAGGTGATGAGCATGTGGCGTTACCGGATAACGCGGCTACTGTCCCGGAAATCCCCGGAGCTGGTGATACCGGCTGAGCTGGCAGCAGAGGGAAGCGGCAGACGGGAATGGAATCGCTTCCTGGCCACCCACTACCGGCGTGGCTGGAATGTCAACGTATCCCGGGTGATGGATAACGCCACGCATGTGGCGGTGTACTTTGGCTCTTACCTGAAAAAGCCGCCGGTGCCAATGAGCCGTCTGGAACACTATGCCGGTCAGGATGAAATTGGTCTGCGTTACAACAGCCACCGCACAAAACGGGAAGAATACCTGGTGATGAGTGGCGATGAGTTCATGGAAAGGTTCTCGTGGCATGTGGCGGATAAGGGGTTCCGTATAGTGAGGTACTACGGTTTCCTGAGTCCGTCGAAACGGCGGTTACTGGAAGAGGTGGTGTACGTCATAACGGAGACGGTGAGAAAAACGGCGATGCAAATCAGGTGGAGAGGGATGTATCAGAGGTTACTGAAGGTTGACCCGCTGAAGTGCATTCTGTGCGGAAGTCAGATGCGTTTTACGGGGCTGAAGCGGGGTTACCGACTGGCAGAGCTGGTCCTGATGCATGAGCGACTGGCACGACAGCAGGTGTGCGGCTGAGAGTCGCAGAGGGGAATTTGCGTCCATTTTACCGGAAACGGAGCAAAAAACCGCCATTCATACCCTGTATCAATCAGTGTCATCCTGTTTAATAGTCGTTTCCGCTCATATGGTGCACAAGAGGTGTTGAAGAAATATCCGTTTTGTGGTGCTTTTTTAGTCTTTTGGGGATTTAAATTCCTATCGAT

>IS91-V110

CGAGTAGGCAGCCTGGCGGCTGCGGCTTGTCATGGCCTGGAATTACCGTTATAAAAAAAGATAATGTCATTGTCTTTCAGGTAGTTATATGGCCCGTTCAGCTAAACCCCGTAAACGCAAACCCGCACCACAAAGAAGCAAACTTCCCCGCTATGTTGTGAAACTTCATCCGGATGATTTTTTTGACGAAGAAGACGCTGAAGTTCTGCGCTTTGATAATTTTGACGATGCCGTTGAGTGCTGCGCTGACCTGGGTATTCCGTTCTTTCTGGATGCAGGAAACAAAAAGCTGGTCTTCTGGTTTGTTCGTGTCGATGACGAAGGGTATCCGGAAATAGCCCGCTGTACGGAGCGGGAGTTTGCGACCATTCTTGCCGGTATCAGCGCCGGCGGTATGTACTGTCCGGAATGCGGCACGGTTCACTGGCCGGACGGAGTCACCCCGCCCTTCTGATGCTTCCCCGTTTTGCCGATATTTTTCAGCAGGGAAACCGCTGGCTTAACTGGCTGGAGAAGCAGCCGGAAGGGTCTGTGCGTCCGGTGGTGACTGAGTCAGTGACAAAAATCATGGCATGCGGGACCACGCTGATGGGCTACACGCAATGGTGCTGTTCGTCACCGGACTGTTGCCACACCAAAAAGGTCTGCTTCCGGTGTAAAAGCCGCTCCTGTCCGCACTGCGGGGTGAAGGCTGGCGCACAGTGGATACAGTATCTGCTGAGTCTGGTTCCCGACTGCCCGTGGCAGCATATTGTGTTCACACTTCCCTGCCAGTACTGGTCCCTGATATTCCACAACCGGTGGTTGCTGGCAGAGATGAGCCGTATCGCAGCGGATGTGATACTGGAAATCTGCCGTCAGGCGGACGTGGAGCCGGGGATATTCACGGTAATCCACACATGGGGACGAGACCAGCAGTGGCATCCGCATATCCATTTATCGACAACTGCCGGTGGTGTGACGTCGGGCCACACCTGGAAAAATCTTCATTTTTACGCCCGTAAGGTGATGAGCATGTGGCGTTACCGGATAACGCGGCTACTGTCCCGGAAATCCCCGGAGCTGGTGATACCGGCTGAGCTGGCAGCAGAGGGAGGCGGCAGACGGGAATGGAATCGCTTCCTGGCCACCCACTACCGGCGTGGCTGGAATGTCAACGTATCCCGGGTGATGGATAACGCCACGCATGTGGCTGTGTACTTTGGCTCTTACCTGAAAAAGCCGCCGGTGCCAATGAGCCGTCTGGAACACTATGCCGGTCAGGATGAAATTGGTCTGCGTTACAACAGCCACCGCACAAAACGGGAAGAATACCTGGTGATGAGTGGCGATGAGTTCATGGAAAGGTTCTCGTGGCATGTGGCGGATAAGGGGTTCCGTATAGTGAGGTACTACGGTTTCCTGAGTCCGTCGAAACGGCGGTTACTGGAAGAGGTGGTGTACGTCATAACGGAGACGGTGAGAAAAACGGCGATGCAAATCAGGTGGAGAGGGATGTATCAGAGGTTACTGAAGGTTGACCCGCTGAAGTGCATTCTGTGCGGAAGTCAGATGCGTTTTACGGGGCTGAAGCGGGGTTACCGACTGGCAGAGCTGGTCCTGATGCATGAGCGACTGGCACGACAGCAGGTGTGCGGCTGAGAGTCGCAGAGGGGAATTTGCGTCCATTTTACCGGAAACGGAGCAAAAAACCGCCATTCATACCCTGTATCAATCAGTGTCATCCTGTTTAATAGTCGTTTCCGCTCATATGGTGCACAAGAGGTGTTGAAGAAACATCCGTTTTGTGGTGCTTTTTTAGTCTTTTGGGGATTTAAATTCCTATCGAT

>IS91-V111

CGAGTAGGCAGCCTGGCGGCTGCGGCTTGTCATGGCCTGAAATTACCGTTATAAAAACAGACAATATCATTGTCTTTCAGGTAGTTATATGTCCCGTTCAGCTAAACCCCGTAAACGAAAACCTGCCCCTCAAAGAAGCAAACTTCCCCGCTATGTCGTGAAGCTTCACGACGATGACTTCTTTGACGAAGAAGACGCAGAAGCTCTGCGCTTTGATAATTTTGACGATGCCGTTGAGTGCTGCGCAGACCTGAATATTCCCTTCTTTGTGGATGCCGGAAACAAAAAGCTGGTCTTCTGGTTTGTTCGTGTCGATGACGAAGGGTATCCTGAAATAGCCCGCTGCACGGAGCGGGAGTTTGCGACCATTCTTGCCGGTATCAGCGCCGGCGGCATGTACTGCCCGGAGTGTGGCACGGTTCACTGGCCGGACGGAGTCCCCCCGCCTTCTGATGCTTCCCCGTTTTGCCGACATTTTTCAGCAGGGTAACCGCTGGCTTAACTGGCTGGAGAAACAGCCGGAAGGTTCAGTGCGTCCGGTGGTGACTGAGTCAGTGACAAAAATCATGGCATGCGGGACCACGCTGATGGGCTACACGCAATGGTGCTGTTCGTCACCGGACTGTTGCCACACCAAAAAGGTCTGCTTCCGGTGTAAAAGCCGCTCCTGTCCGCACTGCGGGGTGAAGGCTGGCGCACAGTGGATACAGTATCTGCTGAGCCTGGTCCCCGACTGCCCGTGGCAGCATATTGTGTTCACACTTCCCTGCCAGTACTGGTCCCTGGTGTTCCACAACCGGTGGTTACTGGCAGAGATGAGCCGCATTGCAGCGGATGTGATACTGGAAATCTGCCATCAGACAGATGTGGAGCCGGGGATATTCACGGTGATCCACACATGGGGGCGTGACCAGCAGTGGCATCCGCATATCCATTTATCGACAACTGCCGGTGGTGTGACGTCGGGCCACACCTGGAAAAATCTTCATTTTTACGCCCGTAAGGTGATGAGCATGTGGCGTTACCGGATAACGCGGCTACTGTCCCGGAAATACCCGGAGCTGGTGATACCGGATGAACTGGCAGTGGAAGGAAACAGCAAACGGGACTGGAATTGCTTCCTGGACACGCATTACCGCCGCGGCTGGAATGTCAACATATCCAGGGTGATGGATAACGCCACACATGTGGCGGTGTACTTCGGCTCTTACCTGAAAAAGCCACCGGTGCCGATGAGTCGGCTGGAGCATTATGCCGGTCAGGATGAAATCGGTCTGCGTTACAACAGTCACCGTACAAAACGGGAAGAATACCTGTTGATGAGTGGAGATGAGTTCATGGAAAGGTTCTCCTGGCATGTAGCAGATAAGGGGTTCCGTATGGTGAGGTACTACGGTTTCCTGAGTCCGGTGAAGCGCCGGTTACTGGAAGATGTTGTGTACGTCATAACGGAGACGGTGAGAAAGACGGCGATGCAAATCAGGTGGAGAGGGATGTATCAGCGGTTACTGAAGGTTGACCCGCTAAAGTGCATCCTGTGCGGATGTCAGATGCGTTTTACGGGGCTGAAGCGGGGCTACCGTCTGACAGAGCTGGTCCTGATGCATGAGCCACTGGCGCAACAGCGGGTGTGCGGCTGAGAGCCGCATCGGGGAATTGCGTCCATTTTCAGGGGAATGGAGTAAAAAACCATCAGTGATATACAGTATCACTCGATAAGATCCATTTAATTGATGGCGGTGCACTCATGGCACGCAGGCAGTGTCGAATAAACATCCGTTTTTGGGTGTTTTTTAATCTTTTTGGGATTTAAATTCCTATCGAT

>IS91-V112

CGAGTAGGCAGCCTGGCGGCTGCGGCTTGTCATGGCCTGGAATTACCGTTATAAAAAAAGATAATGTTATTGTCTTTCAGGTAGTTATATGGCCCGTTCAGCTAAACCCCGTAAACGCAAACCCGCACCACAAAGAAGCAAACTTCCCCGCTATGTTGTGAAACTTCATCCGGATGATTTTTTTGACGAAGAAGACGCTGAAGTTCTGCGCTTTGATAATTTTGACGATGCCGTTGAGTGCTGCGCTGACCTGGGTATTCCGTTCTTTCTGGATGCAGGAAACAAAAAGCTGGTCTTCTGGTTTGTTCGTGTCGATGACGAAGGGTATCCGGAAATAGCCCGCTGTACGGAGCGGGAGTTTGCAACCATTCTTGCCGGTATCAGTGCCGGTGGTATGTACTGCCCGGAATGCGGCACAGTTCACTGGCCGGATGGAGTTACCCCACCCGTCTGATGCTTCCCCGTTTTGCCGATATTTTTCAGCAGGGTAACCGCTGGCTTAACTGGCTGGAGAAACAGCCGGAAGGTTCAGTGCGTCCGGTGGTGACTGAGTCAGTGACAAAAATCATGGCATGCGGGACCACGCTGATGGGCTACACGCAATGGTGCTGTTCATCACCGGACTGTTGCCACACAAAAAAGGTCTGCTTCCGGTGTAAAAGTCGCTCCTGCCCGCACTGCGGAGTGAAGGCTGGCGCACAGTGGATACAGTATCTGCTGAGTCTGGTCCCCGACTGCCCGTGGCAGCATATTGTGTTCACGCTTCCGTGCCAGTACTGGCCCCTGGTGTTCCACAACAGGTGGTTACTGGCAGAGATGAGCCGCATTGCTGCGGATGTGATACTGGAAATCTGCCGCCAGGCAGATGTGGAGGCGGGGATATTCACGGTAATCCACACATGGGGACGAGACCAGCAGTGGCATCCGCACATTCACCTGTCGACAACAGCCGGAGGCGTGACGTCAGGTCACTCCTGGAAAAACCTTCATTTTTACGCCCGTAAGGTGATGAGCATGTGGCGTTACCGGATAACGCGGCTACTGTCCCGGAAATACCCGGAGCTGGTAATACCGGATGAACTGGCAGTGGAAGGAAACAGCAAACGGGACTGGAATCGCTTCCTGGACAGTCATTACCGGCGGGGCTGGAATGTCAACGTATCCAGGGTGATGGATAACGCCACACATGTGGCAGTGTACTCTGGCTCTTACCTGAAAAAGCCGCCGGTGCCGATGAGCCGTCTGGAGCACTATGCAGGTCAGGATGAAATCGGTCTGCGTTACAACAGCCACCGGACAAAACGGGAAGAATACCTGGTGATGAGTGGCGATGAGTTCATGGAAAGGTTCTCCTGGCATGTAGCAGATAAGGGGTTCCGTATGGTGAGGTACTACGGTTTCCTGAGTCCGGTGAAGCGCCGGTTACTGGAAGAGGTGGTGTACGCCATAACGGAGACAGTGAGAAAAACAGCGATACAAATCAGGTGGAGAGGGATGTATCAGAGGTTACTGAAGGTTGACCCGCTGAAGTGCATTCTGTGCGGAAGTCAGATGCGTTTTACGGGGCTGAAGCGGGGTTACCGACTGGCAGAGCTGGTCCTGATGCATGAGCGACTGGCACGACAGCAGGTGTGCGGCTGAGAGCCGCTGAGGGGAAGTTGCGTCCATTTTACCGGAAACGGAGCAAAAAACCGCCATTCATACCCTGTATCAATCAGTGTCATCCTGTTTAATAGTCGTTTCCGCTCATATGGTGCACAAGGGGTGTTGAAGAAACATCCGTTTTGTGATGCTTTTGTAGTCTTTTGGGGATTTAAATTCCTATCGAT

>IS91-V113

CGAGTAGGCAGCCTGGCGGCTGCGGCTTGTCATGGCCTGGAATTACCGTTATAAAAAAAGATAATGTTATTGTCTTTCAGGTAGTTATATGGCCCGTTCAGCTAAACCCCGTAAACGCAAACCCGCACCACAAAGAAGCAAACTTCCCCGCTATGTTGTGAAACTTCATCCGGATGATTTTTTTGACGAAGAAGACGCTGAAGTTCTGCGCTTTGATAATTTTGACGATGCCGTTGAGTGCTGCGCTGACCTGGGTATTCCGTTCTTTCTGGATGCAGGAAACAAAAAGCTGGTCTTCTGGTTTGTTCGTGTCGATGACGAAGGGTATCCGGAAATAGCCCGCTGTACGGAGCGGGAGTTTGCAACCATTCTTGCCGGTATCAGTGCCGGTGGTATGTACTGCCCGGAATGCGGCACAGTTCACTGGCCGGATGGAGTTACCCCACCCGTCTGATGCTTCCCCGTTTTGCCGATATTTTTCAGCAGGGTAACCGCTGGCTTAACTGGCTGGAGAAACAGCCGGAAGGTTCAGTGCGTCCGGTGGTGACTGAGTCAGTGACAAAAATCATGGCATGCGGGACCACGCTGATGGGCTACACGCAATGGTGCTGTTCATCACCGGACTGTTGCCACACAAAAAAGGTCTGCTTCCGGTGTAAAAGTCGCTCCTGCCCGCACTGCGGAGTGAAGGCTGGCGCACAGTGGATACAGTATCTGCTGAGTCTGGTCCCCGACTGCCCGTGGCAGCATATTGTGTTCACGCTTCCGTGCCAGTACTGGCCCCTGGTGTTCCACAACAGGTGGTTACTGGCAGAGATGAGCCGCATTGCTGCGGATGTGATACTGGAAATCTGCCGCCAGGCAGATGTGGAGGCGGGGATATTCACGGTAATCCACACATGGGGACGAGACCAGCAGTGGCATCCGCACATTCACCTGTCGACAACAGCCGGAGGCGCGACGTCAGGTCACTCCTGGAAAAACCTTCATTTTTACGCCCGTAAGGTGATGAGCATGTGGCGTTACCGGATAACGCGGCTACTGTCCCGGAAATACCCGGAGCTGGTAATACCGGATGAACTGGCAGTGGAAGGAAACAGCAAACGGGACTGGAATCGCTTCCTGGACAGTCATTACCGGCGGGGCTGGAATGTCAACGTATCCAGGGTGATGGATAACGCCACACATGTGGCAGTGTACTCTGGCTCTTACCTGAAAAAGCCGCCGGTGCCGATGAGCCGTCTGGAGCACTATGCAGGTCAGGATGAAATCGGTCTGCGTTACAACAGCCACCGGACAAAACGGGAAGAATACCTGGTGATGAGTGGCGATGAGTTCATGGAAAGGTTCTCCTGGCATGTAGCAGATAAGGGGTTCCGTATGGTGAGGTACTACGGTTTCCTGAGTCCGGTGAAGCGCCGGTTACTGGAAGAGGTGGTGTACGCCATAACGGAGACAGTGAGAAAAACAGCGATACAAATCAGGTGGAGAGGGATGTATCAGAGGTTACTGAAGGTTGACCCGCTGAAGTGCATTCTGTGCGGAAGTCAGATGCGTTTTACGGGGCTGAAGCGGGGTTACCGACTGGCAGAGCTGGTCCTGATGCATGAGCGACTGGCACGACAGCAGGTGTGCGGCTGAGAGCCGCAGAGGGGAAGTTGCGTCCATTTTACCGGAAACGGAGCAAAAAACCGCCATTCATACCCTGTATCAATCAGTGTCATCCTGTTTAATAGTCGTTTCCGCTCATATGGTGCACAAGGGGTGTTGAAGAAACATCCGTTTTGTGATGCTTTTGTAGTCTTTTGGGGATTTAAATTCCTATCGAT

>IS91-V114

CGAGTAGGCAGCCTGGCGGCTGCGGCTTGTCATGGCCTGAAATTACCGTTATAAAAACAGACAATATCATTGTCTTTCAGGTAGTTATATGGCCCGTTCAGCTAAACCCCGTAAACGCAAACCCGCACCACAAAGAAGCAAACTTCCCCGCTATGTTGTGAAACTTCATCCGGATGATTTTTTTGACGAAGAAGACGCTGAAGTTCTGCGCTTTGATAATTTTGACGATGCCGTTGAGTGCTGCGCAGACCTGAATATTCCCTTCTTTGTGGATGCCGGAAACAAAAAGCTGGTCTTCTGGTTTGTTCGTGTCGATGACGAAGGGTATCCTGAAATAGCCCGCTGCACGGAGCGGGAGTTTGCGACCATTCTTGCCGGTATCAGCGCCGGCGGCATGTACTGCCCGGAGTGTGGCACGGTTCACTGGCCGGACGGAGTCCCCCCGCCCTTCTGATGCTTCCCCGTTTTGCCGACATTTTTCAGCAGGGAAACCGCTGGCTTAACTGGCTGGAGAAACAACCGGAAGGTTCAGTGCGTCCGGTAGTCATTGAGTCGGTGACAAAAATCATGGCGTGCGGAACCACGCTGATGGGGTACACACAGTGGTGCTGTTCATCTCCGGACTGTTGCCACACAAAAAAGGTCTGCTTCCGGTGTAAAAGTCGCTCCTGCCCGCACTGCGGAGTGAAGGCTGGCGCACAGTGGATACAGTATCTGCTGAGTCTGGTTCCCGACTGTCCGTGGCAGCATATTGTGTTCACACTTCCCTGCCAGTACTGGTCCCTGGTGTTCCACAACCGGTGGTTACTGGCAGAGATGAGCCGCATTGCTGCGGATGTGATACAGGAAATCTGCCGCCAGGCAGATGTGGTGCCGGGGATATTCACGGTCATCCACACATGGGGACGTGACCAGCAGTGGCATCCGCACATTCACCTGTCGACAACGGCCGGCGGCGTGACACCAGACCACACCTGGAAAAACCTTCATTTTTACGCCCGTAAGGTGATGAGCATGTGGCGTTACAGGATAACGTGGTTACTGTCACGAAAATACCCGGAGCTGGTGATACCGGATGCGCTGGCAGTTGAAGGAAGCAGCAGACGGGACTGGAATCGCTTCCTGGACAGTCATTACCGGCGGGGCTGGAATGTCAACGTATCCCGGGTGATGGATAACGCCACACATGTGGCGGTGTACTTCGGCTCTTACCTGAAAAAACCGCCGGTGCCGATGAGCCGTCTGGAGCACTATGCCGGTCAGGATGAAATTGGTCTGCGTTACAACAGTCACCGGACAAAACGGGAAGAATACCTGGTGATGAGTGGTGATGAGTTTATGGAAAGGTTCTCCTGGCATGTGGCGGATAAGGGGTTCCGTATGGTGAGGTACTACGGTTTCCTGAGTCCGGTAAAGCGCCGGTTACTGGAAGATGTTGTGTACGTCATAACGGAGACGGTGAGAAAGACGGCGATGCAAATCAGGTGGAGAGGGATGTATCAGCGGTTACTGAAGGTTGACCCGCTAAAGTGCATCCTGTGCGGATGTCAGATGCGTTTTACGGGGCTGAAGCGGGGTTACCGACTGGCAGAGCTGGTCCTGATGCATGAGCGACTGGCACGACAGCAGGTGTGCGGCTGAGAGCCGCAGAGGGGAAGTTGCGTCCATTTTACCGGAAACGGAGCAAAAAACCGCCATTCATACCCTGTATCAATCAGTGTCATCCTGTTTAATAGTCGTTTCCGCTCATATGGTGCACAAGGGGTGTTGAAGAAATATCCGTTTTGTGGTGCTTTTTTAGTCTTTTGGGGATTTAAATTCCTATCGAT

>IS91-V115

CGAGTAGGCAGCCTGGCGGCTGCGGCTTGTCATGGTCTGGAATTACCGTTATAAAAAAAGATAATATCATTGTCTTTCAGGTAGTTATATGGCCCGTTCAGCTAAACCCCGTAAACGCAAACCCGCACCACAAAGAAGCAAACTTCCCCGCTATGTTGTGAAACTTCATCCGGATGATTTTTTTGACGAAGAAGAAGCTGAAATTCTGCGCTTTGATAATTTTGACGATGCTGTTGAGTGCTGCGCTGACCTGGGTATTCCGTTCTTTCTGGATGCGGGAAACAAAAAGCTGGTCTTCTGGTTTGTTCGTGTTGATGACGAAGGGTATCCGGAAATAGCCCGCTGTACGGAGCGGGAGTTTGCGACCATTCTTGCCGGTATCAGCGCCGGCGGTATGTACTGCCAGGAATGCGGCACTGTTCACTGGCCGGATGGCGTTACCCCGCCCTTCTGATGCTTCCCCGTTTTGCAGACATCTTTCAGCAGGGAAACCGCTGGCTTAACTGGCTGGAAAAACAGCCGGAAGGTTCAGTGCGTCCGGTGGTGACTGAGTCGGTGACAAAAATCATGGCGTGCGGAACCACGCTGATGGGGTACACGCAGTGGTGTTGTTCATCCCCGGACTGCTGCCATACAAAAAAGGTCTGCTTCCGGTGTAAAAGTCGCTCCTGCCCACACTGCGGAGTGAAGGCCGGCGCACAGTGGATACAGTATCTGCTGAGTCTGGTCCCCGACTGCCCGTGGCAGCATATTGTGTTCACGCTTCCGTGCCAGTACTGGTCCCTGGTGTTCCACAACAGGTGGTTACTGGCAGAGATGAGTCGTATTGCTGCGGATGTGATACTGGAAATCTGCCGCCAGGCAGATGTGGAGCCTGGGATATTTACGGTAATCCACACATGGGGGCGTGACCAGCAGTGGCATCCGCATATCCATTTATCGACAACTGCCGGTGGTGTGACGTCGGGTCACACCAGGAAAAACCTTCATTTTTACGCCCGTAAGGTGATGAGCATGTGGCGTTACCGGATAACGCGGTTACTGTCACGGAAATACCCGGAGCTGGTGATACCGGATGAGCTGGCCGTTGAGGGAAGCAGCAGACGGGACTGGAATCGCTTCCTGGACACGCATTACCGGCGGGGCTGGAATGTCAACGTATCCAGGGTGATGGATAACGCCACACATGTGGCAGTGTACTTTGGCTCTTACCTGAAAAAGCCACCGGTGCCGATGAGTCGGCTGGAGCATTATGCCGGTCAGGATGAAATCGGTCTGCGTTACAACAGTCACCGTACAAAACGGGAAGAATACCTGTTGATGAGTGGAGATGAGTTCATGGAAAGGTTCTCCTGGCATGTAGCAGATAAGGGGTTCCGTATGGTGAGGTACTACGGTTTCCTGAGTCCGGTGAAGCGCCGCTTACTGGAAGAAGTTGTGTACGTCATAACGGAGACGGTGAGAAAAACGGCGATGCAAATCAGGTGGAGAGGGATGTATCAGCGGTTACTGAAGGTTGACCCGCTGAAGTGCATTCTGTGCGGAAGTCAGATGCGTTTTACGGGGCTGAAGCGGGGTTACCGACTGGCAGAGCTGGTCCTGATGCATGAGAGACTGGCACGACAGCAGGTGTGCGGCTGAGAGCCGCAGAGGGGAAGCTACGTCCATTTTGAAGGAAACGGAGCAAAAAAGCGCCATTCATCCCCTGTATCAATCAGTGCCCCCCTGTTTAATAGTCATTTCCGTTCATATGGTGCACAAGGGGTGTTGAAGAAACATCCGTTTTGTGGTGCTTTTTTAGTCTTTTGGGGATTTAAATTCCTATCGAT

>IS91-V116

CGAGTAGGCAGCCTGGCGGCTGCGGCTTGTCATGGTCTGAGATTACCGTTATAAAAACAGGCAATATCATTGTCTTTCAGGTGGTTATATGGCCCGTTCAGCTAAACCCCGTAAACGAAAACCTTCCCCACAAAGAAGCAAACTTCCCCGCTATGTTGTGAAACTTCATCCGGATGATTTTTTTGACGAAGAAGACGCTGAAGTTCTGCGCTTTGATAATTTTGACGATGCCGTTGAGTGCTGCGCTGACCTGGGTATTCCGTTCTTTCTGGATGCAGGAAACAAAAAGCTGGTCTTCTGGTTTGTTCGTGTCGATGACGAAGGGTATCCGGAAATAGCCCGCTGTACGGAGCGGGAGTTTGCAACCATTCTTGCCGGTATCAGTGCCGGTGGTATGTACTGCCCGGAATGCGGCACAGTTCACTGGCCGGATGGCGTTACCCCACCCGTCTGATGCTTCCCCGTTTTGCCGATATTTTTCAGCAGGGTAACCGCTGGCTTAACTGGCTGGAGAAACAGCCGGAAGGTTCAGTGCGTCCGGTGGTGACTGAGTCAGTGACAAAAATCATGGCATGCGGGACCACGCTGATGGGCTACACGCAATGGTACTGTTCATCACCGGACTGTTGCCACACAAAAAAGGTCTGCTTCCGGTGTAAAAGTCGCTCCTGCCCGCACTGCGGGGTGAAGGCTGGCGCACAGTGGATACAGTATCTGCTGAGTCTGGTCCCCGACTGCCCGTGGCAGCATATTGTGTTCACACTTCCCTGCCAGTACTGGCCCCTGATATTCCACAACAGGTGGTTACTGGCAGAGATGAGCCGCATTGCTGCGAATGTGATACTGGAAATCTGCCGTCAGGCGGACGTGGAGCCGGGGATATTCACGGTAATCCACACATGGGGGCGTGACCAGCAGTGGCATCCGCATATCCATTTATCGACAACTGCCGGTGGTGTGACGTCGGGCCACACCTGGAAAAATCTTCATTTTTACGCCCGTAAGGTGATGAGCATGTGGCGTTACCGGATAACGCGGCTACTGTCCCGGAAATACCCGGAGCTGGTAATACCGGATGAACTGGCAGTGGAAGGAAACAGCAAACGGGACTGGAATCGCTTTCTGGACACGCATTACCGCCGCGGCTGGAATGTCAACATATCCAGGGTGATGGATAACGCCACACATGTGGCGGTGTACTTCGGCTCTTACCTGAAAAAGCCACCGGTGCCGATGAGTCGGCTGGAGCATTATGCCGGTCAGGATGAAATCGGTCTGCGTTACAACAGTCACCGTACAAAACGGGAAGAATACCTGTTGATGAGTGGAGATGAGTTCATGGAAAGGTTCTCCTGGCATGTAGCAGATAAGGGGGTCCGTATGGTGAGGTACTACGGTTTCCTGAGTCCGGTGAAGCGCCGCTTACTGGAAGAGGTGGTGTACGTCATAACGGAGACGGTGAGAAAAACGGCGATGCAAATCAGGTGGAGAGGGATGTATCAGCGGTTACTGAAGGTTGACCCGCTGAAATGCGTCCTGTGCGGAAGTCAGATGCGGTTTACGGGGCTGAAGCGGGGATACCGTCTGGCAGAGCTGGTTATGATGCATGAGCCGCTGGCCCGAATGCAGTGTTGCGGCTGAGAGCCGCAGAGGGGAAGTTGCGTCCATTTTACGGGGAACGGAGCAAAAAACCACCATTCATACCCTTTATCAATCAGTGTTATCCTGTTTAATAGTCGTTTCCGTTCATATGGTGCATAAGGAGTGTTGAAGAAATATCCGTTTTGTGGTGTTTTTTAATCTTTTGGGGGTTTTAATTCCTATTGAT

>IS91-V117

CGAGTAGGCAGCCTGGCGGCTGCGGCTTGTCATGGTCTGAGATTACCGTTATAAAAACAGGCAATATCATTGTCTTTCAGGTGGTTATATGGCCCGTTCAGCTAAACCCCGTAAACGAAAACCTTCCCCACAAAGAAGCAAACTTCCCCGCTATGTTGTGAAACTTCATCCGGATGATTTTTTTGACGAAGAAGACGCTGAAGTTCTGCGCTTTGATAATTTTGACGATGCCGTTGAGTGCTGCGCTGACCTGGGTATTCCGTTCTTTCTGGATGCAGGAAACAAAAAGCTGGTCTTCTGGTTTGTTCGTGTCGATGACGAAGGGTATCCGGAAATAGCCCGCTGTACGGAGCGGGAGTTTGCAACCATTCTTGCCGGTATCAGTGCCGGTGGTATGTACTGCCCGGAATGCGGCACAGTTCACTGGCCGGATGGCGTTACCCCACCCGTCTGATGCTTCCCCGTTTTGCCGATATTTTTCAGCAGGGTAACCGCTGGCTTAACTGGCTGGAGAAACAGCCGGAAGGTTCAGTGCGTCCGGTGGTGACTGAGTCAGTGACAAAAATCATGGCATGCGGGACCACGCTGATGGGCTACACGCAATGGTACTGTTCATCACCGGACTGTTGCCACACAAAAAAGGTCTGCTTCCGGTGTAAAAGTCGCTCCTGCCCGCACTGCGGGGTGAAGGCTGGCGCACAGTGGATACAGTATCTGCTGAGTCTGGTCCCCGACTGCCCGTGGCAGCATATTGTGTTCACACTTCCCTGCCAGTACTGGCCCCTGATATTCCACAACAGGTGGTTACTGGCAGAGATGAGCCGCATTGCTGCGAATGTGATACTGGAAATCTGCCGTCAGGCGGACGTGGAGCCGGGGATATTCACGGTAATCCACACATGGGGGCGTGACCAGCAGTGGCATCCGCATATCCATTTATCGACAACTGCCGGTGGTGTGACGTCGGGCCACACCTGGAAAAATCTTCATTTTTACGCCCGTAAGGTGATGAGCATGTGGCGTTACCGGATAACGCGGCTACTGTCCCGGAAATACCCGGAGCTGGTAATACCGGATGAACTGGCAGTGGAAGGAAACAGCAAACGGGACTGGAATCGCTTTCTGGACACGCATTACCGCCGCGGCTGGAATGTCAACATATCCAGGGTGATGGATAACGCCACACATGTGGCGGTGTACTTCGGCTCTTACCTGAAAAAGCCACCGGTGCCGATGAGTCGGCTGGAGCATTATGCCGGTCAGGATGAAATCGGTCTGCGTTACAACAGTCACCGTACAAAACGGGAAGAATACCTGTTGATGAGTGGAGATGAGTTCATGGAAAGGTTCTCCTGGCATGTAGCAGATAAGGGGTTCCGTATGGTGAGGTACTACGGTTTCCTGAGTCCGGTGAAGCGCCGCTTACTGGAAGAGGTGGTGTACGTCATAACGGAGACGGTGAGAAAAACGGCGATGCAAATCAGGTGGAGAGGGATGTATCAGCGGTTACTGAAGGTTGACCCGCTGAAATGCGTCCTGTACGGAAGTCAGATGCGGTTTACGGGGCTGAAGCGGGGATACCGTCTGGCAGAGCTGGTTATGATGCATGAGCCGCTGGCCCGAATGCAGTGTTGCGGCTGAGAGCCGCAGAGGGGAAGTTGCGTCCATTTTACGGGGAACGGAGCAAAAAACCACCATTCATACCCTTTATCAATCAGTGTTATCCTGTTTAATAGTCGTTTCCGTTCATATGGTGCATAAGGAGTGTTGAAGAAATATCCGTTTTGTGGTGTTTTTTAATCTTTTGGGGGTTTTAATTCCTATTGAT

>IS91-V118

CGAGTAGGCAGCCTGGCGGCTGCGGCTTGTCATGGTCTGAGATTACCGTTATAAAAACAGGCAATATCATTGTCTTTCAGGTGGTTATATGGCCCGTTCAGCTAAACCCCGTAAACGAAAACCTTCCCCACAAAGAAGCAAACTTCCCCGCTATGTTGTGAAACTTCATCCGGATGATTTTTTTGACGAAGAAGACGCTGAAGTTCTGCGCTTTGATAATTTTGACGATGCCGTTGAGTGCTGCGCTGACCTGGGTATTCCGTTCTTTCTGGATGCAGGAAACAAAAAGCTGGTCTTCTGGTTTGTTCGTGTCGATGACGAAGGGTATCCGGAAATAGCCCGCTGTACGGAGCGGGAGTTTGCAACCATTCTTGCCGGTATCAGTGCCGGTGGTATGTACTGCCCGGAATGCGGCACAGTTCACTGGCCGGATGGCGTTACCCCACCCGTCTGATGCTTCCCCGTTTTGCCGATATTTTTCAGCAGGGTAACCGCTGGCTTAACTGGCTGGAGAAACAGCCGGAAGGTTCAGTGCGTCCGGTGGTGACTGAGTCAGTGACAAAAATCATGGCATGCGGGACCACGCTGATGGGCTACACGCAATGGTACTGTTCATCACCGGACTGTTGCCACACAAAAAAGGTCTGCTTCCGGTGTAAAAGTCGCTCCTGCCCGCACTGCGGGGTGAAGGCTGGCGCACAGTGGATACAGTATCTGCTGAGTCTGGTCCCCGACTGCCCGTGGCAGCATATTGTGTTCACACTTCCCTGCCAGTACTGGCCCCTGATATTCCACAACAGGTGGTTACTGGCAGAGATGAGCCGCATTGCTGCGAATGTGATACTGGAAATCTGCCGTCAGGCGGACGTGGAGCCGGGGATATTCACGGTAATCCACACATGGGGGCGTGACCAGCAGTGGCATCCGCATATCCATTTATCGACAACTGCCGGTGGTGTGACGTCGGGCCACACCTGGAAAAATCTTCATTTTTACGCCCGTAAGGTGATGAGCATGTGGCGTTACCGGATAACGCGGCTACTGTCCCGGAAATACCCGGAGCTGGTAATACCGGATGAACTGGCAGTGGAAGGAAACAGCAAACGGGACTGGAATCGCTTTCTGGACACGCATTACCGCCGCGGCTGGAATGTCAACATATCCAGGGTGATGGATAACGCCACACATGTGGCGGTGTACTTCGGCTCTTACCTGAAAAAGCCACCGGTGCCGATGAGTCGGCTGGAGCATTATGCCGGTCAGGATGAAATCGGTCTGCGTTACAACAGTCACCGTACAAAACGGGAAGAATACCTGTTGATGAGTGGAGATGAGTTCATGGAAAGGTTCTCCTGGCATGTAGCAGATAAGGGGTTCCGTATGGTGAGGTACTACGGTTTCCTGAGTCCGGTGAAGCGCCGCTTACTGGAAGAGGTGGTGTACGTCATAACGGAGACGGTGAGAAAAACGGCGATGCAAATCAGGTGGAGAGGGATGTATCAGCGGTTACTGAAGGTTGACCCGCTGAAATGCGTCCTGTACGGAAGTCAGATGCGGTTTACGGGGCTGAAGCGGGGATACCGTCTGGCAGAGCTGGTTATGATGCATGAGCCGCTGGCCCGAATGCAGTGTTGCGGCTGAGAGCCGCAGAGGGGAAGTTGCGTCCATTTTACGGGGAACGGAGCAAAAAACCACCATTCATACCCTTTATCAATCAGTGTTATCCTGTTTAATAGTCGTTTCCGTTCATATGGTGCATAAGGAGTGTTGAAGAAATATCCGTTTTGTGGTGTTTTTTTAATCTTTTGGGGGTTTTAATTCCTATTGAT

>IS91-V119

CGAGTAGGCAGCCTGGCGGCTGCGGCTTGTCATGGTCTGAGATTACCGTTATAAAAACAGGCAATATCATTGTCTTTCAGGTGGTTATATGGCCCGTTCAGCTAAACCCCGTAAACGAAAACCTTCCCCACAAAGAAGCAAACTTCCCCGCTATGTTGTGAAACTTCATCCGGATGATTTTTTTGACGAAGAAGACGCTGAAGTTCTGCGCTTTGATAATTTTGACGATGCCGTTGAGTGCTGCGCTGACCTGGGTATTCCGTTCTTTCTGGATGCAGGAAACAAAAAGCTGGTCTTCTGGTTTGTTCGTGTCGATGACGAAGGGTATCCGGAAATAGCCCGCTGTACGGAGCGGGAGTTTGCAACCATTCTTGCCGGTATCAGTGCCGGTGGTATGTACTGCCCGGAATGCGGCACAGTTCACTGGCCGGATGGCGTTACCCCACCCGTCTGATGCTTCCCCGTTTTGCCGATATTTTTCAGCAGGGTAACCGCTGGCTTAACTGGCTGGAGAAACAGCCGGAAGGTTCAGTGCGTCCGGTGGTGACTGAGTCAGTGACAAAAATCATGGCATGCGGGACCACGCTGATGGGCTACACGCAATGGTACTGTTCATCACCGGACTGTTGCCACACAAAAAAGGTCTGCTTCCGGTGTAAAAGTCGCTCCTGCCCGCACTGCGGGGTGAAGGCTGGCGCACAGTGGATACAGTATCTGCTGAGTCTGGTCCCCGACTGCCCGTGGCAGCATATTGTGTTCACACTTCCCTGCCAGTACTGGCCCCTGATATTCCACAACAGGTGGTTACTGGCAGAGATGAGCCGCATTGCTGCGAATGTGATACTGGAAATCTGCCGTCAGGCGGACGTGGAGCCGGGGATATTCACGGTAATCCACACATGGGGGCGTGACCAGCAGTGGCATCCGCATATCCATTTATCGACAACTGCCGGTGGTGTGACGTCGGGCCACACCTGGAAAAATATTCATTTTTACGCCCGTAAGGTGATGAGCATGTGGCGTTACCGGATAACGCGGCTACTGTCCCGGAAATACCCGGAGCTGGTAATACCGGATGAACTGGCAGTGGAAGGAAACAGCAAACGGGACTGGAATCGCTTTCTGGACACGCATTACCGCCGCGGCTGGAATGTCAACATATCCAGGGTGATGGATAACGCCACACATGTGGCGGTGTACTTCGGCTCTTACCTGAAAAAGCCACCGGTGCCGATGAGTCGGCTGGAGCATTATGCCGGTCAGGATGAAATCGGTCTGCGTTACAACAGTCACCGTACAAAACGGGAAGAATACCTGTTGATGAGTGGAGATGAGTTCATGGAAAGGTTCTCCTGGCATGTAGCAGATAAGGGGTTCCGTATGGTGAGGTACTACGGTTTCCTGAGTCCGGTGAAGCGCCGCTTACTGGAAGAGGTGGTGTACGTCATAACGGAGACGGTGAGAAAAACGGCGATGCAAATCAGGTGGAGAGGGATGTATCAGCGGTTACTGAAGGTTGACCCGCTGAAATGCGTCCTGTACGGAAGTCAGATGCGGTTTACGGGGCTGAAGCGGGGATACCGTCTGGCAGAGCTGGTTATGATGCATGAGCCGCTGGCCCGAATGCAGTGTTGCGGCTGAGAGCCGCAGAGGGGAAGTTGCGTCCATTTTACGGGGAACGGAGCAAAAAACCACCATTCATACCCTTTATCAATCAGTGTTATCCTGTTTAATAGTCGTTTCCGTTCATATGGTGCATAAGGAGTGTTGAAGAAATATCCGTTTTGTGGTGTTTTTTAATCTTTTGGGGGTTTTAATTCCTATTGAT

>IS91-V120

CGAGTAGGCAGCCTGGCGGCTGCGGCTTGTCATGGTCTGAGATTACCGTTATAAAAACAGGCAATATCATTGTCTTTCAGGTGGTTATATGGCCCGTTCAGCTAAACCCCGTAAACGAAAACCTTCCCCACAAAGAAGCAAACTTCCCCGCTATGTTGTGAAACTTCATCCGGATGATTTTTTTGACGAAGAAGACGCTGAAGTTCTGCGCTTTGATAATTTTGACGATGCCGTTGAGTGCTGCGCTGACCTGGGTATTCCGTTCTTTCTGGATGCAGGAAACAAAAAGCTGGTCTTCTGGTTTGTTCGTGTCGATGACGAAGGGTATCCGGAAATAGCCCGCTGTACGGAGCGGGAGTTTGCAACCATTCTTGCCGGTATCAGTGCCGGTGGTATGTACTGCCCGGAATGCGGCACAGTTCACTGGCCGGATGGCGTTACCCCACCCGTCTGATGCTTCCCCGTTTTGCCGATATTTTTCAGCAGGGTAACCGCTGGCTTAACTGGCTGGAGAAACAGCCGGAAGGTTCAGTGCGTCCGGTGGTGACTGAGTCAGTGACAAAAATCATGGCATGCGGGACCACGCTGATGGGCTACACGCAATGGTACTGTTCATCACCGGACTGTTGCCACACAAAAAAGGTCTGCTTCCGGTGTAAAAGTCGCTCCTGCCCGCACTGCGGGGTGAAGGCTGGCGCACAGTGGATACAGTATCTGCTAAGTCTGGTCCCCGACTGCCCGTGGCAGCATATTGTGTTCACACTTCCCTGCCAGTACTGGCCCCTGATATTCCACAACAGGTGGTTACTGGCAGAGATGAGCCGCATTGCTGCGAATGTGATACTGGAAATCTGCCGTCAGGCGGACGTGGAGCCGGGGATATTCACGGTAATCCACACATGGGGGCGTGACCAGCAGTGGCATCCGCATATCCATTTATCGACAACTGCCGGTGGTGTGACGTCGGGCCACACCTGGAAAAATCTTCATTTTTACGCCCGTAAGGTGATGAGCATGTGGCGTTACCGGATAACGCGGCTACTGTCCCGGAAATACCCGGAGCTGGTAATACCGGATGAACTGGCAGTGGAAGGAAACAGCAAACGGGACTGGAATCGCTTTCTGGACACGCATTACCGCCGCGGCTGGAATGTCAACATATCCAGGGTGATGGATAACGCCACACATGTGGCGGTGTACTTCGGCTCTTACCTGAAAAAGCCACCGGTGCCGATGAGTCGGCTGGAGCATTATGCCGGTCAGGATGAAATCGGTCTGCGTTACAACAGTCACCGTACAAAACGGGAAGAATACCTGTTGATGAGTGGAGATGAGTTCATGGAAAGGTTCTCCTGGCATGTAGCAGATAAGGGGTTCCGTATGGTGAGGTACTACGGTTTCCTGAGTCCGGTGAAGCGCCGCTTACTGGAAGAGGTGGTGTACGTCATAACGGAGACGGTGAGAAAAACGGCGATGCAAATCAGGTGGAGAGGGATGTATCAGCGGTTACTGAAGGTTGACCCGCTGAAATGCGTCCTGTACGGAAGTCAGATGCGGTTTACGGGGCTGAAGCGGGGATACCGTCTGGCAGAGCTGGTTATGATGCATGAGCCGCTGGCCCGAATGCAGTGTTGCGGCTGAGAGCCGCAGAGGGGAAGTTGCGTCCATTTTACGGGGAACGGAGCAAAAAACCACCATTCATACCCTTTATCAATCAGTGTTATCCTGTTTAATAGTCGTTTCCGTTCATATGGTGCATAAGGAGTGTTGAAGAAATATCCGTTTTGTGGTGTTTTTTAATCTTTTGGGGGTTTTAATTCCTATTGAT

>IS91-V121

CGAGTAGGCAGCCTGGCGGCTGCGGCTTGTCATGGTCTGAGATTACCGTTATAAAAACAGGCAATATCATTGTCTTTCAGGTGGTTATATGGCCCGTTCAGCTAAACCCCGTAAACGAAAACCTTCCCCACAAAGAAGCAAACTTCCCCGCTATGTTGTGAAACTTCATCCGGATGATTTTTTTGACGAAGAAGACGCTGAAGTTCTGCGCTTTGATAATTTTGACGATGCCGTTGAGTGCTGCGCTGACCTGGGTATTCCGTTCTTTCTGGATGCAGGAAACAAAAAGCTGGTCTTCTGGTTTGTTCGTGTCGATGACGAAGGGTATCCGGAAATAGCCCGCTGTACGGAGCGGGAGTTTGCAACCATTCTTGCCGGTATCAGTGCCGGTGGTATGTACTGCCCGGAATGCGGCACAGTTCACTGGCCGGATGGCGTTACCCCACCCGTCTGATGCTTCCCCGTTTTGCCGATATTTTTCAGCAGGGTAACCGCTGGCTTAACTGGCTGGAGAAACAGCCGGAAGGTTCAGTGCGTCCGGTGGTGACTGAGTCAGTGACAAAAATCATGGCATGTGGGACCACGCTGATGGGCTACACGCAATGGTACTGTTCATCACCGGACTGTTGCCACACAAAAAAGGTCTGCTTCCGGTGTAAAAGTCGCTCCTGCCCGCACTGCGGGGTGAAGGCTGGCGCACAGTGGATACAGTATCTGCTGAGTCTGGTCCCCGACTGCCCGTGGCAGCATATTGTGTTCACACTTCCCTGCCAGTACTGGCCCCTGATATTCCACAACAGGTGGTTACTGGCAGAGATGAGCCGCATTGCTGCGAATGTGATACTGGAAATCTGCCGTCAGGCGGACGTGGAGCCGGGGATATTCACGGTAATCCACACATGGGGGCGTGACCAGCAGTGGCATCCGCATATCCATTTATCGACAACTGCCGGTGGTGTGACGTCGGGCCACACCTGGAAAAATCTTCATTTTTACGCCCGTAAGGTGATGAGCATGTGGCGTTACCGGATAACGCGGCTACTGTCCCGGAAATACCCGGAGCTGGTAATACCGGATGAACTGGCAGTGGAAGGAAACAGCAAACGGGACTGGAATCGCTTTCTGGACACGCATTACCGCCGCGGCTGGAATGTCAACATATCCAGGGTGATGGATAACGCCACACATGTGGCGGTGTACTTCGGCTCTTACCTGAAAAAGCCACCGGTGCCGATGAGTCGGCTGGAGCATTATGCCGGTCAGGATGAAATCGGTCTGCGTTACAACAGTCACCGTACAAAACGGGAAGAATACCTGTTGATGAGTGGAGATGAGTTCATGGAAAGGTTCTCCTGGCATGTAGCAGATAAGGGGGTCCGTATGGTGAGGTACTACGGTTTCCTGAGTCCGGTGAAGCGCCGCTTACTGGAAGAGGTGGTGTACGTCATAACGGAGACGGTGAGAAAAACGGCGATGCAAATCAGGTGGAGAGGGATGTATCAGCGGTTACTGAAGGTTGACCCGCTGAAATGCGTCCTGTGCGGAAGTCAGATGCGGTTTACGGGGCTGAAGCGGGGATACCGTCTGGCAGAGCTGGTTATGATGCATGAGCCGCTGGCCCGAATGCAGTGTTGCGGCTGAGAGCCGCAGAGGGGAAGTTGCGTCCATTTTACGGGGAACGGAGCAAAAAACCACCATTCATACCCTTTATCAATCAGTGTTATCCTGTTTAATAGTCGTTTCCGTTCATATGGTGCATAAGGAGTGTTGAAGAAATATCCGTTTTGTGGTGTTTTTTAATCTTTTGGGGGTTTTAATTCCTATTGAT

>IS91-V122

CGAGTAGGCAGCCTGGCGGCTGCGGCTTGTCATGGTCTGAGATTACCGTTATAAAAACAGGCAATATCATTGTCTTTCAGGTGGTTATATGGCCCGTTCAGCTAAACCCCGTAAACGAAAACCTTCCCCACAAAGAAGCAAACTTCCCCGCTATGTTGTGAAACTTCATCCGGATGATTTTTTTGACGAAGAAGACGCTGAAGTTCTGCGCTTTGATAATTTTGACGATGCCGTTGAGTGCTGCGCTGACCTGGGTATTCCGTTCTTTCTGGATGCAGGAAACAAAAAGCTGGTCTTCTGGTTTGTTCGTGTCGATGACGAAGGGTATCCGGAAATAGCCCGCTGTACGGAGCGGGAGTTTGCAACCATTCTTGCCGGTATCAGTGCCGGTGGTATGTACTGCCCGGAATGCGGCACAGTTCACTGGCCGGATGGCGTTACCCCACCCGTCTGATGCTTCCCCGTTTTGCCGATATTTTTCAGCAGGGTAACCGCTGGCTTAACTGGCTGGAGAAACAGCCGGAAGGTTCAGTGCGTCCGGTGGTGACTGAGTCAGTGACAAAAATCATAGCATGCGGGACCACGCTGATGGGCTACACGCAATGGTACTGTTCATCACCGGACTGTTGCCACACAAAAAAGGTCTGCTTCCGGTGTAAAAGTCGCTCCTGCCCGCACTGCGGGGTGAAGGCTGGCGCACAGTGGATACAGTATCTGCTGAGTCTGGTCCCCGACTGCCCGTGGCAGCATATTGTGTTCACACTTCCCTGCCAGTACTGGCCCCTGATATTCCACAACAGGTGGTTACTGGCAGAGATGAGCCGCATTGCTGCGAATGTGATACTGGAAATCTGCCGTCAGGCGGACGTGGAGCCGGGGATATTCACGGTAATCCACACATGGGGGCGTGACCAGCAGTGGCATCCGCATATCCATTTATCGACAACTGCCGGTGGTGTGACGTCGGGCCACACCTGGAAAAATCTTCATTTTTACGCCCGTAAGGTGATGAGCATGTGGCGTTACCGGATAACGCGGCTACTGTCCCGGAAATACCCGGAGCTGGTAATACCGGATGAACTGGCAGTGGAAGGAAACAGCAAACGGGACTGGAATCGCTTTCTGGACACGCATTACCGCCGCGGCTGGAATGTCAACATATCCAGGGTGATGGATAACGCCACACATGTGGCGGTGTACTTCGGCTCTTACCTGAAAAAGCCACCGGTGCCGATGAGTCGGCTGGAGCATTATGCCGGTCAGGATGAAATCGGTCTGCGTTACAACAGTCACCGTACAAAACGGGAAGAATACCTGTTGATGAGTGGAGATGAGTTCATGGAAAGGTTCTCCTGGCATGTAGCAGATAAGGGGGTCCGTATGGTGAGGTACTACGGTTTCCTGAGTCCGGTGAAGCGCCGCTTACTGGAAGAGGTGGTGTACGTCATAACGGAGACGGTGAGAAAAACGGCGATGCAAATCAGGTGGAGAGGGATGTATCAGCGGTTACTGAAGGTTGACCCGCTGAAATGCGTCCTGTGCGGAAGTCAGATGCGGTTTACGGGGCTGAAGCGGGGATACCGTCTGGCAGAGCTGGTTATGATGCATGAGCCGCTGGCCCGAATGCAGTGTTGCGGCTGAGAGCCGCAGAGGGGAAGTTGCGTCCATTTTACGGGGAACGGAGCAAAAAACCACCATTCATACCCTTTATCAATCAGTGTTATCCTGTTTAATAGTCGTTTCCGTTCATATGGTGCATAAGGAGTGTTGAAGAAATATCCGTTTTGTGGTGTTTTTTAATCTTTTGGGGGTTTTAATTCCTATTGAT

>IS91-V123

CGAGTAGGCAGCCTGGCGGCTGCGGCTTGTCATGGTCTGAGATTACCGTTATAAAAACAGGCAATATCATTGTCTTTCAGGTGGTTATATGGCCCGTTCAGCTAAACCCCGTAAACGAAAACCTTCCCCACAAAGAAGCAAACTTCCCCGCTATGTTGTGAAACTTCATCCGGATGATTTTTTTGACGAAGAAGACGCTGAAGTTCTGCGCTTTGATAATTTTGACGATGCCGTTGAGTGCTGCGCTGACCTGGGTATTCCGTTCTTTCTGGATGCAGGAAACAAAAAGCTGGTCTTCTGGTTTGTTCGTGTCGATGACGAAGGGTATCCGGAAATAGCCCGCTGTACGGAGCGGGAGTTTGCAACCATTCTTGCCGGTATCAGTGCCGGTGGTATGTACTGCCCGGAATGCGGCACAGTTCACTGGCCGGATGGCGTTACCCCACCCGTCTGATGCTTCCCCGTTTTGCCGATATTTTTCAGCAGGGTAACCGCTGGCTTAACTGGCTGGAGAAACAGCCGGAAGGTTCAGTGCGTCCGGTGGTGACTGAGTCAGTGACAAAAATCATGGCATGCGGGACCACGCTGATGGGCTACACGCAATGGTACTGTTCATCACCGGACTGTTGCCACACAAAAAAGGTCTGCTTCCGGTGTAAAAGTCGCTCCTGCCCGCACTGCGGGGTGAAGGCTGGCGCACAGTGGATACAGTATCTGCTGAGTCTGGTCCCCGACTGCCCGTGGCAGCATATTGTGTTCACACTTCCCTGCCAGTACTGGCCCCTGATATTCCACAACAGGTGGTTACTGGCAGAGATGAGCCGCATTGCTGCGAATGTGATACTGGAAATCTGCCGTCAGGCGGACGTGGAGCCGGGGATATTCACGGTAATCCACACATGGGGGCGTGACCAGCAGTGGCATCCGCATATCCATTTATCGACAACTGCCGGTGGTGTGACGTCGGGCCACACCTGGAAAAATCTTCATTTTTACGCCCGTAAGGTGATGAGCATGTGGCGTTACCGGATAACGCGGCTACTGTCCCGGAAATACCCGGAGCTGGTAATACCGGATGAACTGGCAGTGGAAGGAAACAGCAAACGGGACTGGAATCGCTTTCTGGACACGCATTACCGCCGCGGCTGGAATGTCAACATATCCAGGGTGATGGATAACGCCACACATGTGGCGGTGTACTTCGGCTCTTACCTGAAAAAGCCACCGGTGCCGATGAGTCGGCTGGAGCATTATGCCGGTCAGGATGAAATCGGTCTGCGTTACAACAGTCACCGTACAAAACGGGAAGAATACCTGTTGATGAGTGGAGATGAGTTCATGGAAAGGTTCTCCTGGCATGTAGCAGATAAGGGGGGCCGTATGGTGAGGTACTACGGTTTCCTGAGTCCGGTGAAGCGCCGCTTACTGGAAGAGGTGGTGTACGTCATAACGGAGACGGTGAGAAAAACGGCGATGCAAATCAGGTGGAGAGGGATGTATCAGCGGTTACTGAAGGTTGACCCGCTGAAATGCGTCCTGTGCGGAAGTCAGATGCGGTTTACGGGGCTGAAGCGGGGATACCGTCTGGCAGAGCTGGTTATGATGCATGAGCCGCTGGCCCGAATGCAGTGTTGCGGCTGAGAGCCGCAGAGGGGAAGTTGCGTCCATTTTACGGGGAACGGAGCAAAAAACCACCATTCATACCCTTTATCAATCAGTGTTATCCTGTTTAATAGTCGTTTCCGTTCATATGGTGCATAAGGAGTGTTGAAGAAATATCCGTTTTGTGGTGTTTTTTAATCTTTTGGGGGTTTTAATTCCTATTGAT

>IS91-V124

CGAGTAGGCAGCCTGGCGGCTGCGGCTTGTCATGGTCTGAGATTACCGTTATAAAAACAGGCAATATCATTGTCTTTCAGGTGGTTATATGGCCCGTTCAGCTAAACCCCGTAAACGAAAACCTTCCCCACAAAGAAGCAAACTTCCCCGCTATGTTGTGAAACTTCATCCGGATGATTTTTTTGACGAAGAAGACGCTGAAGTTCTGCGCTTTGATAATTTTGACGATGCCGTTGAGTGCTGCGCTGACCTGGGTATTCCGTTCTTTCTGGATGCAGGAAACAAAAAGCTGGTCTTCTGGTTTGTTCGTGTCGATGACGAAGGGTATCCGGAAATAGCCCGCTGTACGGAGCGGGAGTTTGCAACCATTCTTGCCGGTATCAGTGCCGGTGGTATGTACTGCCCGGAATGCGGCACAGTTCACTGGCCGGATGGCGTTACCCCACCCGTCTGATGCTTCCCCGTTTTGCCGATATTTTTCAGCAGGGTAACCGCTGGCTTAACTGGCTGGAGAAACAGCCGGAAGGTTCAGTGCGTCCGGTGGTGACTGAGTCAGTGACAAAAATCATGGCATGCGGGACCACGCTGATGGGCTACACGCAATGGTACTGTTCATCACCGGACTGTTGCCACACAAAAAAGGTCTGCTTCCGGTGTAAAAGTCGCTCCTGCCCGCACTGCGGGGTGAAGGCTGGCGCACAGTGGATACAGTATCTGCTGAGTCTGGTCCCCGACTGCCCGTGGCAGCATATTGTGTTCACACTTCCCTGCCAGTACTGGCCCCTGATATTCCACAACAGGTGGTTACTGGCAGAGATGAGCCGCATTGCTGCGAATGTGATACTGGAAATCTGCCGTCAGGCGGACGTGGAGCCGGGGATATTCACGGTAATCCACACATGGGGGCGTGACCAGCAGTGGCATCCGCATATCCATTTATCGACAACTGCCGGTGGTGTGACGTCGGGCCACACCTGGAAAAATCTTCATTTTTACGCCCGTAAGGTGATGAGCATGTGGCGTTACCGGATAACGCGGCTACTGTCCCGGAAATACCCGGAGCTGGTAATACCGGATGAACTGGCAGTGGAAGGAAACAGCAAACGGGACTGGAATCGCTTTCTGGACACGCATTACCGCCGCGGCTGGAATGTCAACATATCCAGGGTGATGGATAACGCCACACATGTGGCGGTGTACTTCGGCTCTTACCTGAAAAAGCCACCGGTGCCGATGAGTCGGCTGGAGCATTATGCCGGTCAGGATGAAATCGGTCTGCGTTACAACAGTCACCGTACAAAACGGGAAGAATACCTGTTGATGAGTGGAGATGAGTTCATGGAAAGGTTCTCCTGGCATGTAGCAGATAAGGGGTTCCGTATGGTGAGGTACTACGGTTTCCTGAGTCCGGTGAAGCGCCGCTTACTGGAAGAGGTGGTGTACGTCATAACGGAGACGGTGAGAAAAACGGCGATGCAAATCAGGTGGAGAGGGATGTATCAGCGGTTACTGAAGGTTGACCCGCTGAAATGCGTCCTGTACGGAAGTCAGATGCGGTTTACGGGGCTGAAGCGGGGATACCGTCTGGCAGAGCTGGTTATGATGCATGAGCCGCTGGCCCGAATGCAGTGTTGCGGCTGAGAGCCGCAGAGGGGAAGTTGCGTCCATTTTACGGGGAACGGAGCAAAAAACCACCATTCATACCCTTTATCAATCTGTGTTATCCTGTTTAATAGTCGTTTCCGTTCATATGGTGCATAAGGAGTGTTGAAGAAATATCCGTTTTGTGGTGTTTTTTAATCTTTTGGGGGTTTTAATTCCTATTGAT

>IS91-V125

CGAGTAGGCAGCCTGGCGGCTGCGGCTTGTCATGGTCTGAGATTACCGTTATAAAAACAGGCAATATCATTGTCTTTCAGGTGGTTATATGGCCCGTTCAGCTAAACCCCGTAAACGAAAACCTTCCCCACAAAGAAGCAAACTTCCCCGCTATGTTGTGAAACTTCATCCGGATGATTTTTTTGACGAAGAAGACGCTGAAGTTCTGCGCTTTGATAATTTTGACGATGCCGTTGAGTGCTGCGCTGACCTGGGTATTCCGTTCTTTCTGGATGCAGGAAACAAAAAGCTGGTCTTCTGGTTTGTTCGTGTCGATGACGAAGGGTATCCGGAAATAGCCCGCTGTACGGAGCGGGAGTTTGCAACCATTCTTGCCGGTATCAGTGCCGGTGGTATGTACTGCCCGGAATGCGGCACAGTTCACTGGCCGGATGGCGTTACCCCACCCGTCTGATGCTTCCCCGTTTTGCCGATATTTTTCAGCAGGGTAACCGCTGGCTTAACTGGCTGGAGAAACAGCCGGAAGGTTCAGTGCGTCCGGTGGTGACTGAGTCAGTGACAAAAATCATGGCATGCGGGACCACGCTGATGGGCTACACGCAATGGTACTGTTCATCACCGGACTGTTGCCACACAAAAAAGGTCTGCTTCCGGTGTAAAAGTCGCTCCTGCCCGCACTGCGGGGTGAAGGCTGGCGCACAGTGGATACAGTATCTGCTGAGTCTGGTCCCCGACTGCCCGTGGCAGCATATTGTGTTCACACTTCCCTGCCAGTACTGGCCCCTGATATTCCACAACAGGTGGTTACTGGCAGAGATGAGCCGCATTGCTGCGAATGTGATACTGGAAATCTGCCGTCAGGCGGACGTGGAGCCGGGGATATTCACGGTAATCCACACATGGGGGCGTGACCAGCAGTGGCATCCGCATATCCATTTATCGACAACTGCCGGTGGTGTGACGTCGGGCCACACCTGGAAAAATCTTCATTTTTACGCCCGTAAGGTGATGAGCATGTGGCGTTACCGGATAACGCGGCTACTGTCCCGGAAATACCCGGAGCTGGTAATACCGGATGAACTGGCAGTGGAAGGAAACAGCAAACGGGACTGGAATCGCTTTCTGGACACGCATTACCGCCGCGGCTGGAATGTCAACATATCCAGGGTGATGGATAACGCCACACATGTGGCGGTGTACTTCGGCTCTTACCTGAAAAAGCCACCGGTGCCGATGAGTCGGCTGGAGCATTATGCCGGTCAGGATGAAATCGGTCTGCGTTACAACAGTCACCGTACAAAACGGGAAGAATACCTGTTGATGAGTGGAGATGAGTTCATGGAAAGGTTCTCCTGGCATGTAGCAGATAAGGGGTTCCGTATGGTGAGGTACTACGGTTTCCTGAGTCCGGTGAAGCGCCGCTTACTGGAAGAGGTGGTGTACGTCATAACGGAGACGGTGAGAAAAACGGCGATGCAAATCAGGTGGAGAGGGATGTATCAGCGGTTACTGAAGGTTGACCCGCTGAAATGCGTCCTGTACGGAAGTCAGATGCGGTTTACGGGGCTGAAGCGGGGATACCGTCTGGCAGAGCTGGTTATGATGCATGAGCCGCTGGCCCGAATGCAGTGTTGCGGCTGAGAGCCGCAGAGGGGAAGTTGCGTCCATTTTACGGGGAACGGAGCAAAAAACCACCATTCATACCCTTTATCAATCAGTGTTATCCCGTTTAATAGTCGTTTCCGTTCATATGGTGCATAAGGAGTGTTGAAGAAATATCCGTTTTGTGGTGTTTTTTAATCTTTTGGGGGTTTTAATTCCTATTGAT

>IS91-V126

CGAGTAGGCAGCCTGGCGGCTGCGGCTTGTCATGGTCTGAGATTACCGTTATAAAAACAGGCAATATCATTGTCTTTCAGGTGGTTATATGGCCCGTTCAGCTAAACCCCGTAAACGAAAACCTTCCCCACAAAGAAGCAAACTTCCCCGCTATGTTGTGAAACTTCATCCGGATGATTTTTTTGACGAAGAAGACGCTGAAGTTCTGCGCTTTGATAATTTTGACGATGCCGTTGAGTGCTGCGCTGACCTGGGTATTCCGTTCTTTCTGGATGCAGGAAACAAAAAGCTGGTCTTCTGGTTTGTTCGTGTCGATGACGAAGGGTATCCGGAAATAGCCCGCTGTACGGAGCGGGAGTTTGCAACCATTCTTGCCGGTATCAGTGCCGGTGGTATGTACTGCCCGGAATGCGGCACAGTTCACTGGCCGGATGGCGTTACCCCACCCGTCTGATGCTTCCCCGTTTTGCCGATATTTTTCAGCAGGGTAACCGCTGGCTTAACTGGCTGGAGAAACAGCCGGAAGGTTCAGTGCGTCCGGTGGTGACTGAGTCAGTGACAAAAATCATGGCATGCGGGACCACGCTGATGGGCTACACGCAATGGTACTGTTCATCACCGGACTGTTGCCACACAAAAAAGGTCTGCTTCCGGTGTAAAAGTCGCTCCTGCCCGCACTGCGGGGTGAAGGCTGGCGCACAGTGGATACAGTATCTGCTGAGTCTGGTCCCCGACTGCCCGTGGCAGCATATTGTGTTCACACTTCCCTGCCAGTACTGGCCCCTGATATTCCACAACAGGTGGTTACTGGCAGAGATGAGCCGCATTGCTGCGAATGTGATACTGGAAATCTGCCGTCAGGCGGACGTGGAGCCGGGGATATTCACGGTAATCCACACATGGGGGCGTGACCAGCAGTGGCATCCGCATATCCATTTATCGACAACTGCCGGTGGTGTGACGTCGGGCCACACCTGGAAAAATCTTCATTTTTACGCCCGTAAGGTGATGAGCATGTGGCGTTACCGGATAACGCGGCTACTGTCCCGGAAATACCCGGAGCTGGTAATACCGGATGAACTGGCAGTGGAAGGAAACAGCAAACGGGACTGGAATCGCTTTCTGGACACGCATTACCGCCGCGGCTGGAATGTCAACATATCCAGGGTGATGGATAACGCCACACATGTGGCGGTGTACTTCGGCTCTTACCTGAAAAAGCCACCGGTGCCGATGAGTCGGCTGGAGCATTATGCCGGTCAGGATGAAATCGGTCTGCGTTACAACAGTCACCGTACAAAACGGGAAGAATACCTGTTGATGAGTGGAGATGAGTTCATGGAAAGGTTCTCCTGGCATGTAGCAGATAAGGGGTTCCGTATGGTGAGGTACTACGGTTTCCTGAGTCCGGTGAAGCGCCGCTTACTGGAAGAGGTGGTGTACGTCATAACGGAGACGGTGAGAAAAACGGCGATGCAAATCAGGTGGAGAGGGATGTATCAGCGGTTACTGAAGGTTGACCCGCTGAAATGCGTCCTGTACGGAAGTCAGATGCGGTTTACGGGGCTGAAGCGGGGATACCGTCTGGCAGAGCTGGTTATGATGCATGAGCCGCTGGCCCGAATGCAGTGTTGCGGCTGAGAGCCGCAGAGGGGAAGTTGCGTCCATTTTACGGGGAACGGAGCAAAAAACCACCATTCATACCCTTTATCAATCAGTGTTACCCTGTTTAATAGTCGTTTCCGTTCATATGGTGCATAAGGAGTGTTGAAGAAATATCCGTTTTGTGGTGTTTTTTAATCTTTTGGGGGTTTTAATTCCTATTGAT

>IS91-V127

CGAGTAGGCAGCCTGGCGGCTGCGGCTTGTCATGGTCTGAGATTACCGTTATAAAAACAGGCAATATCATTGTCTTTCAGGTGGTTATATGGCCCGTTCAGCTAAACCCCGTAAACGAAAACCTTCCCCACAAAGAAGCAAACTTCCCCGCTATGTTGTTAAACTTCATCCGGATGATTTTTTTGACGAAGAAGACGCTGAAGTTCTGCGCTTTGATAATTTTGACGATGCCGTTGAGTGCTGCGCTGACCTGGGTATTCCGTTCTTTCTGGATGCAGGAAACAAAAAGCTGGTCTTCTGGTTTGTTCGTGTCGATGACGAAGGGTATCCGGAAATAGCCCGCTGTACGGAGCGGGAGTTTGCAACCATTCTTGCCGGTATCAGTGCCGGTGGTATGTACTGCCCGGAATGCGGCACAGTTCACTGGCCGGATGGCGTTACCCCACCCGTCTGATGCTTCCCCGTTTTGCCGATATTTTTCAGCAGGGTAACCGCTGGCTTAACTGGCTGGAGAAACAGCCGGAAGGTTCAGTGCGTCCGGTGGTGACTGAGTCAGTGACAAAAATCATGGCATGCGGGACCACGCTGATGGGCTACACGCAATGGTACTGTTCATCACCGGACTGTTGCCACACAAAAAAGGTCTGCTTCCGGTGTAAAAGTCGCTCCTGCCCGCACTGCGGGGTGAAGGCTGGCGCACAGTGGATACAGTATCTGCTGAGTCTGGTCCCCGACTGCCCGTGGCAGCATATTGTGTTCACACTTCCCTGCCAGTACTGGCCCCTGATATTCCACAACAGGTGGTTACTGGCAGAGATGAGCCGCATTGCTGCGAATGTGATACTGGAAATCTGCCGTCAGGCGGACGTGGAGCCGGGGATATTCACGGTAATCCACACATGGGGGCGTGACCAGCAGTGGCATCCGCATATCCATTTATCGACAACTGCCGGTGGTGTGACGTCGGGCCACACCTGGAAAAATCTTCATTTTTACGCCCGTAAGGTGATGAGCATGTGGCGTTACCGGATAACGCGGCTACTGTCCCGGAAATACCCGGAGCTGGTAATACCGGATGAACTGGCAGTGGAAGGAAACAGCAAACGGGACTGGAATCGCTTTCTGGACACGCATTACCGCCGCGGCTGGAATGTCAACATATCCAGGGTGATGGATAACGCCACACATGTGGCGGTGTACTTCGGCTCTTACCTGAAAAAGCCACCGGTGCCGATGAGTCGGCTGGAGCATTATGCCGGTCAGGATGAAATCGGTCTGCGTTACAACAGTCACCGTACAAAACGGGAAGAATACCTGTTGATGAGTGGAGATGAGTTCATGGAAAGGTTCTCCTGGCATGTAGCAGATAAGGGGGTCCGTATGGTGAGGTACTACGGTTTCCTGAGTCCGGTGAAGCGCCGCTTACTGGAAGAGGTGGTGTACGTCATAACGGAGACGGTGAGAAAAACGGCGATGCAAATCAGGTGGAGAGGGATGTATCAGCGGTTACTGAAGGTTGACCCGCTGAAATGCGTCCTGTGCGGAAGTCAGATGCGGTTTACGGGGCTGAAGCGGGGATACCGTCTGGCAGAGCTGGTTATGATGCATGAGCCGCTGGCCCGAATGCAGTGTTGCGGCTGAGAGCCGCAGAGGGGAAGTTGCGTCCATTTTACGGGGAACGGAGCAAAAAACCACCATTCATACCCTTTATCAATCAGTGTTATCCTGTTTAATAGTCGTTTCCGTTCATATGGTGCATAAGGAGTGTTGAAGAAATATCCGTTTTGTGGTGTTTTTTAATCTTTTGGGGGTTTTAATTCCTATTGAT

>IS91-V128

CGAGTAGGCAGCCTGGCGGCTGCGGCTTGTCATGGTCTGAGATTACCGTTATAAAAACAGGCAATATCATTGTCTTTCAGGTGGTTATATGGCCCGTTCAGCTAACCCCCGTAAACGAAAACCTTCCCCACAAAGAAGCAAACTTCCCCGCTATGTTGTGAAACTTCATCCGGATGATTTTTTTGACGAAGAAGACGCTGAAGTTCTGCGCTTTGATAATTTTGACGATGCCGTTGAGTGCTGCGCTGACCTGGGTATTCCGTTCTTTCTGGATGCAGGAAACAAAAAGCTGGTCTTCTGGTTTGTTCGTGTCGATGACGAAGGGTATCCGGAAATAGCCCGCTGTACGGAGCGGGAGTTTGCAACCATTCTTGCCGGTATCAGTGCCGGTGGTATGTACTGCCCGGAATGCGGCACAGTTCACTGGCCGGATGGCGTTACCCCACCCGTCTGATGCTTCCCCGTTTTGCCGATATTTTTCAGCAGGGTAACCGCTGGCTTAACTGGCTGGAGAAACAGCCGGAAGGTTCAGTGCGTCCGGTGGTGACTGAGTCAGTGACAAAAATCATGGCATGCGGGACCACGCTGATGGGCTACACGCAATGGTACTGTTCATCACCGGACTGTTGCCACACAAAAAAGGTCTGCTTCCGGTGTAAAAGTCGCTCCTGCCCGCACTGCGGGGTGAAGGCTGGCGCACAGTGGATACAGTATCTGCTGAGTCTGGTCCCCGACTGCCCGTGGCAGCATATTGTGTTCACACTTCCCTGCCAGTACTGGCCCCTGATATTCCACAACAGGTGGTTACTGGCAGAGATGAGCCGCATTGCTGCGAATGTGATACTGGAAATCTGCCGTCAGGCGGACGTGGAGCCGGGGATATTCACGGTAATCCACACATGGGGGCGTGACCAGCAGTGGCATCCGCATATCCATTTATCGACAACTGCCGGTGGTGTGACGTCGGGCCACACCTGGAAAAATATTCATTTTTACGCCCGTAAGGTGATGAGCATGTGGCGTTACCGGATAACGCGGCTACTGTCCCGGAAATACCCGGAGCTGGTAATACCGGATGAACTGGCAGTGGAAGGAAACAGCAAACGGGACTGGAATCGCTTTCTGGACACGCATTACCGCCGCGGCTGGAATGTCAACATATCCAGGGTGATGGATAACGCCACACATGTGGCGGTGTACTTCGGCTCTTACCTGAAAAAGCCACCGGTGCCGATGAGTCGGCTGGAGCATTATGCCGGTCAGGATGAAATCGGTCTGCGTTACAACAGTCACCGTACAAAACGGGAAGAATACCTGTTGATGAGTGGAGATGAGTTCATGGAAAGGTTCTCCTGGCATGTAGCAGATAAGGGGTTCCGTATGGTGAGGTACTACGGTTTCCTGAGTCCGGTGAAGCGCCGCTTACTGGAAGAGGTGGTGTACGTCATAACGGAGACGGTGAGAAAAACGGCGATGCAAATCAGGTGGAGAGGGATGTATCAGCGGTTACTGAAGGTTGACCCGCTGAAATGCGTCCTGTACGGAAGTCAGATGCGGTTTACGGGGCTGAAGCGGGGATACCGTCTGGCAGAGCTGGTTATGATGCATGAGCCGCTGGCCCGAATGCAGTGTTGCGGCTGAGAGCCGCAGAGGGGAAGTTGCGTCCATTTTACGGGGAACGGAGCAAAAAACCACCATTCATACCCTTTATCAATCAGTGTTATCCTGTTTAATAGTCGTTTCCGTTCATATGGTGCATAAGGAGTGTTGAAGAAATATCCGTTTTGTGGTGTTTTTTAATCTTTTGGGGGTTTTAATTCCTATTGAT

>IS91-V129

CGAGTAGGCAGCCTGGCGGCTGCGGCTTGTCATGGTCTGAGATTACCGTTATAAAAACAGGCAATATCATTGTCTTTCAGGTGGTTATATGGCCCGTTCAGCTAAACCCCGTAAACGAAAACCTTCCCCACAAAGAAGCAAACTTCCCCGCTATGTTGTGAAACTTCATCCGGATGATTTTTTTGACGAAGAAGACGCTGAAGTTCTGCGCTTTGATAATTTTGACGATGCCGTTGAGTGCTGCGCTGACCTGGGTATTCCGTTCTTTCTGGATGCAGGAAACAAAAAGCTGGTCTTCTGGTTTGTTCGTGTCGATGACGAAGGGTATCCGGAAATAGCCCGCTGTACGGAGCGGGAGTTTGCAACCATTCTTGCCGGTATCAGTGCCGGTGGTATGTACTGCCCGGAATGCGGCACAGTTCACTGGCCGGATGGCGTTACCCCACCCGTCTGATGCTTCCCCGTTTTGCCGATATTTTTCAGCAGGGTAACCGCTGGCTTAACTGGCTGGAGAAACAGCCGGAAGGTTCAGTGCGTCCGGTGGTGACTGAGTCAGTGACAAAAATCATGGCATGCGGGACCACGCTGATGGGCTACACGCAATGGTACTGTTCATCACCGGACTGTTGCCACACAAAAAAGGTCTGCTTCCGGTGTAAAAGTCGCTCCTGCCCGCACTGCGGGGTGAAGGCTGGCGCACAGTGGATACAGTATCTGCTGAGTCTGGTCCCCGACTGCCCGTGGCAGCATATTGTGTTCACACTTCCCTGCCAGTACTGGCCCCTGATATTCCACAACAGGTGGTTACTGGCAGAGATGAGCCGCATTGCTGCGAATGTGATACTGGAAATCTGCCGTCAGGCGGACGTGGAGCCGGGGATATTCACGGTAATCCACACATGGGGGCGTGACCAGCAGTGGCATCCGCATATCCATTTATCGACAACTGCCGGTGGTGTGACGTCGGGCCACACCTGGAAAAATATTCATTTTTACGCCCGTAAGGTGATGAGCATGTGGCGTTACCGGATAACGCGGCTACTGTCCCGGAAATACCCGGAGCTGGTAATACCGGATGAACTGGCAGTGGAAGGAAACAGCAAACGGGACTGGAATCGCTTTCTGGACACGCATTACCGCCGCGGCTGGAATGTCAACATATCCAGGGTGATGGATAACGCCACACATGTGGCGGTGTACTTCGGCTCTTACCTGAAAAAGCCACCGGTGCCGATGAGTCGGCTGGAGCATTATGCCGGTCAGGATGAAATCGGTCTGCGTTACAACAGTCACCGTACAAAACGGGAAGAATACCTGTTGATGAGTGGAGATGAGTTCATGGAAAGGTTCTCCTGGCATGTAGCAGATAAGGGGTTCCGTATGGTGAGGTACTACGGTTTCCTGAGTCCGGTGAAGCGCCGCTTACTGGAAGAGGTGGTGTACGTCATAACGGAGACGGTGAGAAAAACGGCGATGCAAATCAGGTGGAGAGGGATGTATCAGCGGTTACTGAAGGTTGACCCGCTGAAATGCGTCCTGTACGGAAGTCAGATGCGGTTTACGGGGCTGAAGCGGGGATACCGTCTGGCAGAGCTGGTTATGATGCATGAGCCGCTGGCCCGAATGCAGTGTTGCGGCTGAGAGCCGCAGAGGGGAAGTTGCGTCCATTTTACGGGGAACGGAGCAAAAAACCACCATCCATACCCTTTATCAATCAGTGTTATCCTGTTTAATAGTCGTTTCCGTTCATATGGTGCATAAGGAGTGTTGAAGAAATATCCGTTTTGTGGTGTTTTTTAATCTTTTGGGGGTTTTAATTCCTATTGAT

>IS91-V130

CGAGTAGGCAGCCTGACGGCTGCGGCTTGTCATGGTCTGAGATTACCGTTATAAAAACAGGCAATATCATTGTCTTTCAGGTGGTTATATGGCCCGTTCAGCTAAACCCCGTAAACGAAAACCTTCCCCACAAAGAAGCAAACTTCCCCGCTATGTTGTGAAACTTCATCCGGATGATTTTTTTGACGAAGAAGACGCTGAAGTTCTGCGCTTTGATAATTTTGACGATGCCGTTGAGTGCTGCGCTGACCTGGGTATTCCGTTCTTTCTGGATGCAGGAAACAAAAAGCTGGTCTTCTGGTTTGTTCGTGTCGATGACGAAGGGTATCCGGAAATAGCCCGCTGTACGGAGCGGGAGTTTGCAACCATTCTTGCCGGTATCAGTGCCGGTGGTATGTACTGCCCGGAATGCGGCACAGTTCACTGGCCGGATGGCGTTACCCCACCCGTCTGATGCTTCCCCGTTTTGCCGATATTTTTCAGCAGGGTAACCGCTGGCTTAACTGGCTGGAGAAACAGCCGGAAGGTTCAGTGCGTCCGGTGGTGACTGAGTCAGTGACAAAAATCATGGCATGCGGGACCACGCTGATGGGCTACACGCAATGGTACTGTTCATCACCGGACTGTTGCCACACAAAAAAGGTCTGCTTCCGGTGTAAAAGTCGCTCCTGCCCGCACTGCGGGGTGAAGGCTGGCGCACAGTGGATACAGTATCTGCTGAGTCTGGTCCCCGACTGCCCGTGGCAGCATATTGTGTTCACACTTCCCTGCCAGTACTGGCCCCTGATATTCCACAACAGGTGGTTACTGGCAGAGATGAGCCGCATTGCTGCGAATGTGATACTGGAAATCTGCCGTCAGGCGGACGTGGAGCCGGGGATATTCACGGTAATCCACACATGGGGGCGTGACCAGCAGTGGCATCCGCATATCCATTTATCGACAACTGCCGGTGGTGTGACGTCGGGCCACACCTGGAAAAATATTCATTTTTACGCCCGTAAGGTGATGAGCATGTGGCGTTACCGGATAACGCGGCTACTGTCCCGGAAATACCCGGAGCTGGTAATACCGGATGAACTGGCAGTGGAAGGAAACAGCAAACGGGACTGGAATCGCTTTCTGGACACGCATTACCGCCGCGGCTGGAATGTCAACATATCCAGGGTGATGGATAACGCCACACATGTGGCGGTGTACTTCGGCTCTTACCTGAAAAAGCCACCGGTGCCGATGAGTCGGCTGGAGCATTATGCCGGTCAGGATGAAATCGGTCTGCGTTACAACAGTCACCGTACAAAACGGGAAGAATACCTGTTGATGAGTGGAGATGAGTTCATGGAAAGGTTCTCCTGGCATGTAGCAGATAAGGGGTTCCGTATGGTGAGGTACTACGGTTTCCTGAGTCCGGTGAAGCGCCGCTTACTGGAAGAGGTGGTGTACGTCATAACGGAGACGGTGAGAAAAACGGCGATGCAAATCAGGTGGAGAGGGATGTATCAGCGGTTACTGAAGGTTGACCCGCTGAAATGCGTCCTGTACGGAAGTCAGATGCGGTTTACGGGGCTGAAGCGGGGATACCGTCTGGCAGAGCTGGTTATGATGCATGAGCCGCTGGCCCGAATGCAGTGTTGCGGCTGAGAGCCGCAGAGGGGAAGTTGCGTCCATTTTACGGGGAACGGAGCAAAAAACCACCATTCATACCCTTTATCAATCAGTGTTATCCTGTTTAATAGTCGTTTCCGTTCATATGGTGCATAAGGAGTGTTGAAGAAATATCCGTTTTGTGGTGTTTTTTAATCTTTTGGGGGTTTTAATTCCTATTGAT

>IS91-V131

CGAGTAGGCAGCCTGGCGGCTGCGGCTTGTCATGGTCTGAGATTACCGTTATAAAAACAGGCAATATCATTGTCTTTCAGGTGGTTATATGGCCCGTTCAGCTAAACCCCGTAAACGAAAACCTTCCCCACAAAGAAGCAAACTTCCCCGCTATGTTGTGAAACTTCATCCGGATGATTTTTTTGACGAAGAAGACGCTGAAGTTCTGCGCTTTGATAATTTTGACGATGCCGTTGAGTGCTGCGCTGACCTGGGTATTCCGTTCTTTCTGGATGCAGGAAACAAAAAGCTGGTCTTCTGGTTTGTTCGTGTCGATGACGAAGGGTATCCGGAAATAGCCCGCTGTACGGAGCGGGAGTTTGCAACCATTCTTGCCGGTATCAGTGCCGGTGGTATGTACTGCCCGGAATGCGGCACAGTTCACTGGCCGGATGGCGTTACCCCACCCGTCTGATGCTTCCCCGTTTTGCCGATATTTTTCAGCAGGGTAACCGCTGGCTTAACTGGCTGGAGAAACAGCCGGAAGGTTCAGTGCGTCCGGTGGTGACTGAGTCAGTGACAAAAATCATGGCATGCGGGACCACGCTGATGGGCTACACGCAATGGTACTGTTCATCACCGGACTGTTGCCACACAAAAAAGGTCTGCTTCCGGTGTAAAAGTCGCTCCTGCCCGCACTGCGGGGTGAAGGCTGGCGCACAGTGGATACAGTATCTGCTGAGTCTGGTCCCCGACTGCCCGTGGCAGCATATTGTGTTCACACTTCCCTGCCAGTACTGGCCCCTGATATTCCACAACAGGTGGTTACTGGCAGAGATGAGCCGCATTGCTGCGAATGTGATACTGGAAATCTGCCGTCAGGCGGACGTGGAGCCGGGGATATTCACGGTAATCCACACATGGGGGCGTGACCAGCAGTGGCATCCGCATATCCATTTATCGACAACTGCCGGTGGTGTGACGTCGGGCCACACCTGGAAAAATCTTCATTTTTACGCCCGTAAGGTGATGAGTATGTGGCGTTACCGGATAACGCGGCTACTGTCCCGGAAATACCCGGAGCTGGTAATACCGGATGAACTGGCAGTGGAAGGAAACAGCAAACGGGACTGGAATCGCTTTCTGGACACGCATTACCGCCGCGGCTGGAATGTCAACATATCCAGGGTGATGGATAACGCCACACATGTGGCGGTGTACTTCGGCTCTTACCTGAAAAAGCCACCGGTGCCGATGAGTCGGCTGGAGCATTATGCCGGTCAGGATGAAATCGGTCTGCGTTACAACAGTCACCGTACAAAACGGGAAGAATACCTGTTGATGAGTGGAGATGAGTTCATGGAAAGGTTCTCCTGGCATGTAGCAGATAAGGGGTTCCGTATGGTGAGGTACTACGGTTTCCTGAGTCCGGTGAAGCGCCGCTTACTGGAAGAGGTGGTGTACGTCATAACGGAGACGGTGAGAAAGACGGCGATGCAAATCAGGTGGAGAGGGATGTATCAGCGGTTACTGAAGGTTGACCCGCTGAAATGCGTCCTGTACGGAAGTCAGATGCGGTTTACGGGGCTGAAGCGGGGATACCGTCTGGCAGAGCTGGTTATGATGCATGAGCCGCTGGCCCGAATGCAGTGTTGCGGCTGAGAGCCGCAGAGGGGAAGTTGCGTCCATTTTACGGGGAACGGAGCAAAAAACCACCATTCATACCCTTTATCAATCAGTGTTATCCTGTTTAATAGTCGTTTCCGTTCATATGGTGCATAAGGAGTGTTGAAGAAATATCCGTTTTGTGGTGTTTTTTAATCTTTTGGGGGTTTTAATTCCTATTGAT

>IS91-V132

CGAGTAGGCAGCCTGGCGGCTGCGGCTTGTCATGGTCTGAGATTACCGTTATAAAAACAGGCAATATCATTGTCTTTCAGGTGGTTATATGGCCCGTTCAGCTAAACCCCGTAAACGAAAACCTTCCCCACAAAGAAGCAAACTTCCCCGCTATGTTGTGAAACTTCATCCGGATGATTTTTTTGACGAAGAAGACGCTGAAGTTCTGCGCTTTGATAATTTTGACGATGCCGTTGAGTGCTGCGCTGACCTGGGTATTCCGTTCTTTCTGGATGCAGGAAACAAAAAGCTGGTCTTCTGGTTTGTTCGTGTCGATGACGAAGGGTATCCGGAAATAGCCCGCTGTACGGAGCGGGAGTTTGCAACCATTCTTGCCGGTATCAGTGCCGGTGGTATGTACTGCCCGGAATGCGGCACAGTTCACTGGCCGGATGGCGTTACCCCACCCGTCTGATGCTTCCCCGTTTTGCCGATATTTTTCAGCAGGGTAACCGCTGGCTTAACTGGCTGGAGAAACAGCCGGAAGGTTCAATGCGTCCGGTGGTGACTGAGTCAGTGACAAAAATCATGGCATGCGGGACCACGCTGATGGGCTACACGCAATGGTACTGTTCATCACCGGACTGTTGCCACACAAAAAAGGTCTGCTTCCGGTGTAAAAGTCGCTCCTGCCCGCACTGCGGGGTGAAGGCTGGCGCACAGTGGATACAGTATCTGCTGAGTCTGGTCCCCGACTGCCCGTGGCAGCATATTGTGTTCACACTTCCCTGCCAGTACTGGCCCCTGATATTCCACAACAGGTGGTTACTGGCAGAGATGAGCCGCATTGCTGCGAATGTGATACTGGAAATCTGCCGTCAGGCGGACGTGGAGCCGGGGATATTCACGGTAATCCACACATGGGGGCGTGACCAGCAGTGGCATCCGCATATCCATTTATCGACAACTGCCGGTGGTGTGACGTCGGGCCACACCTGGAAAAATATTCATTTTTACGCCCGTAAGGTGATGAGCATGTGGCGTTACCGGATAACGCGGCTACTGTCCCGGAAATACCCGGAGCTGGTAATACCGGATGAACTGGCAGTGGAAGGAAACAGCAAACGGGACTGGAATCGCTTTCTGGACACGCATTACCGCCGCGGCTGGAATGTCAACATATCCAGGGTGATGGATAACGCCACACATGTGGCGGTGTACTTCGGCTCTTACCTGAAAAAGCCACCGGTGCCGATGAGTCGGCTGGAGCATTATGCCGGTCAGGATGAAATCGGTCTGCGTTACAACAGTCACCGTACAAAACGGGAAGAATACCTGTTGATGAGTGGAGATGAGTTCATGGAAAGGTTCTCCTGGCATGTAGCAGATAAGGGGTTCCGTATGGTGAGGTACTACGGTTTCCTGAGTCCGGTGAAGCGCCGCTTACTGGAAGAGGTGGTGTACGTCATAACGGAGACGGTGAGAAAAACGGCGATGCAAATCAGGTGGAGAGGGATGTATCAGCGGTTACTGAAGGTTGACCCGCTGAAATGCGTCCTGTACGGAAGTCAGATGCGGTTTACGGGGCTGAAGCGGGGATACCGTCTGGCAGAGCTGGTTATGATGCATGAGCCGCTGGCCCGAATGCAGTGTTGCGGCTGAGAGCCGCAGAGGGGAAGTTGCGTCCATTTTACGGGGAACGGAGCAAAAAACCACCATTCATACCCTTTATCAATCAGTGTTATCCTGTTTAATAGTCGTTTCCGTTCATATGGTGCATAAGGAGTGTTGAAGAAATATCCGTTTTGTGGTGTTTTTTAATCTTTTGGGGGTTTTAATTCCTATTGAT

>IS91-V133

CGAGTAGGCAGCCTGGCGGCTGCGGCTTGTCATGGTCTGAGATTACCGTTATAAAAACAGGCAATATCATTGTCTTTCAGGTGGTTATATGGCCCGTTCAGCTAAACCCCGTAAACGAAAACCTTCCCCACAAAGAAGCAAACTTCCCCGCTATGTTGTGAAACTTCATCCGGATGATTTTTTTGACGAAGAAGACGCTGAAGTTCTGCGCTTTGATAATTTTGACGATGCCGTTGAGTGCTGCGCTGACCTGGGTATTCCGTTCTTTCTGGATGCAGGAAACAAAAAGCTGGTCTTCTGGTTTGTTCGTGTCGATGACGAAGGGTATCCGGAAATAGCCCGCTGTACGGAGCGGGAGTTTGCAACCATTCTTGCCGGTATCAGTGCCGGTGGTATGTACTGCCCGGAATGCGGCACAGTTCACTGGCCGGATGGCGTTACCCCACCCGTCTGATGCTTCCCCGTTTTGCCGATATTTTTCAGCAGGGTAACCGCTGGCTTAACTGGCTGGAGAAACAGCCGGAAGGTTCAGTGCGTCCGGTGGTGACTGAGTCAGTGACAAAAATCATGGCATGCGGGACCACGCTGATGGGCTACACGCAATGGTACTGTTCATCACCGGACTGTTGCCACACAAAAAAGGTCTGCTTCCGGTGTAAAAGTCGCTCCTGCCCGCACTGCGGGGTGAAGGCTGGCGCACAGTGGATACAGTATCTGCTGAGTCTGGTCCCCGACTGCCCGTGGCAGCATATTGTGTTCACACTTCCCTGCCAGTACTGGCCCCTGATATTCCACAACAGGTGGTTACTGGCAGAGATGAGCCGCATTGCTGCGAATGTGATACTGGAAATCTGCCGTCAGGCGGACGTGGAGCCGGGGATATTCACGGTAATCCACACATGGGGGCGTGACCAGCAGTGGCATCCGCATATCCATTTATCGACAACTGCCGGTGGTGTGACGTCGGGCCACACCTGGAAAAATATTCATTTTTACGCCCGTAAGGTGATGAGCATGTGGCGTTACCGGATAACGCGGCTACTGTCCCGGAAATACCCGGAGCTGGTAATACCGGATGAACTGGCAGTGGAAGGAAACAGCAAACGGGACTGGAATCGCTTTCTGGACACGCATTACCGCCGCGGCTGGAATGTCAACATATCCAGGGTGATGGATAACGCCACACATGTGGCGGTGTACTTCGGCTCTTACCTGAAAAAGCCACCGGTGCCGATGAGTCGGCTGGAGCATTATGCCGGTCAGGATGAAATCGGTCTGCGTTACAACAGTCACCGTACAAAACGGGAAGAATACCTGTTGATGAGTGGAGATGAGTTCATGGAAAGGTTCTCCTGGCATGTAGCAGATAAGGGGTTCCGTATGGTGAGGTACTACGGTTTCCTGAGTCCGGTGAAGCGCCGCTTACTGGAAGAGGTGGTGTACGTCATAACGGAGACGGTGAGAAAAACGGCGATGCAAATCAGGTGGAGAGGGATGTATCAGCGGTTACTGAAGGTTGACCCGCTGAAATGCGTCCTGTACGGAAGTCAGATGCGGTTTACGGGGCTGAAGCGGGGATACCGTCTGGCAGAGCTGGTTATAATGCATGAGCCGCTGGCCCGAATGCAGTGTTGCGGCTGAGAGCCGCAGAGGGGAAGTTGCGTCCATTTTACGGGGAACGGAGCAAAAAACCACCATTCATACCCTTTATCAATCAGTGTTATCCTGTTTAATAGTCGTTTCCGTTCATATGGTGCATAAGGAGTGTTGAAGAAATATCCGTTTTGTGGTGTTTTTTAATCTTTTGGGGGTTTTAATTCCTATTGAT

>IS91-V134

CGAGTAGGCAGCCTGGCGGCTGCGGCTTGTCATGGTCTGAGATTACCGTTATAAAAACAGGCAATATCATTGTCTTTCAGGTGGTTATATGGCCCGTTCAGCTAAACCCCGTAAACGAAAACCTTCCCCACAAAGAAGCAAACTTCCCCGCTATGTTGTGAAACTTCATCCGGATGATTTTTTTGACGAAGAAGACGCTGAAGTTCTGCGCTTTGATAATTTTGACGATGCCGTTGAGTGCTGCGCTGACCTGGGTATTCCGTTCTTTCTGGATGCAGGAAACAAAAAGCTGGTCTTCTGGTTTGTTCGTGTCGATGACGAAGGGTATCCGGAAATAGCCCGCTGTACGGAGCGGGAGTTTGCAACCATTCTTGCCGGTATCAGTGCCGGTGGTATGTACTGCCCGGAATGCGGCACAGTTCACTGGCCGGATGGCGTTACCCCACCCGTCTGATGCTTCCCCGTTTTGCCGATATTTTTCAGCAGGGTAACCGCTGGCTTAACTGGCTGGAGAAACAGCCGGAAGGTTCAGTGCGTCCGGTGGTGACTGAGTCAGTGACAAAAATCATGGCATGCGGGACCACGCTGATGGGCTACACGCAATGGTACTGTTCATCACCGGACTGTTGCCACACAAAAAAGGTCTGCTTCCGGTGTAAAAGTCGCTCCTGCCCGCACTGCGGGGTGAAGGCTGGCGCACAGTGGATACAGTATCTGCTGAGTCTGGTCCCCGACTGCCCGTGGCAGCATATTGTGTTCACACTTCCCTGCCAGTACTGGCCCCTGATATTCCACAACAGGTGGTTACTGGCAGAGATGAGCCGCATTGCTGCGAATGTGATACTGGAAATCTGCCGTCAGGCGGACGTGGAGCCGGGGATATTCACGGTAATCCACACATGGGGGCGTGACCAGCAGTGGCATCCGCATATCCATTTATCGACAACTGCCGGTGGTGTGACGTCGGGCCACACCTGGAAAAATCTTCATTTTTACGCCCGTAAGGTGATGAGCATGTGGCGTTACCGGATAACGCGGCTACTGTCCCGGAAATACCCGGAGCTGGTAATACCGGATGAACTGGCAGTGGAAGGAAACAGCAAACGGGACTGGAATCGCTTTCTGGACACGCATTACCGCCGCGGCTGGAATGTCAACATATCCAGGGTGATGGATAACGCCACACATGTGGCGGTGTACTTCGGCTCTTACCTGAAAAAGCCACCGGTGCCGATGAGTCGGCTGGAGCATTATGCCGGTCAGGATGAAATCGGTCTGCGTTACAACAGTCACCGTACAAAACGGGAAGAATACCTGTTGATGAGTGGAGATGAGTTCATGGAAAGGTTCTCCTGGCATGTAGCAGATAAGGGGTTCCGTATGGTGAGGTACTACGGTTTCCTGAGTCCGGTGAAGCGCCGCTTACTGGAAGAGGTGGTGTACGTCATAACGGAGACGGTGAGAAAAACGGCGATGCAAATCAGGTGGAGAGGGATGTATCAGCGGTTACTGAAGGTTAACCCGCTGAAATGCGTCCTGTACGGAAGTCAGATGCGGTTTACGGGGCTGAAGCGGGGATACCGTCTGGCAGAGCTGGTTATGATGCATGAGCCGCTGGCCCGAATGCAGTGTTGCGGCTGAGAGCCGCAGAGGGGAAGTTGCGTCCATTTTACGGGGAACGGAGCAAAAAACCACCACTCATACCCTTTATCAATCAGTGTTATCATGTTTAATAGTCGTTTCCGTTCATATGGTGCATAAGGAGTGTTGAAGAAATATCCGTTTTGTGGTGTTTTTTAATCTTTTGGGGGTTTTAATTCCTATTGAT
